# Phylogenomics and metabolic engineering reveal a conserved gene cluster in Solanaceae plants for withanolide biosynthesis

**DOI:** 10.1101/2024.09.27.614867

**Authors:** Samuel Edward Hakim, Nancy Choudhary, Karan Malhotra, Jian Peng, Ahmed Arafa, Arne Bültemeier, Ronja Friedhoff, Maximilian Bauer, Claus-Peter Witte, Marco Herde, Philipp Heretsch, Boas Pucker, Jakob Franke

## Abstract

Withanolides are steroidal lactones from nightshade (Solanaceae) plants. Of the over 1,200 known representatives, many possess potent biological activities, but their drug potential has not been fully realised up until now. A central obstacle is the limited availability of minor withanolides, caused by a lack of knowledge about the underlying biosynthetic pathways. Here, we combine phylogenomics with metabolic engineering to overcome this limitation. By sequencing the genome of the medicinal plant and archetypical withanolide producer ashwagandha (*Withania somnifera*) and comparing the genome sequences of nine Solanaceae species, we discovered a conserved gene cluster for withanolide biosynthesis, consisting of two sub-gene clusters which differ in their expression patterns. To investigate the functions of the encoded enzymes, we established metabolic engineering platforms in yeast (*Saccharomyces cerevisiae*) and the model plant *Nicotiana benthamiana*. This allowed us to reconstitute the first three oxidative steps of withanolide biosynthesis, catalysed by the cytochrome P450 monooxygenases CYP87G1, CYP88C7, and CYP749B2, leading to the aglycone of the known compound withanoside V. Our work sets the basis for the biotechnological production of withanolides in heterologous hosts and will therefore help to fully harness the drug potential of these plant steroids in the future.

## Introduction

Plants are well-known for their extensive capabilities to produce structurally complex metabolites with potent biological activities. Even though many of these specialised metabolites have been studied extensively at a chemical level, the genetic basis for their biosynthesis is often still unknown, representing a major hurdle for biotechnological improvement of medicinal plants and development of microbial production systems alike. Such a lack of knowledge at the gene level also exists for withanolide biosynthesis. Withanolides are steroidal lactones occurring in several members of the nightshade family (Solanaceae). Named after the first discovery of these compounds in the medicinal plant *Withania somnifera* (ashwagandha), approx. 1,200 withanolides are nowadays known from 22 genera of Solanaceae^1,2^. These metabolites have played a role in traditional medicine since ancient times^3,4^. For example, *W*. *somnifera* has been used in traditional Indian medicine, Ayurveda, as an anti-stress agent for millennia; these stress-relieving properties are also supported by modern placebo-controlled studies^5–7^. Detailed pharmacological studies underline the broad spectrum of biological activities of withanolides^4,8–12^ and withanolide-inspired drug candidates^13^. In stark contrast, only a single enzyme specific to withanolide biosynthesis has been identified and characterised so far^14^, hampering efforts to engineer withanolide metabolism *in planta* and produce withanolides biotechnologically^15^. This enzyme, sterol Δ^24^-isomerase (24ISO)^14^ catalyses the isomerisation of 24-methylenecholesterol (**1**), an intermediate of the general phytosterol pathway, to the withanolide-specific key intermediate 24-methyldesmosterol (**2**) (Fig. 1). As such, the 24ISO reaction represents the key committed step at which withanolide biosynthesis branches from phytosterol and brassinosteroid biosynthesis^14^. Several genes and enzymes in the general phytosterol pathway upstream of 24ISO were investigated in withanolide-producing plants^16–19^. In contrast, the biosynthetic pathway downstream of 24ISO, responsible for the conversion of 24-methyldesmosterol (**2**) to withanolides, is not yet known. While several putative withanolide biosynthesis gene candidates were tested by virus-induced gene silencing^20,21^, no clear biochemical activity has been reported for them so far.

**Fig. 1.**
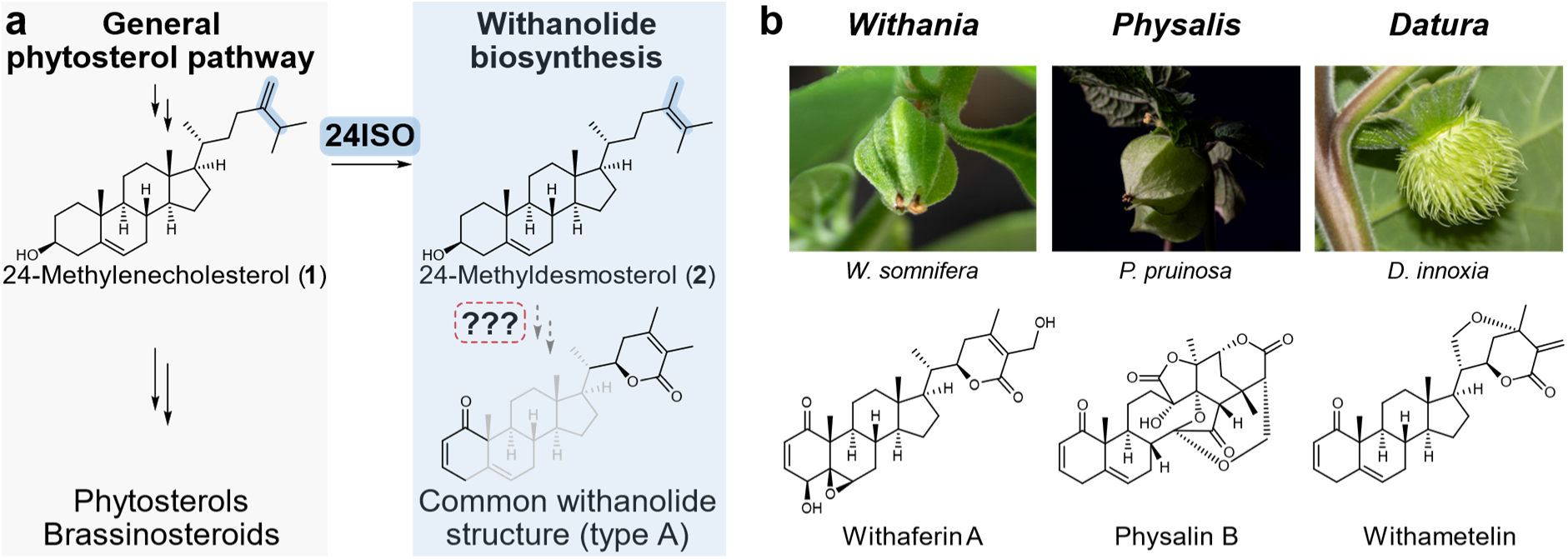
Withanolides are steroidal lactones from Solanaceae plants derived from the general phytosterol pathway intermediate 24-methylenecholesterol (1). **a** 24-Methylenecholesterol (1) is converted by the only known withanolide biosynthetic enzyme sterol Δ^24^-isomerase (24ISO)^14^ into 24-methyldesmosterol **(2).** The enzymes for all subsequent biosynthetic steps are not known. b Representative genera and species of withanolide-producing plants and characteristic compounds from them. Photographs by Jakob Maximilian Horz, TU Braunschweig, Germany.

While the elucidation of plant biosynthetic pathways has been a slow and tedious process for a long time, advances in sequencing techniques over the past 20 years have dramatically accelerated the pace of gene function discovery in plants^22^. Most commonly, transcriptomic data is used to identify biosynthetic genes based on specific expression patterns in different growth stages, tissues or even cell types^23–26^. Genomic data was traditionally considered to be less important for pathway elucidation, because – in contrast to microorganisms^27^ – many biosynthetic genes in plants discovered early were not physically clustered^28–30^. However, this perspective is getting continuously challenged, as more plant genome sequences become available^31^. Indeed, many examples now show that biosynthetic gene clusters in plants are relatively common and enable efficient pathway elucidation^30^, as demonstrated recently for saponin, alkaloid, and terpenoid biosynthesis^32–35^.

During the discovery of the only withanolide-specific pathway gene, *24ISO*, Knoch *et al*. reported possible clustering of this gene with other genes common for specialised metabolism^14^, but their analysis was still strongly limited by the lack of high-quality genome sequences of Solanaceae plants in 2018. A comparison of *Capsicum annuum*, *Solanum melongena,* and *Petunia inflata* indicated the co-occurrence of *24ISO* with genes encoding cytochrome P450 monooxygenases and oxoglutarate-dependent dioxygenases^14^. However, none of the species of this comparison is known as a producer of canonical withanolides. In recent years, several high-quality genome sequences of withanolide-producing plants have been released. These include for example *Physalis floridana* (*Physalis pubescens*)^36^, *Physalis grisea*^37^, *Physalis pruinosa*^37^, *Datura stramonium*^34,38^, and *Datura wrightii*^39^. In this work, we now revisit the previous hypothesis of possible gene clustering in the context of withanolide biosynthesis. By sequencing the genome of the archetypical withanolide producer *Withania somnifera* and synteny analyses with other Solanaceae genome sequences, we reveal a conserved gene cluster in withanolide-producing plants that harbours the withanolide pathway gene *24ISO* and multiple other genes typical for specialised metabolism. To overcome previous obstacles in functional validation of withanolide pathway genes, we employed metabolic engineering in the model organisms yeast (*Saccharomyces cerevisiae*) and *Nicotiana benthamiana* to successfully establish two independent platforms for withanolide pathway reconstitution. This enabled the characterisation of three cytochrome P450 monooxygenases that oxidise 24-methyldesmosterol (**2**) to construct the pivotal δ-lactone ring of withanolides. Our discovery of a conserved gene cluster for withanolide biosynthesis in Solanaceae plants and the development of synthetic biology systems will enable full elucidation and engineering of withanolide biosynthesis in the future, to further harness the drug potential of withanolides.

## Results

### Genome assembly of *Withania somnifera*

To explore the genomic context of withanolide biosynthesis, we generated a genome assembly of *W. somnifera*, known as a prolific producer of withanolides. Building on Oxford Nanopore Sequencing^40^, we generated an assembly comprising 102 contigs with an N50 length of 71 Mb and a total assembly size of 2.8 Gbp (Supplementary Table 1). A total of 35,118 protein-encoding genes were predicted based on homology and transcriptome data, with an average gene length of 4,978 bp and an average coding sequence (CDS) length of 1,266 bp. BUSCO analysis^41^ using Solanales_odb10 dataset revealed 96.6% complete homologs in the predicted proteins of *W. somnifera*. Additionally, we re-annotated the chromosome-scale genome sequences of *P. grisea* and *P. pruinosa,* resulting in improved gene models and higher BUSCO completeness compared to the previous annotations^37^. The predicted protein-coding genes of both genome sequences achieved 97.5% completeness in the BUSCO analysis.

The 12 pseudochromosomes from the chromosome-scale assembly of *P. pruinosa*^37^ were well represented by the top 36 largest contigs in our *W. somnifera* assembly (accounting for ∼87% of the total assembly). A Circos plot revealed large overall synteny between both species (Fig. 2). Although the relative amount of repeats is comparable in *P. pruinosa* (77.59%) and *W. somnifera* (76.02%), *W. somnifera* has more terminal inverted repeats (TIR) (13.13% vs 1.7%).

**Fig. 2.**
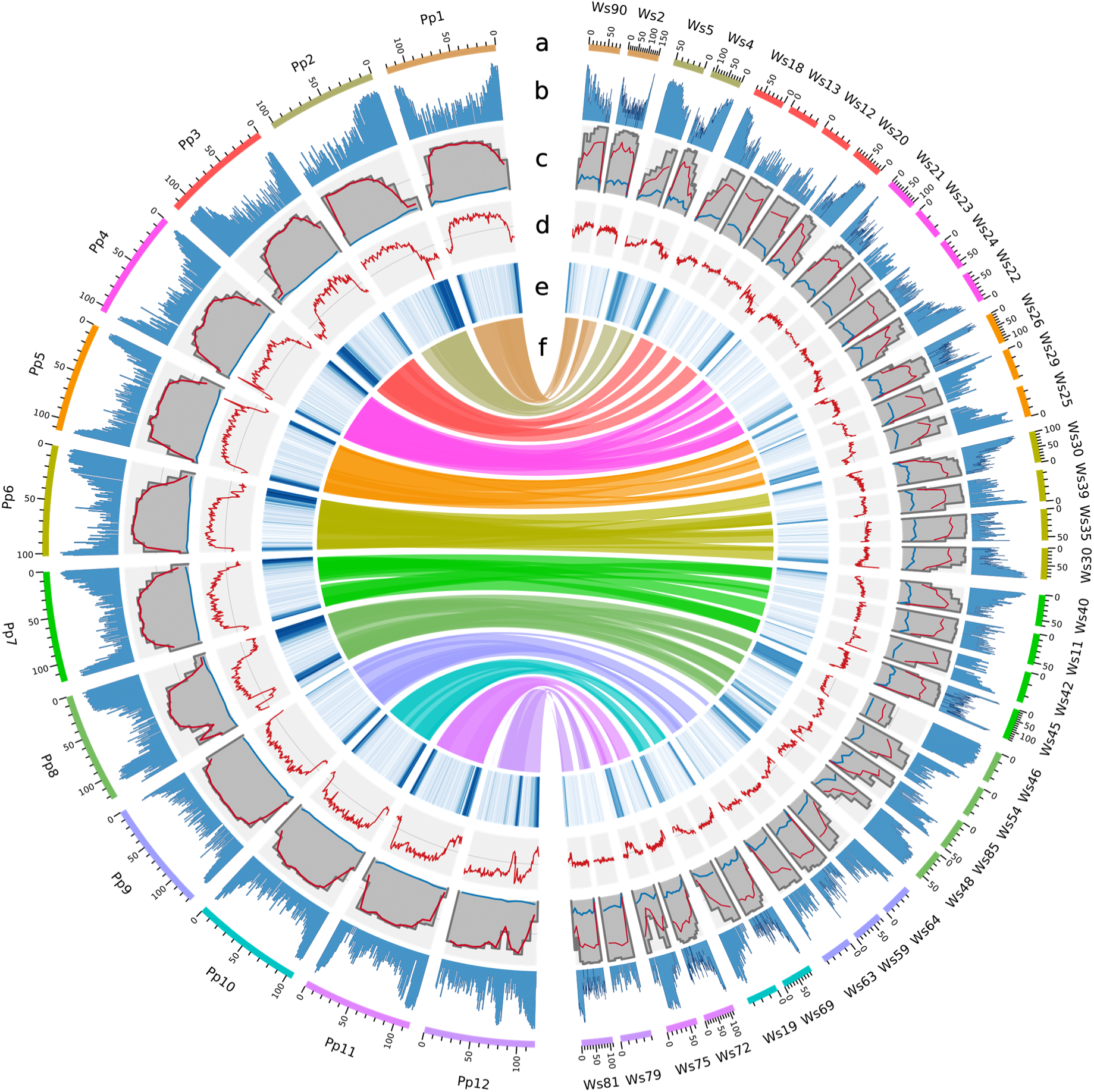
Circos plot comparing important genomic features of the newly assembled genome sequence of *Withania somnifera* (*Ws*, right) with that of *Physalis pruinosa* (*Pp*, left). **a** Genomic landscape of the 36 *W. somnifera* pseudochromosomes (right) and 12 *P. pruinosa* pseudochromosomes (left). All density information is calculated in non-overlapping 1-Mbp windows. **b** Tandem repeats density. **c** Percentage of transposable elements (TEs), calculated in 10-Mbp non-overlapping windows. Total TEs in grey, long-terminal repeat (LTR) retrotransposons in red and terminal inverted repeats (TIR) in blue. **d** GC content. **e** Distribution of protein-coding genes. **f** Links between syntenic regions of both genomes.

### Phylogenomics discovery of a putative withanolide biosynthetic gene cluster

With our high-quality assembly of the *W. somnifera* genome sequence in hands, we next set out to systematically explore the genomic context surrounding *24ISO*, the only previously reported withanolide biosynthesis gene. To search for a possible conserved gene cluster, we analysed highly continuous genome sequences of the following withanolide-producing plants: *Physalis floridana*^36^, *Physalis grisea*^37^, *Physalis pruinosa*^37^, *Datura stramonium*^34,38^, and *Datura wrightii*^39^. For comparison, we included genome sequences of *Solanum lycopersicum*^42^, *Solanum tuberosum*^43^, and *Nicotiana tabacum*^44^, which are phylogenetically related, but not known as withanolide producers. Using the experimentally characterised *24ISO* genes^14^ as a bait, we identified their genomic positions and further orthologous genes as a starting point for synteny comparison. All genome sequences of withanolide-producing plants analysed here contained two copies of *24ISO* in close proximity. In *W. somnifera*, a third *24ISO* copy was additionally found on a different contig. No *24ISO* orthologue was found in *S. lycopersicum*, *S. tuberosum*, and *N. tabacum*. Then, we compared the synteny of the genomic regions surrounding these *24ISO* orthologues (Fig. 3, Supplementary Fig. 1). Genomes of withanolide producers contained a syntenic region that was absent in the non-producers *S. lycopersicum* and *S. tuberosum*. We then deduced the classes of encoded enzymes in this syntenic region by their Pfam domains. Strikingly, all conserved genes belong to gene families common in plant specialised metabolism, most importantly cytochrome P450 monooxygenases (CYPs), 2-oxoglutarate-dependent dioxygenases (ODDs), short-chain dehydrogenases/reductases (SDRs), and acyltransferases (AT); less expected was the occurrence of sulfotransferase (ST) genes. While sulfotransferases are well-known in the biosynthesis of certain specialised metabolites such as glucosinolates^45^, a possible link to withanolide biosynthesis is not yet known. A few withanolides bearing 3-*O*-sulphate groups are known, however^46,47^.

**Fig. 3.**
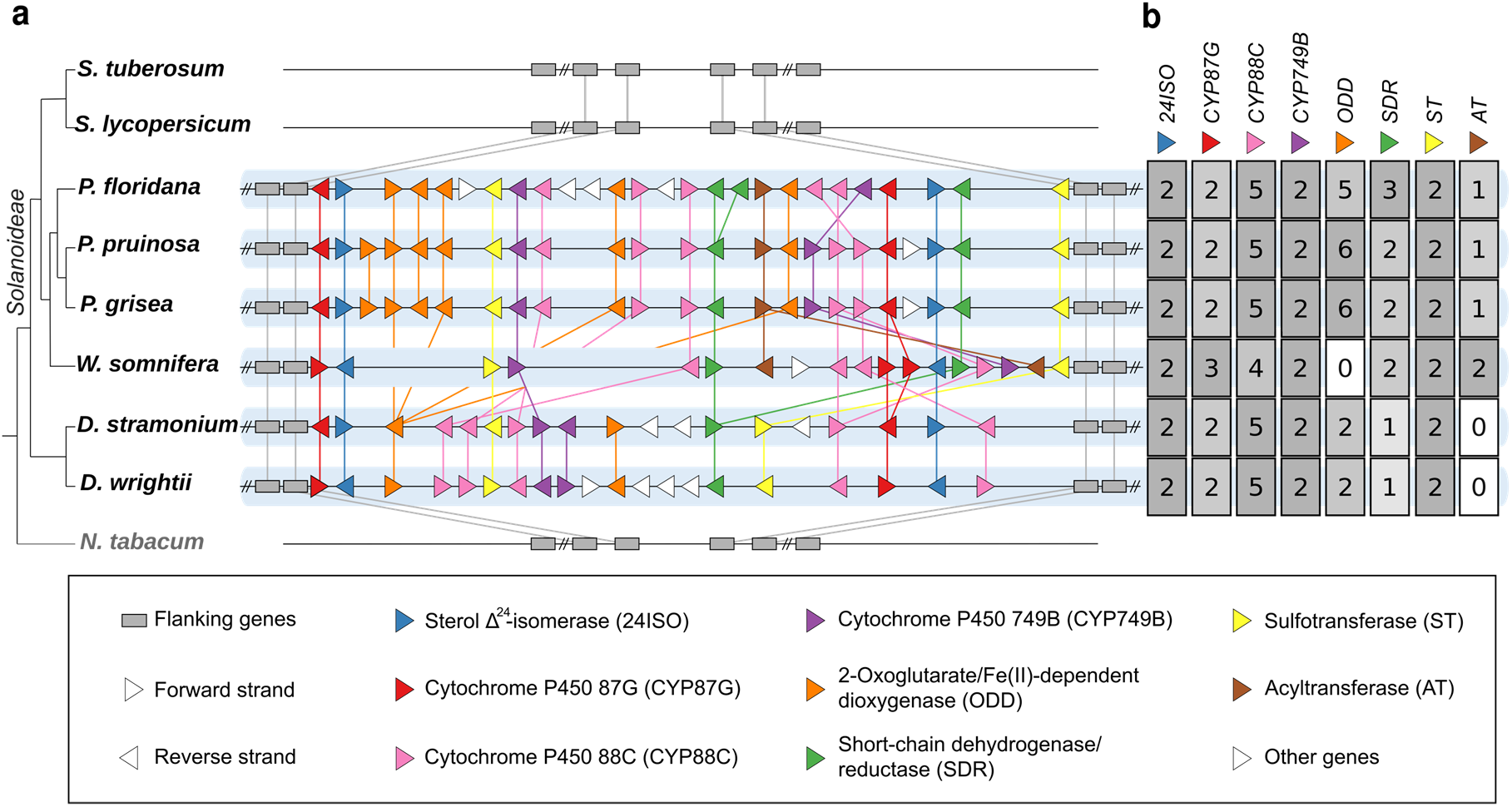
Syntenic biosynthetic gene clusters containing *24ISO* in withanolide-producing Solanaceae plants. **a** Synteny plot. Gene lengths are unified for clarity. *N. tabacum* is included as an outgroup of the non-withanolide producing subfamily Solanoideae. **b** Heatmap summary of gene copy numbers of each gene family. Background colour is normalised to the highest number per column. Expression information about genes in the cluster are displayed in Supplementary Fig. 2-4.

Cytochrome P450 monooxygenases (CYPs) are key players in plant specialised metabolism and take a central role in triterpenoid and steroid biosynthetic pathways^26,48–53^. Past studies demonstrated that several CYP families, e.g., CYP716, are particularly likely to be involved in triterpenoid biosynthesis, whereas many other CYP families contain no known representative acting on such substrates^48,54^. The CYPs in the putative withanolide biosynthetic gene cluster fall into three different CYP families, namely CYP87, CYP88 (both part of the CYP85 clan) and CYP749 (part of the CYP72 clan). Both clans, particularly CYP85, are well-known as hotspots of CYPs involved in triterpenoid and steroid metabolism^48^. Two known members of the CYP88 family act on diterpenoids, namely CYP88A3 and CYP88A4 from gibberellin biosynthesis^55^, but all four other CYP88s participate in pentacyclic triterpenoid^56^ and cucurbitacin^57,58^ biosynthesis. Likewise, all members of the CYP87^57,59^ and CYP749^60^ families act on steroid and triterpenoid substrates. A CYP749 from *W. somnifera* was previously identified by a transcriptome-based approach and associated with withanolide biosynthesis based on virus-induced gene silencing, which resulted in a strong change in the withanolide profile^20^. Employing a similar approach, a recent study in *P. angulata* implied a role of a CYP749 and a CYP88 in the biosynthesis of physalin-type withanolides^61^. The biochemical functions of these CYPs as well as the genomic locations of the underlying genes have been unknown so far. We performed a BLAST search of these previously suggested candidates against our assembly of *W. somnifera*. Indeed, the closest homologues of these CYPs are encoded in the gene clusters described here (Supplementary Table 2).

Next, we analysed published RNA-seq datasets to verify that the genes in the gene cluster are not only physically clustered, but also co-expressed. Surprisingly, we observed two clearly separated groups of expression patterns, both in *W. somnifera* and in *D. stramonium* (Supplementary Fig. 2-4). This suggests that the withanolide gene cluster is separated into two sub-gene clusters, which are differentially regulated. The sub-gene cluster separation was also examined by phylogenetic analyses (Supplementary Fig. 5-10).

Taken together, our phylogenomics analysis in combination with gene silencing studies by others^20,61^ strongly suggest that withanolide biosynthesis involves a conserved gene cluster.

### Development of a yeast platform for production of withanolide pathway intermediates

Next, we wanted to support our discovery of a putative withanolide biosynthetic gene cluster by elucidating the biochemical functions of pathway enzymes. Reconstitution of withanolide biosynthesis in heterologous hosts has remained an unsolved problem up until now. The extremely low polarity and the limited accessibility of the last known intermediate 24-methyldesmosterol (**2**) prevents an efficient use as a substrate for enzyme assays *in vitro*. An alternative would be to produce 24-methyldesmosterol (**2**) *in vivo*, but no efficient metabolic engineering strategy has been reported so far. Previous studies showed that its precursor, 24-methylenecholesterol (**1**), can be produced in yeast (*Saccharomyces cerevisiae*) by deleting the ergosterol biosynthesis genes *ERG4* and *ERG5* and adding a gene encoding a Δ7 reductase from plants or animals, in order to hijack the sterol metabolism of yeast to produce plant-like sterols^62–64^. We decided to utilise and expand this strategy to set up a platform for functional evaluation of withanolide biosynthesis genes in yeast (Fig. 4a).

**Fig. 4.**
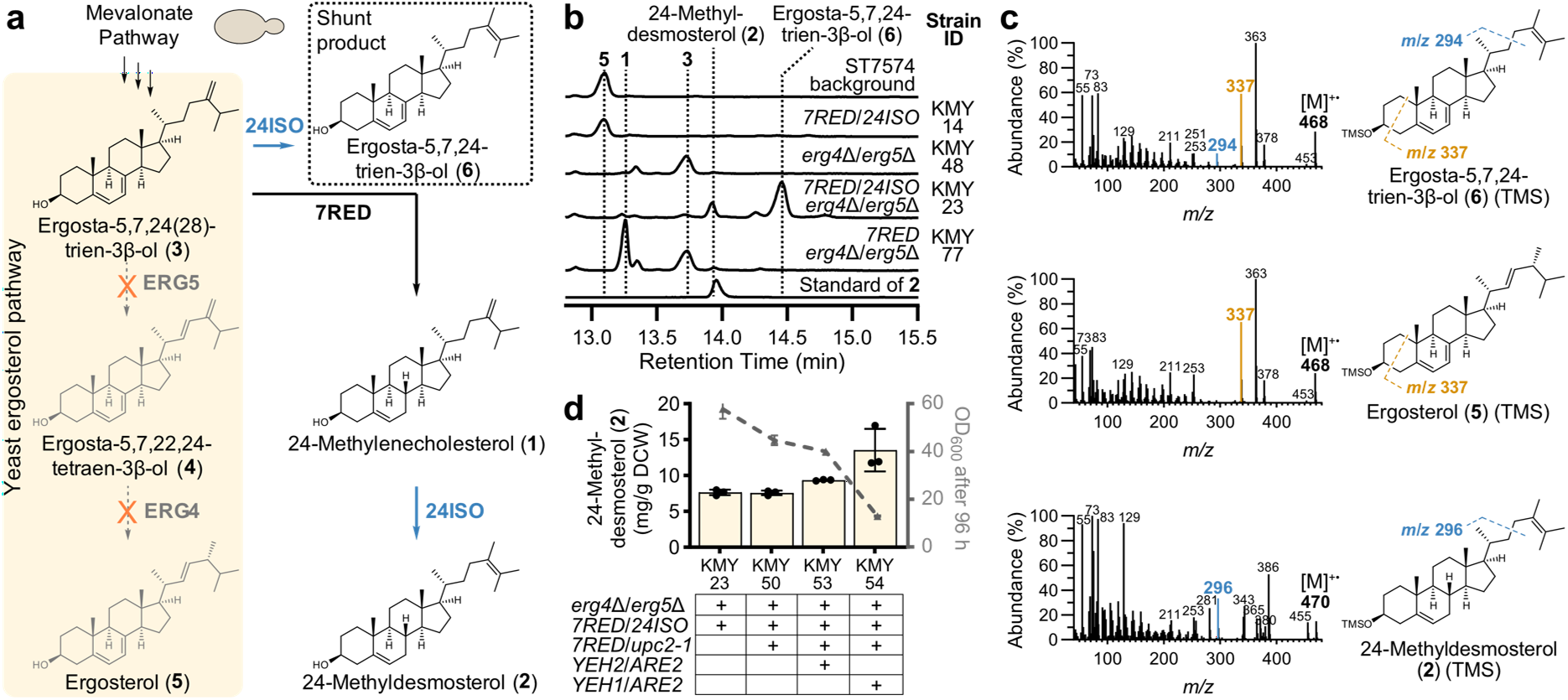
Metabolic engineering of yeast for producing the key intermediate 24-methyldesmosterol (2). **a** Yeast engineering strategy to divert flux from ergosterol (**5**) biosynthesis to 24-methyldesmosterol (**2**). **b** GCMS total ion current chromatograms showing production of 24-methyldesmosterol (**2**) and the shunt product ergosta-5,7,24-trien-3β-ol (**6**) in engineered yeast strain KMY23. **c** Comparison of electron impact mass spectra of ergosta-5,7,24-trien-3β-ol (**6**) and related compounds as TMS ethers supporting the proposed structure of **6**. **d** Further yeast engineering to improve 24-methyldesmosterol (**2**) production. DCW: Dry cell weight.

With the help of established CRISPR/Cas techniques^65^, we inserted *Physalis peruviana* orthologues of the known plant genes *sterol Δ7 reductase* (*7RED*, also known as *DWF5*)^66^ and *24ISO*^14^ into the genome of the prototrophic *S. cerevisiae* strain ST7574, which is derived from CEN.PK113-7D and contains a *cas9* gene^65^. The resulting strain KMY14 only produced trace amounts of 24-methylenecholesterol (**1**) and 24-methyldesmosterol (**2**) (Fig. 4b). This result was in line with previous reports showing that under native conditions the ergosterol biosynthetic enzymes ERG4 and ERG5 efficiently consume the shared intermediate ergosta-5,7,24(28)-trien-3β-ol (**3**) via ergosta-5,7,22,24(28)-tetraen-3β-ol (**4**) to ergosterol (**5**)^14,63,64^. Upon deletion of *ERG4* and *ERG5* in the background strain ST7574 (strain KMY48), the required shared intermediate ergosta-5,7,24(28)-trien-3β-ol (**3**) was formed as the major sterol as reported in literature^67,68^. To redirect this intermediate to the desired product 24-methyldesmosterol (**2**), we deleted *ERG4* and *ERG5* in the *7RED*/*24ISO*-expressing strain KMY14 to generate KMY23. Gratifyingly, KMY23 produced 24-methyldesmosterol (**2**) at a level of 7.6 mg/g dry cell weight (DCW) (Fig. 4b). Besides 24-methyldesmosterol (**2**), a major peak **6** at 14.5 min was observed in yeast, with a mass difference of −2 compared with 24-methyldesmosterol (**2**) (Fig. 4b). The mass spectrum of this compound **6** was almost identical to that of ergosterol (**5**) (Fig. 4c), but the difference in retention time indicated that **6** must be an isomer of ergosterol (**5**) (Fig. 4b). A striking difference in their mass spectra was the presence of a fragment at *m*/*z* 294 for **6** that was absent for ergosterol (**5**); in the mass spectrum of 24-methyldesmosterol (**2**), this fragment was shifted to *m*/*z* 296, matching the mass difference of the molecular ions. Previous studies reported that this fragment is indicative of sterols with a Δ^24(28)^ or Δ^24(25)^ double bond and is formed by a McLafferty rearrangement involving allylic cleavage of the C-22(23) bond^68–71^. The mass difference of −2 suggested that **6** must possess an additional double bond in the ABCD ring system. Ergosterol (**5**) and **6** both exhibit a major fragment at *m*/*z* 337 ([M−131]^+^), which is characteristic of sterols with Δ^5,7^ dienes^72,73^; therefore, the additional double bond of **6** is most likely positioned at C-7,8. We therefore propose that compound **6** is ergosta-5,7,24-trien-3β-ol (Fig. 4c). This Δ^7^ analogue of 24-methyldesmosterol (**2**) would be formed if ergosta-5,7,24(28)-trien-3β-ol (**3**) is converted by 24ISO before reduction by 7RED can take place. This hypothesis was also supported by the fact that upon deletion of *24ISO* the peak for ergosta-5,7,24-trien-3β-ol (**6**) completely disappeared (strain KMY77); instead, a mixture of the 7RED product 24-methylenecholesterol (**1**) and non-reduced ergosta-5,7,24(28)-trien-3β-ol (**3**) was observed (Fig. 4b). The occurrence of the non-reduced products ergosta-5,7,24-trien-3β-ol (**6**) in KMY23 and ergosta-5,7,24(28)-trien-3β-ol (**3**) in KMY77 therefore indicates a bottleneck caused by limited activity of *P. peruviana* 7RED in our yeast system.

To facilitate subsequent gene discovery, we next wanted to improve the production of 24-methyldesmosterol (**2**) in yeast. First, we added an additional copy of *P. peruviana 7RED* to improve the conversion of ergosta-5,7,24(28)-trien-3β-ol (**3**) to 24-methylenecholesterol (**1**); simultaneously, we overexpressed the *upc2-1* allele, encoding transcription factor mutant UPC2^G888A^ involved in the regulation of sterol metabolism and reported to boost the biosynthesis of sterols^74,75^. The resulting strain KMY50, however, did not show elevated levels of 24-methyldesmosterol (**2**). Previous studied showed that manipulation of sterol homeostasis, i.e., the balance of sterol acylation and sterol ester hydrolysis, can improve the production of sterols in yeast^76^. We therefore co-expressed the acyl-CoA:sterol acyltransferase gene *ARE2* either with the sterol ester hydrolase gene *YEH2* (KMY53) or with *YEH1* (KMY54). In both cases, an improvement in 24-methyldesmosterol (**2**) levels was noted. KMY53 produced 9.3 mg/g DCW, whereas KMY54 reached 13.5 mg/g DCW (Fig. 4d). However, the final OD_600_ after cultivation for 96 h in shake flasks was severely reduced, particularly for strain KMY54. In summary, our data show that the key intermediate 24-methyldesmosterol (**2**) can be produced in engineered yeast, but further strain improvement is required to improve product levels, reduce the amounts of shunt product **6**, and overcome growth defects.

### Engineering of *Nicotiana benthamiana* for production of 24-methyldesmosterol

Due to these unresolved limitations of yeast as a platform for withanolide pathway reconstitution, we alternatively envisioned to generate a plant-based system using the popular model organism *Nicotiana benthamiana*^77^. In contrast to yeast, plants natively produce 24-methylenecholesterol (**1**) as a transient intermediate en route to phytosterols and brassinosteroids.^78^ We therefore expected that usage of a plant host would enable us to circumvent the undesired yeast shunt product ergosta-5,7,24-trien-3β-ol (**6**) linked with insufficient 7RED activity. Furthermore, transient expression in *N. benthamiana* provides the added benefit of easy and flexible “mix-and-match” co-expression, which would drastically accelerate the pace of testing different gene combinations^77^. We first transiently expressed only *24ISO* in *N. benthamiana*; while this caused changes in the metabolic profile, no 24-methyldesmosterol (**2**) was detected. This suggested that insufficient amounts of 24-methylenecholesterol (**1**) accumulate under native conditions in *N. benthamiana*. We therefore sought to increase the formation of 24-methylenecholesterol (**1**) and hence 24-methyldesmosterol (**2**) by metabolic engineering. In principle, this could be achieved by two complementary strategies: Either by blocking the pathway downstream of 24-methylenecholesterol (**1**) controlled by the reductase DWF1^79^ (strategy I), or overexpressing its upstream phytosterol pathway (strategy II) (Fig. 5a). We first attempted to silence *DWF1* by virus-induced gene silencing^80^. In accordance with previous reports^79^, this caused a very strong dwarf phenotype and sick-looking plants (Supplementary Fig. 11a), probably due to the effects on brassinosteroid levels. After transient overexpression of *24ISO* in *DWF1*-silenced *N. benthamiana* plants, small quantities of 24-methyldesmosterol (**2**) could be detected (Supplementary Fig. 11b). Nonetheless, the experimental challenges of coordinating the relatively slow systemic gene silencing with the relatively fast local transient overexpression and the severe phenotypic effects on the plants prompted us to test upstream pathway overexpression as an alternative strategy (Fig. 5a). First, we co-expressed *24ISO* with a gene encoding truncated, feedback-insensitive hydroxymethylglutaryl coenzyme A reductase (tHMGR), which is known as a very efficient booster for the mevalonate pathway and triterpenoid production^49^. In these samples, we observed a very small peak below the limit of quantification corresponding to 24-methyldesmosterol (**2**) (Fig. 5b). We concluded that further reactions of the phytosterol pathway limit the 24-methyldesmosterol (**2**) yield.

**Fig. 5.**
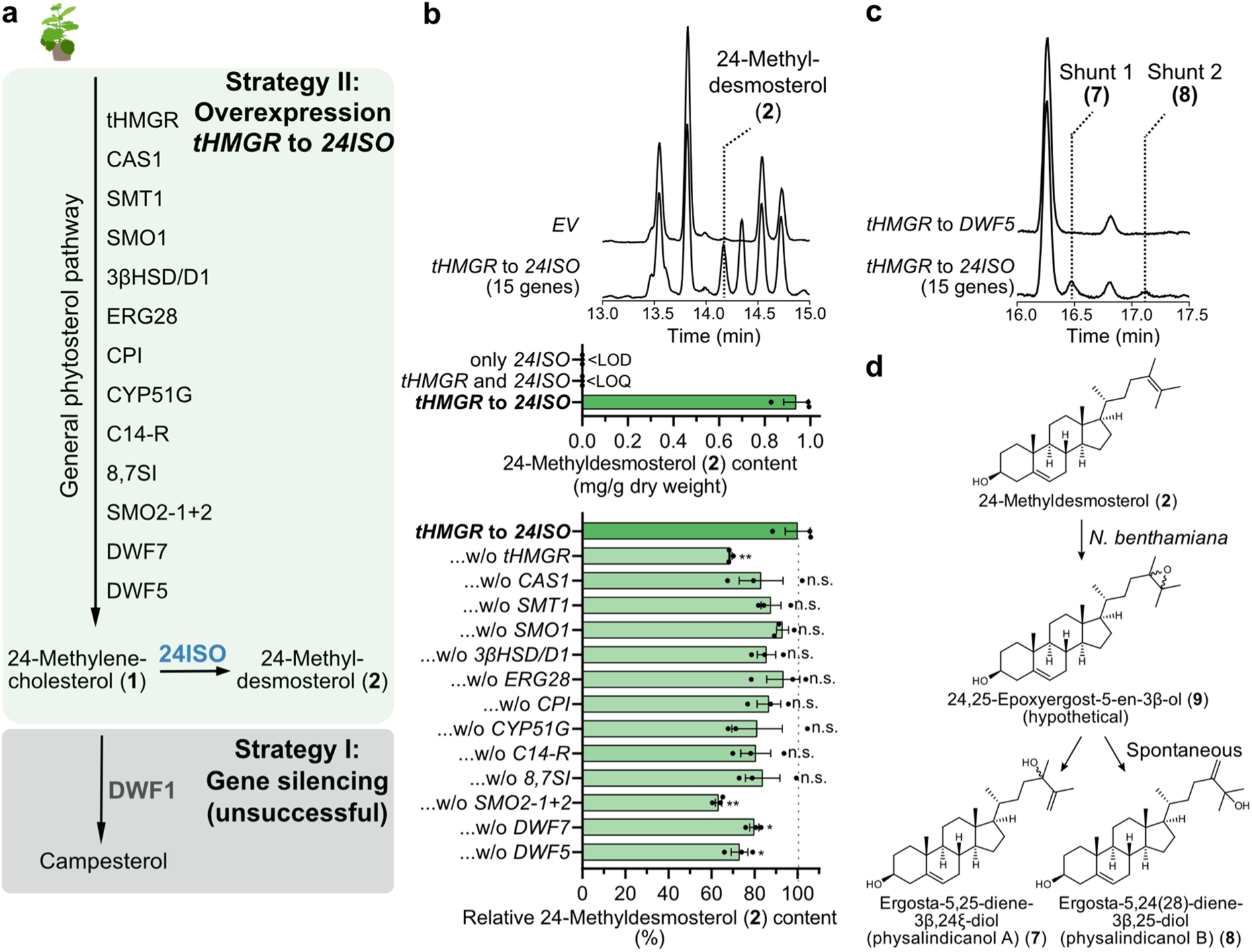
Metabolic engineering in *Nicotiana benthamiana* for producing the key intermediate 24-methyldesmosterol (2). **a** Overview of the two different engineering strategies that were attempted in this work. **b** Effect of phytosterol pathway gene overexpression (strategy II) on production of 24-methyldesmosterol (**2**). Total ion current (TIC) GC-MS chromatograms of representative samples from transient expression in *N. benthamiana*. The bottom bar plot indicates relative 24-methyldesmosterol (**2**) levels when a single gene of the 15-gene set from *tHMGR* to *24ISO* was left out. Bar plots show means ± SEM and data points of three biological replicates. * p < 0.05, ** p < 0.01, NS not significant by Student’s *t* test; LOD: limit of detection, LOQ, limit of quantification. **c** GC-MS TIC chromatograms showing formation of two shunt products **7** and **8** dependent on *24ISO* in *N. benthamiana*. **d** Proposed formation of shunt products **7** and **8** by background epoxidation of the Δ^24^ double bond in *N. benthamiana*.

As no comprehensive information is available which enzymes en route to 24-methylenecholesterol (**1**) are rate-limiting, we decided to transiently overexpress the full gene set for the phytosterol pathway reported from *Arabidopsis thaliana* (Supplementary Fig. 12) in addition to *tHMGR* and *24ISO* in *N. benthamiana*. This comprised a total of 15 genes. Gratifyingly, this approach led to a drastic increase in 24-methyldesmosterol (**2**) formation to 0.9 mg/g dry weight (Fig. 5b). Notably, the byproduct ergosta-5,7,24-trien-3β-ol (**6**) was not observed, supporting our hypothesis that the 7RED bottleneck encountered in yeast can be circumvented by using a plant host. To understand which upstream pathway genes had the largest effect on 24-methyldesmosterol (**2**) titres, we performed a leave-one-out experiment (Fig. 5b). Statistically significant negative effects on 24-methyldesmosterol (**2**) levels were observed when *tHMGR*, *SMO2-1* plus *SMO2-2*, *DWARF5*, or *DWARF7* were not included, suggesting that these represent major bottlenecks towards 24-methylenecholesterol (**1**). The chromatograms of *N. benthamiana* plants producing 24-methyldesmosterol (**2**) also contained two new product peaks dependent on the presence of *24ISO* with a mass difference of +88 compared with 24-methyldesmosterol (**2**), which would fit to an extra trimethylsilyl (TMS)-bearing oxygen (Fig. 5c). Analysis of fragmentation patterns suggested that these compounds were oxidised in the side chain (Supplementary Fig. 13-15, Supplementary Table 3). The two compounds **7** and **8** were successfully isolated (isolated yield 0.03 and 0.03 mg/g dry weight, respectively) and fully characterised by NMR spectroscopy (Supplementary Fig. 16, Supplementary Table 4-5). Surprisingly, in comparison to 24-methyldesmosterol (**2**), the Δ^24^ double bond of **7** and **8** was shifted to Δ^25^ or Δ^24(28)^, respectively. In agreement with the fragmentation analysis, NMR data showed that both products **7** and **8** contained an additional hydroxy group in allylic position of the double bond in the side chain. Hence, the systematic name of **7** is ergosta-5,25-diene-3β,24ξ-diol, while **8** is ergosta-5,24(28)-diene-3β,25-diol. Both compounds were isolated before from the withanolide-producing plant *Physalis minima* var. *indica* and named physalindicanol A (**7**) and B (**8**), respectively^81^. The occurrence of these regioisomeric allylic alcohols can be explained by background epoxidation of the Δ^24^ double bond of 24-methyldesmosterol (**2**)^81^ in *N. benthamiana*, generating hypothetical intermediate **9**; either *in vivo* or during workup, this epoxide can then be opened to allylic alcohols **7** and **8** (Fig. 5d).

We concluded that, despite the undesirable background epoxidation, our metabolic engineering strategy in *N. benthamiana* reaching 0.9 mg/g dry weight of 24-methyldesmosterol (**2**) was suitable for further elucidation of withanolide biosynthesis.

### Elucidation of oxidative lactone formation in withanolide biosynthesis

After establishing heterologous platforms capable of producing the last known withanolide biosynthesis intermediate 24-methyldesmosterol (**2**), we turned our attention to screening genes from the gene cluster (Fig. 3) to identify the next steps in withanolide biosynthesis. Labelling studies demonstrated that withanolide biosynthesis proceeds by oxidative assembly of the side chain lactone^82,83^. From all oxidative enzymes, only the cytochrome P450 monooxygenases CYP87G, CYP88C, and CYP749B were conserved within all withanolide-producing species (Fig. 3) and therefore prioritised. For simplification, orthologues with >90% sequence identity were excluded from the initial screening. The genes of the three CYPs *Pp*CYP87G-a, *Pp*CYP88C-d, and *Pp*CYP749B-b from *Physalis pruinosa* were expressed in different combinations in our yeast and *N. benthamiana* platforms producing 24-methyldesmosterol (**2**); these CYPs were named *Pp*CYP87G1, *Pp*CYP88C7, and *Pp*CYP749B2, respectively (Fig. 6a). In yeast, a cytochrome P450 reductase gene from *Arabidopsis thaliana* was included.

**Fig. 6.**
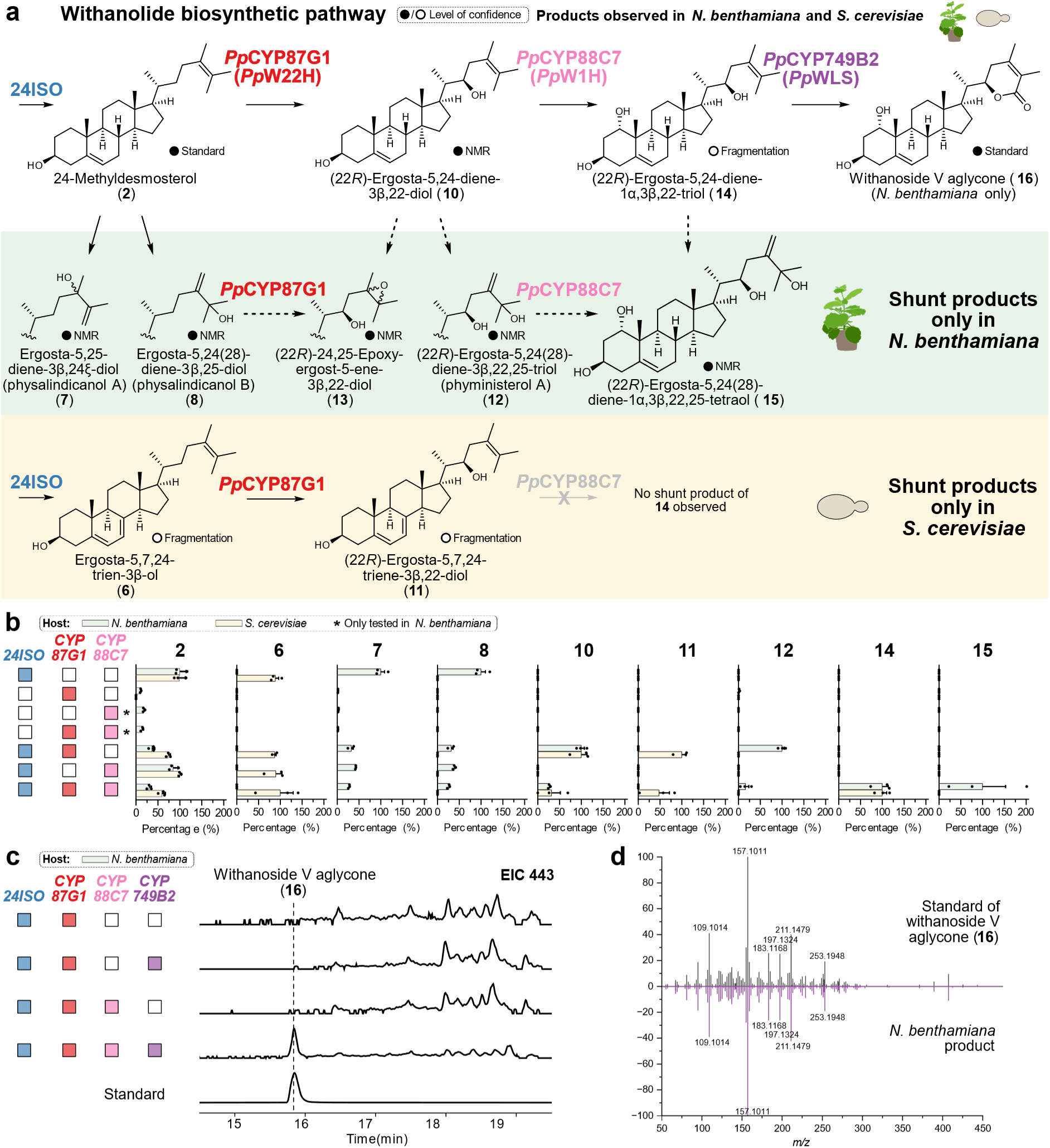
Elucidation of CYP-catalysed oxidative steps leading to withanoside V aglycone (16). **a** Proposed formation of intermediates and shunt products in *N. benthamiana* and *S. cerevisiae* along the biochemical pathway to withanoside V aglycone (**16**). Dashed arrows indicate alternative pathways to *N. benthamiana* shunt products. **b** Relative levels of identified compounds in GC-MS chromatograms of *N. benthamiana* (green) and *S. cerevisiae* (yellow). For each compound, peak areas were normalised to internal standard and sample dry weight and converted to relative amounts with the highest mean value set to 100%. Bar plots show means ± SEM and data points of three biological replicates. **c** Extracted ion LC-MS chromatograms of *m*/*z* 443 (low resolution) from *CYP* co-expression in *N. benthamiana* leading to formation of withanoside V aglycone (**16**) in comparison to an authentic reference compound. **d** Mirror plot comparing high resolution MS/MS spectra (35 eV) of authentic withanoside V aglycone (**16**) and the product from *N. benthamiana*. *Pp*: *Physalis pruinosa;* W22H: withanolide 22-hydroxylase; W1H: withanolide 1-hydroxylase; WLS: withanolide lactone synthase.

Of the three tested CYP genes, only *PpCYP87G1* showed activity detectable by GC-MS in yeast and *N. benthamiana*. In both hosts, a new product **10** with a mass shift of +88 was detected (Fig. 6b). Analysis of the mass spectrum from electron impact ionisation suggested that **10** might be a hydroxylation product of 24-methyldesmosterol (**2**) at C-22 (Supplementary Fig. 13-15, Supplementary Table 3). We isolated the shared product **10** from *N. benthamiana* (0.04 mg/g dry weight isolated yield) and elucidated its structure by NMR spectroscopy (Supplementary Fig. 16, Supplementary Table 4-5). In agreement with our GC-MS fragmentation analysis, **10** contained an additional hydroxy group at C-22. Due to the flexibility and free rotation of the side chain, the stereochemistry at C-22 could not be confidently deduced by nuclear Overhauser effect spectroscopy (NOESY). Instead, we compared the coupling pattern of H-22 of our isolated compound **10** with two pairs of structurally related C-22 epimers that were obtained by total synthesis (Supplementary Fig. 17). For the two synthetic 22*S* epimers, doublets of doublets were observed at H-22, in contrast to apparent doublets of triplets for the two synthetic *22R* epimers. As compound **10** also showed an apparent doublet of triplets for H-22, C-22 was assigned as 22*R*, which is also the configuration expected for natural withanolides. In conclusion, compound **10** was confirmed to be (22*R*)-ergosta-5,24-diene-3β,22-diol (22*R*-hydroxy-24-methyldesmosterol). Besides shared product **10**, we also observed further compounds from *PpCYP87G1* expression occurring exclusively either in yeast or in *N. benthamiana*. In yeast, a second compound **11** with a mass shift of −2 compared with **10** was produced, while in *N. benthamiana* further compounds **12** and **13** with mass differences of +88 and +16 compared with **10** were detected (Fig. 6b). As the exclusive occurrence in either yeast or *N. benthamiana* as well as the mass shifts were in excellent agreement with the occurrence of shunt products related to 24-methyldesmosterol (**2**) observed during our metabolic engineering efforts (Fig. 4b and Fig. 5c), we strongly suspected that these products reflected the same background reactions and not additional enzymatic activity of *Pp*CYP87G1. To confirm this, we isolated the putative shunt products **12** and **13** from *N. benthamiana* (0.04 and 0.01 mg/g dry weight isolated yields). NMR spectroscopy indicated that both products contained a hydroxy group at C-22 (Supplementary Fig. 16, Supplementary Table 4-5). In addition, compound **12** possessed the same Δ^24(28)^ double bond and C-25 hydroxy group as shunt product **8**. Compound **13** did not contain any olefinic carbons in the side chain, but instead two carbons at 62.6 and 65.6 ppm, respectively, which imply that **13** contains an epoxy group. The isolation of epoxide **13** strongly supports our hypothesis that the rearranged allylic alcohols are derived from Δ^24^ epoxidation (Fig. 5d). Compound **12**, named (22*R*)-ergosta-5,24(28)-diene-3β,22,25-triol was isolated before from *Physalis minima* and has been named phyministerol A^84^; our NMR data of **12** are in very good agreement with the published data (Supplementary Table 6). Compound **13** or (22*R*)-24,25-epoxyergost-5-ene-3β,22-diol is a new compound, but steroids with matching side chains have been isolated from the nightshade plant *Petunia hybrida* (Supplementary Fig. 18)^85^.

Next, we co-expressed *PpCYP87G1* with the other two CYP genes in yeast and *N. benthamiana*. Only for the combination of *PpCYP87G1* and *PpCYP88C7* we observed new compounds by GC-MS (Fig. 6b). One product **14** with a mass shift of +88 compared with the *Pp*CYP87G1 product **10** was found in both heterologous hosts. In *N. benthamiana*, extra peaks with an additional +88 mass shift were observed; no extra peak was detected in yeast. The mass spectra of all *Pp*CYP88C7-dependent peaks showed a *m*/*z* 217 fragment, which has been reported as a diagnostic ion for 1,3-dihydroxylated cyclohexanes^86^ (Supplementary Fig. 13-15, Supplementary Table 3). The fragmentation pattern therefore suggested that *Pp*CYP88C7 carries out hydroxylation of C-1. Although we could not purify sufficient amounts of **14** for NMR spectroscopy, we successfully obtained shunt product **15** from *N. benthamiana* (isolated yield 0.01 mg/g dry weight). NMR data indicated that the side chain of **15** was identical to shunt product (22*R*)-ergosta-5,24(28)-diene-3β,22,25-triol (phyministerol A) (**12**). However, one of the other carbons exhibited a drastic downfield shift to 73.1 ppm, indicating an additional hydroxy substituent. An HMBC correlation from methyl group H-19 and COSY correlations with H-2 unambiguously confirmed that the new hydroxy group was located at C-1 (Supplementary Fig. 16). A NOE correlation between methyl group H-19 and H-1 indicated an α configuration of C-1 (Supplementary Fig. 19). Previously isolated steroids from nightshade plants with a hydroxy group at C-1, such as withanosides, also possess a C-1 α configuration^87^.

Lastly, only when *PpCYP87G1*, *PpCYP88C7*, and *PpCYP749B2* were co-expressed, a new peak was observed by LC-MS from *N. benthamiana* samples that matched the aglycon of the known withanolide glycoside withanoside V (**16**) ^87^ in terms of retention time, high-resolution mass and MS/MS fragmentation (Fig. 6c and Fig. 6d). This compound was not observed when samples were saponified before LC-MS analysis. Co-expression of *PpCYP87G1* and *PpCYP749B2* without *PpCYP88C7* did not result in lactone formation, suggesting that lactone formation in the side chain cannot take place before C-1 hydroxylation.

In summary, our data provide support for a three-stage oxidative assembly of the characteristic lactone ring of withanolides (Fig. 6a): First, C-22 is hydroxylated by *Pp*CYP87G1; second, hydroxylation of C-1 is carried out by *Pp*CYP88C7; lastly, the lactone ring is generated by oxidation of C-26 and simultaneous lactonisation by *Pp*CYP749B2. Therefore, we propose the trivial names withanolide 22-hydroxylase (W22H) for *Pp*CYP87G1, withanolide 1-hydroxylase (W1H) for *Pp*CYP88C7, and withanolide lactone synthase (WLS) for *Pp*CYP749B2. Together, these three CYPs are therefore responsible for the biosynthesis of the key δ-lactone ring and for preparing the enone formation in the A ring of withanolides.

## Discussion

In this work, we identified a gene cluster for withanolide biosynthesis in Solanaceae plants. The gene cluster reported here adds to several previously reported cases of gene clustering in specialised metabolism in Solanaceae plants, for example for biosynthesis of glycoalkaloids, modified fatty acids, acyl sugars, or terpenoids^23,53,88,89^. As such, the Solanaceae family represents an increasingly intriguing model system to study gene clustering in plants. The function of the gene cluster described here for withanolide biosynthesis is supported by our phylogenomics analysis, the biochemical activity of *Pp*CYP87G1, *Pp*CYP88C7, and *Pp*CYP749B2 in yeast and *N. benthamiana* enabled by metabolic engineering, and gene silencing results by others^20,61^. Similar to examples from other plants^90^, our synteny analyses highlight the plasticity of plant biosynthetic gene clusters. Even in closely related species of the same genus, changes in gene order and orientation as well as gene duplications are not unusual. How these differences are relevant for increasing the structural diversity of withanolides will need to be addressed in the future. A unique feature of the withanolide gene cluster is the consistent occurrence of sub-gene clusters, as supported by gene expression (Supplementary Fig. 2-4) and phylogenetic analyses (Supplementary Fig. 5-10). Gene duplications followed by neofunctionalization are common drivers of evolution^30^. In the case of withanolide biosynthesis, it appears that such an evolutionary event has occurred not only on the single gene level, but at a whole gene cluster level. Future studies will need to carefully address functional differences between the sub-gene clusters to gain further insights into the evolution and biological role of withanolides in nightshade plants.

Elucidating and engineering the biosynthetic pathways of specialised steroids from plants still represents a major challenge. While many other compound classes such as alkaloids or other terpenoids are nowadays readily accessible in yeast or *N. benthamiana,* progress for steroids has been lagging behind. Only recently, the first successful metabolic engineering examples of steroidal natural products derived from the more common precursor cholesterol were published^52,91–93^. These have been facilitated by the fact that cholesterol natively accumulates in *N. benthamiana*, albeit at low levels. In contrast, our data shows that the production of relevant levels of 24-methylenecholesterol (**1**) or 24-methyldesmosterol (**2**) in yeast or *N. benthamiana* requires substantial additional engineering of sterol metabolism. In yeast, lowering the formation of shunt products arising from insufficient activity of 7RED will be critical. The production of withanolide pathway intermediates in *N. benthamiana* appeared more efficient, even though an extensive set of genes had to be co-expressed. For maximum production of 24-methyldesmosterol (**2**) in *N. benthamiana*, we had to transiently overexpress 15 genes at the same time. To our knowledge, this setup is amongst the most complex examples reported to date^94–96^. Once again, this highlights the tremendous power of efficient and facile co-expression in *N. benthamiana* for gene discovery and pathway engineering in plants^77^. Our metabolic engineering efforts now enable further elucidation of withanolide biosynthesis, in order to close the gap from traditional medicine to modern drug development and production.

## Methods

### Plant materials

*W. somnifera* (XX-0-STGAL—22/1985) plants were grown in the Botanical Garden of TU Braunschweig in a greenhouse with natural light at approximately 20 °C. Genes from *P. pruinosa* were obtained as synthetic genes from GeneWiz (Azenta Life Sciences) based on the published genome sequence^37^ and our re-annotation without handling of the plant.

### Genome sequencing, assembly and annotation

DNA extraction, quality control, short fragment depletion, and DNA repair for the Oxford Nanopore sequencing was performed as previously described^97^. Sequencing was performed via MinION Mk1B on R9.4.1 flow cells. Basecalling was performed with Guppy v 6.4.6+ae70e8f in high accuracy mode. A genome sequence was constructed with NextDenovo2 v2.5.2 (read_cutoff=1k, genome_size = 3g, and seed depth = 30)^98^. RagTag v2.1.0^99^ was used to reorder the contigs in the assembly based on homology with the close relative *P. pruinosa* genome assembly for direct comparison.

RNA-seq reads (Supplementary Table 7) were aligned with HISAT 2.2.1 ^100^ using default parameters to generate hints for the gene prediction. A homology-based gene prediction of the *W. somnifera* genome was performed using GeMoMa v1.9^101,102^, based on annotations of six Solanoideae sub-family species (*Physalis floridana*^36^, *Datura stramonium*^34^, *Iochroma cyaneum*^103^, *Atropa belladonna*^34^, *Lycium barbarum*^104^, and *Solanum lycopersicum*^42^). For each of these six species, extracted coding sequences were aligned to the *W. somnifera* genome sequence with MMseqs2^105^ (version 15-6f452) using default parameters suggested by GeMoMa. For predicting gene models, GeMoMa utilised protein-coding gene alignments, intron position conservation, and RNA-seq data for refining intron boundaries. The resulting six gene annotation sets were filtered and merged using GeMoMa Annotation Filter (GAF) with parameters f=“start==’M’ and stop==’*’ and (isNaN(score) or score/aa>=3.50)” and atf=“tie==1 or sumWeight>1”. GeMoMa was also used to annotate the genome sequences of *P. pruinosa* and *P. grisea* using identical parameters, with a modified filtering step, f=“start==’M’ and stop==’*’ and (isNaN(score) or score/aa>=4.00)” and atf=“tie==1 and sumWeight>1” for both species. BUSCO v5.6.1^106^ with the Solanales dataset was used to assess the completeness of the genome assemblies and annotations. The sources of genomic data sets are listed in Supplementary Table 8.

### Identification of transposable elements

Transposable elements (TEs) and tandem repeats were annotated using Extensive *de-*novo TE Annotator^107^ (EDTA) v2.2.1 and Tandem Repeats Finder (TRF)^108^ v4.10.0, respectively, as described in the Supplementary Methods.

### Synteny analysis

JCVI/MCscan^109^ v1.4.14 was employed to compare the local synteny among the genome sequences of known withanolide-producing species, *P. floridana*, *P. pruinosa*, *P. grisea*, *W. somnifera*, *D. stramonium* and *D. wrightii*. Non-withanolide producing species, *S. lycopersicum*, *S. tuberosum* and *N. tabacum* were also included. The sources of genomic data sets are listed in Supplementary Table 8. The region surrounding the *P. pruinosa 24ISO* gene was selected to identify microsynteny among these species. Connection of genes between species was validated and manually revised based on phylogenetic trees (Supplementary Fig. 5-10).

### Phylogenetic analyses

Phylogenetic trees were constructed using a total of 32 Solanaceae species (Supplementary Table 8). For each gene family, *Physalis pruinosa* gene sequences from the biosynthetic gene cluster were used as queries in BLASTp^110^ searches against the polypeptide sequences of these 32 plant species, and the top BLAST hits were selected. This was performed using the Python script collect_best_blast_hits.py available at https://github.com/bpucker/ApiaceaeFNS1^111^. To differentiate between related gene families, additional outgroup sequences were included: *SSR1* and *SSR2* for *24ISO*; flavonoid biosynthesis genes *F3’H* and *F3’5’H* for *CYP*s; *DFR* for *SDR*s; and *FLS*, *F3H*, and *AOP* for *ODD*s. Additional plant sequences for *ST*s and *AT*s were also incorporated into their respective trees. Sequences were aligned using MAFFT^112^ v7.520 with default settings and the L-INS-i accuracy-oriented method. Alignments with occupancy below 10% were removed using pxclsq from PHYX^113^. The final phylogenetic trees were generated using IQ-TREE^114^ v1.6.12, with 1,000 bootstrap replicates. The best-fit evolutionary models were inferred using ModelFinder^115^: JTT+R5 for 24ISO, JTT+F+R10 for CYPs, JTT+R9 for SDRs, JTT+F+R6 for STs and ATs, and JTT+F+R9 for ODDs. Phylogenetic trees were visualised and annotated using iTOL^116^ v6.8.

### Expression analysis

To evaluate the expression of genes from identified gene clusters in withanolide-producing species analysed in this study, publicly available paired-end RNA-seq datasets were collected. However, sufficient datasets with diverse tissue samples were only available for *D. stramonium* and *W. somnifera*. The FASTQ files for both species were retrieved from the Sequence Read Archive (SRA) using prefetch and fasterq-dump from NCBI SRA toolkit v3.1.0. Transcript abundance was quantified using Kallisto v0.50.1^117^ based on the coding sequences. The individual count tables were then merged and filtered as previously described^111,118^. Expression heatmaps were generated in R to visualise the expression patterns of the identified gene clusters.

### Metabolic engineering of *S. cerevisiae*

For metabolic engineering of yeast (*S. cerevisiae*), the EasyClone-MarkerFree system was used^65^. All yeast strains generated in this study were derived from *S. cerevisiae* ST7574 (CEN.PK background) expressing *cas9* available on Euroscarf. Biobricks were assembled *via* USER cloning^119^. Amplification of all genes and promoters was performed using Phusion U polymerase (Thermo Fisher Scientific) with primers containing uracil overhangs. Resulting biobricks were assembled into AsiSI/Nb.BsmI (NEB) digested integrative vectors by USER cloning. Assembled synthetic constructs were transformed into NEB 5-alpha competent *E. coli* cells (NEB). Positive clones as judged by colony PCR were verified through Sanger sequencing (Microsynth Seqlab). Linear integration fragments containing the target genes flanked by homology arms for genomic integration were generated by NotI-HF (NEB) digestion. Linearised synthetic constructs were introduced into *S. cerevisiae* using the standard lithium acetate method^120^. Site-specific integrations were confirmed using standardised primers included in the EasyClone-MarkerFree Vector Set. Yeast strains generated in this study are listed in Supplementary Table 9, primers in Supplementary Table 10, plasmids in Supplementary Table 11, and coding sequences of inserted genes in Supplementary Table 12.

For gene deletions, homology-directed repair templates were prepared with High-Fidelity Q5 polymerase (NEB) using a PCR with 10 cycles and primers containing overlapping sequences. Guide RNAs (gRNAs) for deleting *ERG4*, *ERG5* and *24ISO* were designed using CHOPCHOP^121^. The required 20 nt gRNA sequences (Supplementary Table 13) were introduced into pCfB8792 (Addgene Plasmid # 126910) by inverse PCR using primers containing gRNA sequences (Supplementary Table 13) as overhang. The resulting amplicons containing modified gRNA sequences were gel purified (Macherey-Nagel) and treated with DpnI (NEB) for 30 min at 37 °C. DpnI-digested product was ligated using T4 DNA ligase (NEB) for 2 h at room temperature and transformed into NEB 5-alpha competent *E. coli* cells. Sequencing was done to confirm the gRNA sequence from isolated plasmids. Cultures for yeast strains were set in triplicates for extraction of metabolites. Primary cultures were cultivated at 30 °C under continuous shaking at 210 rpm in YPD broth (Carl Roth) containing peptone (20 g/L), yeast extract (10 g/L) and glucose (20 g/L) supplemented with 200 mg/L G418. Overnight primary cultures were used to set secondary cultures with a starting OD_600_ ∼ 0.2 in YPD medium without antibiotics. Secondary cultures of synthetic constructs harbouring galactose-inducible promoters were induced with YP medium containing peptone (20 g/L), yeast extract (10 g/L) supplemented with filter-sterilised galactose (20 g/L). Secondary cultures were grown for 4 days and OD_600_ was measured just before harvesting; then, metabolite profiles were analysed by GC-MS as described below.

### Transient expression in *Nicotiana benthamiana*

Transient expression in *Nicotiana benthamiana* was performed with *N. benthamiana* LAB strain, which was grown from seeds either in a greenhouse with 11 to 16 hours of illumination per day at 21-23 °C as described previously^122^ or in a phytochamber with 16.5/7.5 h photoperiod at 100 μmol m^-2^ s^-1^ light intensity and a temperature of 22 °C during the day and 20 °C during the night. Genes were cloned into the plasmid pHREAC^123^ by Golden Gate cloning or In-Fusion cloning as reported^122^. All primers for metabolic engineering and transient expression in *N. benthamiana* are listed in Supplementary Table 14. The resulting plasmids were transformed into *Agrobacterium tumefaciens* GV3101 by electroporation. The *A. tumefaciens* strains were cultured in LB medium with antibiotics (25 μg/mL gentamicin, 50 μg/mL rifampicin, 50 μg/mL kanamycin) for 2 days at 28 °C with shaking. Cells were harvested, resuspended in MMA infiltration buffer (10 mM MgCl_2_, 10 mM MES, 100 μM acetosyringone), and adjusted to OD_600_ 0.1. For co-expression of multiple genes, the corresponding strains were mixed so that each strain had a final OD_600_ 0.1. The mixtures were syringe-infiltrated into the abaxial side of 4-week-old *N. benthamiana* leaves. For compound purification, vacuum infiltration was used instead as described below. After infiltration, plants were maintained in a greenhouse or phytochamber until further analysis. Leaf samples were harvested 7 days post-infiltration for GC-MS and LC-MS analysis as described below.

### GC-MS sample preparation

GC-MS analysis of yeast strains was performed with cell pellets. Cells were harvested at 5,000 rpm for 5 min and washed twice with sterile water. Harvested cells (1.5 mL) were saponified at 90 °C for 10 min with 500 µL of 20% aqueous KOH in 50% ethanol containing 5 µg/mL of 5α-cholestane as internal standard. To the saponified extracts, 0.5 mm glass beads (Carl Roth, Germany) and either 500 µL hexane or 500 µL ethyl acetate were added depending on the expected polarity window of metabolites. Samples were then lysed in a FastPrep-24TM 5G homogeniser (MP Biomedicals, CA, USA) with predefined settings for *S. cerevisiae*. The organic layer was transferred to a glass vial, and the extraction process repeated a second time. Both organic layers were pooled in a glass vial and concentrated *in vacuo* using an RVC 2-25 CDplus (Christ, Osterode am Harz, Germany) rotary vacuum concentrator with a cooling trap. Dried extracts were derivatised using 1:1 pyridine : (BSTFA+1% TMCS) at 70 °C for 1 h. GC-MS analysis was carried out as described below. For GC-MS analysis of *N. benthamiana* leaf samples after transient expression, 10 leaf disks were excised using a cork borer no. 5 (ø = 10 mm) and lyophilised until their weight remained constant (≈ 20 mg). Then, a 5 mm steel bead was added, and the disks were ground in a MM 400 ball mill (Retsch, Haan, Germany) at 30 Hz for 20 s. Ground leaves were saponified at 70 °C for 1 h with 500 µL of 10% aqueous KOH in 90% ethanol, containing 10 µg/mL of 5α-cholestane as internal standard. Afterwards, 300 µL deionised water was added. The saponified mixture was extracted twice with either 500 µL hexane or 500 µL ethyl acetate depending on the expected polarity window of metabolites. Before extraction with ethyl acetate, ethanol from the saponification solution was evaporated in a rotary vacuum concentrator with a cooling trap. After addition of solvent, samples were vortexed well and centrifuged for 1 min at 10,000 × *g*. Both organic layers were pooled in a glass vial and concentrated *in vacuo* as for yeast samples. Derivatisation was performed for 1 h at 70 °C using 75 µL of 1:1 pyridine : (BSTFA+1% TMCS). Afterwards, the samples were diluted with 100 µL ethyl acetate and centrifuged for 10 min at 10,000 × g. The supernatant was subjected to GC-MS analysis as described below.

Semi-quantification of compounds from GC-MS data (shown in Fig. 6) was performed by integration of peak areas from compound-specific extracted ion chromatograms (EIC). For each compound, the EIC of the most abundant ion was chosen. Compounds **2**, **6**, **7**, **8**, **10**, and **11** were extracted more efficiently with hexane and therefore quantified from hexane extracts. Ethyl acetate extracts were used for integration of compounds **12**, **14**, and **15**. The following EICs were chosen for integration of each compound: 5α-cholestane (internal standard), *m/z* 217; **2**, *m/z* 129; **6**, *m/z* 363; **7**, *m/z* 157; **8**, *m/z* 157; **10**, *m/z* 295; **11**, *m/z* 293; **12**, *m/z* 183; **14**, *m/z* 383; **15**, *m/z* 183. Peak areas were normalised to internal standard and sample dry weight and converted into relative amounts with the highest mean value set to 100%.

### LC-MS sample preparation

LC-MS was used for the detection of oxidised withanolide pathway intermediates. After transient expression, *N. benthamiana* leaf samples were harvested in the same way as described for GC-MS analysis. 600 µL of ethyl acetate were added to the ground leaf powder and vortexed vigorously. The mixture was subsequently centrifuged at 14,000 × *g* for 15 minutes. The supernatant was transferred to a glass vial and concentrated *in vacuo.* Afterwards, the dried extracts were reconstituted in 600 µL ethanol and analysed by LC-MS as described below.

### General chemical methods and chemicals

GC-MS analyses were carried out on a Hewlett Packard HP6890N GC system connected to a mass selective detector 5973N with an OPTIMA 5 MS column (30 m × 0.25 mm i.d., 0.25 µm film, Macherey-Nagel, Düren, Germany). Helium was used as carrier gas at constant flow rate of 1.5 mL/min. Injection volume was 1 µL with a split of 1:5. For the quantification of 24-methyldesmosterol (**2**), the initial oven temperature was set to 100 °C followed by a ramp of 30 °C/min until 275 °C. Thereafter, the temperature was further raised to 290 °C with a ramp of 3 °C/min and held for 4 min, then raised to 300 °C with a ramp of 3 °C/min and held for 3.83 min. The total run time was 22 min. For all other measurements, the initial oven temperature was set to 100 °C followed by a ramp of 30 °C/min until 275 °C. The temperature was then further raised to 300 °C with a ramp of 3 °C/min and held for 15.83 min. The total run time was 30 min. Mass spectra were obtained with a scan range of *m*/*z* 43 to 800, with a solvent delay of 8 minutes after injection.

Analytical and semipreparative LC-MS measurements were performed on an Agilent Infinity II 1260 system consisting of a G7167A autosampler, G7116A column thermostat, G7111B quaternary pump, G7110B make-up pump, G7115A diode array detector, G1364F fraction collector, and G6125B single quadrupole mass spectrometer equipped with an ESI source (positive mode, 4000 V, 12 L min^−1^ drying gas, 350 °C gas temperature). Analytical LC-MS measurements were carried out using a Phenomenex Kinetex 2.6 µm C8 100 Å 4.6 x 150 mm column at 20 °C with 5 mM ammonium acetate in water (mobile phase A) and 5 mM ammonium acetate in MeOH (mobile phase B) and the following gradient: 0-5 min, 10-60% B; 5-12 min, 60-75% B; 12-15 min, 75-100% B; 15-21 min, 100% B; 21-21.1 min, 100-10% B; 21.10-24 min, 10% B. The flow rate was 0.8 mL/min. The injection volume was 5 µL and the total run was 24 min.

High resolution MS/MS measurements were carried out on a Vanquish LC (Thermo Fisher) using a Phenomenex Kinetex 2.6 µm C8 100 Å 4.6 x 150 mm column at 30 °C with a flow rate of 0.8 ml/min and 5 mM ammonium acetate in water (mobile phase A) and 5 mM ammonium acetate in MeOH (mobile phase B) as well as the following gradient: 0-5 min, 70-100% B; 5-5.1 min, 100-70% B; 5.1-7, 70% B. Mass spectra were obtained on an Orbitrap Q Exactive Plus mass spectrometer (Thermo Fisher) operated in PRM mode isolating the precursor with a *m*/*z* of 443 with an isolation window of 1 *m*/*z* followed by fragmentation at 10, 35 and 60 eV and detection of product ions at a resolution of 17,500). The AGC (automatic gain control) target and maximum injection time were set to 2e5 and 100 ms, respectively. The heated ESI (electrospray-ionisation) source was operated at 0 eV CID (collision-induced dissociation), sheath gas flow 60, auxiliary gas flow 17, sweep gas flow 4, spray voltage 3.5 kV, capillary temperature 288 °C, S-lens RF level 50.0 and aux gas heater 475 °C.

Flash chromatography was performed on an Biotage Isolera One with columns and solvents as described in the Supplementary Information. All methods and gradients for compound purification are described in detail in Supplementary Table 15-20.

NMR spectra of isolated compounds were recorded using Bruker spectrometers operating at 400, 500, and 600 MHz for ^1^H NMR, and at 100, 126, and 151 MHz for ^13^C NMR. Experiments were conducted at 298 K with the specified deuterated solvents. Chemical shifts (δ) were referenced relative to the residual solvent signal (CDCl_3_: δ_H_ = 7.26 ppm, δ_C_ = 77.16 ppm; C_5_D_5_N: δ_H_ = 8.74, 7.58, and 7.22 ppm, δ_C_ = 150.35, 135.91, and 123.87 ppm) and expressed in ppm, with coupling constants reported in Hz. Analysis was conducted using TopSpin (version 4.1.3) or MestReNova (version 14.3.1).

An authentic sample of 24-methylenecholesterol (**1**) was provided as a kind gift by Prof. Dr. Hans-Joachim Knölker (TU Dresden, Germany). 24-Methyldesmosterol (**2**) was synthesised according to a procedure by Edwards *et al.*^124^. Withanoside V aglycone (16) was prepared by enzymatic hydrolysis of withanoside V (PhytoLab) as described below.

Synthetic procedures used in the context of determining the C-22 stereochemistry of (22*R*)-ergosta-5,24-diene-3β,22-diol (**10**) are described in the Supplementary Information.

### Preparation of withanoside V aglycone (16)

Withanoside V aglycone (**16**) was prepared by enzymatic hydrolysis of withanoside V (PhytoLab) following a modified protocol by Matsuda *et al.*^87^. A solution of withanoside V (1.2 mg, 1.56 µmol) in sodium acetate buffer (0.4 mL, 0.2 M, pH 5) was treated with cellulase (from *T. reesei*, 4 µL, ≥ 700 U/g, ρ 1.1-1.3 g/mL). The mixture was stirred gently at 50 °C for 24 h and then subjected to an SPE column (Oasis PRiME HLB 1cc (30 mg)). After loading and washing with 5% MeOH (H_2_O:MeOH 95:5), steroids were eluted with 4 mL of 75% acetonitrile (H_2_O:acetonitrile 25:75). The eluate was concentrated under reduced pressure to yield a mixture of the desired aglycone of withanoside V (**16**) with remaining or partially deglycosylated starting material (1.0 mg). This mixture was then purified by semipreparative HPLC to yield 0.4 mg (0.90 µmol, 58%) of withanoside V aglycone (**16**). The identity of the isolated compound was confirmed by comparison of ^1^H NMR spectral data with literature^125^. Semipreparative purification was achieved with a Phenomenex Luna 5 µm C8 100 Å 250 x 100 mm column and a gradient of 5 mM ammonium acetate in water (A) and methanol (B) at a flow rate of 4 mL/min: 0-4 min, 80-100% B; 4-8 min, 100% B; 8-8.1 min, 100-80% B, 8.1-10 min, 80% B.

### Purification of pathway intermediates heterologously produced in *N. benthamiana*

For compound purification, *N. benthamiana* plants were vacuum-infiltrated in a 9.2 L ROTILABO desiccator (Carl Roth, Karlsruhe, Germany) connected to a MZ 2 NT membrane pump (Vacuubrand, Wertheim, Germany) at 30 mbar for 1 min as described previously^126^. 120 and 90 plants were used for the isolation of *Pp*CYP87G1 and *Pp*CYP88C7 (shunt) products, respectively. Leaves were harvested 7 days post infiltration and lyophilised for 3 days until the dry weight remained constant. The dried leaves were ground at room temperature into powder with a blender and extracted with hexane for 24ISO and *Pp*CYP87G1 products and with ethyl acetate for *Pp*CYP87G1+*Pp*CYP88C7 products. The crude extracts were concentrated *in vacuo* and purified by consecutive rounds of chromatography as described in Supplementary Table 15-20. NMR spectra of isolated compounds are shown in Supplementary Fig. 20-59.

## Data availability

Sequences of genes from the withanolide gene clusters shown here are provided as a Supplementary File in xlsx format. Raw sequencing reads are available from the European Nucleotide Archive: *Withania somnifera* genomic data (PRJEB64854) and corresponding RNA-seq data (ERR13615536). The assembled genome sequence and corresponding annotation of *W. somnifera* is available via LeoPARD (https://leopard.tu-braunschweig.de/receive/dbbs_mods_00077979). Re-annotations of the Physalis genome sequences are available via LeoPARD (https://leopard.tu-braunschweig.de/receive/dbbs_mods_00077980) Scripts and commands of the Circos plot generation are described in a GitHub repository (https://github.com/NancyChoudhary28/Withanolide_biosynthesis).

## Acknowledgements

We gratefully acknowledge financial support by the Deutsche Forschungsgemeinschaft (FR 3720/7-1 to J.F.; BP PU 718/2-1 to B.P.; INST 187/741-1 FUGG to C.-P.W.). In addition, work in the group of J.F. was supported by the SMART BIOTECS alliance between the Technische Universität Braunschweig and the Leibniz Universität Hannover, supported by the Ministry of Science and Culture (MWK) of Lower Saxony. J.P is funded by China Scholarship Council (202208320109). A.A. is funded by a scholarship from the Egyptian Ministry of Higher Education (call 2019/2020). P.H. received financial support provided by the European Research Council (ERC Consolidator Grant “RadCrossSyn”), Deutsche Forschungsgemeinschaft (Heisenberg-Program HE7133/8-1), and Boehringer Ingelheim Stiftung (Plus 3 Perspectives Programme). This work was supported by the BMBF-funded de.NBI Cloud within the German Network for Bioinformatics Infrastructure (de.NBI) (031A532B, 031A533A, 031A533B, 031A534A, 031A535A, 031A537A, 031A537B, 031A537C, 031A537D, 031A538A).

We thank Dr. Annika Stein and Stephan Rohrbach (Leibniz University Hannover, Germany) for preliminary work on this project and Dave Biedermann (Leibniz University Hannover, Germany) for synthesis of 24-methyldesmosterol. Prof. Dr. Hans-Joachim Knölker (TU Dresden, Germany) is gratefully acknowledged for providing authentic 24-methylenecholesterol as a kind gift. Umicore is acknowledged for a generous gift of metathesis catalysts. We thank Dr. David Nelson (University of Tennessee, USA) and the P450 nomenclature committee for naming of CYPs. We thank Thorsten Marschall (Botanical Garden of TU Braunschweig, Germany) and Annette Kaiser for excellent technical support in handling *Withania somnifera* propagation. We thank Dr. Gerald Dräger (Leibniz University Hannover, Germany) for excellent X-ray crystallography support. We also thank all members of the research group Plant Biotechnology and Bioinformatics (TU Braunschweig, Germany) for discussion and support. The *Nicotiana benthamiana* icon by Connor-Tansley is licensed under CC-BY 4.0 Unported.

## Author contributions

J.F. and B.P. conceived and supervised the project. S.E.H., N.C., K.M., J.P., A.A., A.B., B.P., and J.F. designed the experiments. N.C. and B.P. performed the assembly and annotation of the *W. somnifera* genome sequence. N.C. performed re-annotations of *Physalis* genome sequences, synteny analyses, phylogenetic analyses, and gene expression analyses. K.M. designed and conducted yeast metabolic engineering experiments. J.P. designed and conducted metabolic engineering in *N. benthamiana*. S.E.H, J.P., and A.B. performed transient expression in *N. benthamiana*. S.E.H. and J.P. purified compounds. A.A. performed NMR analysis. S.E.H., K.M., J.P., and A.B. generated and interpreted chromatographic and mass spectrometric data. S.E.H. optimised and performed deglycosylation of withanoside V. A.B. performed gene silencing in *N. benthamiana*. R.F. performed high molecular weight DNA extraction and nanopore sequencing. C.-P.W. and M.H. designed and performed MS/MS and high-resolution MS measurements. M.B. and P.H. designed synthetic routes. M.B. performed synthesis. S.E.H., N.C., K.M., J.P., A.A., A.B., B.P., and J.F. wrote the manuscript. All authors reviewed and approved the final manuscript.

## Materials & Correspondence

Boas Pucker

Jakob Franke

