## Supplementary Information for "Phylogenomics and metabolic engineering reveal a conserved gene cluster in Solanaceae plants for withanolide biosynthesis"

**Supplementary Methods**

### RNA extraction

RNA from *W. somnifera* root was extracted with the RNA plant and fungi kit (Macherey & Nagel). Homogenization of sample was performed using mortar, pestle and liquid nitrogen. On-column DNA digestion was performed according to the manufacturer's instructions. The extracted RNA quality and quantity was determined using Nanodrop measurement and agarose gel. Paired-end RNA-seq (2 x 150 nt) was conducted using Illumina NovaSeq 6000 (BMKGene).

### Identification of transposable elements

Transposable elements (TEs) were annotated using Extensive *de*-novo TE Annotator[1] (EDTA) v2.2.1 with the following parameters: --genome --cds --anno 1 --sensitive 1 --evaluate 1. EDTA uses various tools like HelitronScanner[2], LTR_FINDER[3,4], LTR_retriever[5], LTRharvest[6], TIR-Learner[7], Generic Repeat Finder[8] and TEsorter[9] to identify and classify novel TEs. Tandem repeats in the genome sequence were identified using Tandem Repeats Finder[10] (TRF) v4.10.0 using parameters 'Match=2, Mismatch=5, Delta=7, PM=80, PI=10, Minscore=5 and MaxPeriod=2000'.

### Virus-induced gene silencing in *N. benthamiana*

Virus-induced gene silencing (VIGS) in *N. benthamiana* was performed following a previously published protocol[11]. Target gene fragments of *NbDWF1* and *NbPDS* (500-600 bp) were designed using the SGN VIGS Tool[12]. Fragments were amplified from *N. benthamiana* leaf cDNA (primers in Supplementary Table 14) and cloned into pTRV2[11] by restriction digestion with EcoRI and XhoI. The resulting plasmid was transformed into *E. coli* Top10 cells, isolated and its sequence confirmed by Sanger sequencing. To enable *Agrobacterium*-mediated gene transfer into *N. benthamiana*, either pTRV1, pTRV2 or pTRV2-gene of interest (GOI) were transformed into *Agrobacterium tumefaciens* GV3101 by electroporation.

*Agrobacterium* strains for VIGS were grown and harvested as described in section "Transient expression in *Nicotiana benthamiana*". All strains were diluted in infiltration buffer to an $OD_{600}$ of 1.0 and strains containing pTRV1 and pTRV2/pTRV2-GOI were mixed in a 1:1 ratio. The mixture was then infiltrated into the leaves of three-week-old *N. benthamiana* plants. These were maintained in a phytochamber for two further weeks until symptoms of photobleaching were clearly visible in *PDS*-silenced control plants.

After these two weeks post inoculation, leaves that had newly emerged were infiltrated with *Agrobacterium tumefaciens* GV3101 containing the plasmid pHREAC-*Pper24ISO* for transient expression. After five days of transient expression in *DWF1*-silenced plants, leaves were harvested and analysed by GC-MS as described. Representative results are shown in Supplementary Fig. 11.

**Synthetic procedures**

To determine the C22 stereochemistry of (22*R*)-ergosta-5,24-diene-3β,22-diol (**10**), two pairs of structurally similar C22 epimers **S7-(22*S*/*R*)** and **S9-(22*S*/*R*)** that differ from **10** only in the oxidation of C26 were synthesised as following:

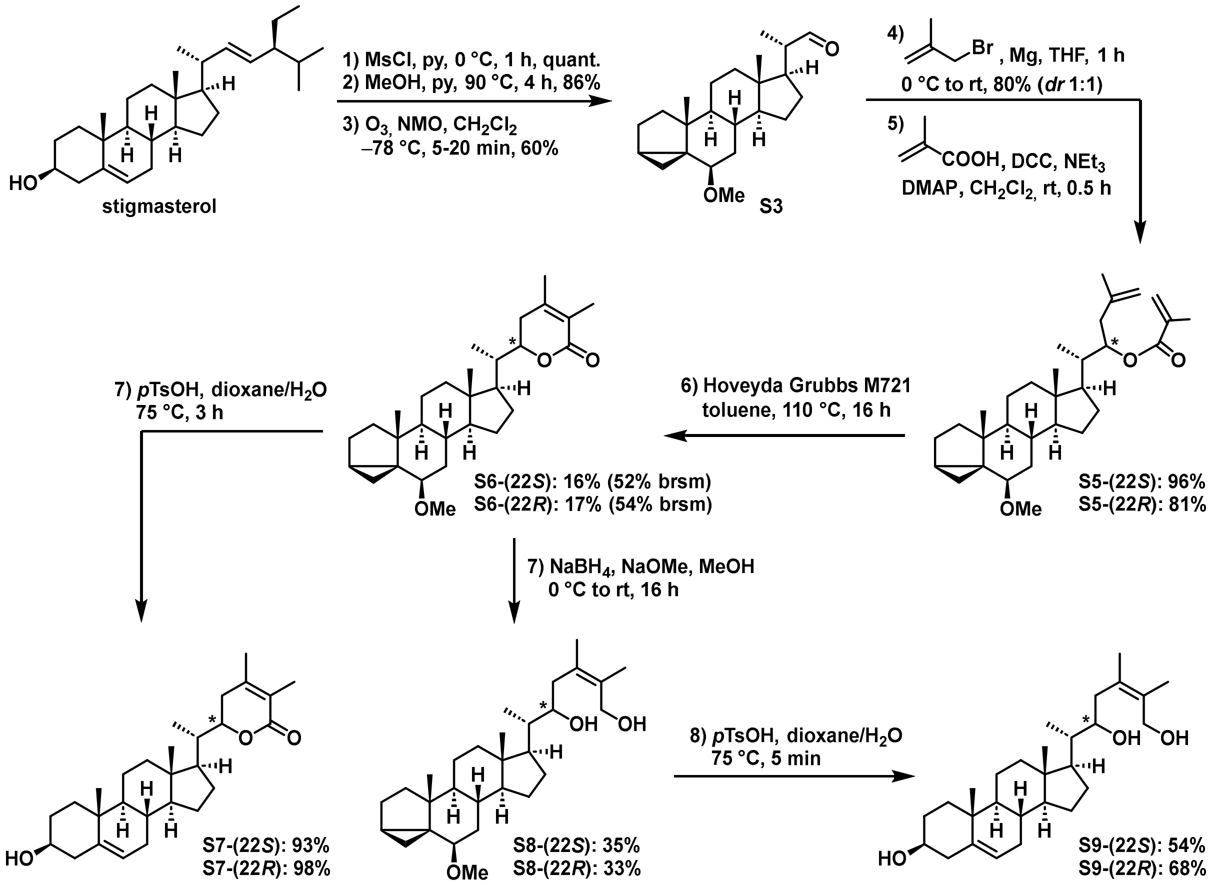


All reactions sensitive to moisture and/or air were carried out under an atmosphere of argon, using heat gun-dried glassware, and anhydrous solvents. Anhydrous dichloromethane, toluene, tetrahydrofuran and diethyl ether were taken from a M. Braun GmbH MB SPS-800 solvent purification system and were stored over 4 Å molecular sieves. Petroleum ether and tetrahydrofuran were purified by distillation. Stigmasterol was acquired from Thermo Scientific in >90% (GC) purity. Hoveyda Grubbs Catalyst® M721 was provided by Umicore. All other solvents (HPLC quality) and commercially available reagents were used without further purification unless otherwise stated.

Concentration under reduced pressure was performed by rotary evaporation at 40 °C and appropriate pressure, followed by exposure to vacuum ($10^{-3}$ mbar) at 25 °C.

Heating experiments were performed in an oil bath.

**Thin layer chromatography:** All reactions were monitored using pre-coated TLC sheets ALUGRAM® Xtra SIL G/$UV_{254}$ (0.2 mm, silica gel, $F_{254}$, aluminium-backed, MACHEREY-NAGEL) with detection by UV light (λ = 254 nm) and/or by staining with an aqueous solution of cerium sulphate and phosphomolybdic acid and heat.

**Flash column chromatography:** silica gel (0.04-0.063 mm, 240-400 mesh) obtained from MACHEREY-NAGEL. The applied petroleum ether fraction had a bp of 40-60 °C. The eluent is given as volume ratios (v/v).

**Automated flash column chromatography:** Combiflash NextGen 300+ used with RediSep® Silver columns by Teledyne ISCO or FlashPure Select columns by Büchi. The eluent is given as volume rations (v/v).

**NMR experiments** were recorded in $CDCl_3$ or methanol-$d_4$ purchased from deutero GmbH. The following NMR spectrometers were used: Bruker Avance I 400 MHz, Bruker Avance III HD 400 MHz, Bruker Avance III 400 MHz with Prodigy BBFO probe head, Bruker Avance III HD 500 MHz with TCI cryo probe head, Bruker Avance Neo 600 MHz with DUL cryo probe head, Bruker Avance IIIHD 600 MHz with Prodigy TCI probe head. Chemical shifts δ are reported in parts per million (ppm) using residual undeuterated solvent ($CDCl_3$: $\delta_H$ = 7.26 ppm, $\delta_C$ = 77.16 ppm; methanol-$d_4$: $\delta_H$ = 3.31 ppm, $\delta_C$ = 49.00 ppm) as an internal reference at 298 K. The given multiplicities are phenomenological; thus, the actual appearance of the signals is stated and not the theoretically expected one. The following abbreviations are used to designate multiplicities: s = singlet, d = doublet, t = triplet, q = quartet, quint = quintet, or combinations of these acronyms. Broad signals will be denoted by addition of the letter "b" e.g., broad singlet is written as "bs". In case no multiplicity could be identified, the chemical shift range of the signal is given (m = multiplet). Peak integrals are given as multiples of protons *Y*H, with *Y* being the number of protons belonging to the given signal. NMR spectra were processed using MestreNova version 15.0.1. NMR spectra of synthetic compounds are shown in Supplementary Fig. 60-89.

**Infrared spectroscopy:** Infrared (IR) spectra were measured on SHIMADZU FT-IR Affinity-1S spectrometer. Wavenumbers $\tilde{\nu}$ are given in $cm^{-1}$ and intensities are as follows: s = strong, m = medium, w = weak.

**High-resolution mass spectra (HRMS)** HR-ESI-MS analysis was performed using either a Waters QTof Premier coupled with a Waters Acquity HPLC incl. a TUV detector or on a Waters LCT Premier mass spectrometer coupled with a Waters Alliance 2695 HPLC. The sample was dissolved in a suitable solvent (approx. 1 mg/mL) and diluted in methanol (1:100). An aliquot of 5 µL was injected in constant flow of methanol and data were recorded in positive or negative ion mode. On the LCT Premier mass spectrometer no HPLC column was installed.

HR-EI-MS analyses were carried out on a Waters Micromass GCT Premier (70 eV) with direct sample inlet.

**Optical rotations** were measured on a Krüss optronic P3000 or a Perkin-Elmer 241 polarimeter at wavelength of $\lambda_{max}$ = 589 nm (sodium D-line) using 100 mm cells. The used solvent is specified for each substance and concentration (g/100 mL) is indicated.

**Melting points (m. p.)** were measured with a MPA100 melting point apparatus by Stanford Research Systems.

**Stigmasteryl mesylate (S1)**

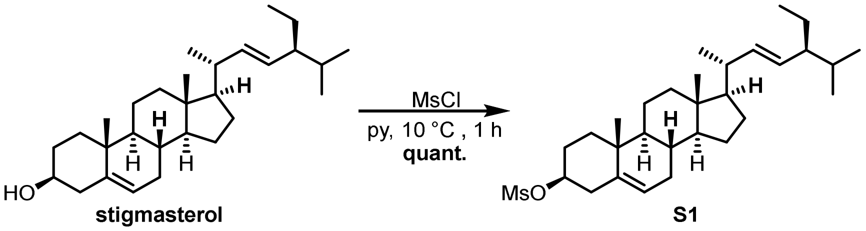


stigmasterol S1

To a stirred solution of stigmasterol (4.50 g, 10.9 mmol, 1.0 equiv.) in pyridine (100 mL) was added methanesulfonyl chloride (4.22 mL, 54.5 mmol, 5.0 equiv.) dropwise at 10 °C. After stirring at 10 °C for 1 h, the reaction mixture was poured into a mixture of ice and water (200 mL) under stirring. The precipitate was filtered off, washed with water (3× 100 mL), and dried under vacuum to give mesylate **S1** (5.35 g, 10.9 mmol, quant.) as a colourless powder, which was used in the next step without further purification.

**TLC** $R_f$ = 0.44 (PE/EtOAc 7:3, CAM).

**m. p.** 124–127 °C.

**$^1$H-NMR** (400 MHz, $CDCl_3$): δ [ppm] = 5.43–5.40 (m, 1H), 5.15 (dd, *J* = 15.2, 8.6 Hz, 1H), 5.02 (dd, *J* = 15.2, 8.6 Hz, 1H), 4.52 (dddd, *J* = 11.0, 11.0, 5.8, 4.7 Hz, 1H), 3.00 (s, 3H), 2.58–2.44 (m, 2H), 2.08–1.94 (m, 4H), 1.90 (dt, *J* = 13.4, 3.6 Hz, 1H), 1.85–1.65 (m, 2H), 1.60–1.36 (m, 8H), 1.32–1.05 (m, 6H), 1.04–1.00 (m, 7H contains 1.03 (d, *J* = 6.5 Hz, 3H), 1.02 (s, 3H)), 0.98–0.89 (m, 1H), 0.84 (d, *J* = 6.4 Hz, 3H), 0.81 (d, *J* = 7.5 Hz, 3H), 0.79 (d, *J* = 7.0 Hz, 3H), 0.70 (s, 3H).

**$^{13}$C-NMR** (101 MHz, $CDCl_3$): δ [ppm] = 138.8, 138.4, 129.5, 124.0, 82.2, 56.9, 56.1, 51.4, 50.1, 42.3, 40.6, 39.7, 39.3, 38.9, 37.1, 36.5, 32.02, 32.01, 31.9, 29.1, 29.0, 25.6, 24.5, 21.4, 21.2, 21.2, 19.3, 19.1, 12.4, 12.2.

**opt. act.** $[\alpha]_D^{24}$ = –43.9 (c=1.20, $CHCl_3$).

**HRMS** (ESI): *m/z* $[M+Na]^+$ calcd. for $[C_{30}H_{50}O_3NaS]^+$: 513.3373; found: 513.3358.

**IR** ṽ [$cm^{-1}$] = 2953 (m), 2932 (m), 2868 (w), 1317 (s), 1172 (s), 964 (m), 939 (s), 925 (s), 879 (m), 871 (s), 561 (m), 529 (s).

All characterization data was consistent with the data reported by Peracaula and co-workers[13].

***i*-Stigmasteryl methyl ether (S2)**

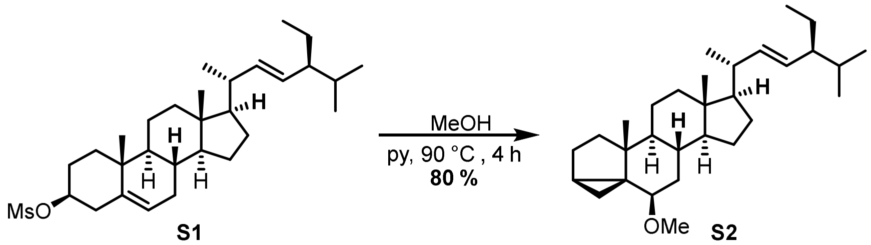


S1 S2

**S2** was synthesized according to the procedure published by Bräse and co-workers[14].

Mesylate **S1** (9.60 g, 19.6 mmol, 1.0 equiv.) was suspended in pyridine (32 mL) and methanol (160 mL). After stirring at 90 °C for 4 h, the solvents were removed under reduced pressure and diethyl ether (100 mL) was added. Remaining solids were separated by filtration and washed with diethyl ether (3x 150 mL). The solution was concentrated under reduced pressure, the resulting oil was adsorbed on silica gel and purified by automated flash column chromatography (silica gel, PE/EtOAc 100:0 to 90:10) yielding *i*-steroid **S2** (7.20 g, 16.9 mmol, 86%) and the isomeric *i*-steroid (1.00 g, 2.34 mmol, 12%) as clear, sticky resins which solidified in the freezer.

**TLC** $R_f$ = 0.55 (PE/EtOAc 9:1, CAM).

**$^1$H-NMR** (400 MHz, $CDCl_3$): $\delta$ [ppm] = 5.15 (dd, *J* = 15.2, 8.6 Hz, 1H), 5.01 (dd, *J* = 15.2, 8.6 Hz, 1H), 3.32 (s, 3H), 2.77 (t, *J* = 2.9 Hz, 1H), 2.10–2.01 (m, 1H), 1.97 (dt, *J* = 12.5, 3.4 Hz, 1H), 1.89 (dt, *J* = 13.5, 3.1 Hz, 1H), 1.81–1.64 (m, 3H), 1.60–1.47 (m, 5H), 1.46–1.36 (m, 3H), 1.30–1.00 (m, 13H, contains 1.02 (s, 3H), 1.01 (d, *J* = 7.0 Hz, 6H)), 0.91–0.78 (m, 12H, contains 0.85 (d, *J*=6.4 Hz, 3H)), 0.73 (s, 3H), 0.65 (dd, *J* = 5.0, 3.8 Hz, 1H), 0.43 (dd, *J* = 8.0, 5.1 Hz, 1H).

**$^{13}$C-NMR** (101 MHz, $CDCl_3$): δ [ppm] = 138.5, 129.4, 82.6, 56.8, 56.7, 56.3, 51.4, 48.2, 43.6, 42.8, 40.7, 40.4, 35.4, 35.2, 33.5, 32.0, 30.6, 29.2, 25.6, 25.1, 24.4, 22.9, 21.6, 21.4, 21.2, 19.4, 19.2, 13.2, 12.6, 12.4.

**opt. act.** $[\alpha]_D^{23}$ = +32.2 (c=1.20, $CHCl_3$).

**HRMS** (EI): *m/z* $[M]^{+\bullet}$ calcd. for $[C_{30}H_{50}O]^{+\bullet}$: 426.3856; found: 426.3848.

**IR** ṽ [$cm^{-1}$] = 2957 (m), 2945 (s), 2928 (s), 2895 (m), 2864 (s), 1454 (m), 1368 (w), 1089 (s), 1080 (s), 1015 (w), 968 (m), 615 (w).

All characterisation data was consistent with the data reported by Bräse and co-workers[14].

**(20*S*)-6*β*-Methoxy-3*α*,5-cyclopregnancarbaldehyde (S3)**

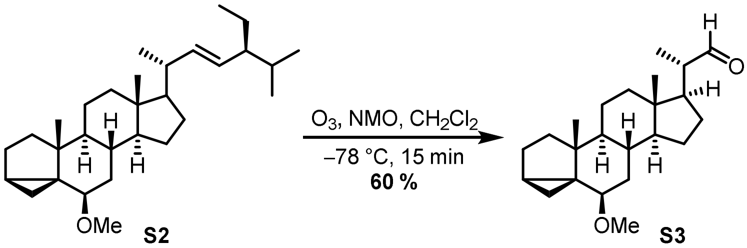


**S3** was synthesized according to the procedure published by Dussault and co-workers[15].

To a stirred solution of *i*-stigmasteryl methyl ether (**S2**, 2.09 g, 4.90 mmol, 1.0 equiv.) in $CH_2Cl_2$ (50 mL) was added *N*-methyl morpholine *N*-oxide (1.72 g, 14.7 mmol, 3.0 equiv.). At –78 °C ozone was bubbled slowly through the stirred solution and the reaction progress was checked by TLC (reaction time 10–15 min). Longer reaction times resulted in oxidation to the carboxylic acid. The reaction mixture was then sparged with $O_2$ for 2-3 min and allowed to warm to ambient temperature, dried over $MgSO_4$ and concentrated under reduced pressure. The resulting oil was adsorbed on silica gel and purified by automated flash column chromatography (silica gel, PE/EtOAc 100:0 to 90:10) yielding aldehyde **S3** as a colourless oil (1.01 g, 2.93 mmol, 60%) which solidified in the freezer.

**TLC** $R_f$ = 0.30 (PE/EtOAc 9:1, CAM).

**$^1$H-NMR** (400 MHz, $CDCl_3$): $\delta$ [ppm] = 9.57 (d, *J* = 3.2 Hz, 1H), 3.31 (s, 3H), 2.77 (t, *J* = 2.8 Hz, 1H), 2.36 (dqd, *J* = 10.0, 6.8, 3.2 Hz, 1H), 1.97–1.81 (m, 3H), 1.81–1.61 (m, 3H), 1.56–1.31 (m, 6H), 1.27–1.14 (m, 2H), 1.11 (d, *J* = 6.8 Hz, 3H), 1.10–1.04 (m, 2H), 1.02 (s, 3H), 0.93–0.78 (m, 3H), 0.76 (s, 3H), 0.68–0.61 (m, 1H), 0.43 (dd, *J* = 8.0, 5.1 Hz, 1H).

**$^{13}$C-NMR** (101 MHz, $CDCl_3$): δ [ppm] = 205.4, 82.4, 56.7, 55.9, 51.3, 49.7, 48.2, 43.5, 43.5, 40.1, 35.3, 35.2, 33.5, 30.6, 27.3, 25.1, 24.7, 22.8, 21.6, 19.4, 13.6, 13.2, 12.7.

**opt. act.** $[\alpha]_D^{23}$ = +31.5 (c=1.08, $CHCl_3$).

**HRMS** (EI): *m/z* $[M]^{+\bullet}$ calcd. for $[C_{23}H_{36}O_2]^{+\bullet}$: 344.2710; found: 344.2716.

**IR** ṽ [cm$^{-1}$] = 2947 (s), 2933 (s), 2913 (m), 2868 (m), 2845 (w), 1721 (s), 1454 (m), 1445 (w), 1382 (w), 1091 (s), 1018 (m), 999 (w), 970 (w), 943 (w), 613 (w).

All characterisation data was consistent with the data reported by Bräse and co-workers[14].

**(22*S*)-24-nor-6*β*-Methoxy-3*α*,5-cyclocholest-25-en-22-ol (S4-(22*S*))**

**(22*R*)-24-nor-6*β*-Methoxy-3*α*,5-cyclocholest-25-en-22-ol (S4-(22*R*))**

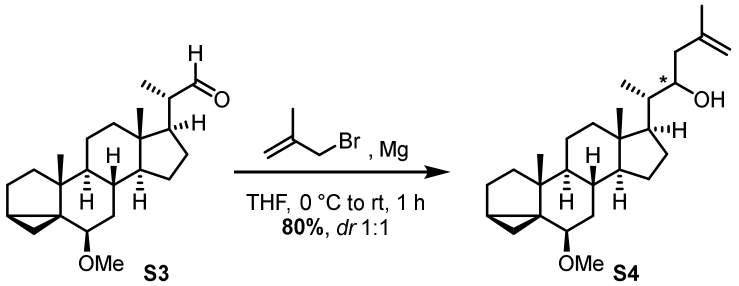


**S4** was synthesized according to the procedure published by Nemoto and co-workers[16].

To a stirred suspension of **S3** (1.41 g, 4.10 mmol, 1.0 equiv.) and Mg powder (499 mg, 20.5 mmol, 5.0 equiv.) in THF (4 mL) was added 3-bromo-2-methylprop-1-ene (1.65 mL, 16.4 mmol, 4.0 equiv.) at 0 °C. After stirring at this temperature for 1 h, the reaction mixture was diluted with $CH_2Cl_2$ (10 mL) and $NH_4Cl$ (sat., 10 mL) was added. The aqueous layer was extracted with $CH_2Cl_2$ (3× 20 mL) and the combined organic phases were dried over $MgSO_4$ and concentrated under reduced pressure. The residue was adsorbed on silica gel and the diastereomers were separated by automated flash column chromatography (silica gel, PE/EtOAc 100:0 to 90:10) to give alcohols **S4** (1.31 g, 3.28 mmol, 80%, *dr* 1:1) as clear oils.

**S4-(22*S*):**

**TLC** $R_f$ = 0.41 (PE/EtOAc 8:2, CAM).

**$^1$H-NMR** (400 MHz, $CDCl_3$): *δ* [ppm] = 4.82 (dt, *J* = 2.9, 1.4 Hz, 1H), 4.78–4.73 (m, 1H), 3.81 (ddd, *J* = 9.2, 4.4, 1.6 Hz, 1H), 3.31 (s, 3H), 2.75 (t, *J* = 2.9 Hz, 1H), 2.24 (dd, *J* = 13.6, 9.1 Hz, 1H), 2.06–1.81 (m, 4H), 1.78–1.57 (m, 6H, contains 1.73 (s, 3H)), 1.54–1.44 (m, 3H), 1.43–1.26 (m, 5H), 1.24–1.01 (m, 7H, contains 1.01 (s, 3H)), 0.91 (d, *J* = 6.6 Hz, 3H), 0.89–0.76 (m, 3H), 0.71 (s, 3H), 0.66–0.60 (m, 1H), 0.41 (dd, *J* = 8.1, 5.1 Hz, 1H).

**$^{13}$C-NMR** (101 MHz, $CDCl_3$): δ [ppm] = 143.3, 113.1, 82.5, 70.3, 56.7, 56.5, 53.0, 48.1, 44.2, 43.5, 42.8, 40.38, 40.36, 35.3, 35.2, 33.5, 30.7, 27.9, 25.1, 24.2, 22.9, 22.6, 21.6, 19.4, 13.2, 12.2, 11.9.

**opt. act.** $[\alpha]_D^{23}$ = +36.3 (c=0.91, $CHCl_3$).

**HRMS** (ESI): *m/z* $[M+Na]^+$ calcd. for $[C_{27}H_{44}O_2Na]^+$: 423.3234; found: 423.3226.

**S4-(22*R*):**

**TLC** $R_f$ = 0.36 (PE/EtOAc 8:2, CAM).

**$^1$H-NMR** (400 MHz, $CDCl_3$): *δ* [ppm] = 4.87–489 (m, 1H), 4.83–4.78 (m, 1H), 3.77 (ddd, *J* = 10.8, 3.5, 2.1 Hz, 1H), 3.32 (s, 3H), 2.76 (t, *J* = 2.9 Hz, 1H), 2.11–2.04 (m, 1H), 2.02–1.93 (m, 2H), 1.89 (dt, *J* = 13.7, 3.2 Hz, 1H), 1.83 (td, *J* = 7.0, 3.4 Hz, 1H), 1.79–1.68 (m, 6H, contains 1.76 (s, 3H)), 1.67–1.58 (m, 2H), 1.55–1.46 (m, 2H), 1.46–1.34 (m, 3H), 1.26–

0.99 (m, 8H, contains 1.02 (s, 3H)), 0.93 (d, $J$ = 6.7 Hz, 3H), 0.91–0.79 (m, 3H), 0.75 (s, 3H), 0.68–0.61 (m, 1H), 0.43 (dd, $J$ = 8.0, 5.1 Hz, 1H).

**$^{13}$C-NMR** (101 MHz, $CDCl_3$): δ [ppm] = 143.6, 113.5, 82.5, 69.8, 56.7, 56.3, 53.4, 48.2, 43.5, 43.3, 40.7, 40.4, 38.9, 35.4, 35.2, 33.5, 30.7, 27.6, 25.1, 24.4, 22.9, 22.3, 21.6, 19.4, 13.2, 12.5, 12.4.

**opt. act.** $[\alpha]_D^{23}$ = +44.7 (c=0.94, $CHCl_3$).

**HRMS** (ESI): *m/z* $[M+Na]^+$ calcd. for $[C_{27}H_{44}O_2Na]^+$: 423.3234; found: 423.3232.

**IR** ṽ [$cm^{-1}$] = 2934 (s), 2866 (m), 1456 (m), 1443 (w), 1373 (m), 1097 (s), 1082 (s), 1057 (s), 1016 (m), 999 (w), 968 (w), 891 (m), 883 (m), 613 (w), 534 (w).

**(22*S*)-24-nor-6*β*-Methoxy-3*α*,5-cyclocholest-25-ene-22-methacrylate (S5-(22*S*))**

**(22*R*)-24-nor-6*β*-Methoxy-3*α*,5-cyclocholest-25-ene-22-methacrylate (S5-(22*R*))**

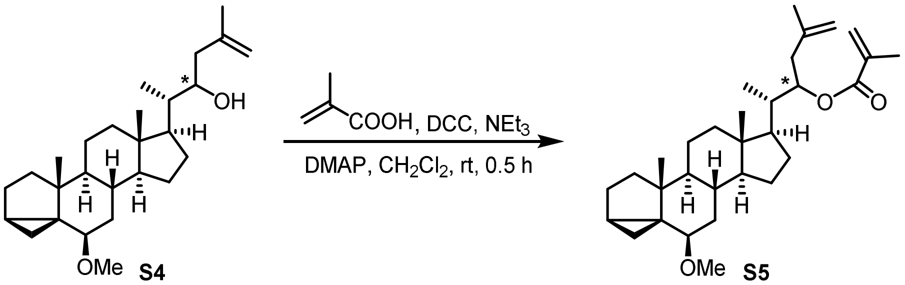


To a stirred solution of alcohol **S4-(22*S*)** (792 mg, 1.98 mmol, 1.0 equiv.) in $CH_2Cl_2$ (20 mL) were added methacrylic acid (0.84 mL, 9.89 mmol, 5.0 equiv.), *N,N'*-dicyclohexyl carbodiimide (2.04 g, 9.89 mmol, 5.0 equiv.), triethylamine (1.38 mL, 9.89 mmol, 5.0 equiv.) and *N,N*-dimethylpyridin-4-amine (24.2 mg, 0.20 mmol, 0.1 equiv.). After stirring for 30 min, the mixture was diluted with $CH_2Cl_2$ (20 mL), washed with HCl (aq., 10% *v/v*, 30 mL) and $NaHCO_3$ (sat. aq., 30 mL) and then was dried over $MgSO_4$. The solvent was removed under reduced pressure, the residue was adsorbed on silica gel and automated flash column chromatography (silica gel, PE/EtOAc 100:0 to 75:25) gave ester **S5-(22*S*)** (885 mg, 1.89 mmol, 96%) as a white amorphous solid.

**S5-(22*S*):**

**TLC** $R_f$ = 0.56 (PE/EtOAc 8:2, UV, CAM).

**$^1$H-NMR** (400 MHz, $CDCl_3$): $\delta$ [ppm] = 6.10–6.05 (m, 1H), 5.52–5.48 (m, 1H), 5.21 (td, $J$ = 7.1, 1.4 Hz, 1H), 4.79–4.73 (m, 1H), 4.73–4.68 (m, 1H), 3.31 (s, 3H), 2.76 (t, $J$ = 2.9 Hz, 1H), 2.37 (dd, $J$ = 13.5, 7.3 Hz, 1H), 2.19 (dd, $J$ = 13.5, 6.8 Hz, 1H), 2.03–1.92 (m, 5H, contains 1.93 (s, 3H)), 1.88 (dt, $J$ = 13.5, 3.1 Hz, 1H), 1.77 (s, 3H), 1.75–1.67 (m, 2H), 1.66–1.55 (m, 2H), 1.50 (td, $J$ = 12.1, 7.7 Hz, 2H), 1.43–1.09 (m, 5H), 1.09–0.98 (m, 8H, contains 1.05 (d, $J$=6.9 Hz, 1H), 1.01 (s, 3H)), 0.92–0.75 (m, 3H), 0.72 (s, 3H), 0.67–0.60 (m, 1H), 0.42 (dd, $J$ = 8.0, 5.1 Hz, 1H).

**$^{13}$C-NMR** (101 MHz, $CDCl_3$): δ [ppm] = 167.1, 142.0, 137.0, 125.1, 113.5, 82.5, 74.2, 56.7, 56.5, 53.0, 48.1, 43.5, 42.9, 40.9, 40.4, 39.0, 35.4, 35.1, 33.5, 30.7, 28.4, 25.1, 24.3, 22.9, 22.4, 21.6, 19.4, 18.6, 13.2, 12.9, 12.2.

**opt. act.** $[\alpha]_D^{23}$ = +21.9 (c=1.11, $CHCl_3$).

**HRMS** (ESI): *m/z* $[M+Na]^+$ calcd. for $[C_{31}H_{48}O_3Na]^+$: 491.3496; found: 491.3501.

**IR** ṽ [$cm^{-1}$] = 2951 (w), 2928 (w), 2909 (w), 2870 (w), 1709 (s), 1447 (w), 1315 (m), 1292 (m), 1159 (s), 1098 (s), 1084 (m), 1059 (w), 1015 (w), 970 (w), 941 (m), 887 (m), 810 (w).

**S5-(22*R*):** starting with 755 mg (1.88 mmol) of **S5-(22*R*)** resulted in a clear oil (714 mg, 1.52 mmol, 81%).

**TLC** $R_f$ = 0.56 (PE/EtOAc 8:2, UV, CAM).

**$^1$H-NMR** (400 MHz, $CDCl_3$): $\delta$ [ppm] = 6.08 (s, 1H), 5.50 (q, $J$ = 1.7 Hz, 1H), 5.16 (dt, $J$ = 10.9, 2.2 Hz, 1H), 4.73 (s, 1H), 4.69 (s, 1H), 3.32 (d, $J$ = 1.3 Hz, 3H), 2.77 (d, $J$ = 2.9 Hz, 1H), 2.24 (dd, $J$ = 14.4, 11.0 Hz, 1H), 2.12 (d, $J$ = 14.3 Hz, 1H), 1.99–1.80 (m, 7H, contains 1.92 (s, 3H))), 1.80–1.69 (m, 5H, contains 1.72 (s, 3H)), 1.68–1.57 (m, 2H), 1.56–1.45 (m, 2H), 1.45–1.35 (m, 2H), 1.27–1.11 (m, 3H), 1.10–1.03 (m, 2H), 1.01 (s, 3H), 0.96 (d, $J$ = 6.7 Hz, 3H), 0.92–0.77 (m, 3H), 0.73 (s, 3H), 0.64 (t, $J$ = 4.4 Hz, 1H), 0.43 (dd, $J$ = 8.1, 5.1 Hz, 1H).

**$^{13}$C-NMR** (101 MHz, $CDCl_3$): δ [ppm] = 167.0, 142.5, 137.0, 125.0, 112.7, 82.5, 74.7, 56.7, 56.3, 53.4, 48.2, 43.5, 43.3, 40.4, 39.3, 36.0, 35.4, 35.2, 33.5, 30.6, 27.3, 25.1, 24.4, 22.9, 22.5, 21.6, 19.4, 18.5, 13.2, 13.2, 12.5.

**opt. act.** $[\alpha]_D^{23}$ = +40.8 (c=0.98, $CHCl_3$).

**HRMS** (ESI): *m/z* $[M+Na]^+$ calcd. for $C_{31}H_{48}O_3Na$: 491.3505; found: 491.3501.

**(22*S*)-6*β*-Methoxy-3*α*,5-cycloergosta-24,25-diene-26,22-lactone (S6-(22*S*))**

**(22*R*)-6*β*-Methoxy-3*α*,5-cycloergosta-24,25-diene-26,22-lactone (S6-(22*R*))**

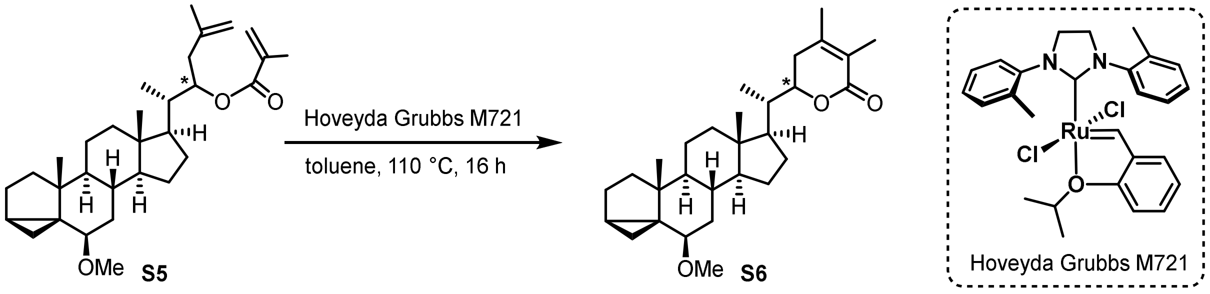


**S6** was synthesized according to the procedure published by Nemoto and co-workers[16].

A solution of ester **S5-(22*S*)** (694 mg, 1.48 mmol, 1.0 equiv.) and catalyst M721 (84.6 mg, 0.15 mmol, 0.1 equiv.) in toluene (148 mL) was stirred for 16 h at 110 °C. The solvent was removed under reduced pressure and automated flash column chromatography (silica gel, PE/EtOAc 100:0 to 65:35) gave lactone **S6-(22*S*)** (105 mg, 0.24 mmol, 16%, 52% brsm) as colourless crystalline solid.

**S6-(22*S*):**

**TLC** $R_f$ = 0.31 (PE/EtOAc 8:2, UV, CAM).

**m. p.** 128–130 °C.

**$^1$H-NMR** (400 MHz, $CDCl_3$): $\delta$ [ppm] = 4.40 (dd, $J$ = 12.4, 3.1 Hz, 1H), 3.31 (d, $J$ = 1.3 Hz, 3H), 2.76 (d, $J$ = 2.9 Hz, 1H), 2.63 (t, $J$ = 15.4 Hz, 1H), 1.97–1.81 (m, 9H, contains 1.92 (s, 3H), 1.87 (s, 3H)), 1.80–1.58 (m, 5H), 1.55–1.46 (m, 3H), 1.44–1.34 (m, 2H), 1.28–1.06 (m, 4H), 1.04–1.00 (m, 6H, contains 1.02 (d, $J$ = 9.8 Hz, 3H), 1.01 (s, 3H)), 0.94–0.77 (m, 4H), 0.71 (s, 3H), 0.64 (t, $J$ = 4.5 Hz, 1H), 0.42 (dd, $J$ = 8.1, 5.1 Hz, 1H).

**$^{13}$C-NMR** (101 MHz, $CDCl_3$): δ [ppm] = 167.3, 149.6, 122.0, 82.5, 78.4, 56.7, 56.3, 51.5, 48.0, 43.5, 42.8, 40.1, 39.6, 35.4, 35.1, 34.1, 33.5, 30.7, 27.9, 25.1, 24.2, 22.9, 21.6, 20.5, 19.4, 13.3, 13.2, 12.7, 12.3.

**opt. act.** $[\alpha]_D^{23}$ = +16.5 (c=1.15, $CHCl_3$).

**HRMS** (ESI): *m/z* $[M+Na]^+$ calcd. for $[C_{29}H_{44}O_3Na]^+$: 463.3183; found: 463.3170.

The absolute stereoconfiguration of **S6-(22*S*)** was determined by X-ray crystallography. A suitable crystal 0.40×0.29×0.04 mm$^3$ was selected and mounted on an 18 mm CryoLoop (20 micron, 0.2 - 0.3 mm, Hampton Research) on an XtaLAB AFC12 (RINC): Kappa single diffractometer. The crystal was kept at a steady temperature during data collection. The structure was solved with the ShelXT 2018/2 [17] structure solution program using the Intrinsic Phasing solution method and by using Olex2 [18] as the graphical interface. The model was refined with version 2019/3 of ShelXL 2019/3 [19] using Least Squares minimisation. X-ray crystallographic data is shown in Supplementary Table 21.

**S6-(22*R*):** starting with 675 mg (1.44 mmol) of **S5-(22*R*)** resulted in a yellowish amorphous solid (107 mg, 0.24 mmol, 17%, 54% brsm).

**TLC** $R_f$ = 0.30 (PE/EtOAc 8:2, UV, CAM).

**$^1$H-NMR** (400 MHz, $CDCl_3$): *δ* [ppm] = 4.37 (dt, *J* = 13.4, 3.5 Hz, 1H), 3.31 (d, *J* = 1.1 Hz, 3H), 2.76 (t, *J* = 3.0 Hz, 1H), 2.43 (t, *J* = 15.5 Hz, 1H), 2.04–1.83 (m, 10H, contains 1.93 (s, 3H), 1.87 (s, 3H)), 1.80–1.57 (m, 4H), 1.56–1.45 (m, 2H), 1.44–1.31 (m, 3H), 1.27–0.97 (m, 11H, contains 1.02 (s, 3H), 1.00 (s, 3H)), 0.93–0.76 (m, 3H), 0.74 (s, 3H), 0.65 (t, *J* = 4.5 Hz, 1H), 0.43 (dd, *J* = 8.1, 5.1 Hz, 1H).

**$^{13}$C-NMR** (101 MHz, $CDCl_3$): δ [ppm] = 167.2, 149.1, 122.1, 82.4, 78.5, 56.7, 56.3, 52.4, 48.2, 43.5, 43.3, 40.3, 39.0, 35.4, 35.2, 33.5, 30.7, 29.7, 27.6, 25.1, 24.4, 22.9, 21.6, 20.7, 19.4, 13.6, 13.2, 12.6, 12.2.

**opt. act.** $[\alpha]_D^{23}$ = +107.1 (c=1.13, $CHCl_3$).

**HRMS** (ESI): *m/z* $[M+Na]^+$ calcd. for $[C_{29}H_{44}O_3Na]^+$: 463.3183; found: 463.3184.

**IR** ṽ [$cm^{-1}$] = 2930 (m), 2864 (w), 1705 (s), 1395 (m), 1379 (m), 1315 (w), 1184 (m), 1146 (w), 1125 (s), 1098 (s), 1082 (m), 1016 (m), 995 (w), 970 (w), 762 (m).

**(22*S*)- 3*β*-Hydroxyergosta-5,24-diene-26,22-lactone (S7-(22*S*))**

**(22*R*)- 3*β*-Hydroxyergosta-5,24-diene-26,22-lactone (S7-(22*R*))**

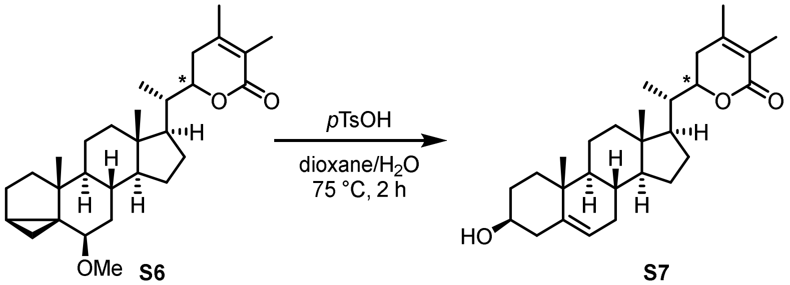


**S7** was synthesized according to the procedure published by Kang and co-workers[20].

Unsaturated lactone **S6-(22*S*)** (19.5 mg, 44.3 mmol, 1.0 equiv.) was dissolved in 1,4-dioxane (0.75 mL) and *p*TsOH (aq., 0.04 M, 0.15 mL) was added. After stirring at 75 °C for 2 h, the mixture was diluted with $CH_2Cl_2$ (5 mL) and $H_2O$ (2 mL). The aqueous layer was extracted with $CH_2Cl_2$ (3× 5 mL), the combined organic phases were dried over $MgSO_4$ and concentrated under reduced pressure. The residue was adsorbed on silica gel and column chromatography (silica gel, PE/EtOAc 50:50) gave alcohol **S7** (17.5 mg, 41.0 mmol, 93%) as white amorphous solid.

**S7-(22*S*):**

**TLC** $R_f$ = 0.20 (PE/EtOAc 6:4, UV, CAM).

**m. p.** 173–176 °C.

**$^1$H-NMR** (600 MHz, MeOD): $\delta$ [ppm] = 5.35 (dt, $J$ = 4.7, 2.0 Hz, 1H), 4.45 (ddd, $J$ = 13.3, 3.6, 1.3 Hz, 1H), 3.44–3.36 (m, 1H), 2.71–2.63 (m, 1H), 2.28–2.17 (m, 2H), 2.08–1.76 (m, 12H, contains 1.98 (s, 3H), 1.86 (s, 3H)), 1.70–1.44 (m, 8H), 1.43–1.34 (m, 1H), 1.26 (td, $J$ = 12.7, 4.3 Hz, 1H), 1.19–1.07 (m, 3H), 1.05 (d, $J$ = 6.3 Hz, 3H), 1.03 (s, 3H), 0.97 (ddd, $J$ = 12.2, 10.8, 5.0 Hz, 1H), 0.75 (s, 3H).

**$^{13}$C-NMR** (151 MHz, MeOD): δ [ppm] = 169.5, 153.3, 142.2, 122.4, 121.9, 80.2, 72.4, 58.0, 53.1, 51.7, 43.4, 43.0, 41.1, 40.5, 38.5, 37.7, 34.6, 33.3, 33.0, 32.3, 28.5, 25.2, 22.2, 20.3, 19.9, 13.7, 12.5, 12.1.

**opt. act.** $[\alpha]_D^{23}$ = –43.9 (c=1.20, $CHCl_3$).

**HRMS** (ESI): *m/z* $[M+Na]^+$ calcd. for $[C_{28}H_{42}O_3Na]^+$: 449.3026; found: 449.3032.

**IR** ṽ [cm$^{-1}$] = 2930 (m), 1694 (s), 1383 (w), 1146 (s), 1146 (s), 1092 (m), 1059 (s), 1022 (m), 1011 (m), 799 (m), 762 (w), 511 (w), 405 (w).

**S7-(22*R*):** starting with 28.3 mg (64.2 mmol) of **S6-(22*R*)** resulted in a white amorphous solid (26.8 mg, 62.8 mmol, 98%).

**TLC** $R_f$ = 0.20 (PE/EtOAc 6:4, UV, CAM).

**m. p.** 200–203 °C.

**$^1$H-NMR** (600 MHz, MeOD): $\delta$ [ppm] = 5.35 (dt, $J$ = 5.7, 1.9 Hz, 1H), 4.41 (ddd, $J$ = 13.4, 3.6, 3.6 Hz, 1H), 3.43–3.35 (m, 1H), 2.52–2.43 (m, 1H), 2.28–2.17 (m, 2H), 2.11 (dd, $J$ = 18.0, 3.5 Hz, 1H), 2.06 (dt, $J$ = 12.7, 3.5 Hz, 1H), 2.03–1.91 (m, 4H, contains 1.99 (s, 3H)), 1.91–1.86 (m, 1H), 1.86–1.83 (m, 3H), 1.82–1.74 (m, 2H), 1.70–1.63 (m, 1H), 1.62–1.45 (m, 5H), 1.44–1.36 (m, 1H), 1.28–1.13 (m, 3H), 1.12–1.05 (m, 2H), 1.05–1.03 (m, 7H, contains 1.04 (s, 3H), 1.03 (d, $J$ = 6.6 Hz, 3H)), 0.97 (ddd, $J$ = 12.0, 10.6, 5.1 Hz, 1H), 0.78 (s, 3H).

**$^{13}$C-NMR** (151 MHz, MeOD): δ [ppm] = 169.5, 152.7, 142.3, 122.3, 122.2, 80.2, 72.4, 57.7, 53.2, 51.7, 43.9, 43.0, 41.0, 40.4, 38.6, 37.7, 33.3, 33.0, 32.3, 30.4, 28.3, 25.4, 22.2, 20.4, 19.9, 13.8, 12.4, 12.1.

**opt. act.** $[\alpha]_D^{23}$ = +41.2 (c=0.98, $CHCl_3$).

**HRMS** (ESI): *m/z* $[M+Na]^+$ calcd. for $[C_{28}H_{42}O_3Na]^+$: 449.3026; found: 449.3015.

**(22*S*)- 6*β*-Methoxy-3*α*,5-cycloergost-24-ene-22,26-diol (S8-(22*S*))**

**(22*R*)- 6*β*-Methoxy-3*α*,5-cycloergost-24-ene-22,26-diol (S8-(22*R*))**

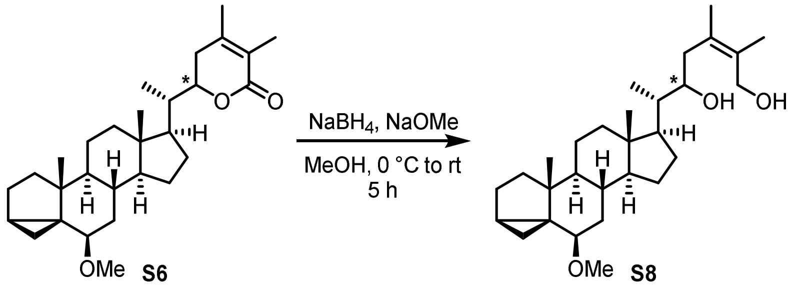


To a solution of unsaturated lactone **S6-(22*S*)** (29.0 mg, 66 mmol, 1.0 equiv.) in MeOH (0.15 mL) were added sodium borohydride (5.0 mg, 132 mmol, 2.0 equiv.) and sodium methoxide (1 mg, 20 mmol, 0.3 equiv.) at 0 °C. After 15 min the ice bath was removed and the solution was allowed to reach ambient temperature while being stirred for 5 h before adding EtOAc (5 mL) and hydrogen chloride (aq., 1 M, 5 mL). The aqueous layer was extracted with EtOAc (3× 10 mL), the combined organic phases were dried over $Na_2SO_4$ and concentrated under reduced pressure. The residue was adsorbed on silica gel and column chromatography (silica gel, PE/EtOAc 50:50) gave diol **S8-(22*S*)** (10.2 mg, 22.9 mmol, 35%) as colourless crystalline solid.

(Note: Reduction with lithium aluminium hydride resulted in unselective reduction yielding the saturated diol. Reduction with DIBAL through the intermediary lactol resulted in similar yields over two steps compared to sodium borohydride.)

**S8-(22*S*):**

**TLC** $R_f$ = 0.24 (PE/EtOAc 7:3, CAM).

**m. p.** 103–107 °C.

**$^1$H-NMR** (400 MHz, $CDCl_3$): *δ* [ppm] = 4.32 (d, *J* = 11.3 Hz, 1H), 3.73 (m, 2H), 3.32 (s, 3H), 2.80–2.69 (m, 2H), 2.56 (bs, 1H), 1.98 (dt, *J* = 12.5, 3.4 Hz, 1H), 1.94–1.83 (m, 2H), 1.78–1.66 (m, 9H, contains 1.80 (s, 3H), 1.70 (s, 3H)), 1.66–1.59 (m, 1H), 1.57–1.47 (m, 2H), 1.45–1.29 (m, 5H), 1.28–1.00 (m, 7H, contains 1.02 (s, 3H)), 0.98–0.93 (m, 3H), 0.92–0.77 (m, 3H), 0.73 (s, 3H), 0.64 (dd, *J* = 5.1, 3.8 Hz, 1H), 0.43 (dd, *J* = 8.0, 5.1 Hz, 1H).

**$^{13}$C-NMR** (101 MHz, $CDCl_3$): δ [ppm] = 131.5, 131.0, 82.5, 70.6, 63.6, 56.7, 56.5, 53.2, 48.1, 43.5, 42.9, 42.1, 40.4, 40.3, 35.3, 35.2, 33.5, 30.7, 27.9, 25.1, 24.3, 22.9, 21.6, 19.4, 19.2, 18.6, 13.2, 12.3, 12.3.

**opt. act.** $[\alpha]_D^{23}$ = +34.0 (c=0.99, $CHCl_3$).

**HRMS** (ESI): *m/z* $[M+Na]^+$ calcd. for $[C_{29}H_{48}O_3Na]^+$: 467.3496; found: 467.3509.

**IR** ṽ [$cm^{-1}$] = 2936 (m), 2905 (w), 2864 (m), 1456 (w), 1381 (w), 1098 (s), 1086 (m), 1011 (m), 1001 (s), 970 (w).

**S8-(22*R*):** starting with 80.0 mg (182 mmol) of **S6-(22*R*)** resulted in a white amorphous solid (27.1 mg, 61.2 mmol, 33%).

**TLC** $R_f$ = 0.18 (PE/EtOAc 7:3, CAM).

**m. p.** 200–202 °C.

**$^1$H-NMR** (400 MHz, $CDCl_3$): *δ* [ppm] = 4.35 (d, *J* = 11.2 Hz, 1H), 3.76 (m, 2H), 3.34 (s, 3H), 2.79 (t, *J* = 2.9 Hz, 1H), 2.52 (dd, *J* = 13.3, 10.9 Hz, 1H), 2.02 (dt, *J* = 12.6, 3.4 Hz, 1H), 1.92 (dt, *J* = 13.4, 3.1 Hz, 1H), 1.86–1.70 (m, 12H, contains 1.83 (s, 3H), 1.72 (s, 3H)), 1.70–1.61 (m, 1H), 1.55 (t, *J* = 6.3 Hz, 1H), 1.52 (t, *J* = 6.7 Hz, 1H), 1.48–1.35 (m, 3H), 1.28–1.07 (m, 5H), 1.04 (s, 3H), 1.00 (d, *J* = 6.7 Hz, 3H), 0.94–0.81 (m, 3H), 0.77 (s, 3H), 0.67 (dd, *J* = 5.1, 3.8 Hz, 1H), 0.45 (dd, *J* = 8.0, 5.1 Hz, 1H).

**$^{13}$C-NMR** (101 MHz, $CDCl_3$): δ [ppm] = 131.9, 130.7, 82.5, 70.1, 63.6, 56.7, 56.3, 53.4, 48.2, 43.5, 43.3, 42.5, 40.4, 35.4, 35.2, 34.4, 33.5, 30.7, 27.7, 25.1, 24.5, 22.9, 21.6, 19.4, 19.0, 18.7, 13.2, 12.4, 12.4.

**opt. act.** $[\alpha]_D^{23}$ = (c=, $CHCl_3$).

**HRMS** (ESI): *m/z* $[M+Na]^+$ calcd. for $[C_{29}H_{48}O_3Na]^+$: 467.3496; found: 467.3493.

**(22*S*)-Ergosta-5,24-diene-3β,22,26-triol (S9-(22*S*))**

**(22*R*)-Ergosta-5,24-diene-3β,22,26-triol (S9-(22*R*))**

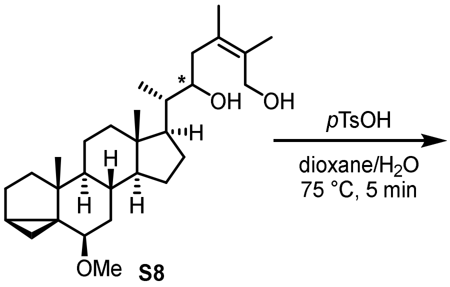


**S9** was synthesized according to a modified procedure published by Kang and co-workers[20].

Unsaturated diol **S8-(22*S*)** (14.8 mg, 33.3 mmol, 1.0 equiv.) was dissolved in 1,4-dioxane (0.45 mL) and 0.15 mL of a *p*TsOH stock solution (aq., 0.03 M, 0.15 equiv.) was added. After stirring at 75 °C for 5 min (longer reaction time resulted in decomposition), the mixture was diluted with EtOAc (5 mL) and $H_2O$ (2 mL). The aqueous layer was extracted with EtOAc (3× 5 mL), the combined organic phases were dried over $Na_2SO_4$ and concentrated under reduced pressure. The residue was adsorbed on silica gel and column chromatography (silica gel, PE/EtOAc 50:50 to 30:70) gave triol **S9-(22*S*)** (7.8 mg, 33.3 mmol, 54%) as white powder.

**S9-(22*S*):**

**TLC** $R_f$ = 0.21 (PE/EtOAc 1:1, CAM).

**m. p.** 164–167 °C.

**$^1$H-NMR** (600 MHz, MeOD): *δ* [ppm] = 5.34 (dd, *J* = 4.6, 2.4 Hz, 1H), 4.16 (d, *J* = 11.5 Hz, 1H), 3.93 (d, *J* = 11.5 Hz, 1H), 3.74 (ddd, *J* = 8.8, 4.6, 1.4 Hz, 1H), 3.43–3.33 (m, 1H), 2.53 (dd, *J* = 13.5, 8.7 Hz, 1H), 2.27–2.17 (m, 2H), 2.07–1.91 (m, 4H), 1.87 (dt, *J* = 13.3, 3.5 Hz, 1H), 1.83–1.75 (m, 4H, contains 1.76 (s, 3H)), 1.72 (s, 3H), 1.69–1.59 (m, 1H), 1.59–1.45 (m, 6H), 1.43–1.36 (m, 1H), 1.36–1.28 (m, 3H), 1.28–1.19 (m, 1H), 1.18–1.05 (m, 3H), 1.03 (s, 3H), 0.99–0.87 (m, 5H, contains 0.97 (d, *J* = 6.7 Hz, 3H)), 0.73 (s, 3H).

**$^{13}$C-NMR** (151 MHz, MeOD): δ [ppm] = 142.2, 131.6, 131.3, 122.4, 72.4, 72.2, 63.6, 58.2, 54.1, 51.7, 43.4, 43.0, 42.4, 41.2, 40.9, 38.6, 37.7, 33.3, 33.0, 32.3, 28.9, 25.3, 22.2, 19.9, 19.4, 17.6, 12.7, 12.2.

**opt. act.** $[\alpha]_D^{23}$ = –30.7 (c=0.25, MeOH).

**HRMS** (ESI): *m/z* $[M+Na]^+$ calcd. for $[C_{28}H_{46}O_3Na]^+$: 453.3339; found: 453.3326.

**IR** ṽ [$cm^{-1}$] = 2963 (m), 2926 (s), 2905 (m), 2879 (m), 2862 (m), 2851 (w), 1038 (s), 1022 (s), 1003 (s), 982 (s), 957 (m), 849 (w), 528 (m), 486 (w), 459 (w).

**S9-(22*R*):** starting with (8.5 mg, 19 mmol) of **S8-(22*R*)** resulted in a white powder (5.6 mg, 13 mmol, 68%).

**TLC** $R_f$ = 0.20 (PE/EtOAc 1:1, CAM).

**m. p.** 135–140 °C.

**$^1$H-NMR** (600 MHz, MeOD): *δ* [ppm] = δ 5.35 (dt, *J* = 5.5, 1.9 Hz, 1H), 4.19 (d, *J* = 11.4 Hz, 1H), 3.85 (d, *J* = 11.5 Hz, 1H), 3.72 (ddd, *J* = 10.8, 3.3, 1.9 Hz, 1H), 3.44–3.35 (m, 1H), 2.44 (dd, *J* = 13.7, 10.8 Hz, 1H), 2.28–2.17 (m, 2H), 2.06 (dt, *J* = 12.6, 3.5 Hz, 1H), 2.02–1.96 (m, 1H), 1.92–1.83 (m, 3H), 1.82–1.74 (m, 4H, contains 1.78 (s, 3)), 1.72 (s, 3H), 1.70–1.63 (m, 1H), 1.60–1.39 (m, 7H), 1.27–1.05 (m, 5H), 1.03 (s, 3H), 0.99 (d, *J* = 6.8 Hz, 3H), 0.98–0.85 (m, 1H), 0.76 (s, 3H).

**$^{13}$C-NMR** (151 MHz, MeOD): δ [ppm] = 142.3, 132.0, 131.6, 122.4, 72.4, 71.3, 63.7, 57.8, 54.5, 51.7, 43.8, 43.7, 43.0, 41.1, 38.6, 37.7, 35.0, 33.3, 33.0, 32.3, 28.7, 25.5, 22.2, 19.9, 19.1, 18.2, 12.8, 12.3.

**opt. act.** $[\alpha]_D^{23}$ = –9.6 (c=0.52, MeOH).

**HRMS** (ESI): *m/z* $[M+Na]^+$ calcd. for $[C_{28}H_{46}O_3Na]^+$: 453.3339; found: 453.3327.

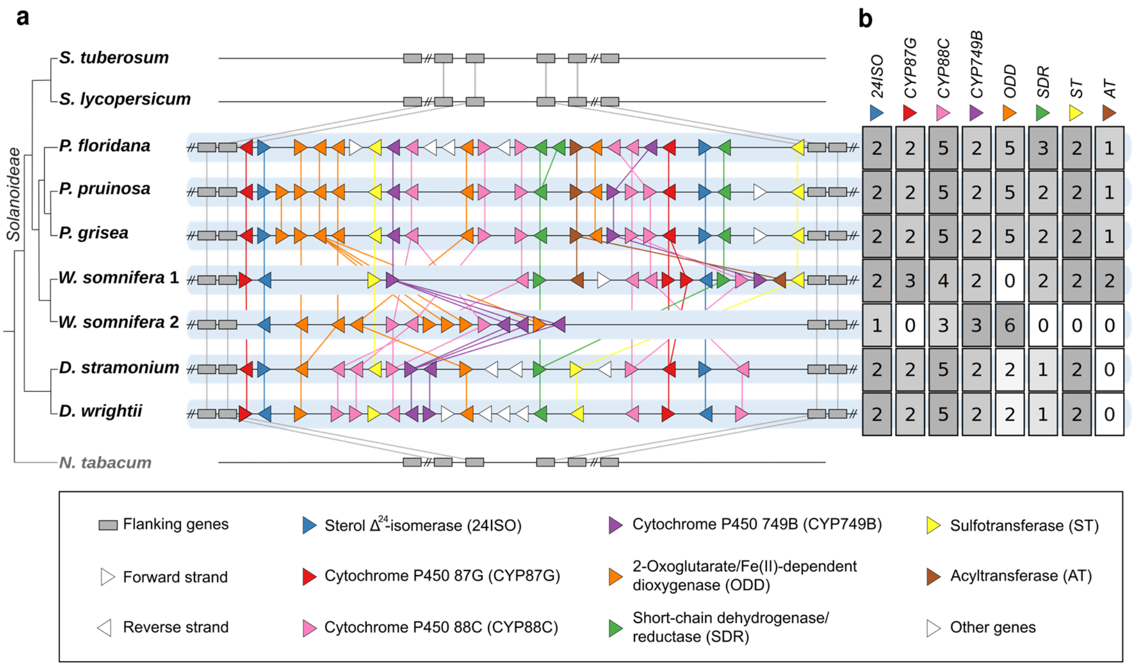


**Supplementary Fig. 1. Syntenic biosynthetic gene clusters containing *24ISO* in withanolide-producing Solanaceae plants, including the second, atypical gene cluster in the genome of *W. somnifera*.**

**a** Synteny plot. Gene lengths are unified for clarity. *N. tabacum* is included as an outgroup of the non-withanolide producing subfamily Solanoideae. **b** Heatmap summary of gene copy numbers of each gene family. Background colour is normalised to the highest number per column.

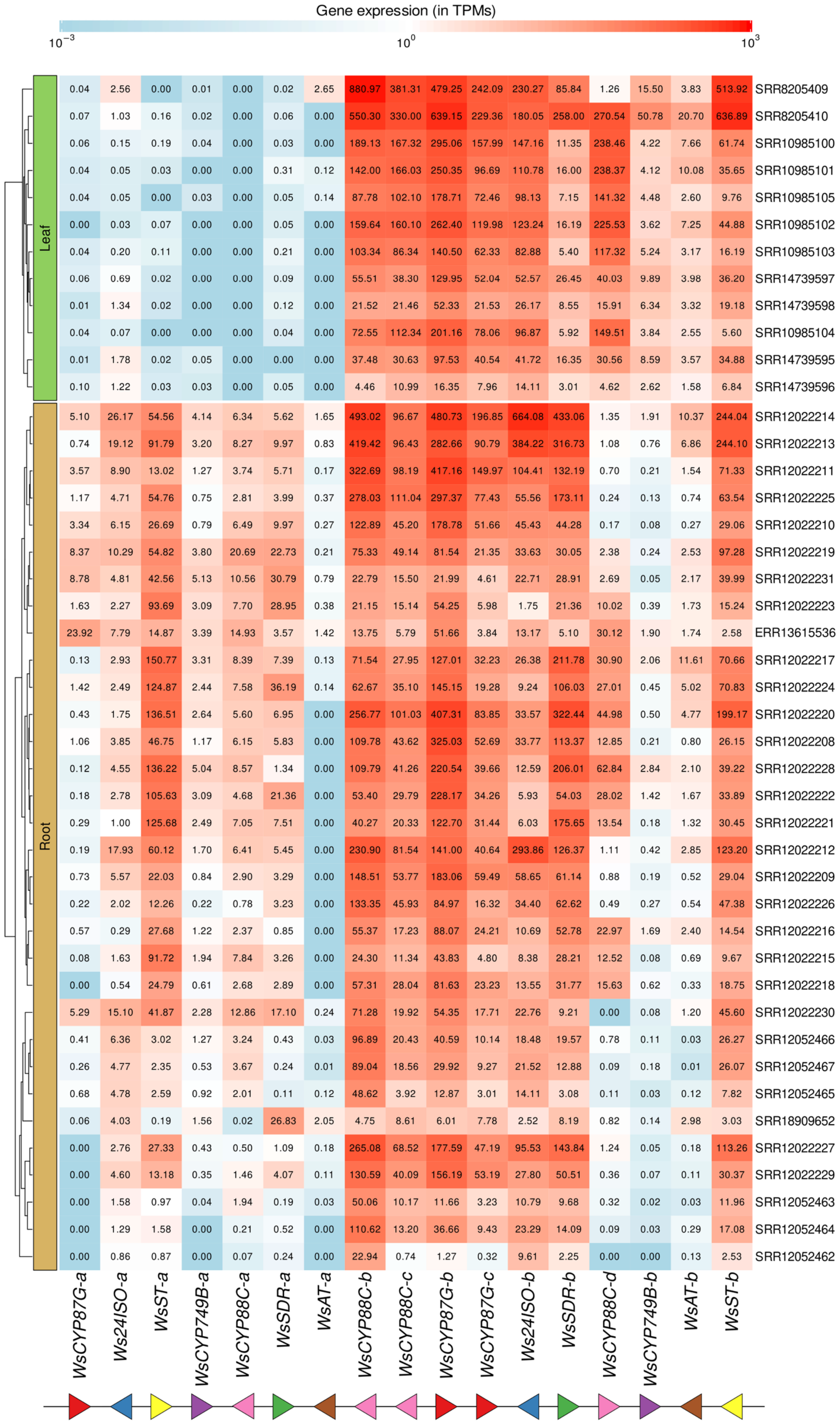


**Supplementary Fig. 2. Genes within the withanolide gene cluster of *W. somnifera* are co-expressed but fall into two distinct expression pattern groups.**

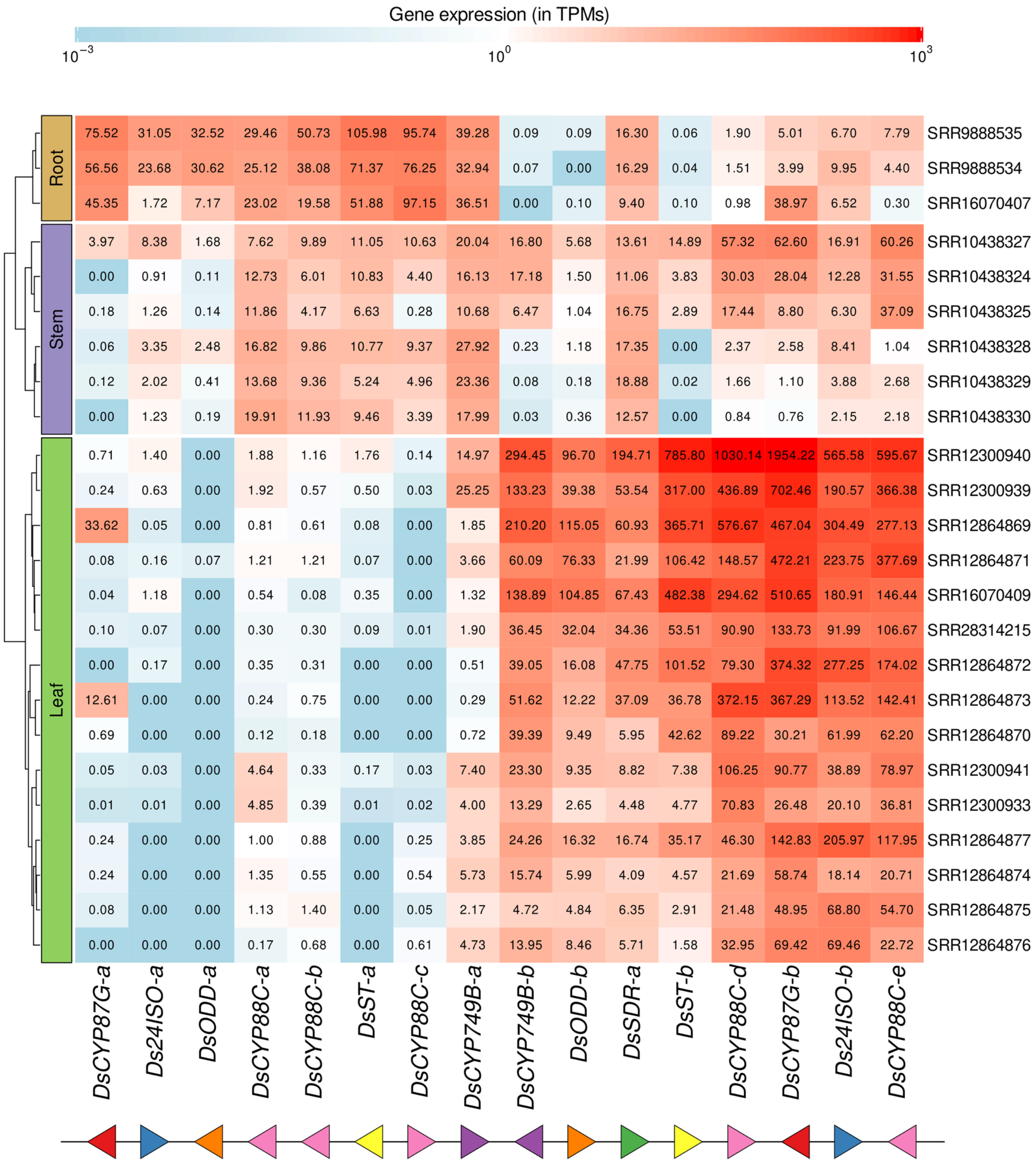


**Supplementary Fig. 3. Genes within the withanolide gene cluster of *D. stramonium* are co-expressed but fall into two distinct expression pattern groups.**

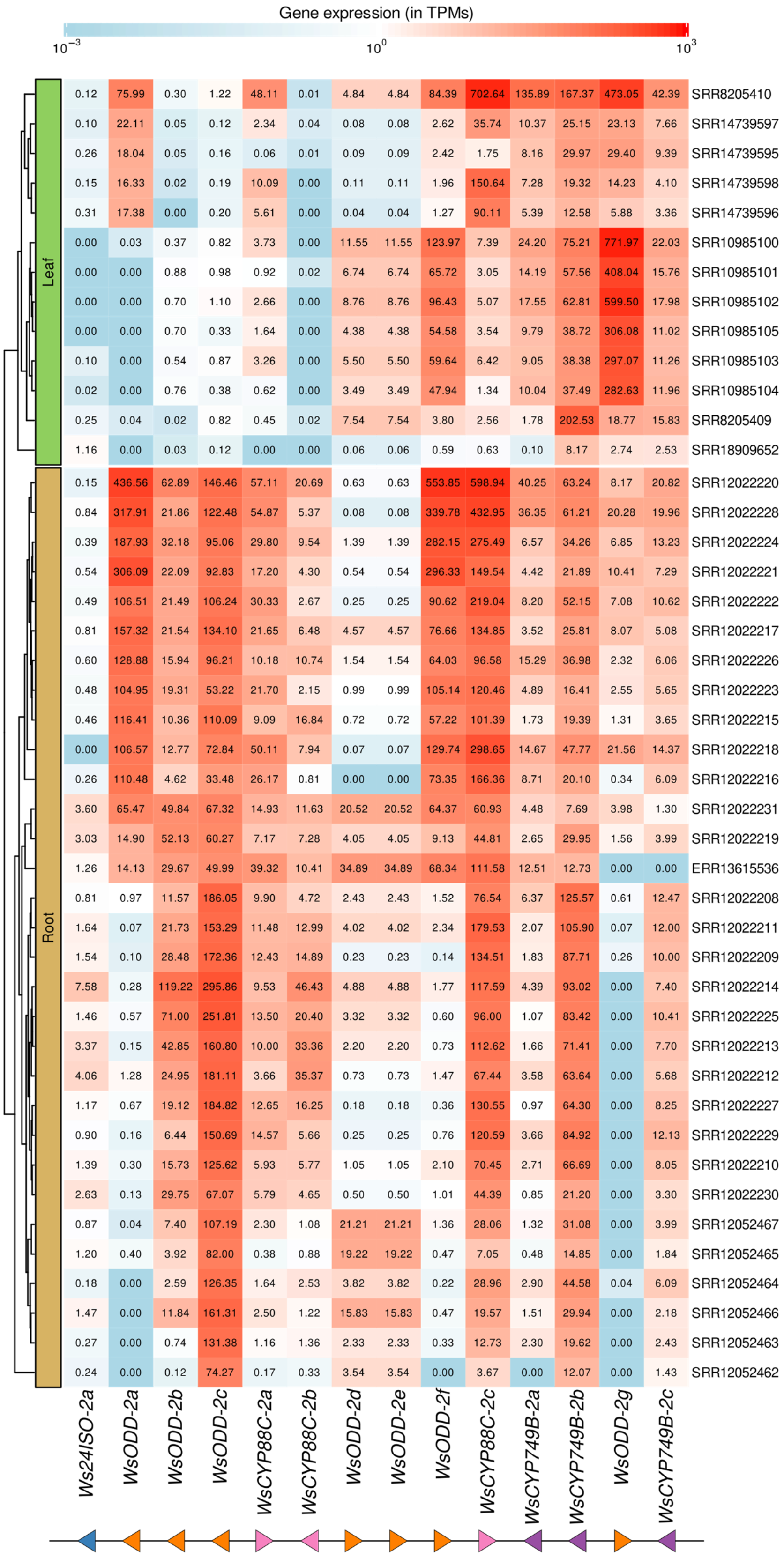


**Supplementary Fig. 4. Expression patterns of genes from the second *24ISO*-containing gene cluster in the genome of *W. somnifera*.**

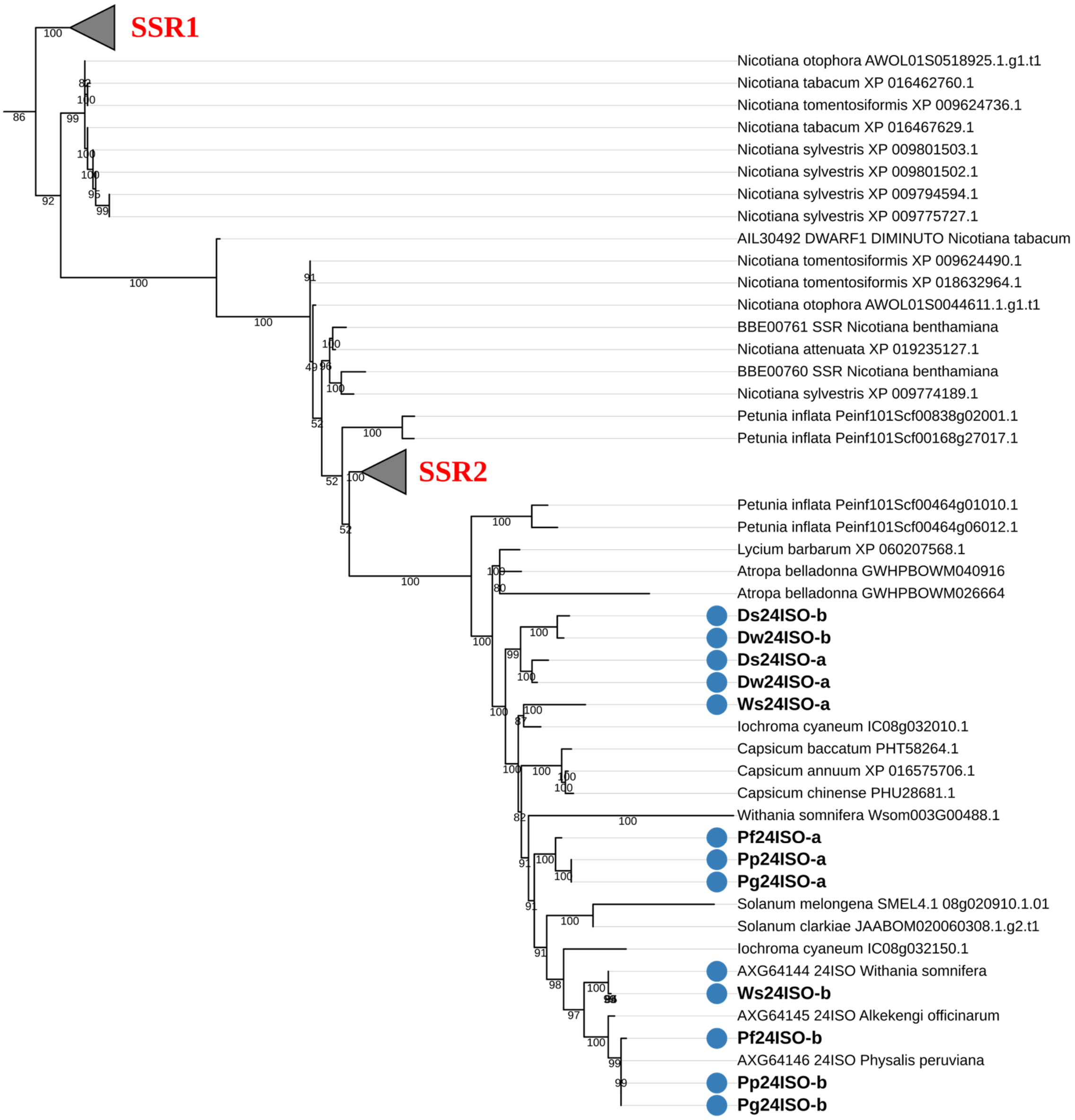


**Supplementary Fig. 5. Phylogenetic tree of $\Delta^{24}$ isomerase genes (*24ISO*) in withanolide gene clusters.**

**Gene sequences from known withanolide-producing Solanaceae species which are part of the gene cluster are marked by blue dots. Gene sequences identified in this study are highlighted in bold. Numbers above the nodes are the bootstrap values of maximum likelihood (ML) analysis based on 1000 replicates. Closely related SSR1 and SSR2 sequences are used as outgroup.**

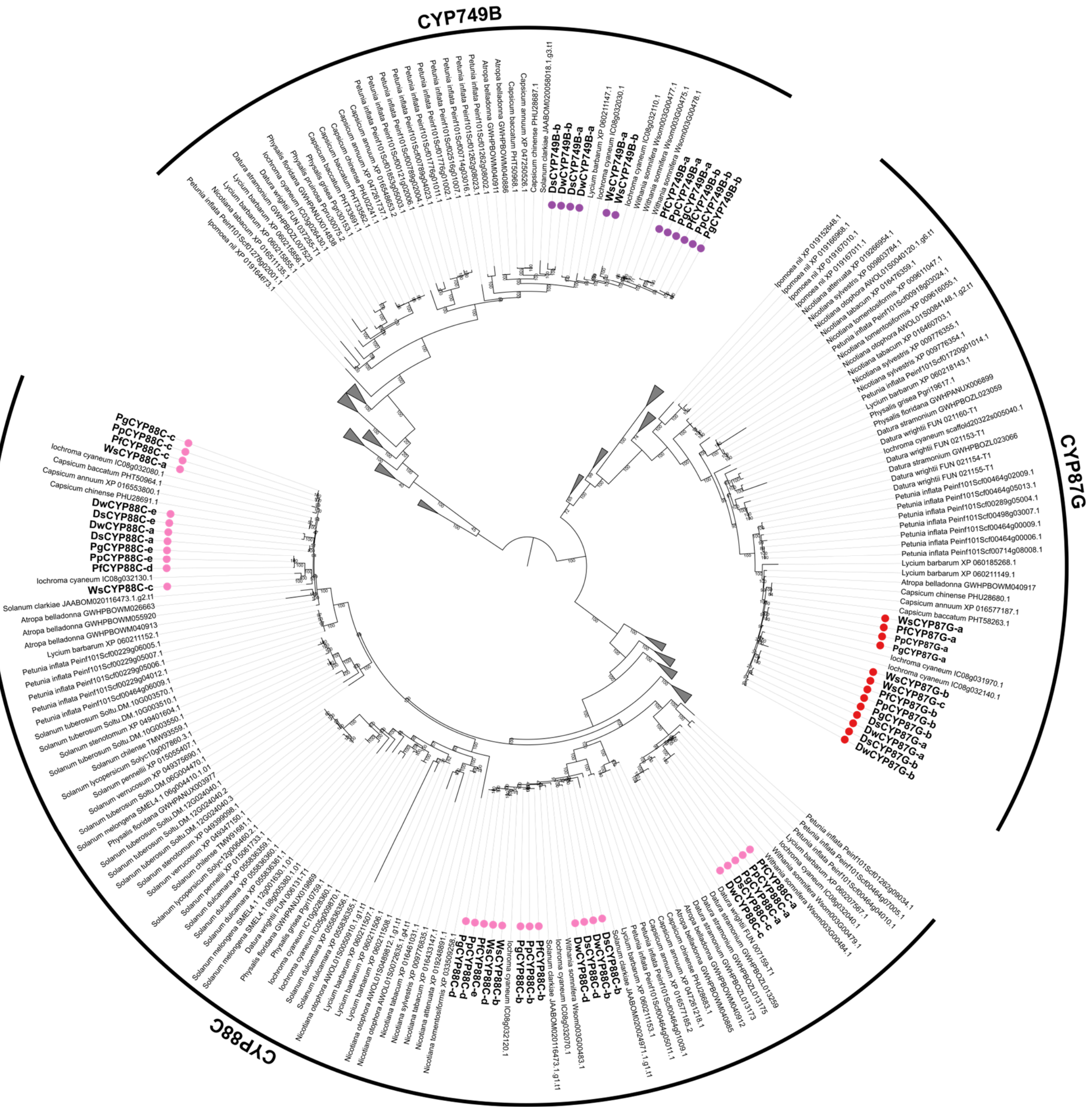


**Supplementary Fig. 6. Phylogenetic tree of cytochrome P450 monooxygenase genes (*CYP*) in withanolide gene clusters.**

**Gene sequences from known withanolide-producing Solanaceae species which are part of the gene cluster are marked by purple, red, and pink dots corresponding to CYP749B, CYP87G, and CYP88C sequences respectively. Gene sequences identified in this study are highlighted in bold. Numbers above the nodes are the bootstrap values of maximum likelihood (ML) analysis based on 1000 replicates.**

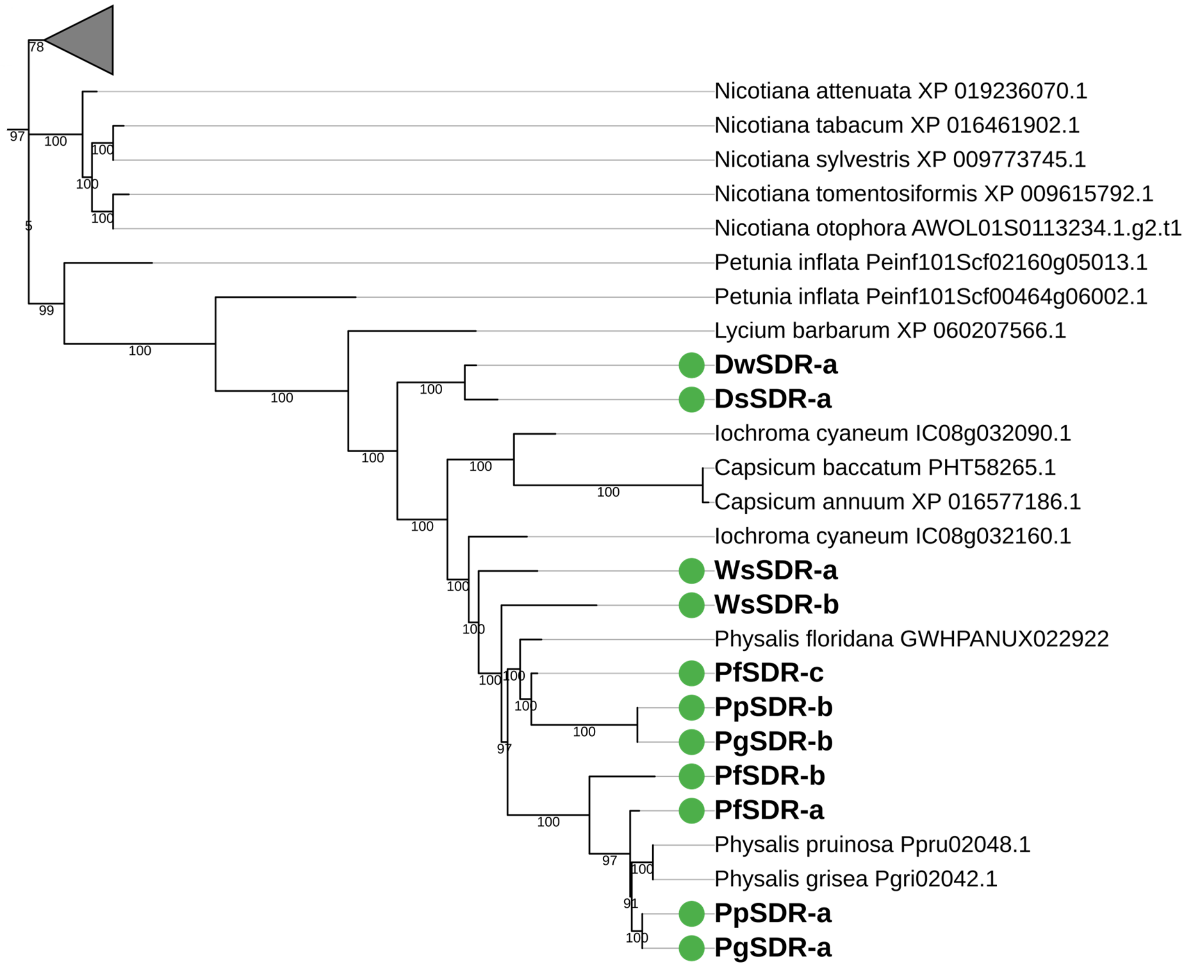


**Supplementary Fig. 7. Phylogenetic tree of short-chain dehydrogenase genes (*SDR*) in withanolide gene clusters.**

**Gene sequences from known withanolide-producing Solanaceae species which are part of the gene cluster are marked by green dots. Gene sequences identified in this study are highlighted in bold. Numbers above the nodes are the bootstrap values of maximum likelihood (ML) analysis based on 1000 replicates.**

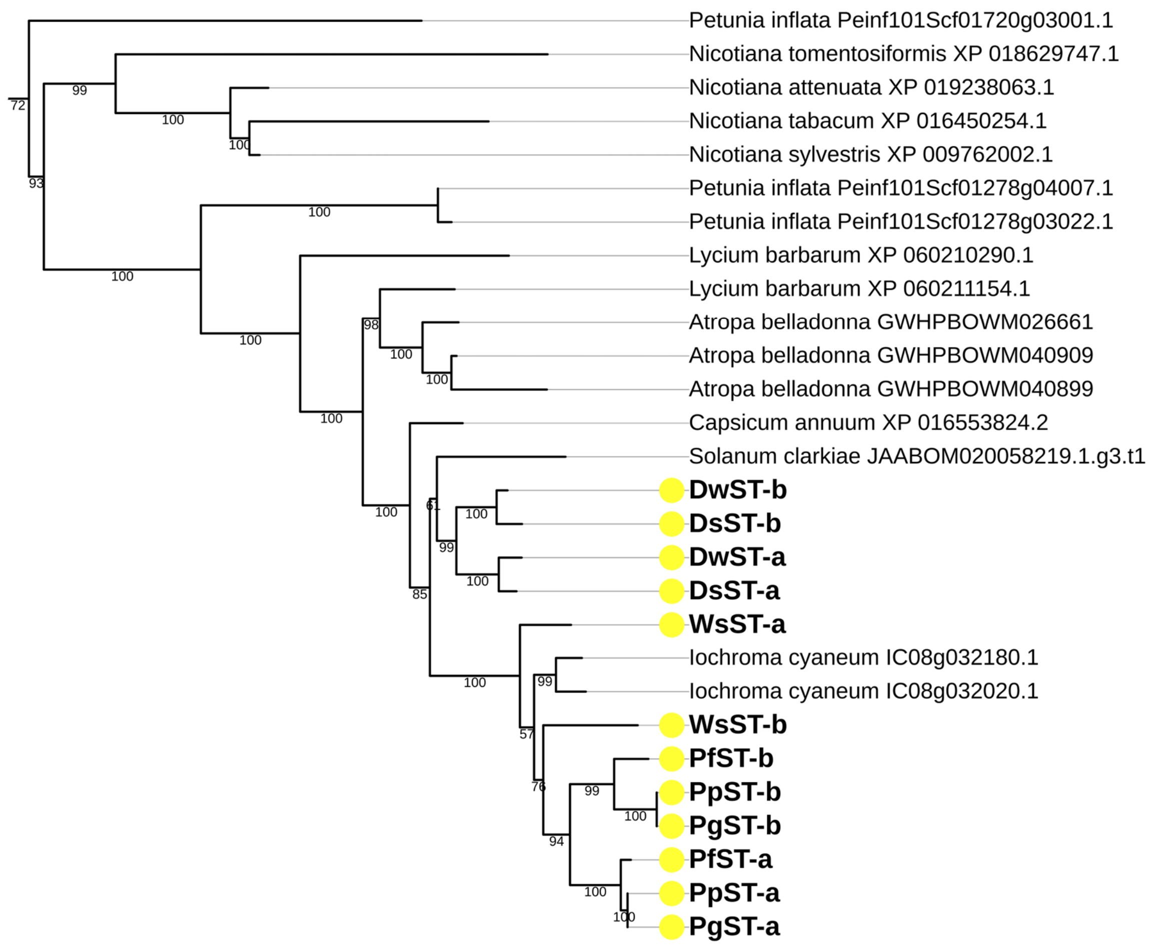


**Supplementary Fig. 8. Phylogenetic tree of sulfotransferase genes (*ST*) in withanolide gene clusters.**

**Gene sequences from known withanolide-producing Solanaceae species which are part of the gene cluster are marked by yellow dots. Gene sequences identified in this study are highlighted in bold. Numbers above the nodes are the bootstrap values of maximum likelihood (ML) analysis based on 1000 replicates.**

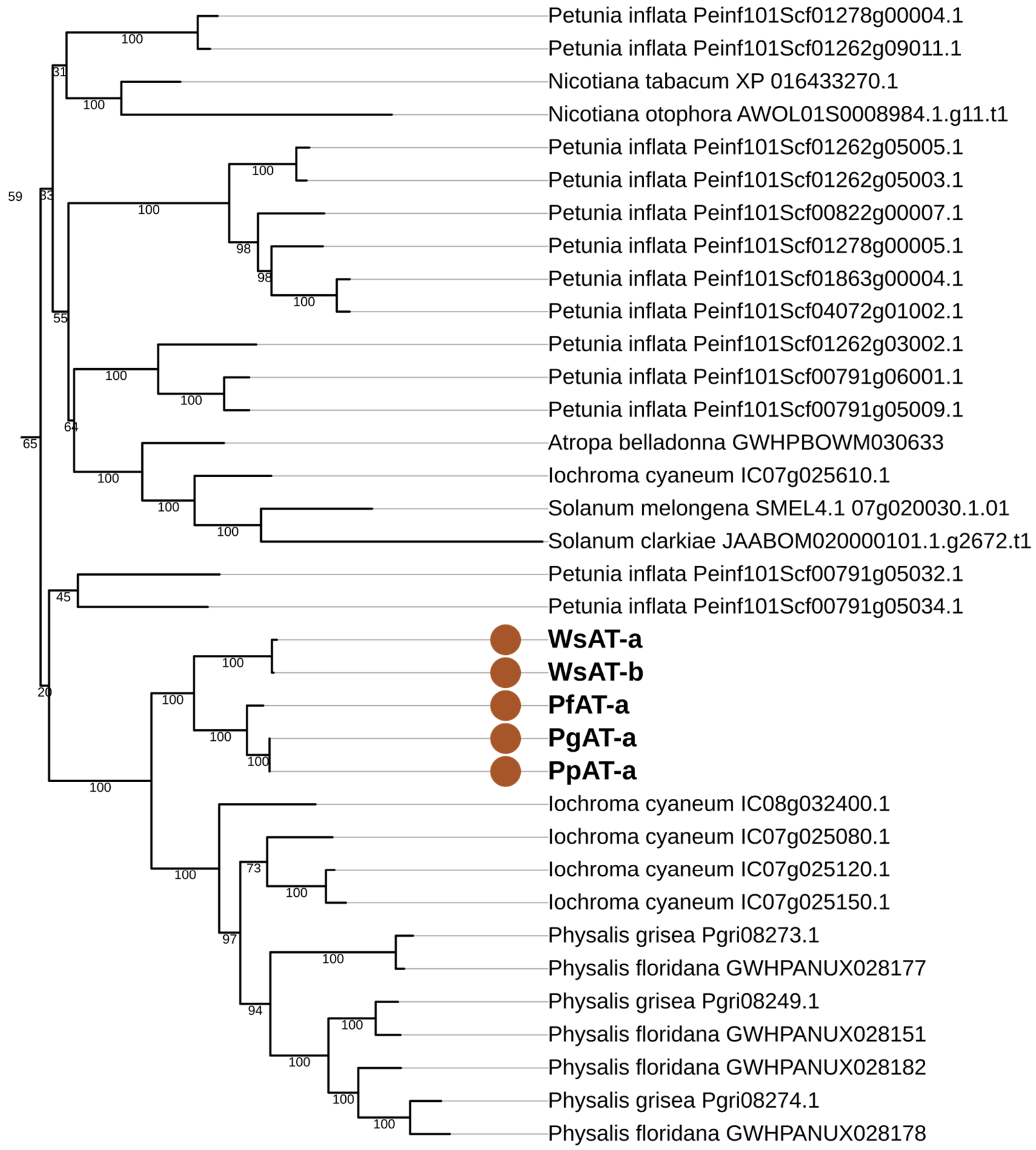


**Supplementary Fig. 9. Phylogenetic tree of acyltransferase genes (*AT*) in withanolide gene clusters.**

**Gene sequences from known withanolide-producing Solanaceae species which are part of the gene cluster are marked by brown dots. Gene sequences identified in this study are highlighted in bold. Numbers above the nodes are the bootstrap values of maximum likelihood (ML) analysis based on 1000 replicates.**

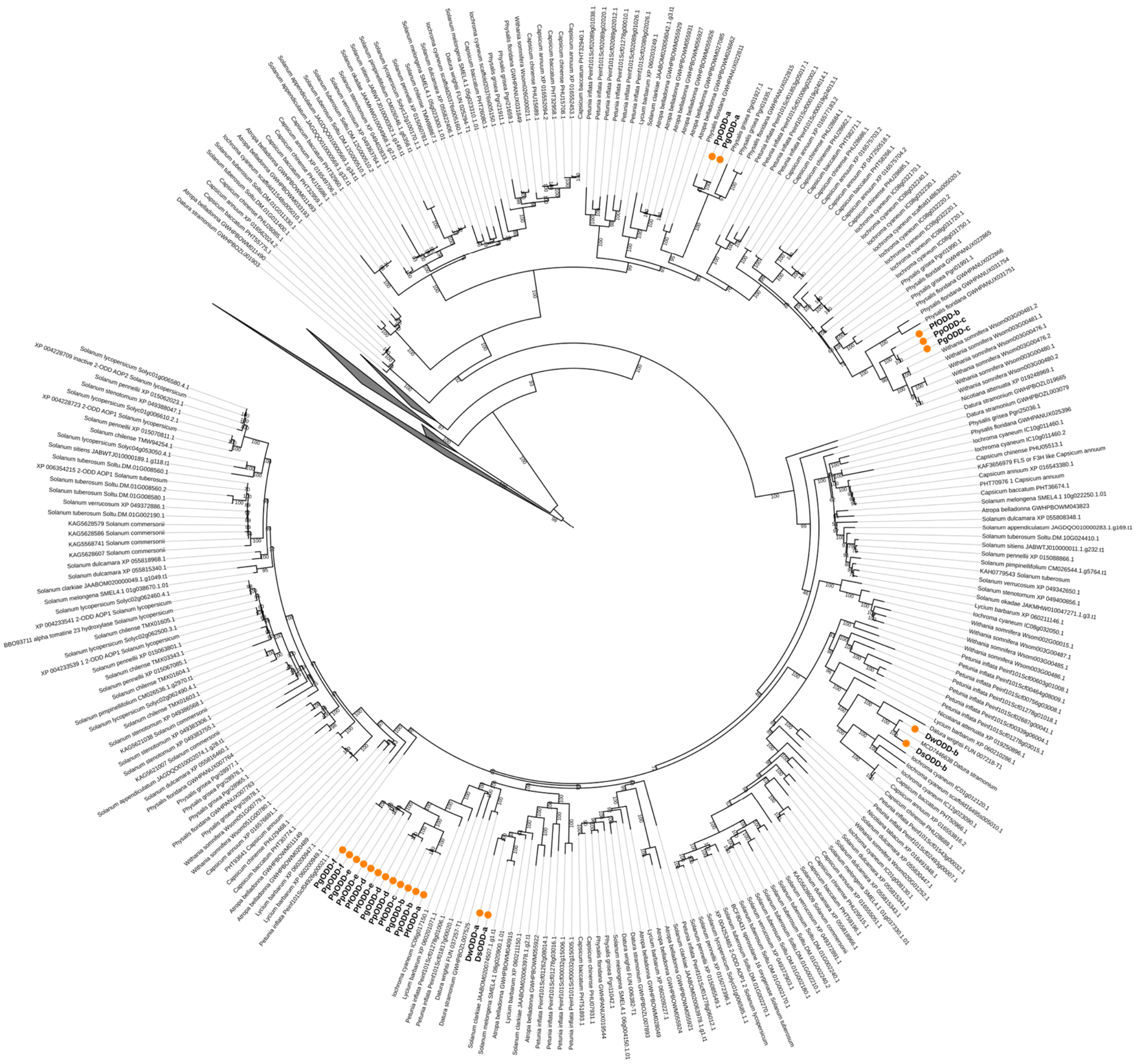

**Supplementary Fig. 10. Phylogenetic tree of 2-oxoglutarate-dependent dioxygenase genes (*ODD*) in withanolide gene clusters.**

**Gene sequences from known withanolide-producing Solanaceae species which are part of the gene cluster are marked by orange dots. Gene sequences identified in this study are highlighted in bold. Numbers above the nodes are the bootstrap values of maximum likelihood (ML) analysis based on 1000 replicates.**

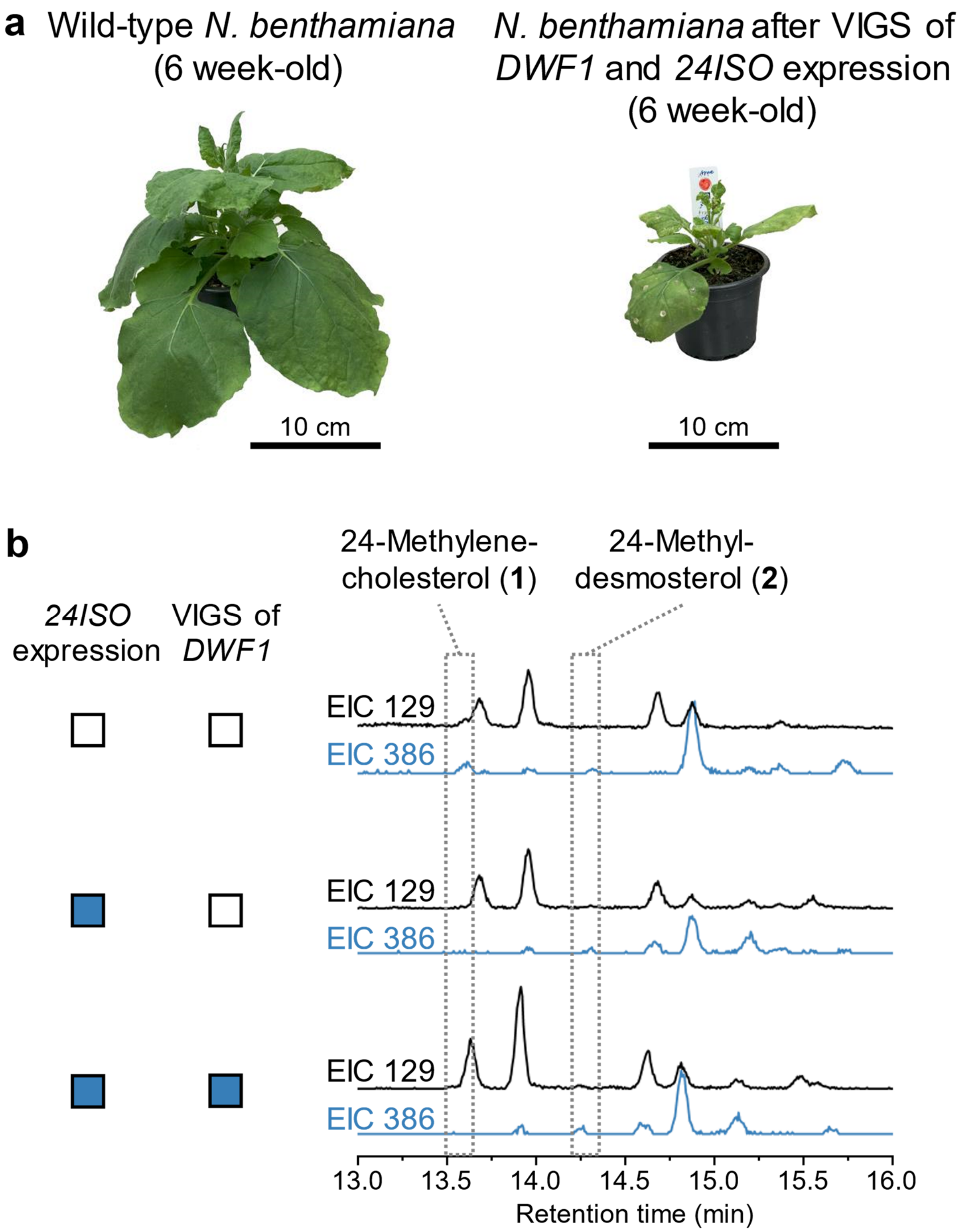


**Supplementary Fig. 11. Combining virus-induced gene silencing of *DWF1* with transient expression of *24ISO* leads to production of trace amounts of 24-methyldesmosterol (2) but comes with a severe dwarf phenotype.**

**a** Comparison of a typical *N. benthamiana* wild-type plant and a representative plant after virus-induced gene silencing of *DWF1* and transient expression of *24ISO*.

**b** Extracted ion chromatograms (EIC) from GC-MS analysis of different combinations of *24ISO* expression and *DWF1* silencing. *m*/*z* 129 is a general A ring fragment of typical steroids, whereas *m*/*z* 386 is specific for 24-methylenecholesterol (**1**) and 24-methyldesmosterol (**2**) (Supplementary Fig. 13-15 and Supplementary Table 3).

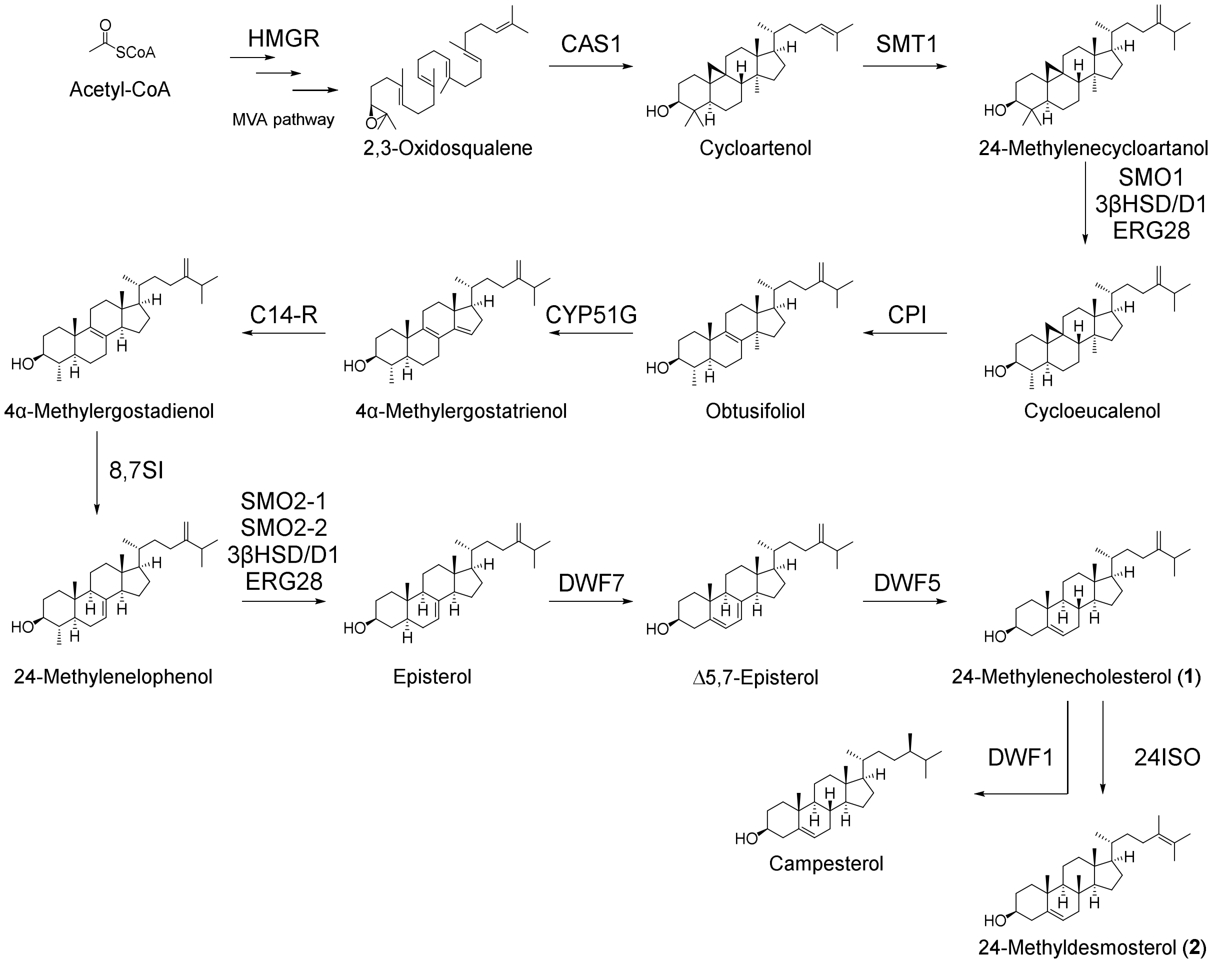


**Supplementary Fig. 12. Full phytosterol pathway transiently overproduced in this work.**

**HMGR:** 3-hydroxy-3-methyl-glutaryl-coenzyme A reductase; **CAS1:** cycloartenol synthase 1; **SMT1:** sterol C-24 methyltransferase; **SMO1 and SMO2-1/2:** sterol C-4 methyl oxidase 1 and 2-1/2-2; **3βHSD/D1:** 3β-hydroxysteroid dehydrogenase/C-4 decarboxylase 1; **ERG28:** ergosterol biosynthetic protein 28; **CPI:** cyclopropylsterol isomerase; **CYP51G:** sterol C-14 demethylase; **C14-R:** sterol C-14 reductase; **8,7SI:** sterol 8,7 isomerase; **DWF7 (DWARF7/STE1):** Δ7 sterol C-5 desaturase; **DWF5 (DWARF5 / 7RED):** sterol Δ7 reductase; **DWF1 (DWARF1/DIM):** sterol C-24 reductase; **24ISO:** sterol Δ24 isomerase.

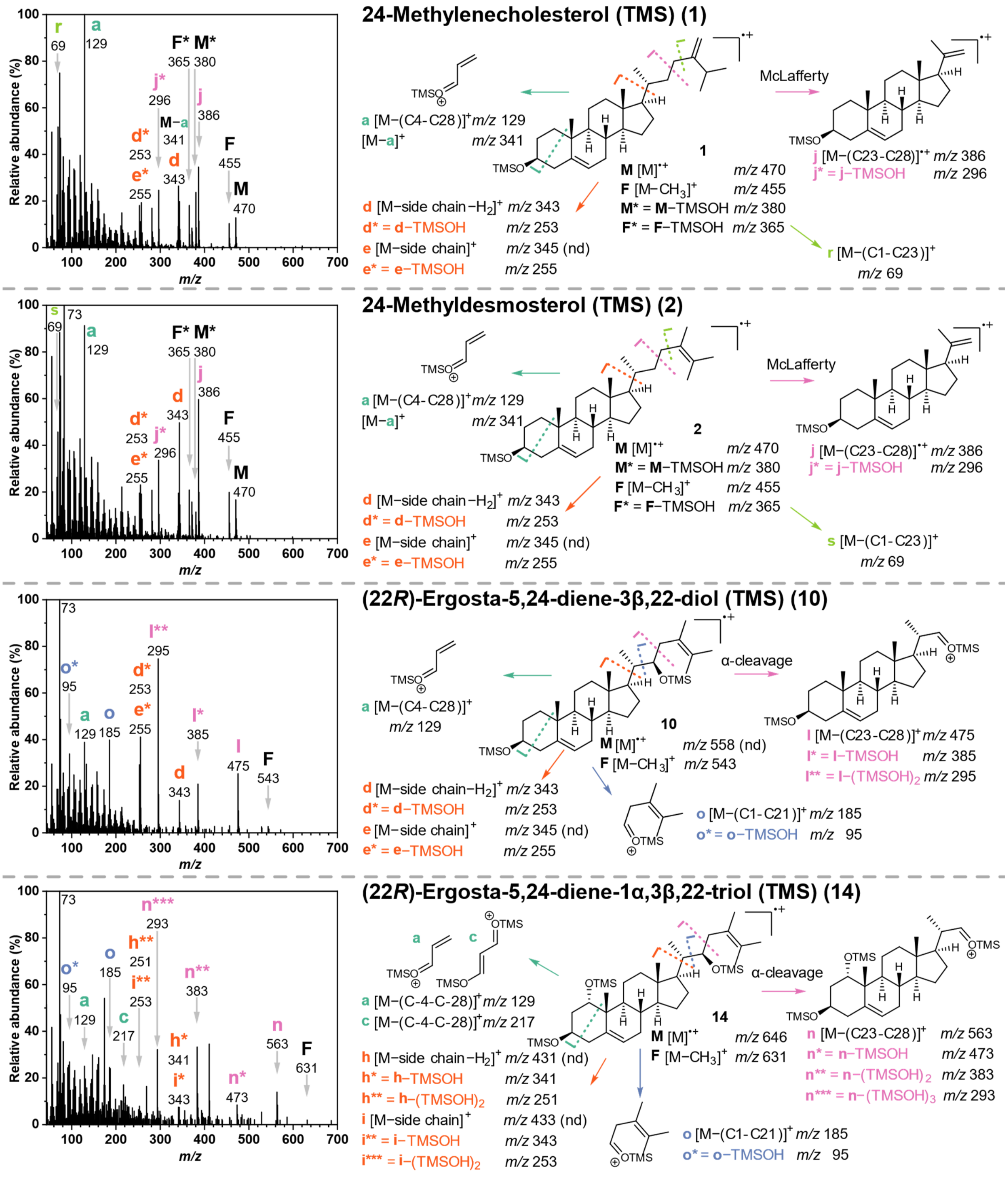


**Supplementary Fig. 13. Electron impact mass spectra and proposed fragmentation routes of main withanolide pathway intermediates from this work. See also Supplementary Table 3.**

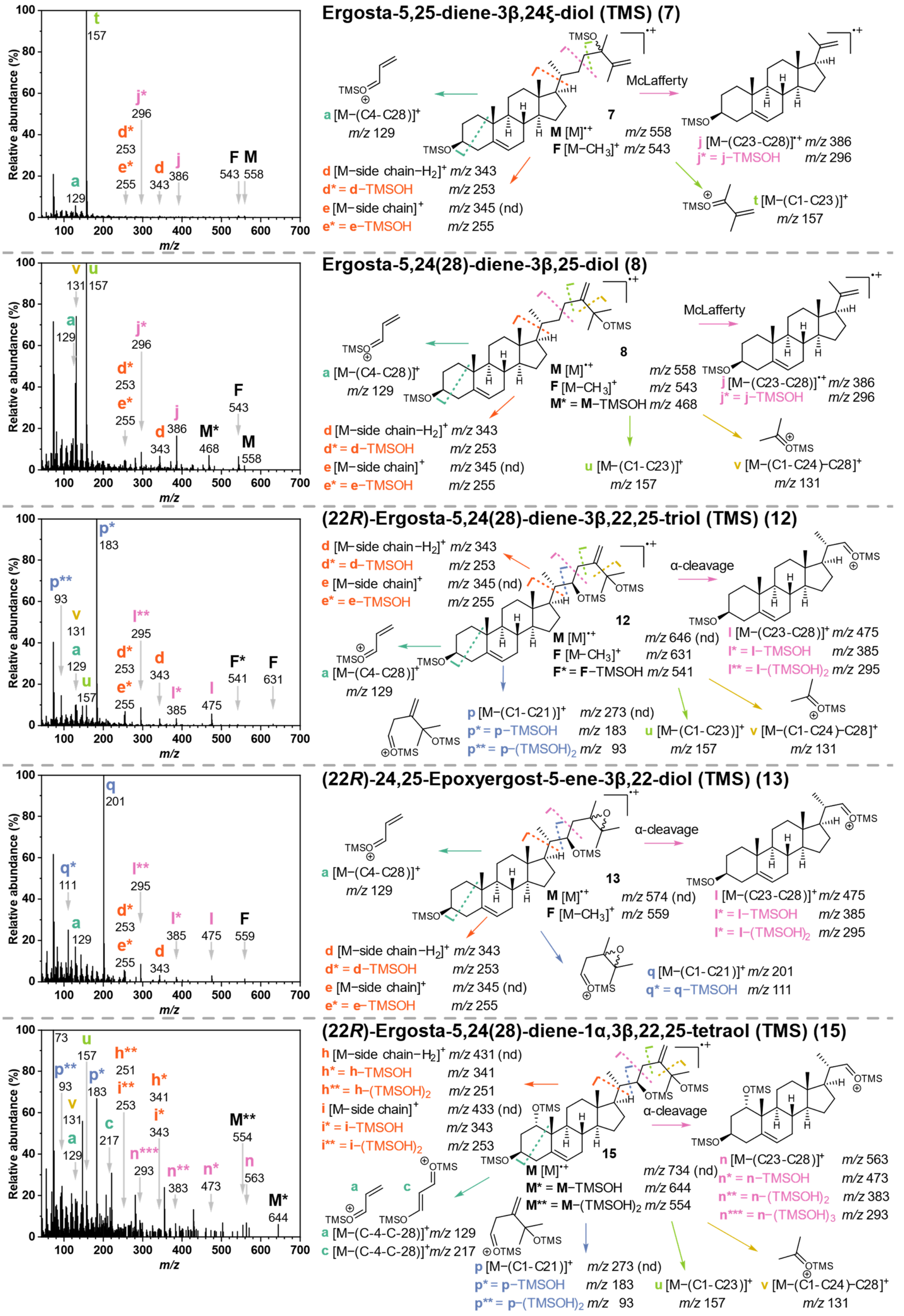


**Supplementary Fig. 14. Electron impact mass spectra and proposed fragmentation routes of *N. benthamiana* shunt products from this work. See also Supplementary Table 3.**

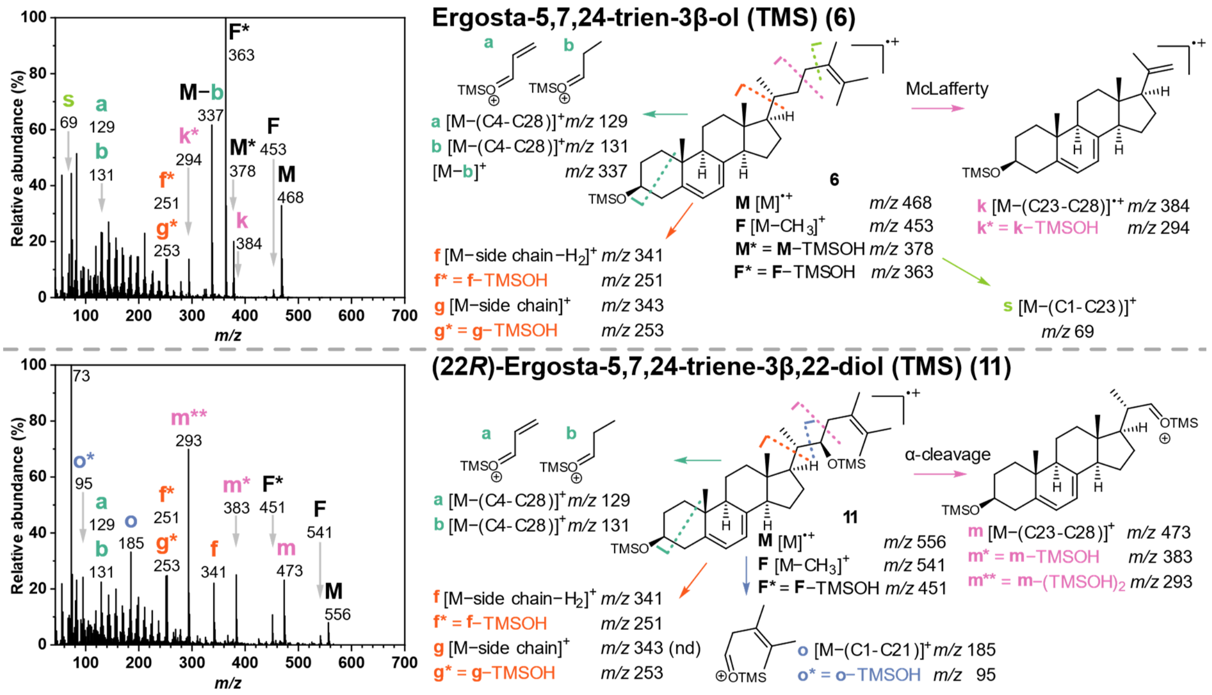


**Supplementary Fig. 15. Electron impact mass spectra and proposed fragmentation routes of *S. cerevisiae* shunt products from this work. See also Supplementary Table 3.**

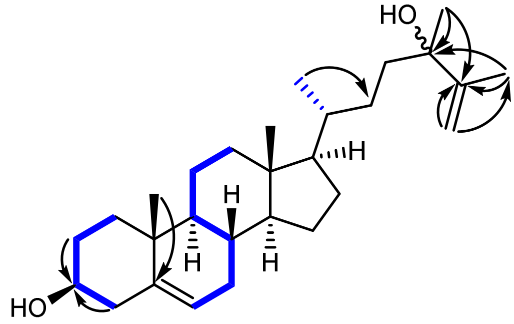

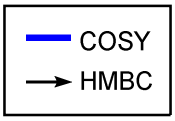


Ergosta-5,25-diene-3β,24ξ-diol (physalindicanol A) (**7**)

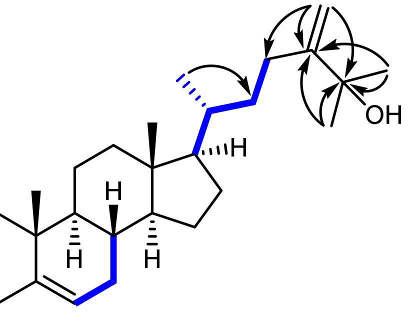

Ergosta-5,24(28)-diene-3β,25-diol (physalindicanol B) (**8**)

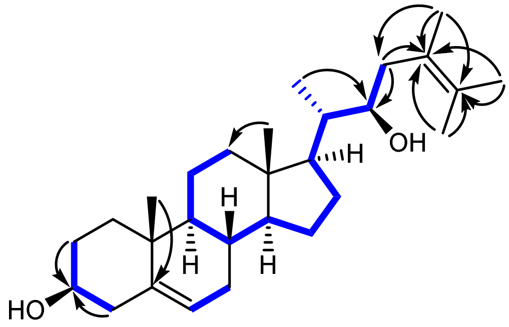

(22*R*)-Ergosta-5,24-diene-3β,22-diol (**10**)

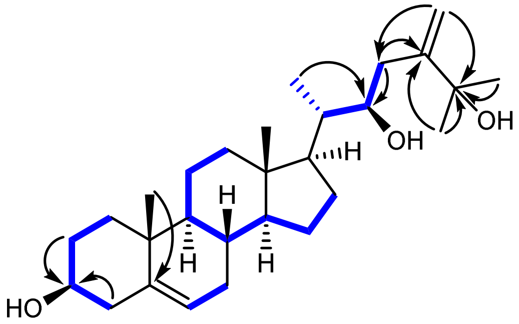

(22*R*)-Ergosta-5,24(28)-diene-3β,22,25-triol (phyministerol A) (**12**)

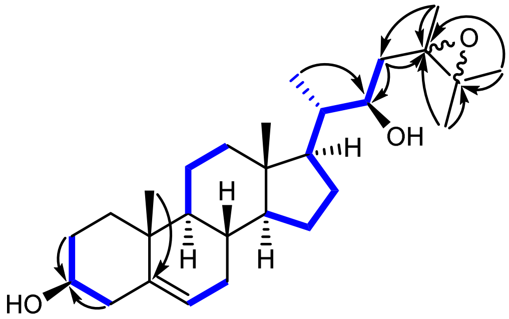

(22*R*)-24,25-Epoxyergost-5-ene-3β,22-diol (**13**)

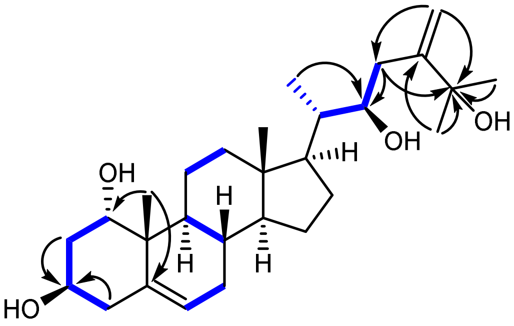

(22*R*)-Ergosta-5,24(28)-diene-1α,3β,22,25-tetraol (**15**)

**Supplementary Fig. 16. Key NMR correlations (COSY and HMBC) for the structure elucidation of isolated compounds.**

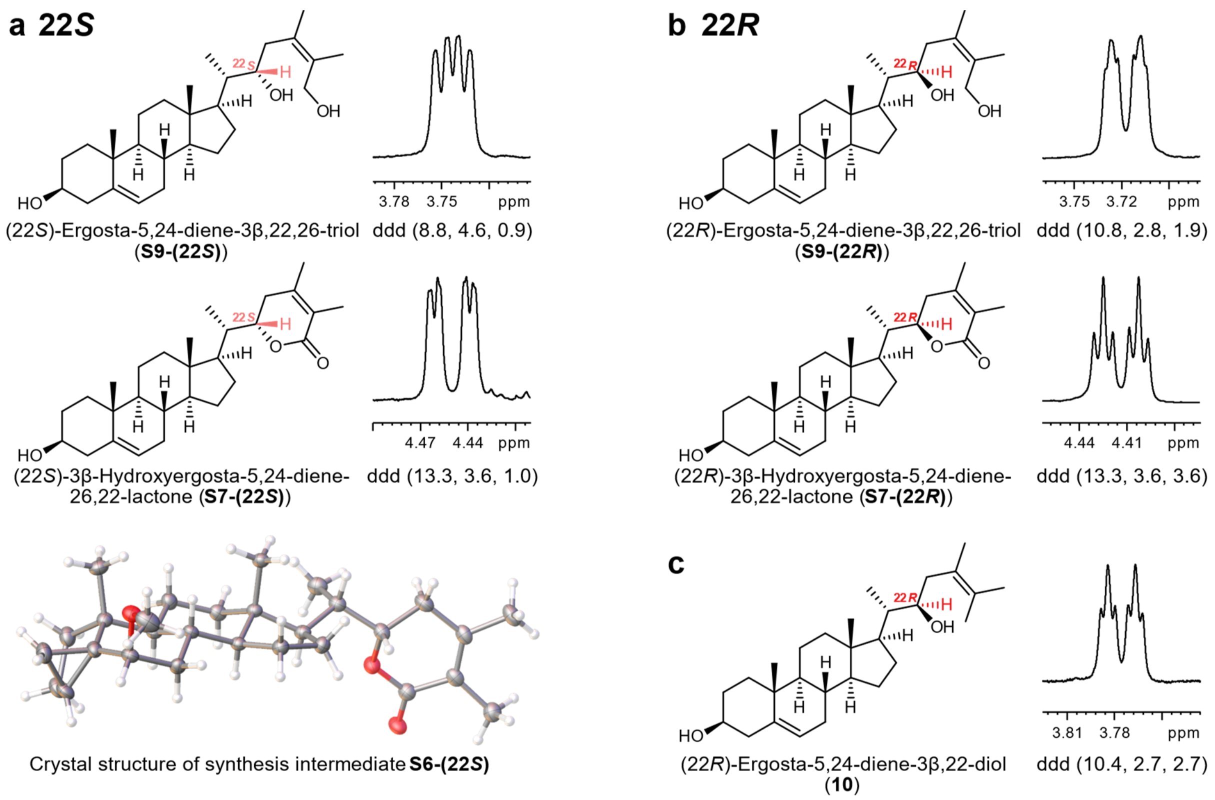


**Supplementary Fig. 17. Determination of C-22 stereochemistry of (22*R*)-ergosta-5,24-diene-3β,22-diol (10) by comparison with H-22 multiplets of synthetic diastereomer pairs S7-(22*S*/*R*) and S9-(22*S*/*R*) (MeOD, 600 MHz, 298 K).**

**a** H-22 multiplets and coupling constants of synthetic 22*S* compounds **S9-(22*S*)** and **S7-(22*S*)**. The 22*S* stereochemistry was confirmed by X-ray analysis of synthesis intermediate **S6-(22*S*)** (see also Supplementary Table 21).

**b** H-22 multiplets and coupling constants of synthetic 22*R* compounds **S9-(22*R*)** and **S7-(22*R*)**.

**c** H-22 multiplet and coupling constants of (22*R*)-ergosta-5,24-diene-3β,22-diol (**10**). The occurrence of two approximately equal minor coupling constants indicates a 22*R* configuration.

| Shunt product **13** from *PpCYP87G1* expression in *N. benthamiana* | Steroids reported from *Petunia hybrida* |
|---|---|
| 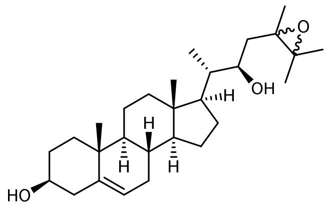 (22*R*)-24,25-Epoxyergost-5-ene-3β,22-diol (**13**) | 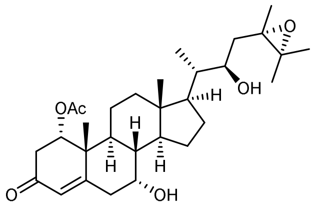 (22*R*,24*S*)-1α-Acetoxy-24,25-epoxy-7α,22-dihydroxyergost-4-en-3-one |
| | 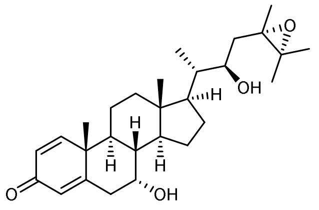 (22*R*,24*S*)-24,25-Epoxy-7α,22-dihydroxyergost-1,4-dien-3-one |

**Supplementary Fig. 18. Structure comparison of shunt product 13 with steroids isolated from *Petunia hybrida* (Solanaceae)[21].**

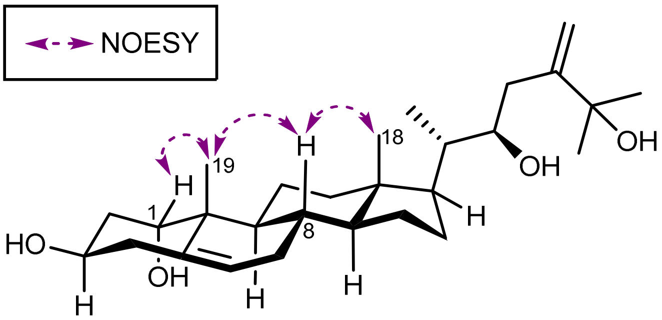


**Supplementary Fig. 19. Key NOESY correlations in support of 1α configuration of (22*R*)-ergosta-5,24(28)-diene-1α,3β,22,25-tetraol (15).**

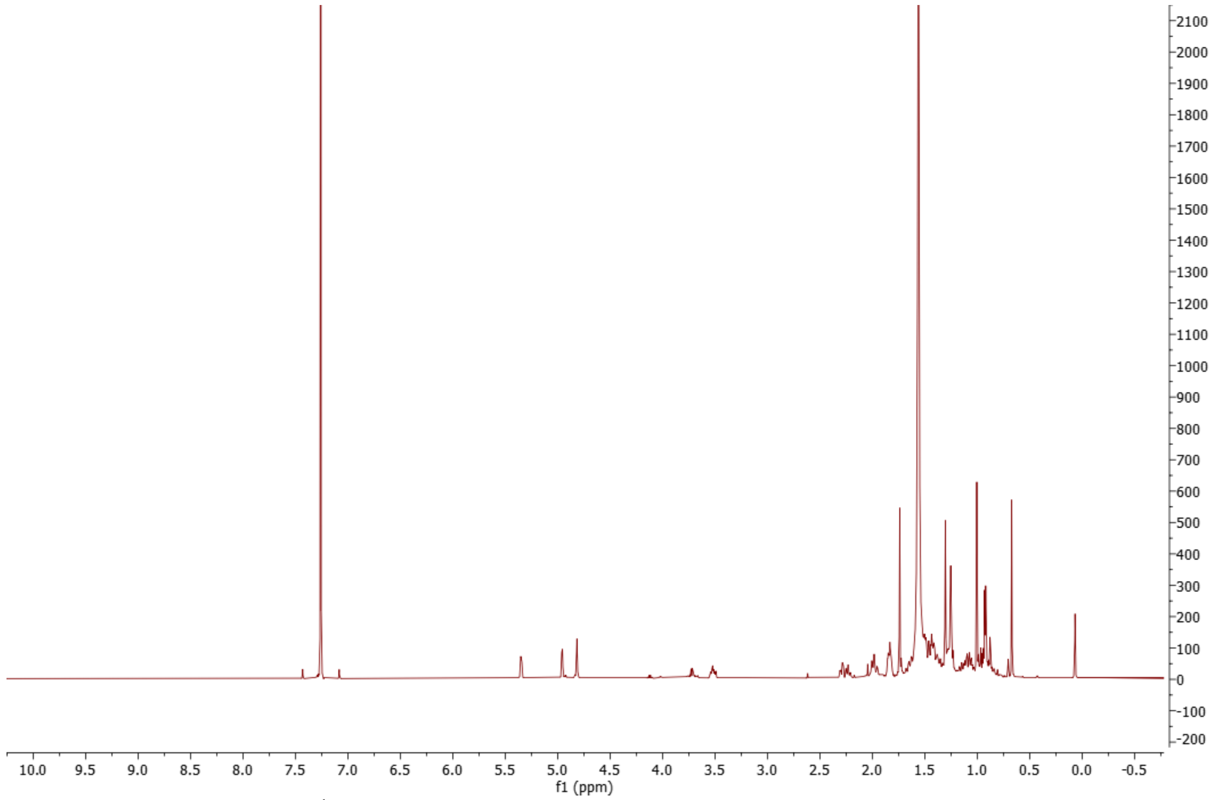


**Supplementary Fig. 20.** **$^{1}$H spectrum of ergosta-5,25-diene-3β,24ξ-diol (physalindicanol A) (7) ($CDCl_3$, 298 K, 600 MHz).**

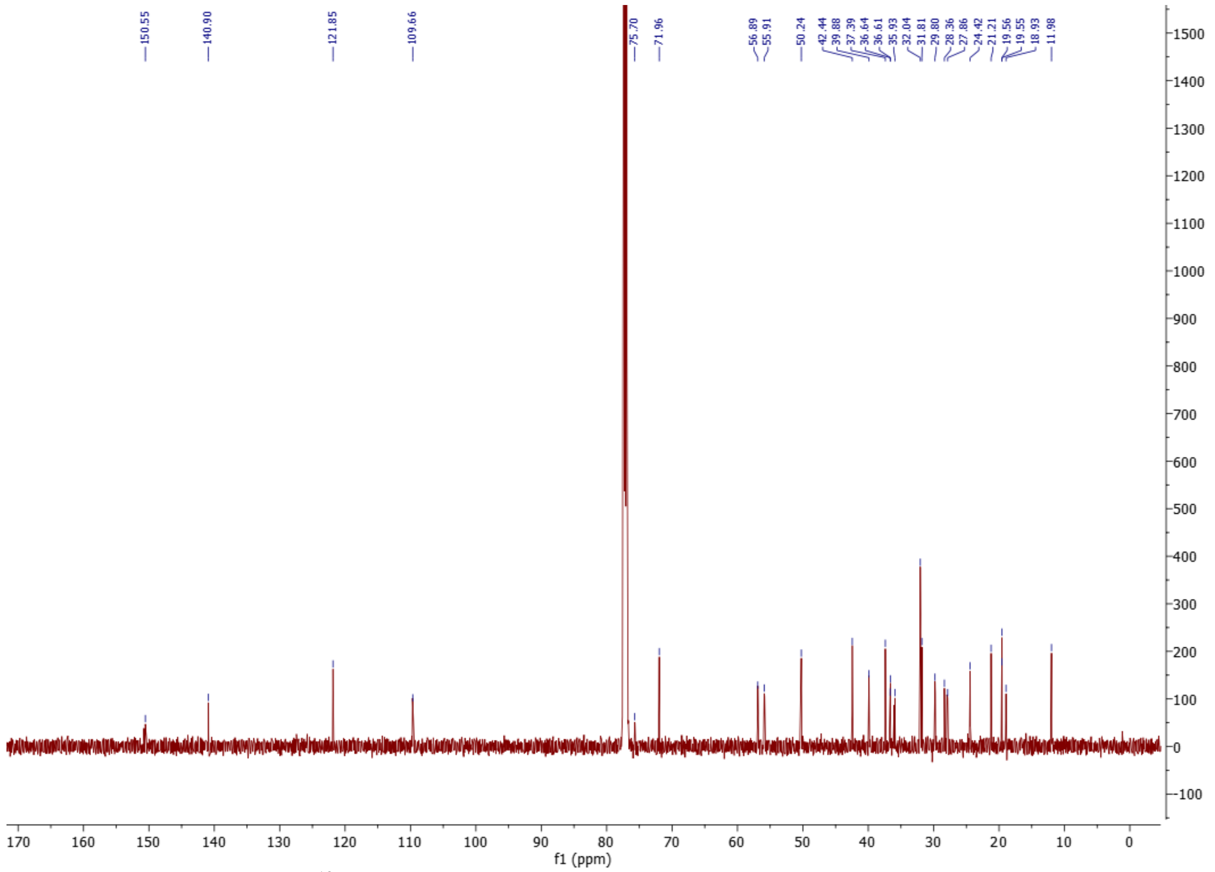


**Supplementary Fig. 21.** **$^{13}$C spectrum of ergosta-5,25-diene-3β,24ξ-diol (physalindicanol A) (7) ($CDCl_3$, 298 K, 151 MHz).**

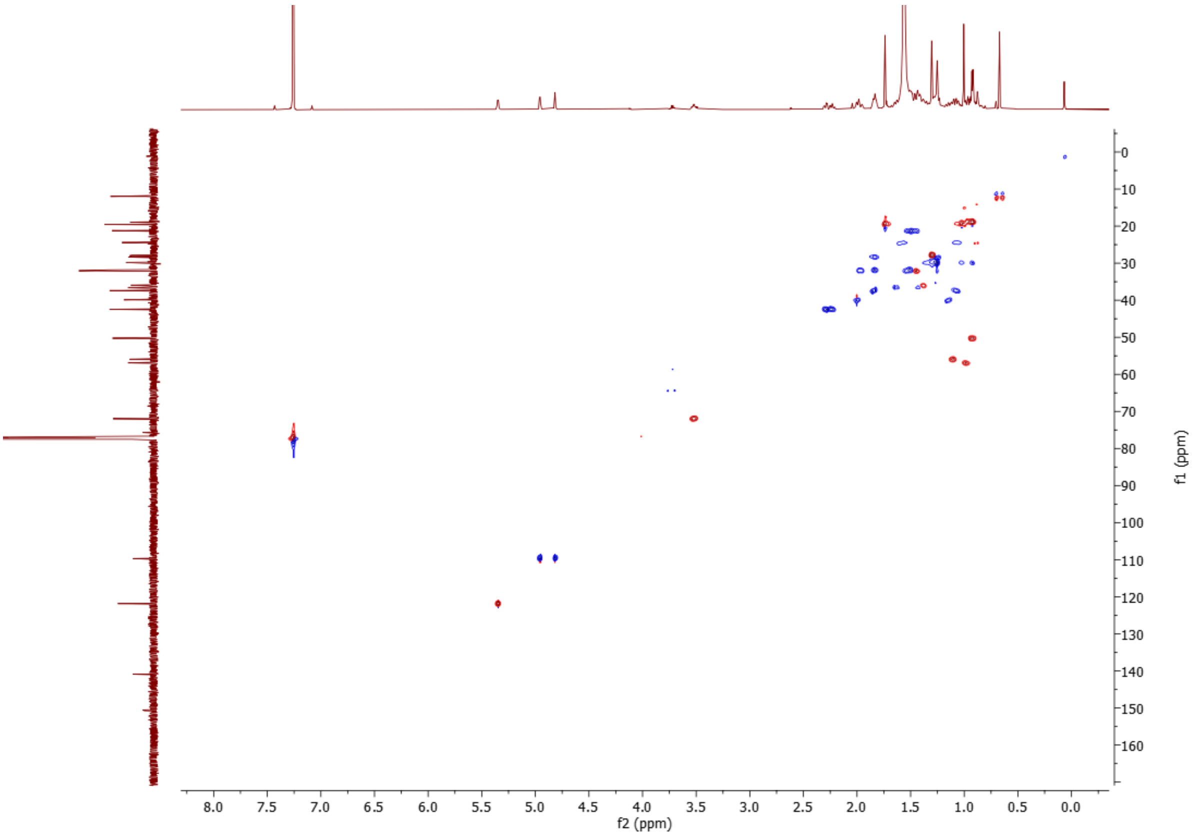


**Supplementary Fig. 22. HSQC spectrum of ergosta-5,25-diene-3β,24ξ-diol (physalindicanol A) (7) ($CDCl_3$, 298 K, 600 MHz).**

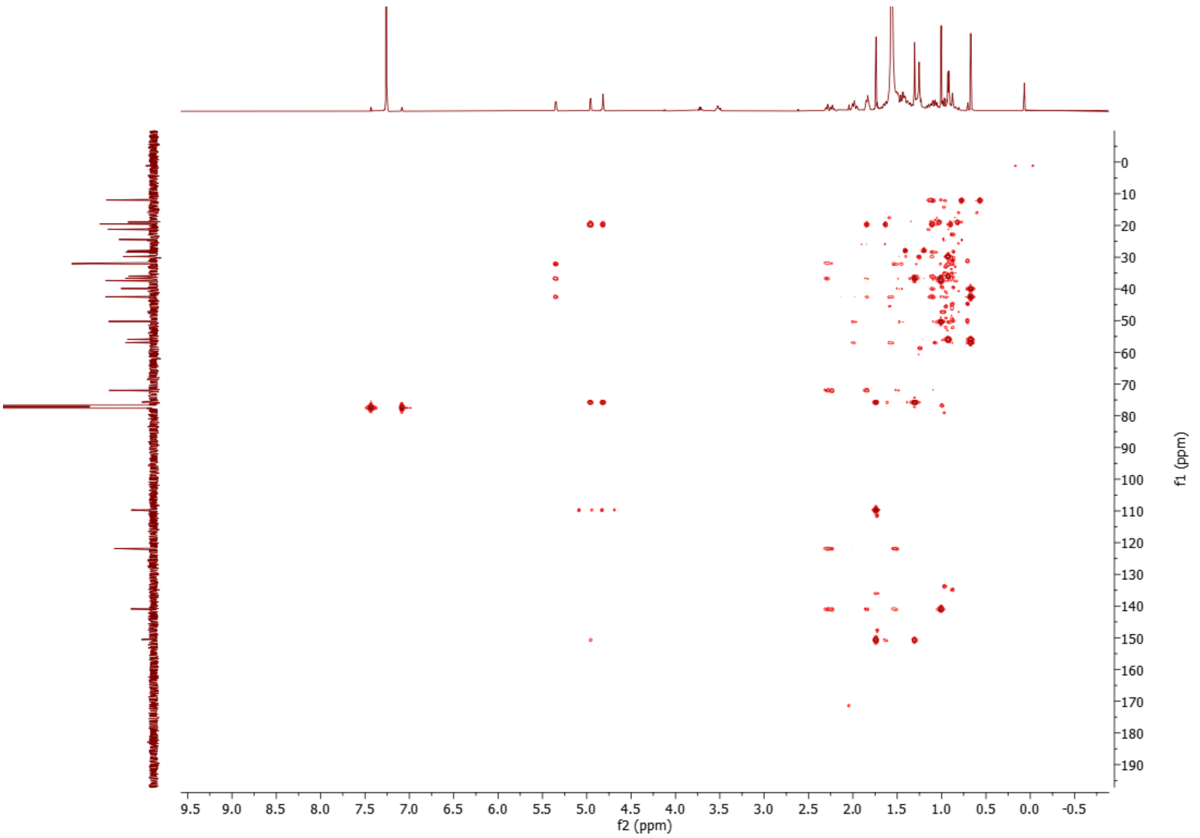


**Supplementary Fig. 23. HMBC spectrum of ergosta-5,25-diene-3β,24ξ-diol (physalindicanol A) (7) ($CDCl_3$, 298 K, 600 MHz).**

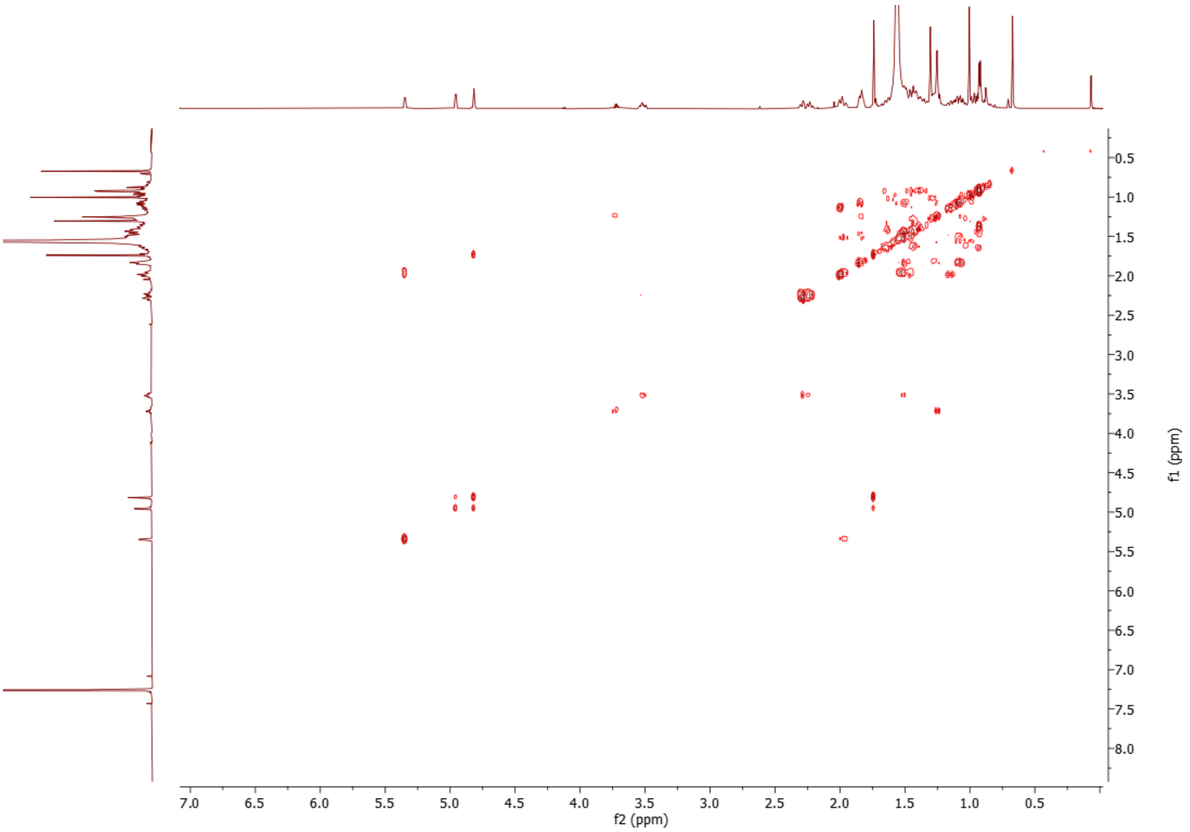


**Supplementary Fig. 24. COSY spectrum of ergosta-5,25-diene-3β,24ξ-diol (physalindicanol A) (7) ($CDCl_3$, 298 K, 600 MHz).**

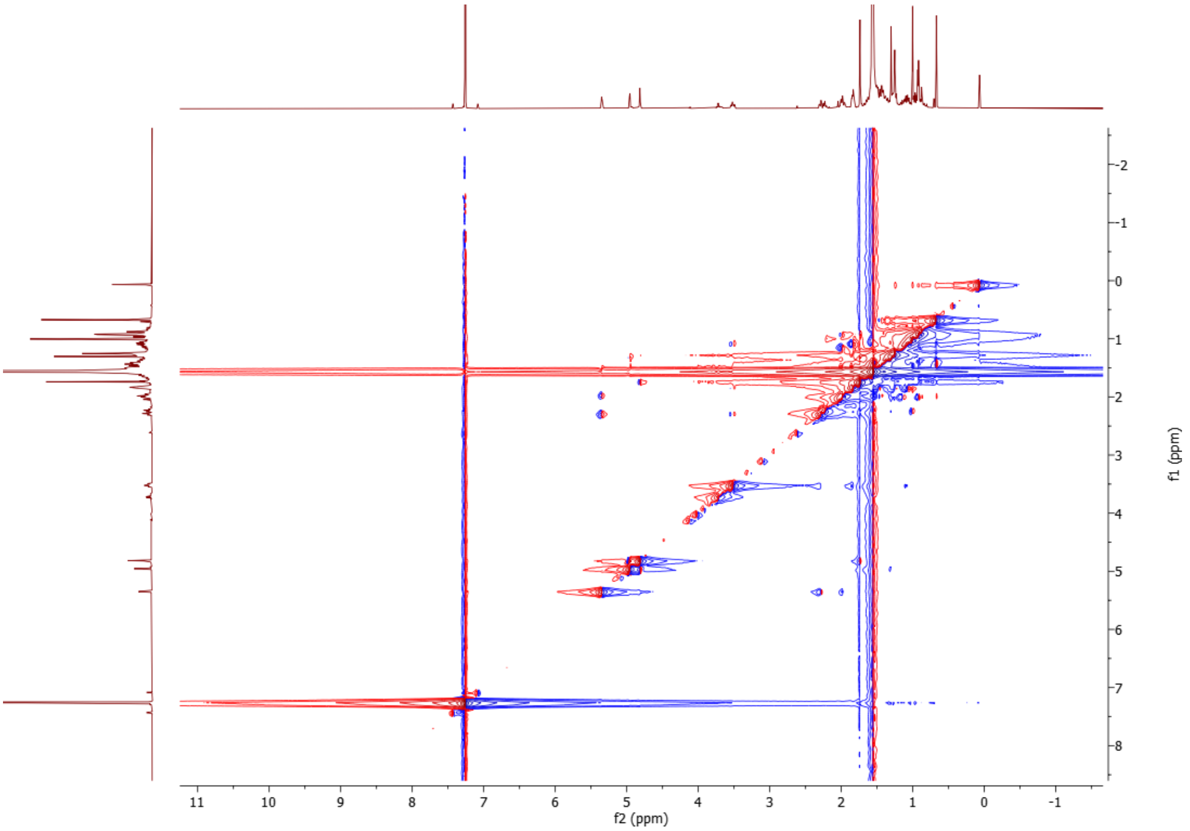


**Supplementary Fig. 25. NOESY spectrum of ergosta-5,25-diene-3β,24ξ-diol (physalindicanol A) (7) ($CDCl_3$, 298 K, 600 MHz).**

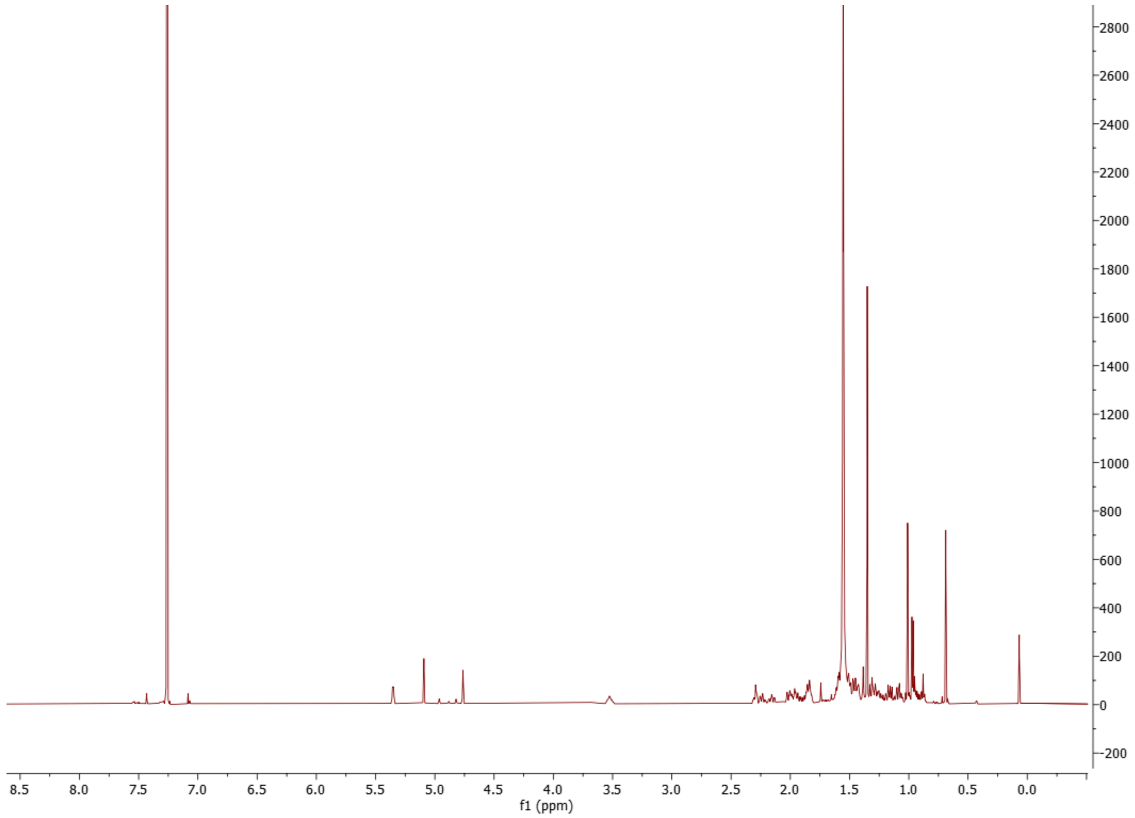

**Supplementary Fig. 26. [1]H spectrum of ergosta-5,24(28)-diene-3β,25-diol (8) (physalindicanol B) ($CDCl_3$, 298 K, 600 MHz).**

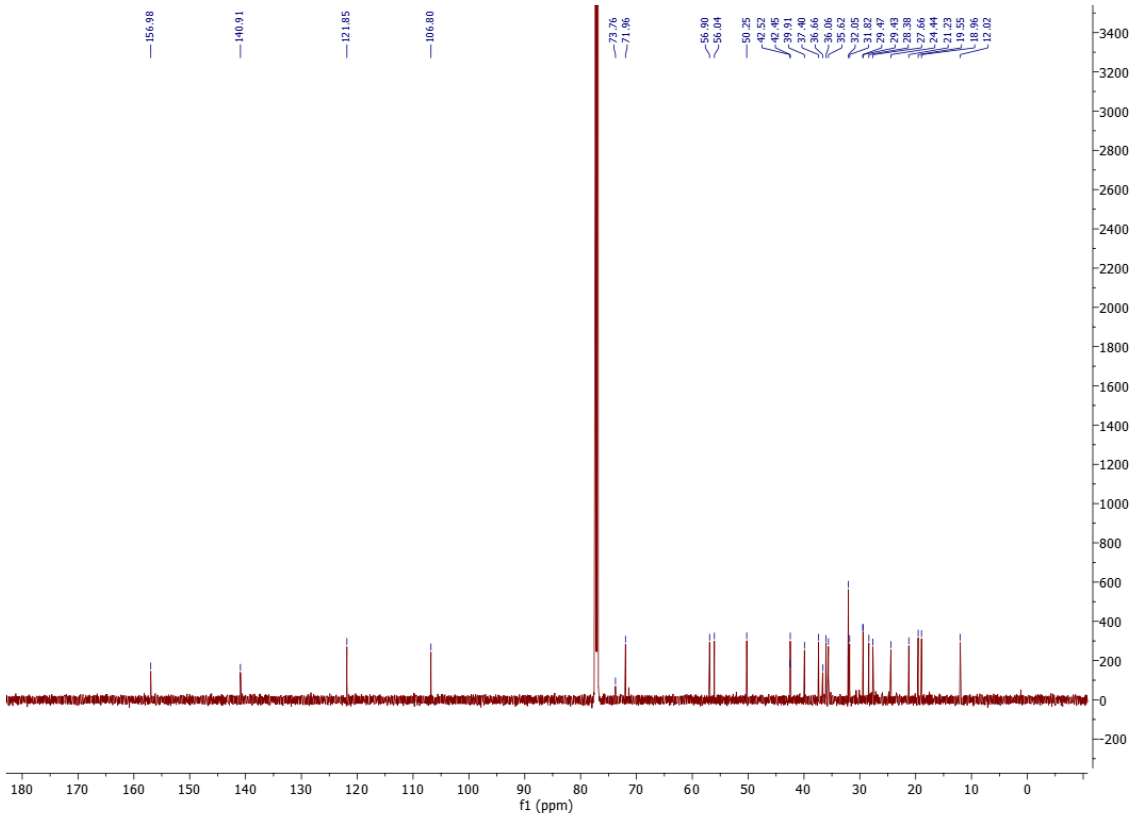

**Supplementary Fig. 27. [13]C spectrum of ergosta-5,24(28)-diene-3β,25-diol (8) (physalindicanol B) ($CDCl_3$, 298 K, 151 MHz).**

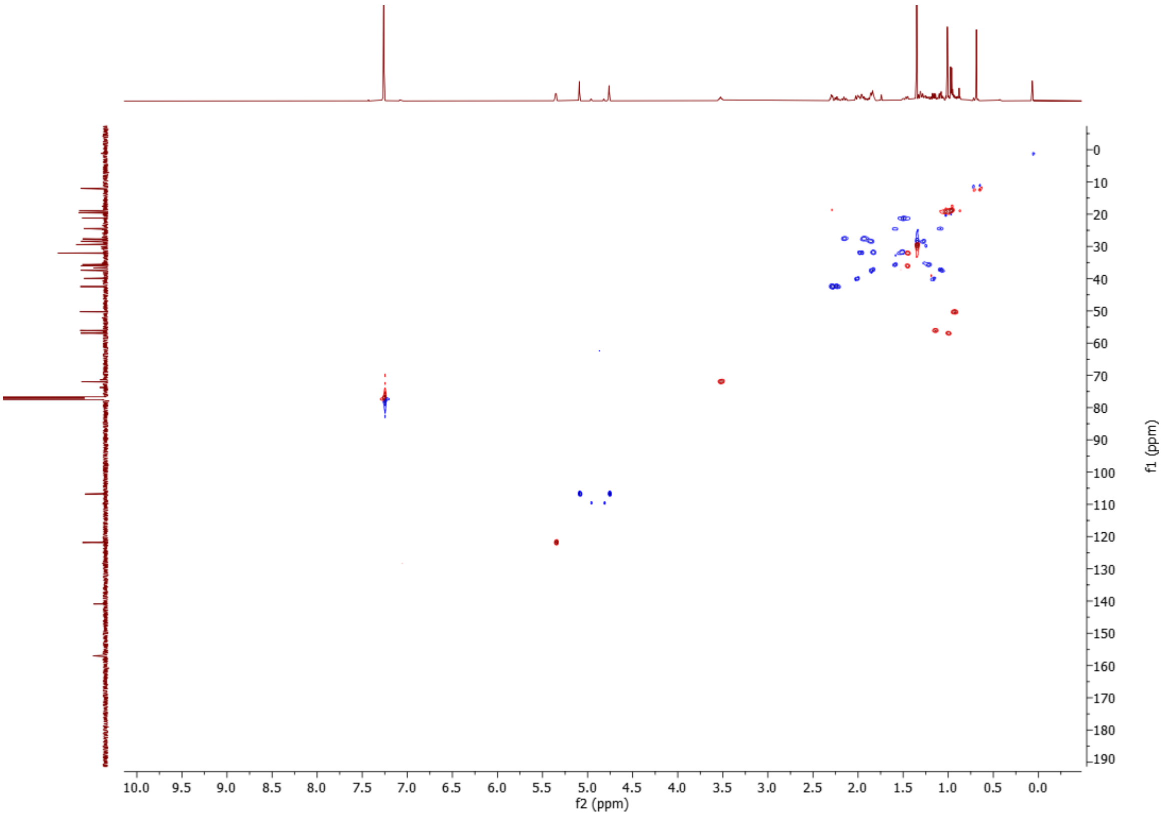


**Supplementary Fig. 28. HSQC spectrum of ergosta-5,24(28)-diene-3β,25-diol (8) (physalindicanol B) ($CDCl_3$, 298 K, 600 MHz).**

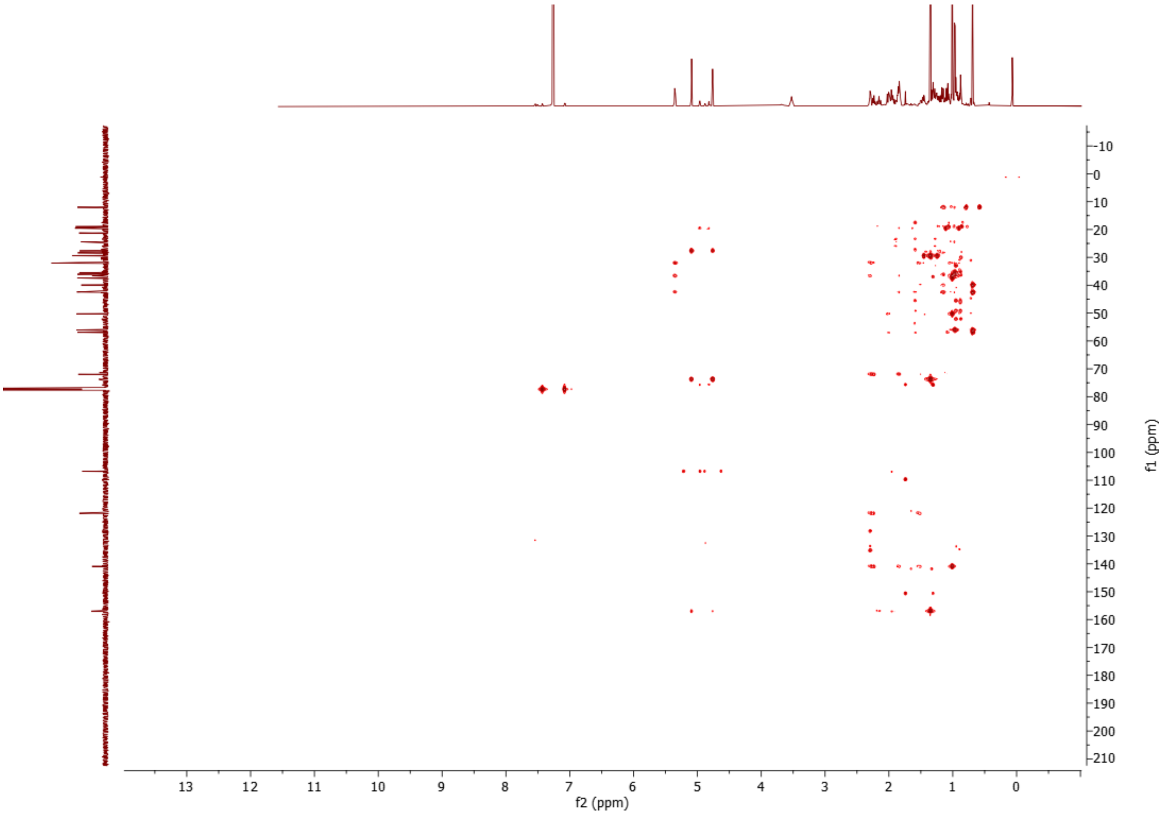


**Supplementary Fig. 29. HMBC spectrum of ergosta-5,24(28)-diene-3β,25-diol (8) (physalindicanol B) ($CDCl_3$, 298 K, 600 MHz).**

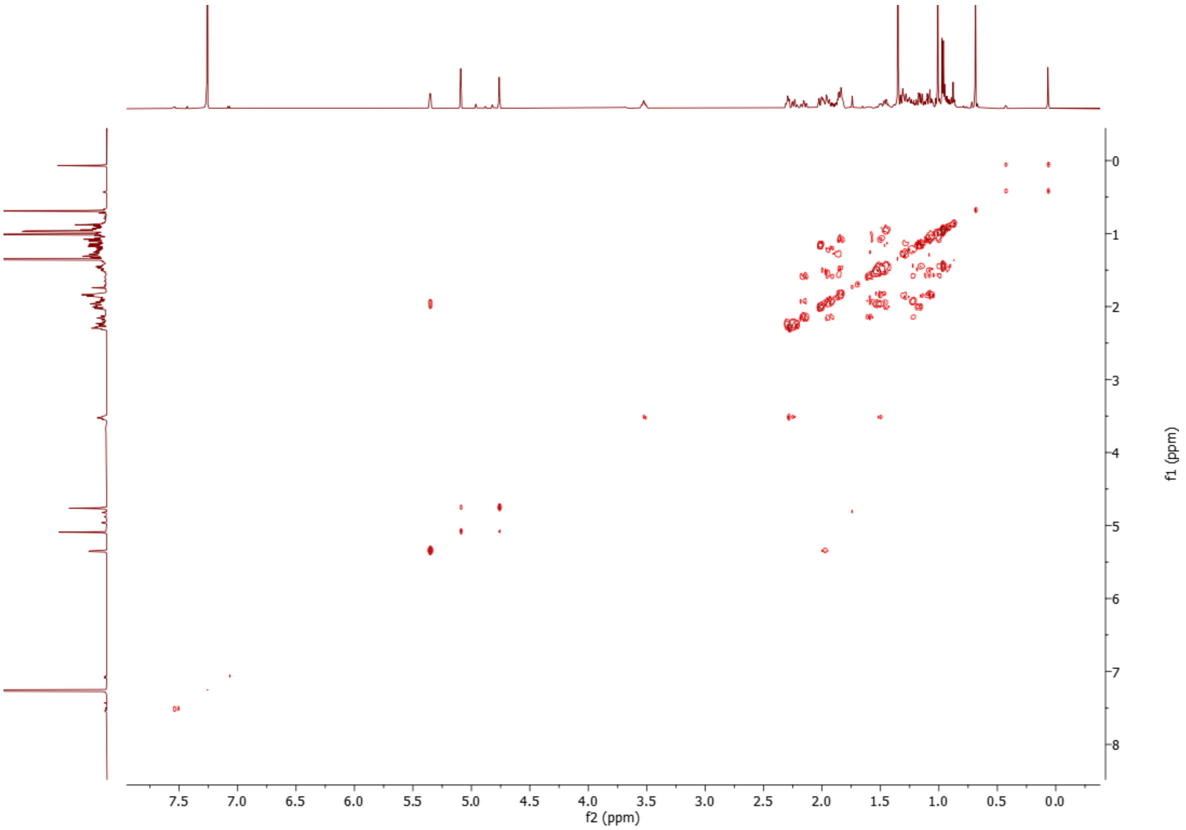


**Supplementary Fig. 30. COSY spectrum of ergosta-5,24(28)-diene-3β,25-diol (8) (physalindicanol B) ($CDCl_3$, 298 K, 600 MHz).**

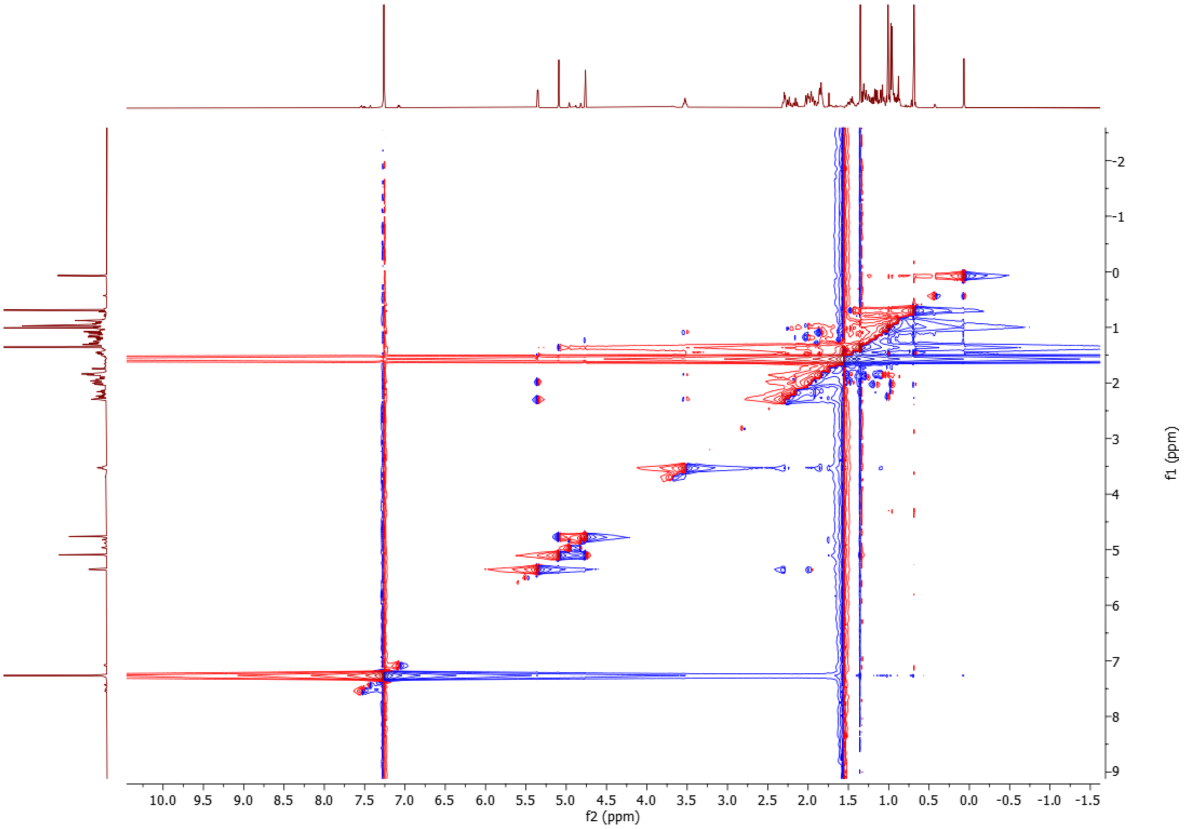


**Supplementary Fig. 31. NOESY spectrum of ergosta-5,24(28)-diene-3β,25-diol (8) (physalindicanol B) ($CDCl_3$, 298 K, 600 MHz).**

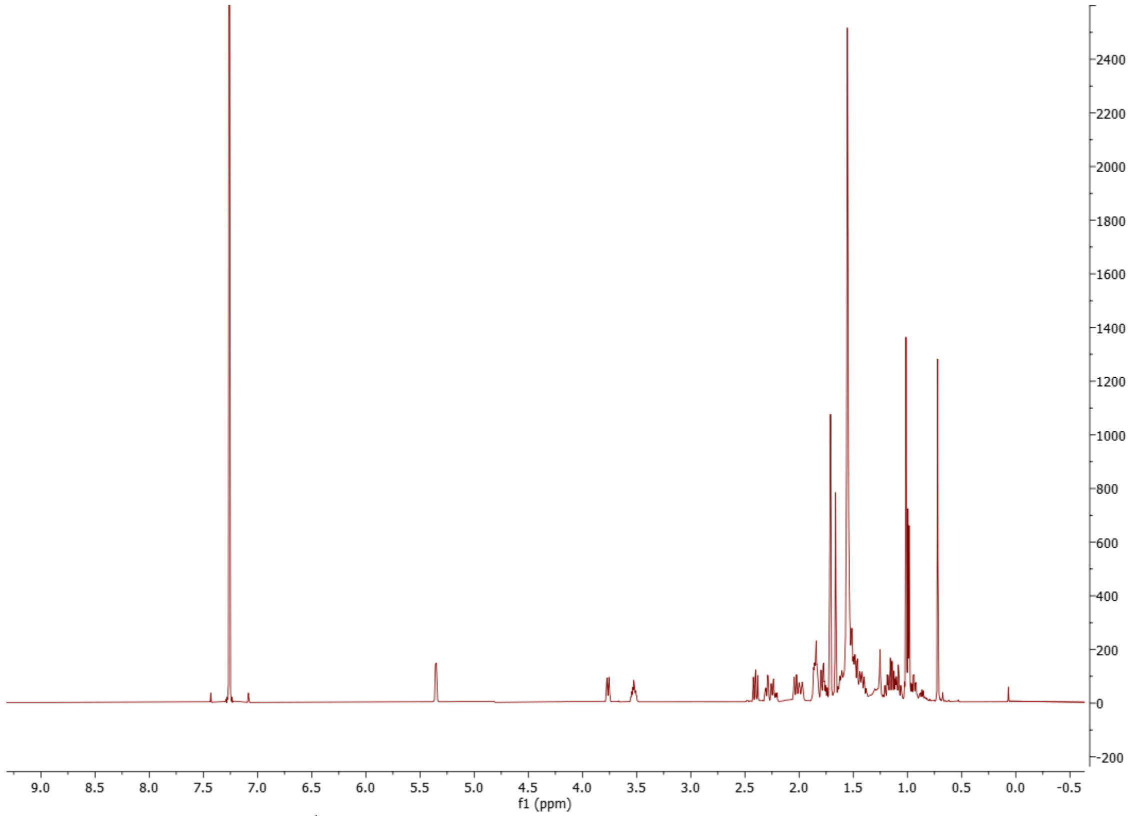


**Supplementary Fig. 32. $^{1}$H spectrum of (22*R*)-ergosta-5,24-diene-3β,22-diol (10) ($CDCl_3$, 298 K, 600 MHz).**

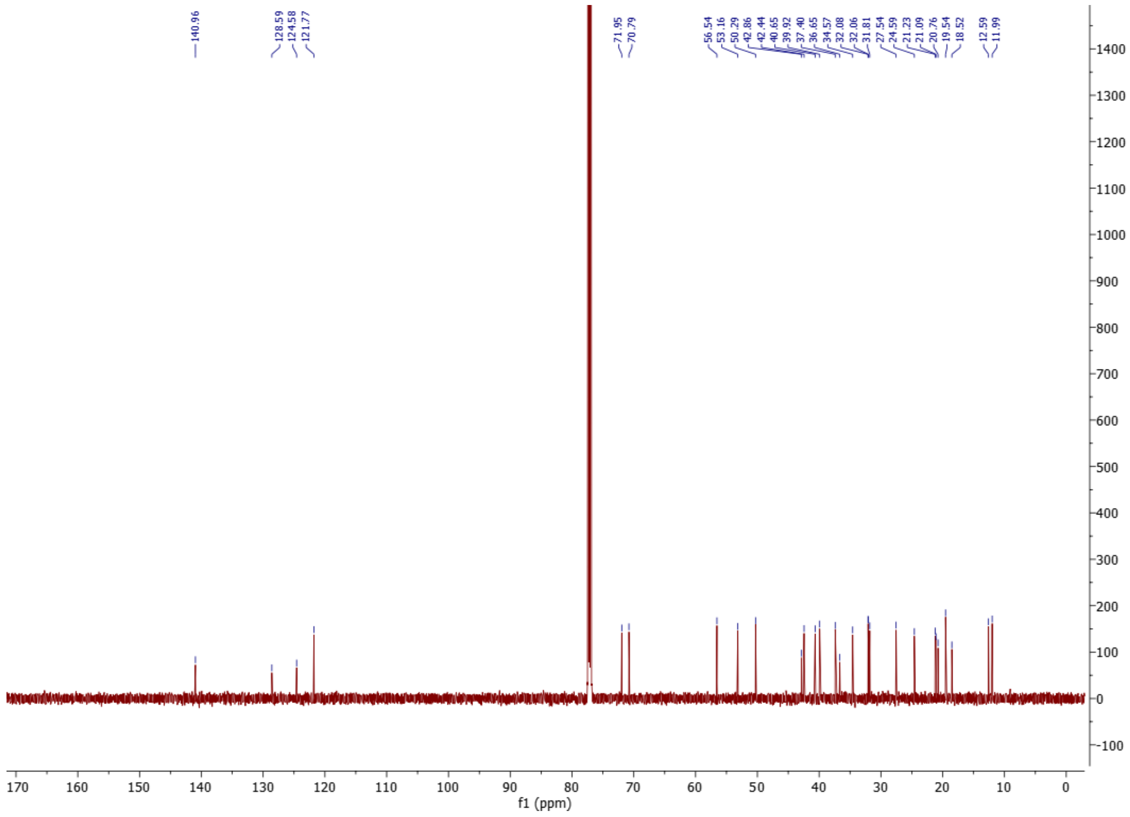


**Supplementary Fig. 33. $^{13}$C spectrum of (22*R*)-ergosta-5,24-diene-3β,22-diol (10) ($CDCl_3$, 298 K, 151 MHz).**

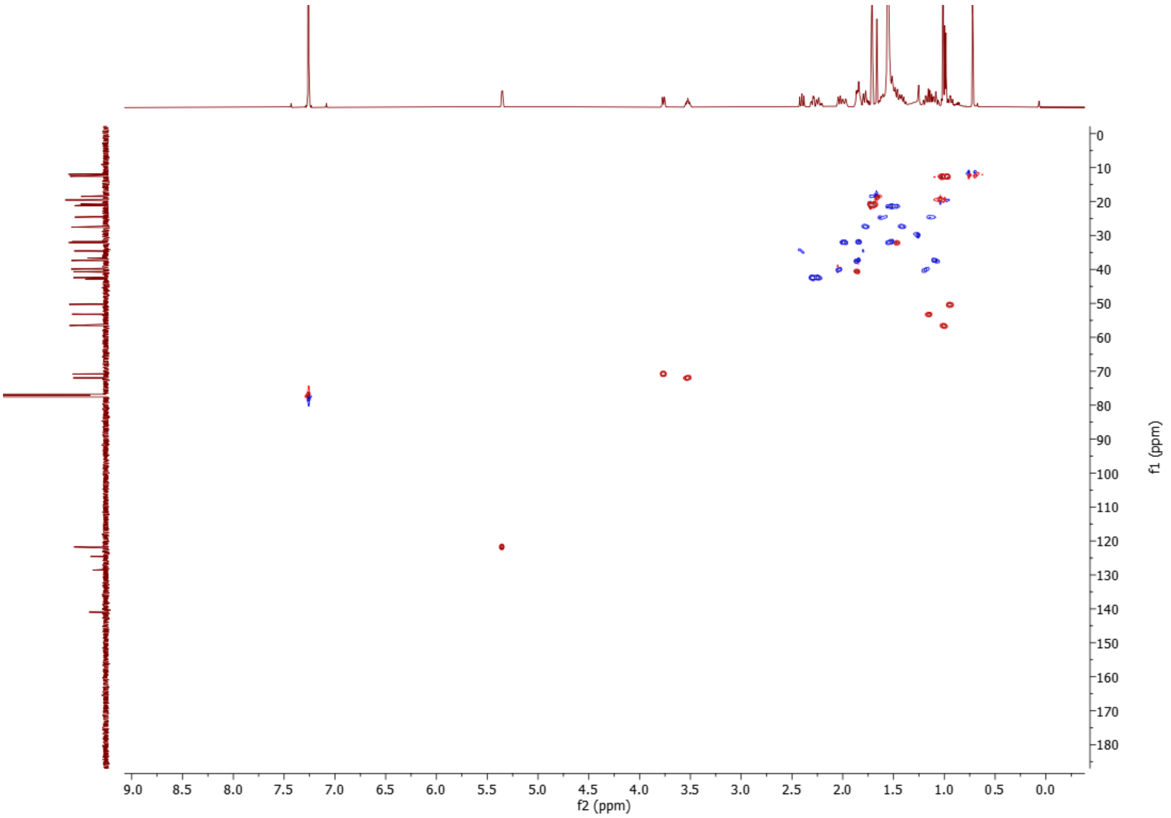


**Supplementary Fig. 34. HSQC spectrum of (22*R*)-ergosta-5,24-diene-3β,22-diol (10) ($CDCl_3$, 298 K, 600 MHz).**

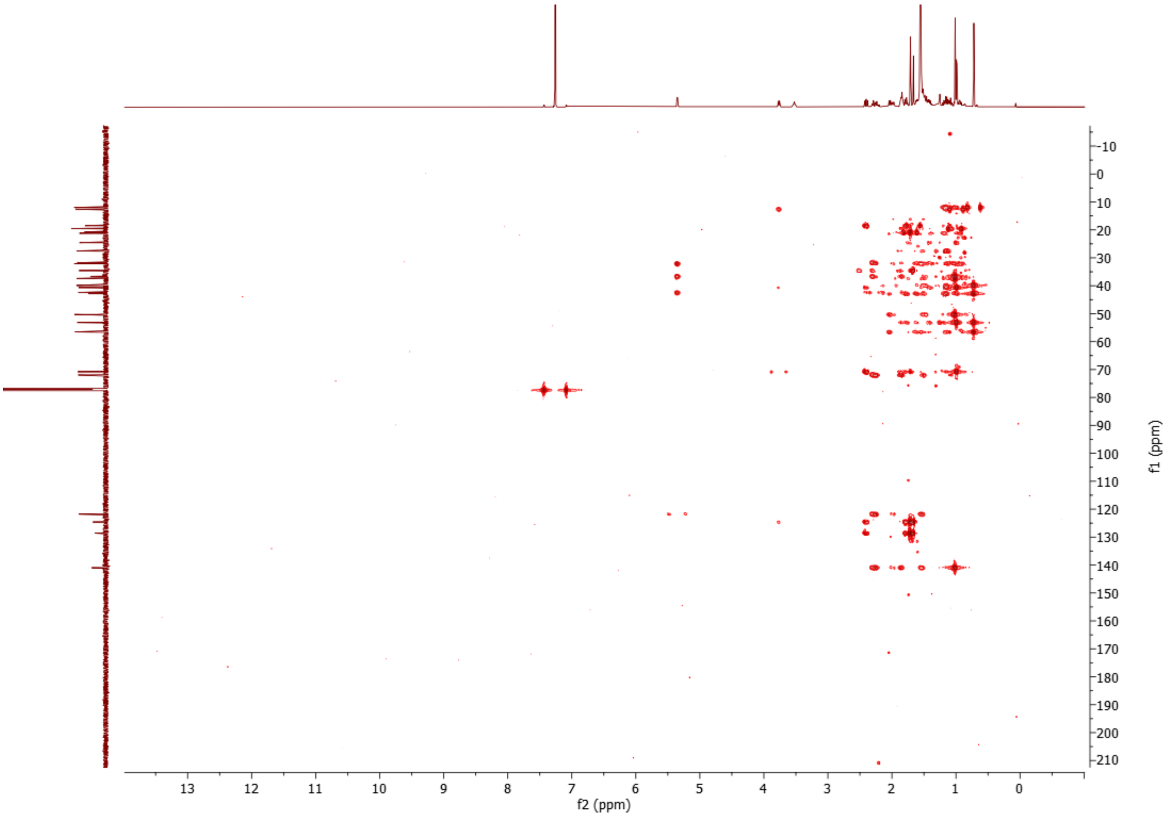


**Supplementary Fig. 35. HMBC spectrum of (22*R*)-ergosta-5,24-diene-3β,22-diol (10) ($CDCl_3$, 298 K, 600 MHz).**

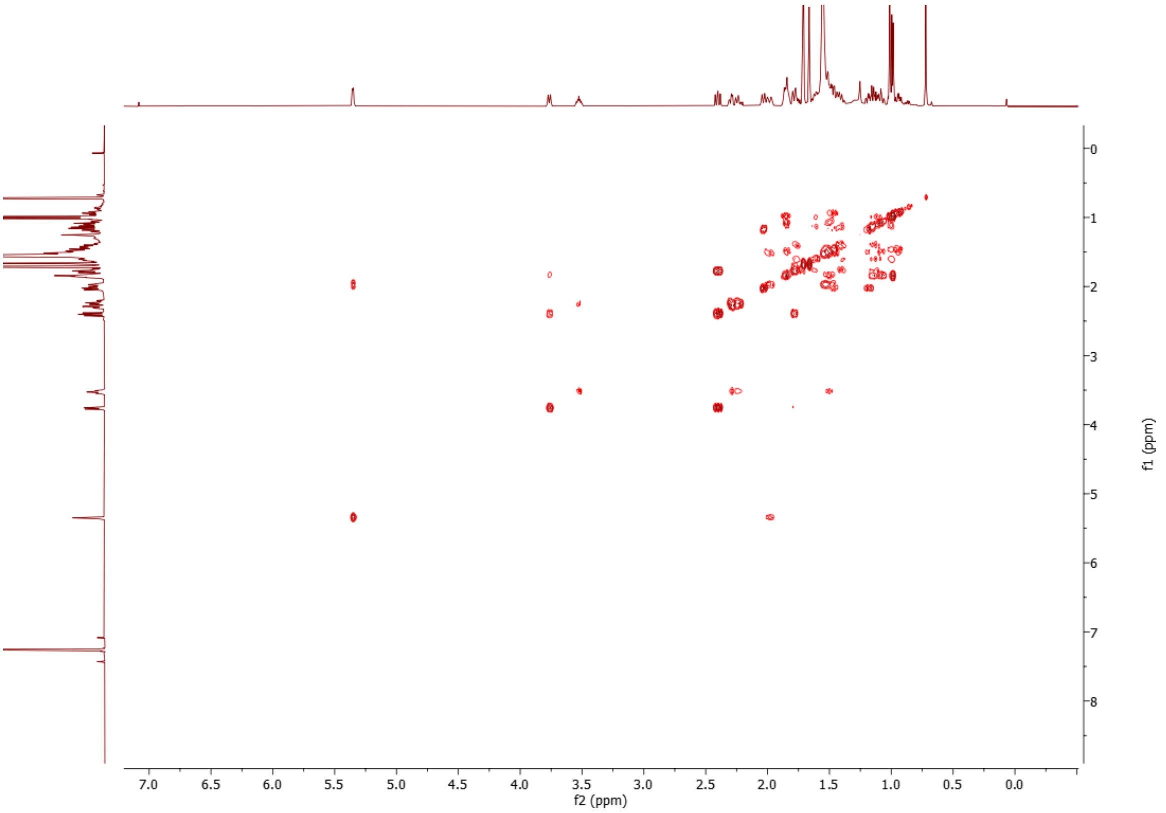


**Supplementary Fig. 36. COSY spectrum of (22*R*)-ergosta-5,24-diene-3β,22-diol (10) ($CDCl_3$, 298 K, 600 MHz).**

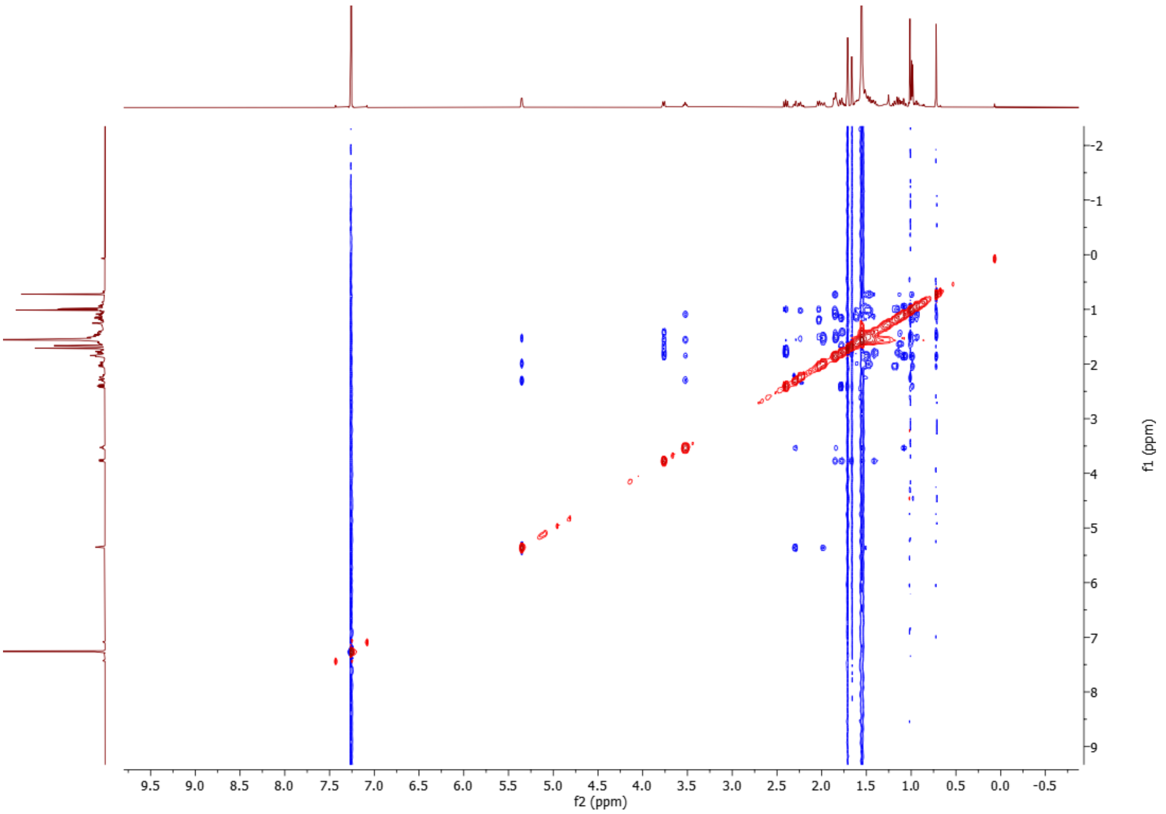


**Supplementary Fig. 37. NOESY spectrum of (22*R*)-ergosta-5,24-diene-3β,22-diol (10) ($CDCl_3$, 298 K, 600 MHz).**

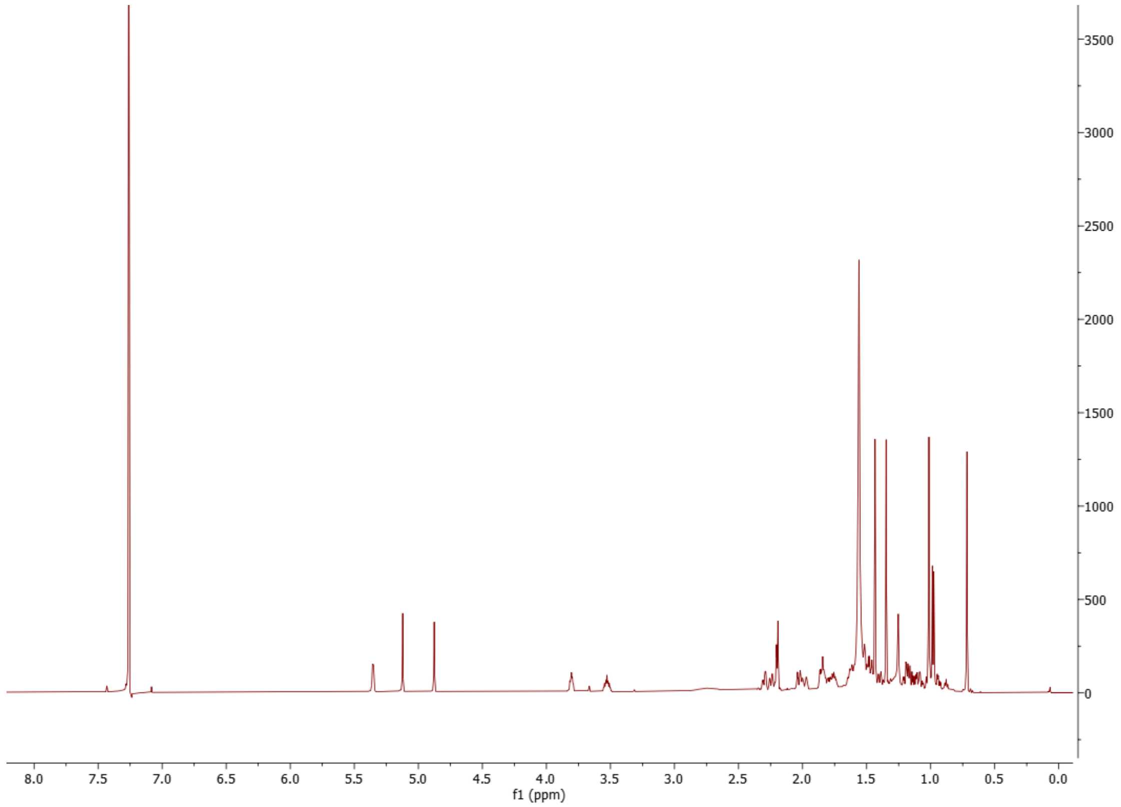


**Supplementary Fig. 38. [1]H spectrum of (22*R*)-ergosta-5,24(28)-diene-3β,22,25-triol (phyministerol A) (12) ($CDCl_3$, 298 K, 600 MHz).**

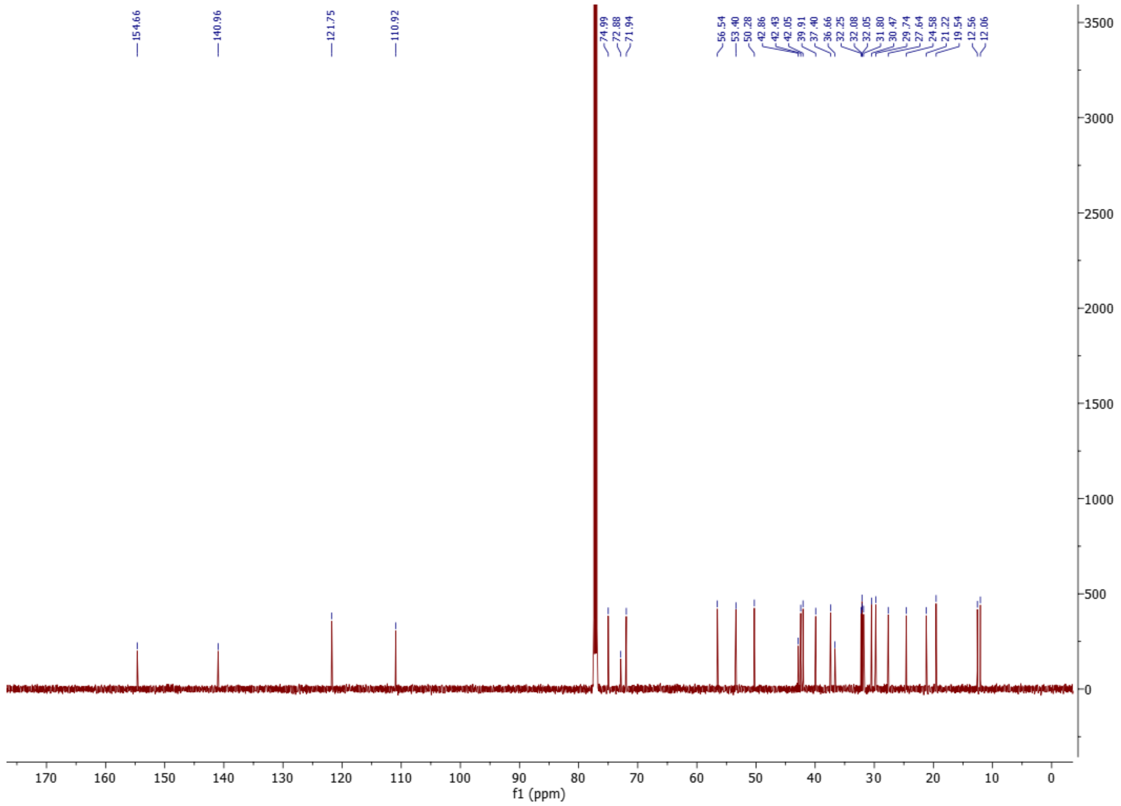


**Supplementary Fig. 39. [13]C spectrum of (22*R*)-ergosta-5,24(28)-diene-3β,22,25-triol (phyministerol A) (12) ($CDCl_3$, 298 K, 151 MHz).**

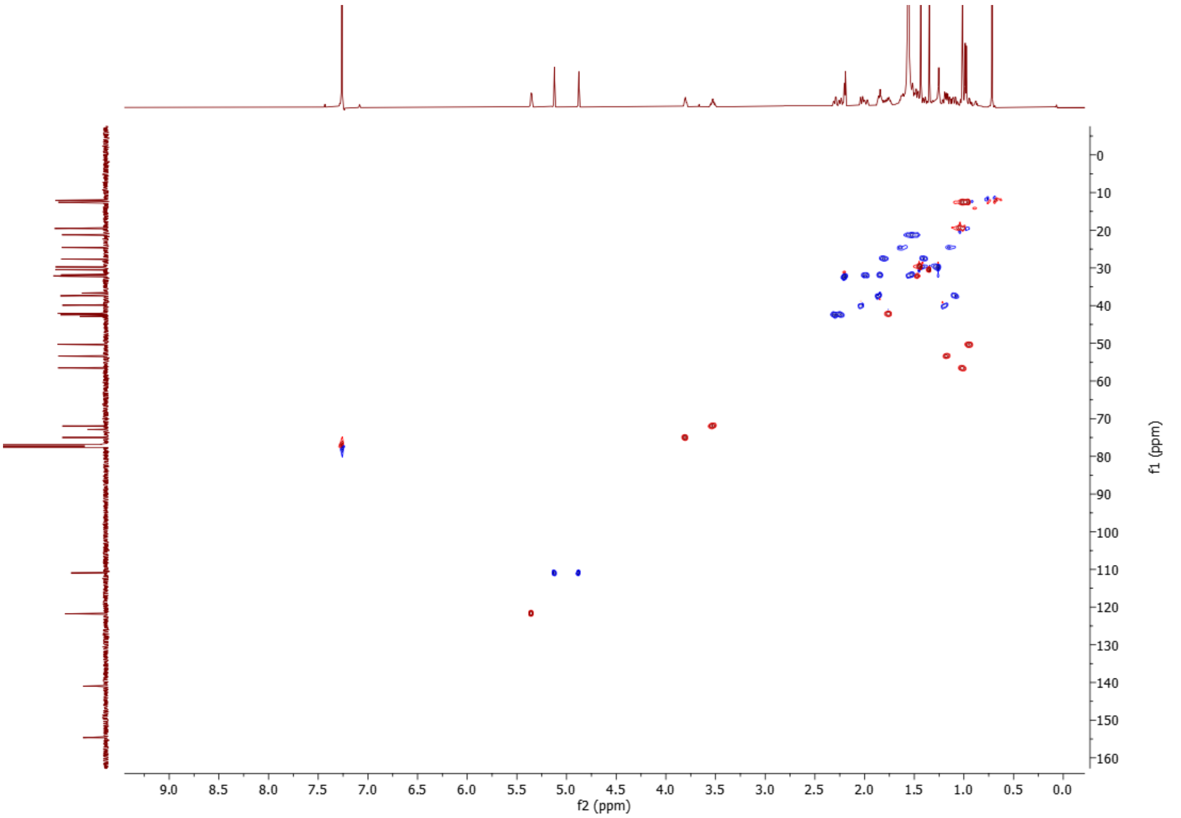


**Supplementary Fig. 40. HSQC spectrum of (22*R*)-ergosta-5,24(28)-diene-3β,22,25-triol (phyministerol A) (12) ($CDCl_3$, 298 K, 600 MHz).**

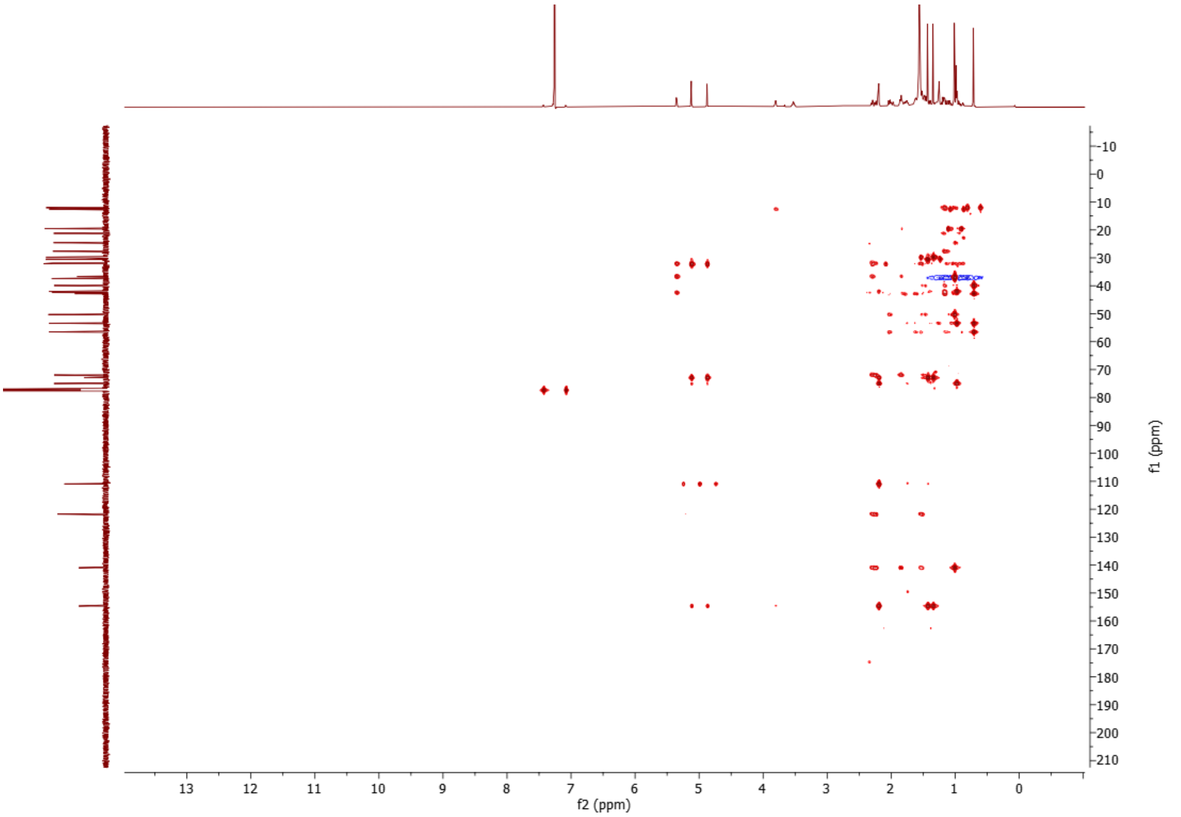


**Supplementary Fig. 41. HMBC spectrum of (22*R*)-ergosta-5,24(28)-diene-3β,22,25-triol (phyministerol A) (12) ($CDCl_3$, 298 K, 600 MHz).**

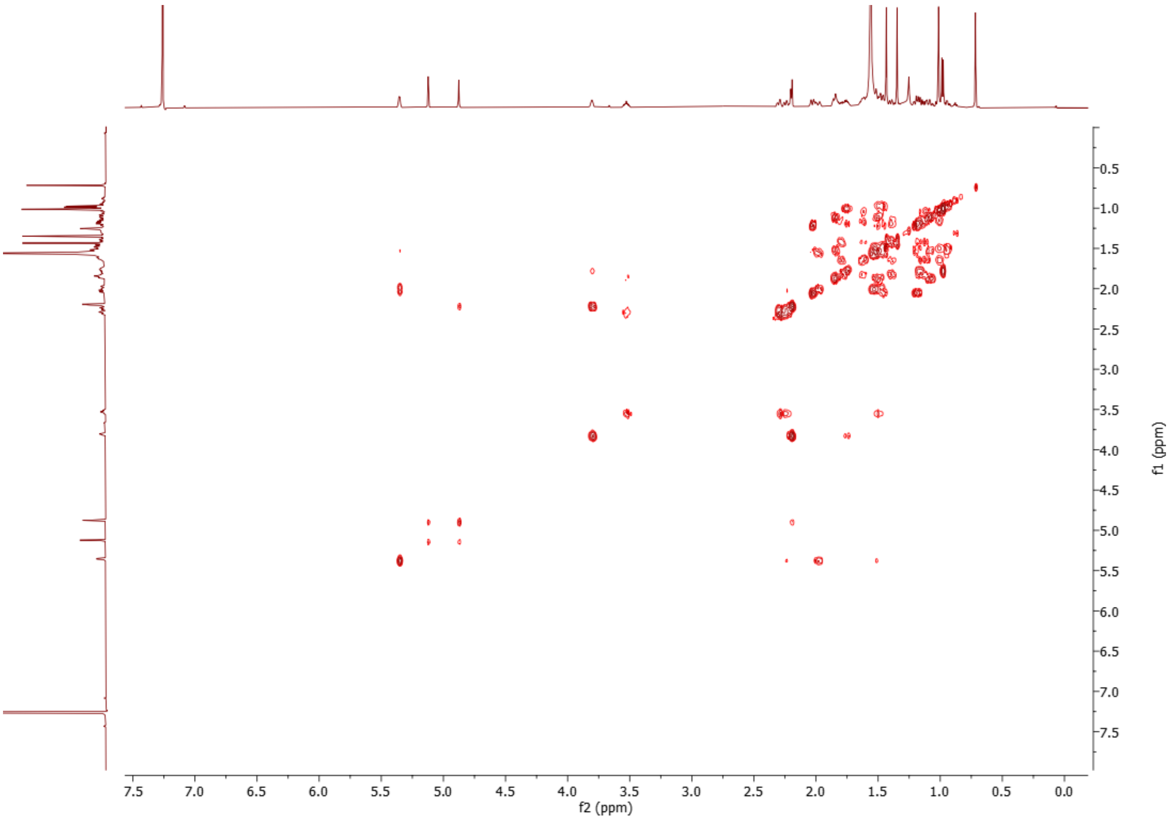


**Supplementary Fig. 42. COSY spectrum of (22*R*)-ergosta-5,24(28)-diene-3β,22,25-triol (phyministerol A) (12) ($CDCl_3$, 298 K, 600 MHz).**

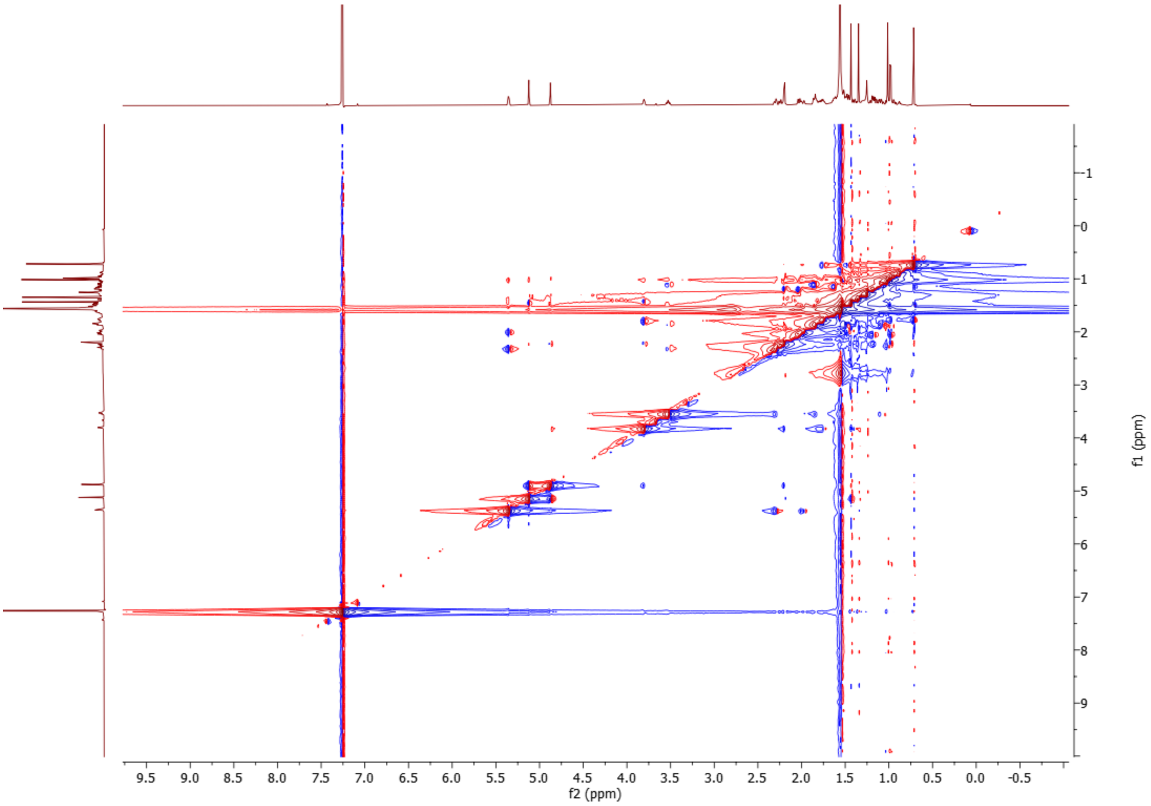


**Supplementary Fig. 43. NOESY spectrum of (22*R*)-ergosta-5,24(28)-diene-3β,22,25-triol (phyministerol A) (12) ($CDCl_3$, 298 K, 600 MHz).**

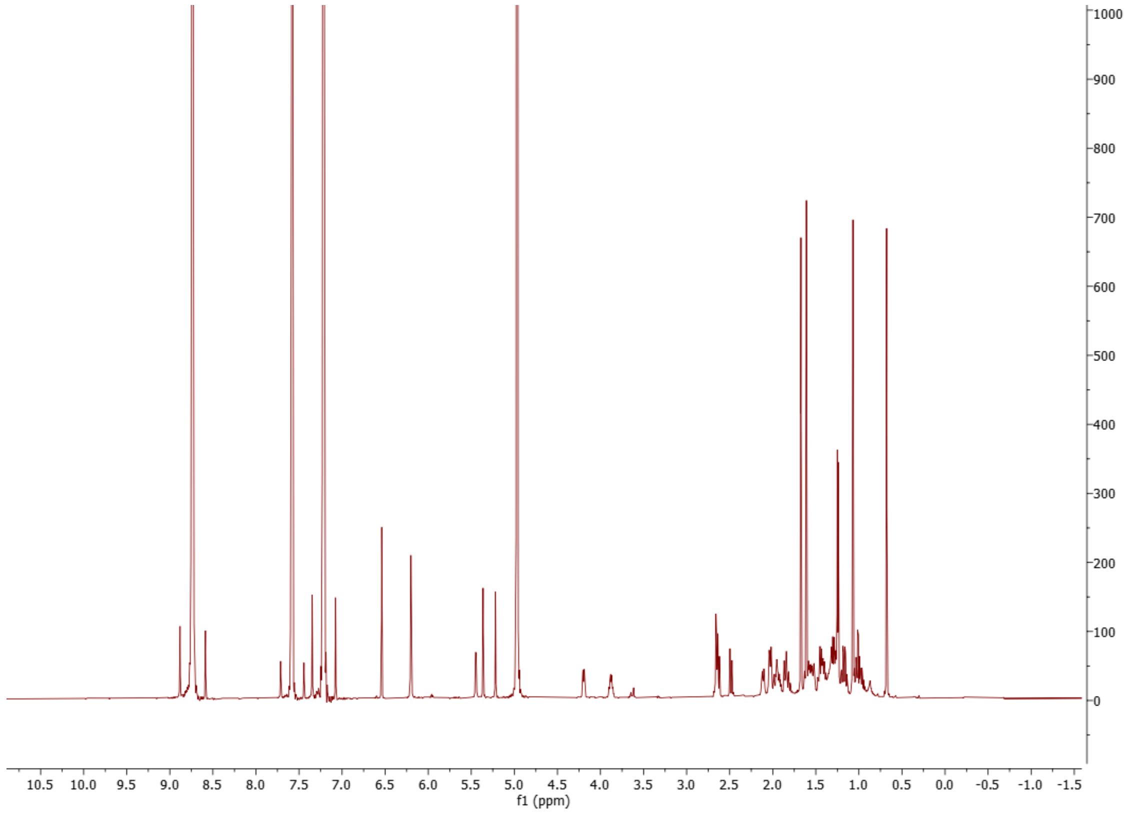


**Supplementary Fig. 44.** $^{1}$**H spectrum of (22*R*)-ergosta-5,24(28)-diene-3β,22,25-triol (phyministerol A) (12) (600 MHz,** $C_5D_5N$**, 298 K).**

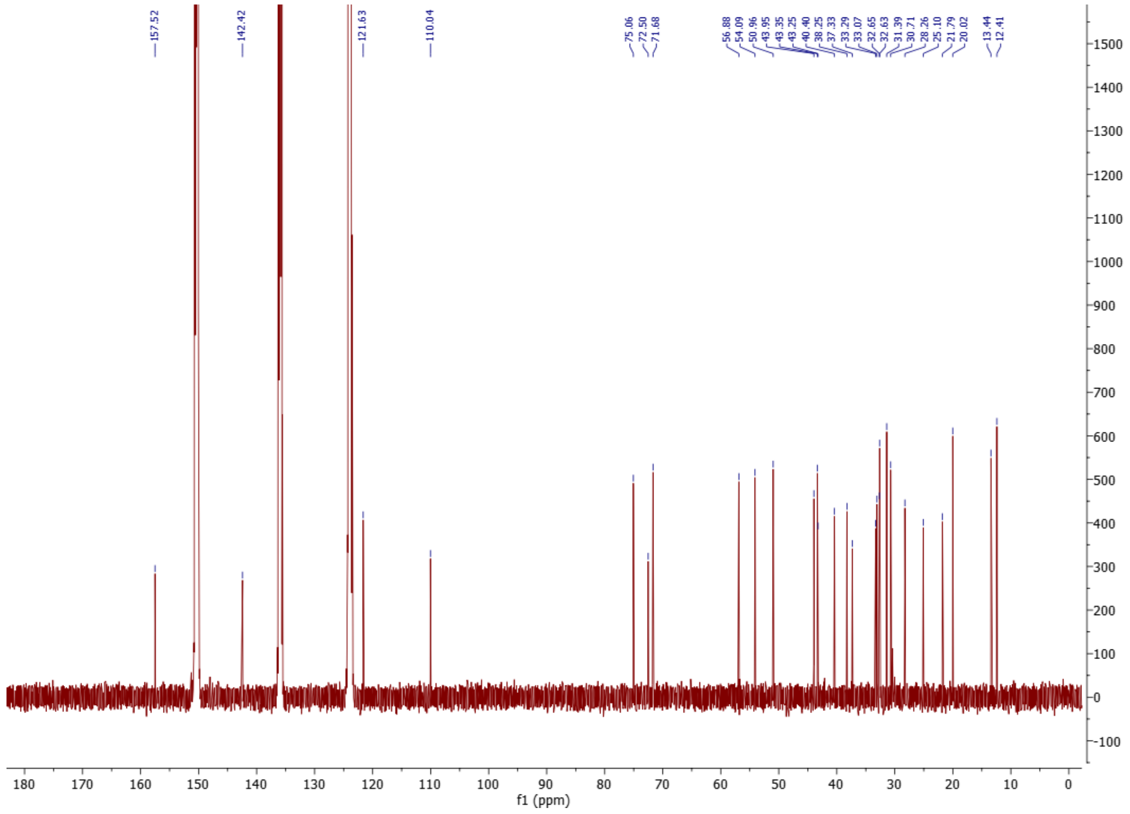


**Supplementary Fig. 45.** $^{13}$**C spectrum of (22*R*)-ergosta-5,24(28)-diene-3β,22,25-triol (phyministerol A) (12) (151 MHz,** $C_5D_5N$**, 298 K).**

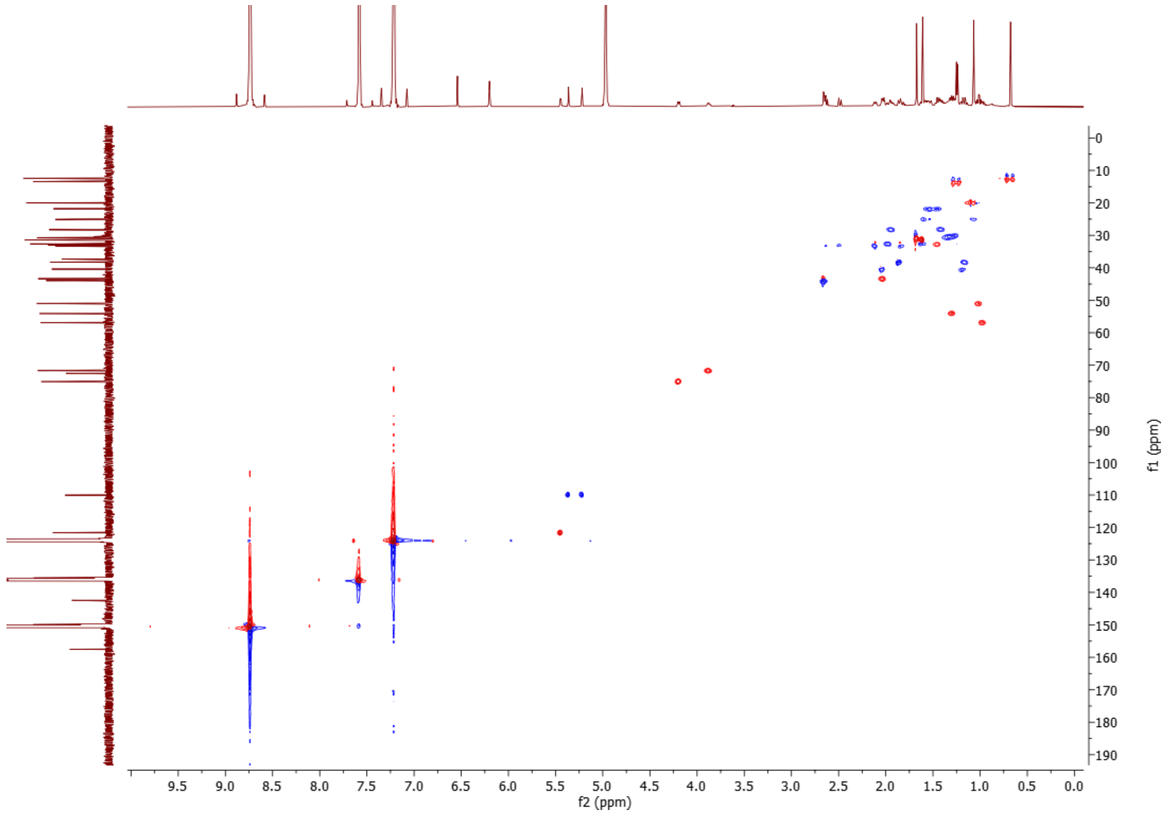


**Supplementary Fig. 46. HSQC spectrum of (22*R*)-ergosta-5,24(28)-diene-3β,22,25-triol (phyministerol A) (12) (600 MHz, $C_5D_5N$, 298 K).**

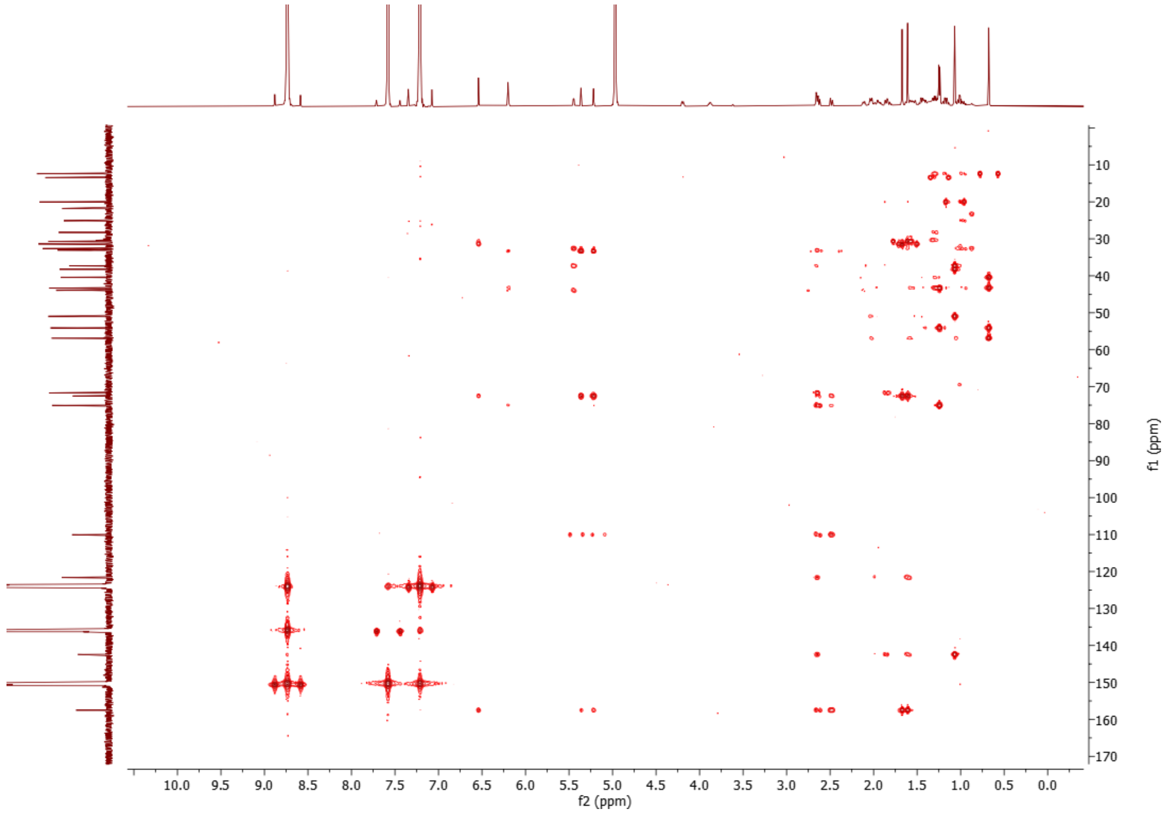


**Supplementary Fig. 47. HMBC spectrum of (22*R*)-ergosta-5,24(28)-diene-3β,22,25-triol (phyministerol A) (12) (600 MHz, $C_5D_5N$, 298 K).**

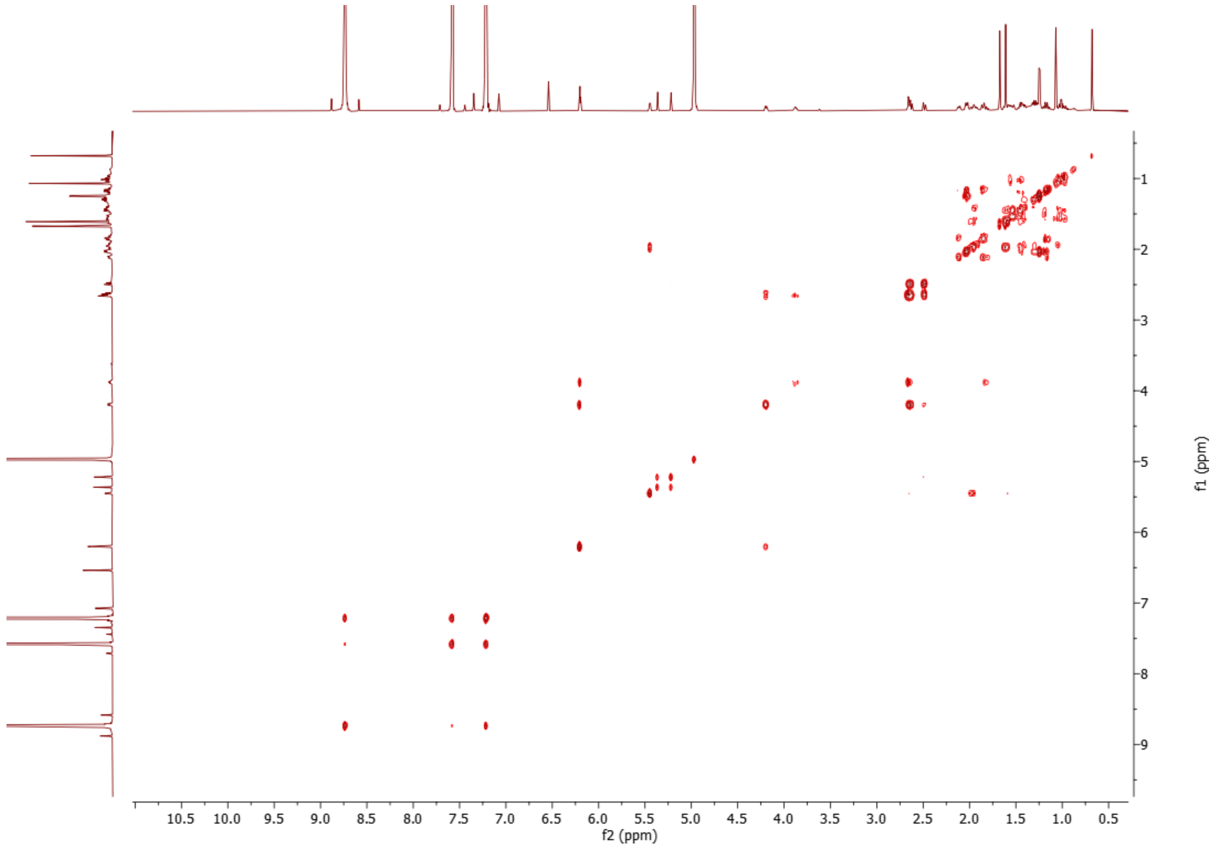


**Supplementary Fig. 48. COSY spectrum of (22*R*)-ergosta-5,24(28)-diene-3β,22,25-triol (phyministerol A) (12) (600 MHz, $C_5D_5N$, 298 K).**

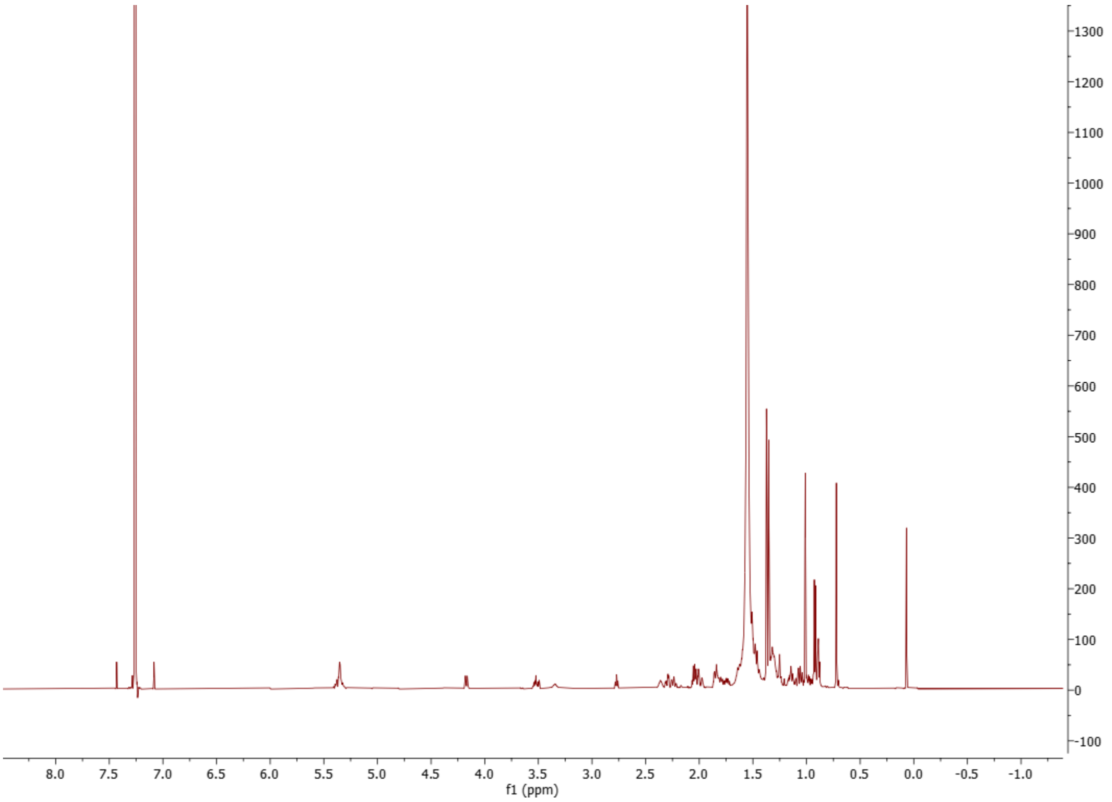


**Supplementary Fig. 49. $^{1}$H spectrum of (22*R*)-24,25-epoxy-ergost-5-ene-3β,22-diol (13) ($CDCl_3$, 298 K, 600 MHz).**

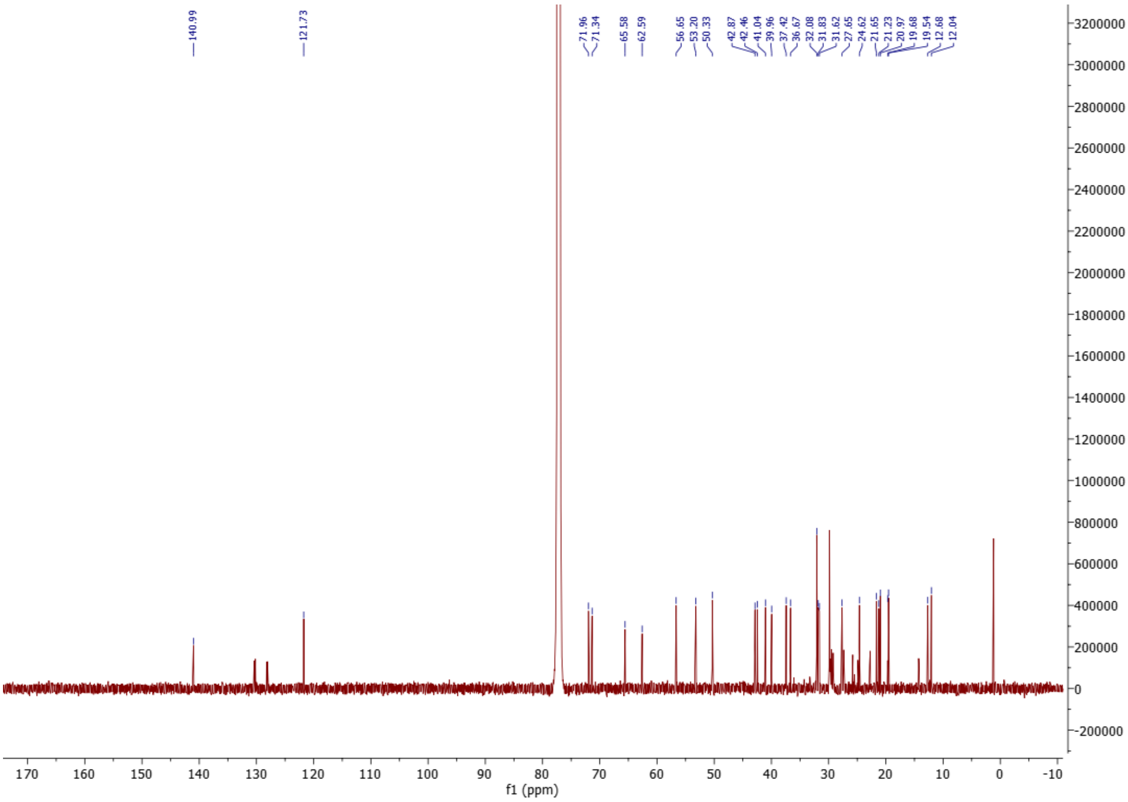


**Supplementary Fig. 50. $^{13}$C spectrum of (22*R*)-24,25-epoxy-ergost-5-ene-3β,22-diol (13) ($CDCl_3$, 298 K, 151 MHz).**

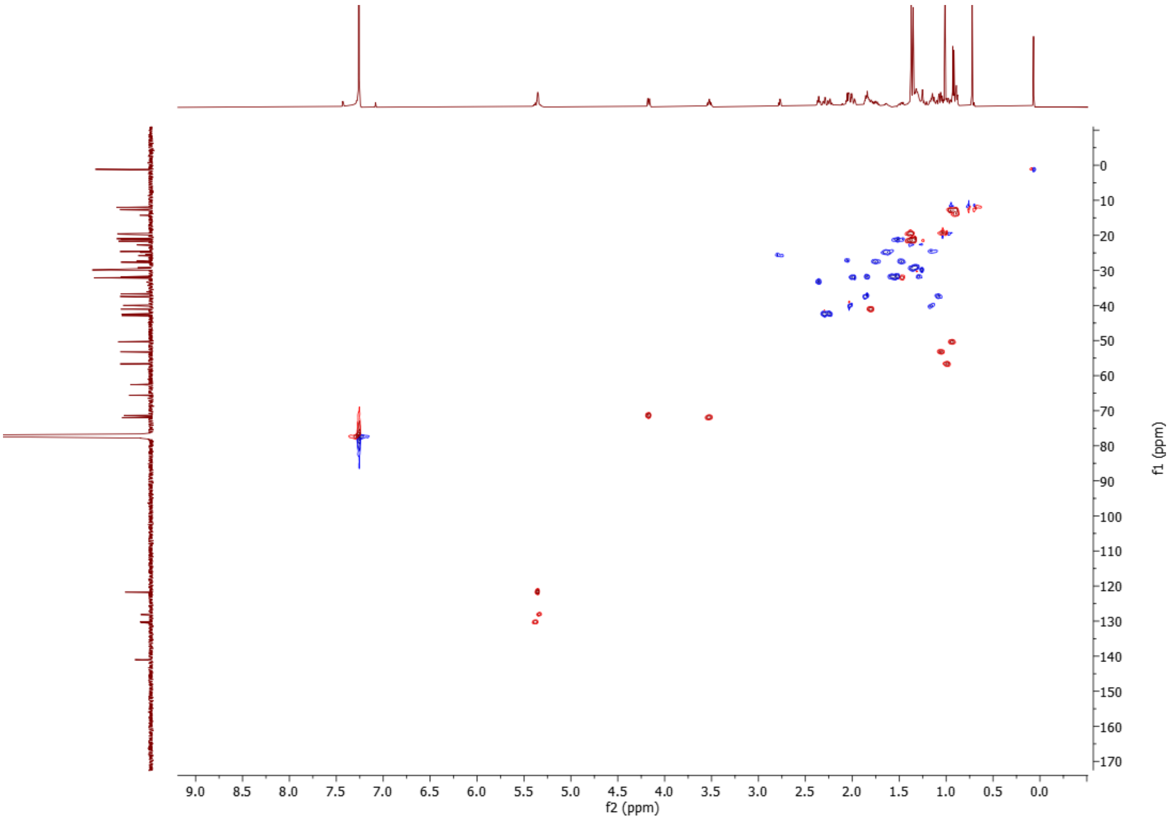


**Supplementary Fig. 51. HSQC spectrum of (22*R*)-24,25-epoxy-ergost-5-ene-3β,22-diol (13) ($CDCl_3$, 298 K, 600 MHz).**

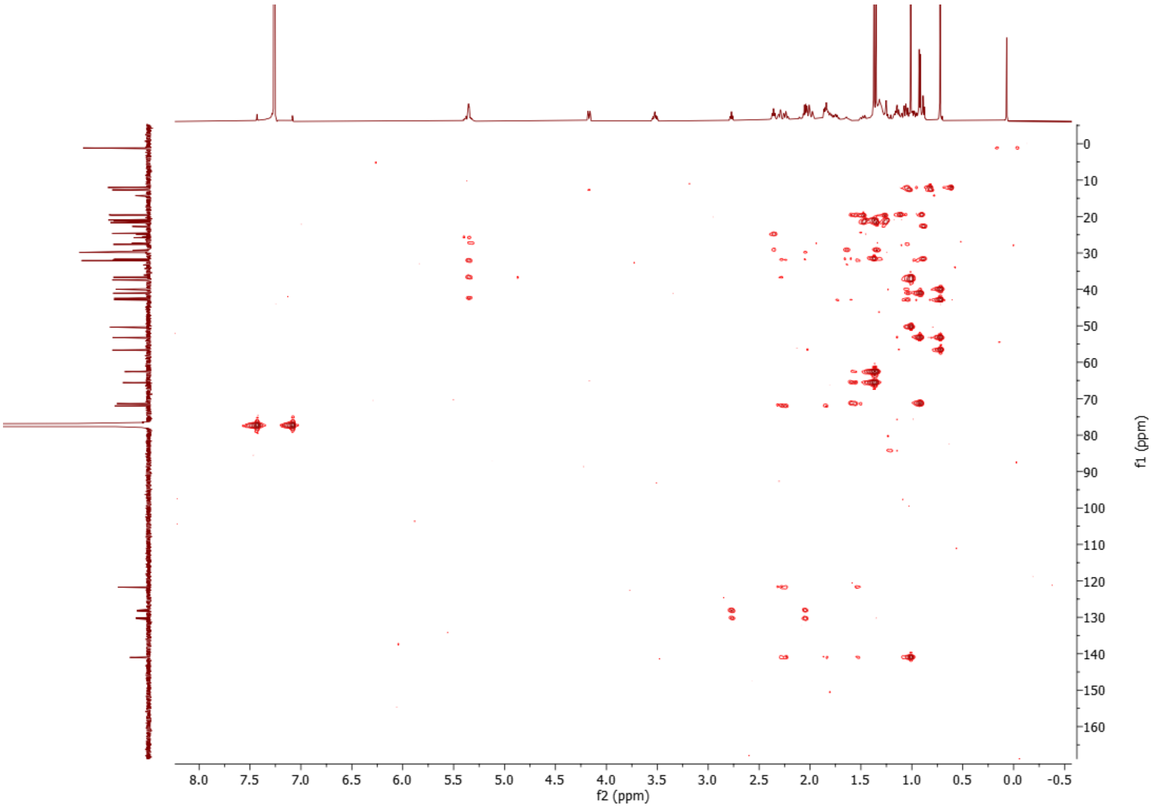


**Supplementary Fig. 52. HMBC spectrum of (22*R*)-24,25-epoxy-ergost-5-ene-3β,22-diol (13) ($CDCl_3$, 298 K, 600 MHz).**

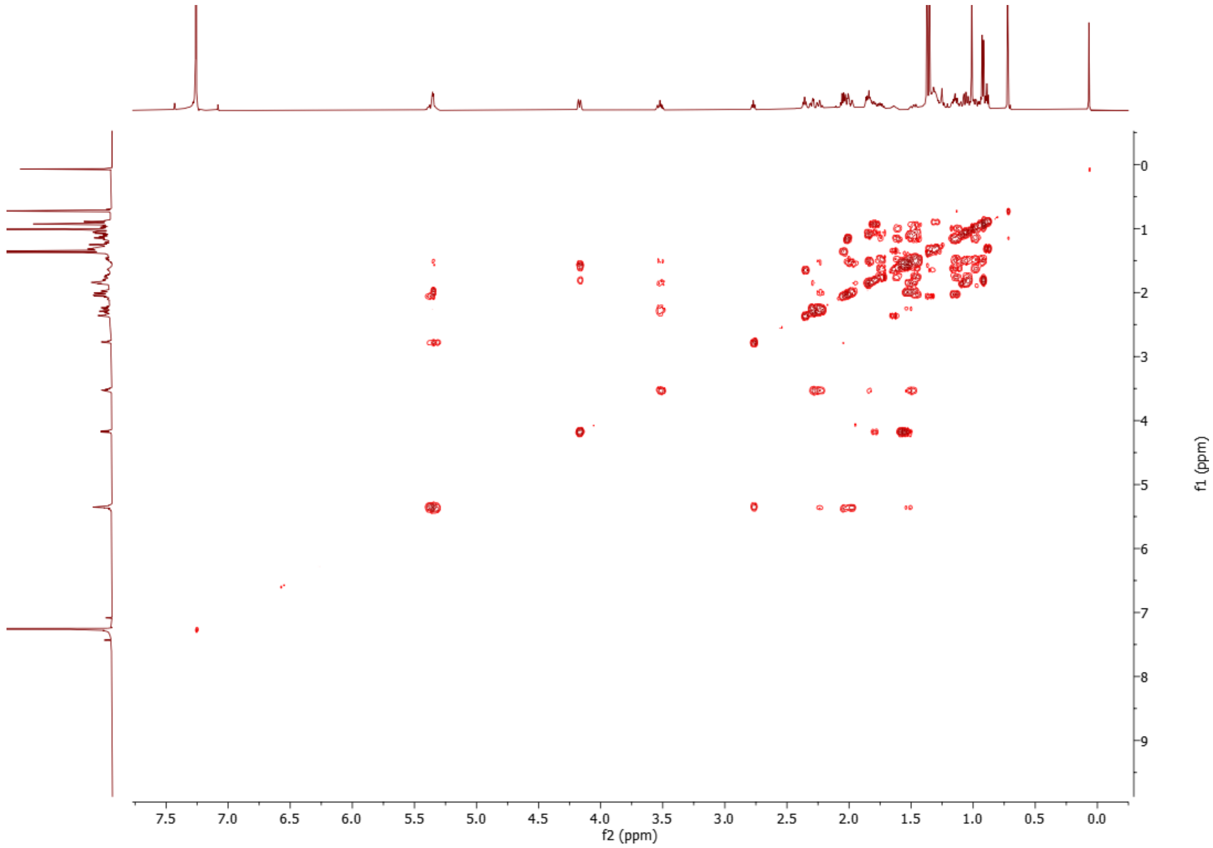


**Supplementary Fig. 53. COSY spectrum of (22*R*)-24,25-epoxy-ergost-5-ene-3β,22-diol (13) ($CDCl_3$, 298 K, 600 MHz).**

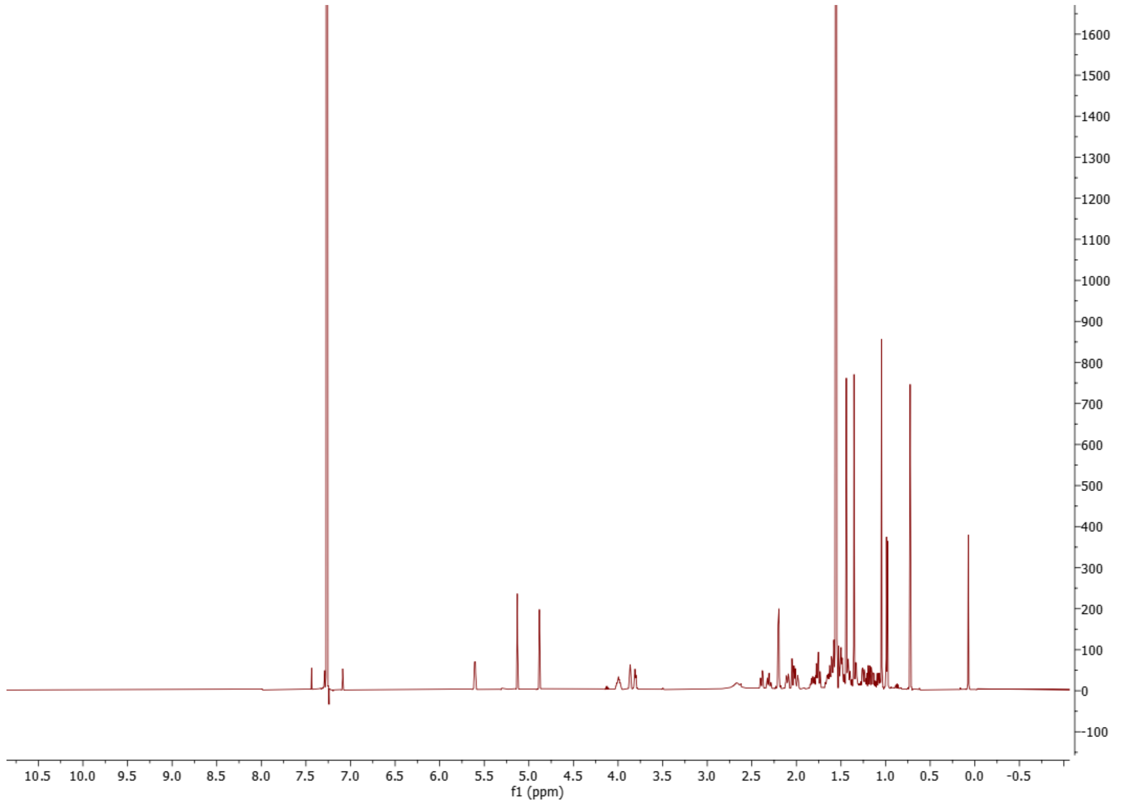


Supplementary Fig. 54. $^{1}H$ spectrum of (22*R*)-ergosta-5,24(28)-diene-1α,3β,22,25-tetraol (15) ($CDCl_3$, 298 K, 600 MHz).

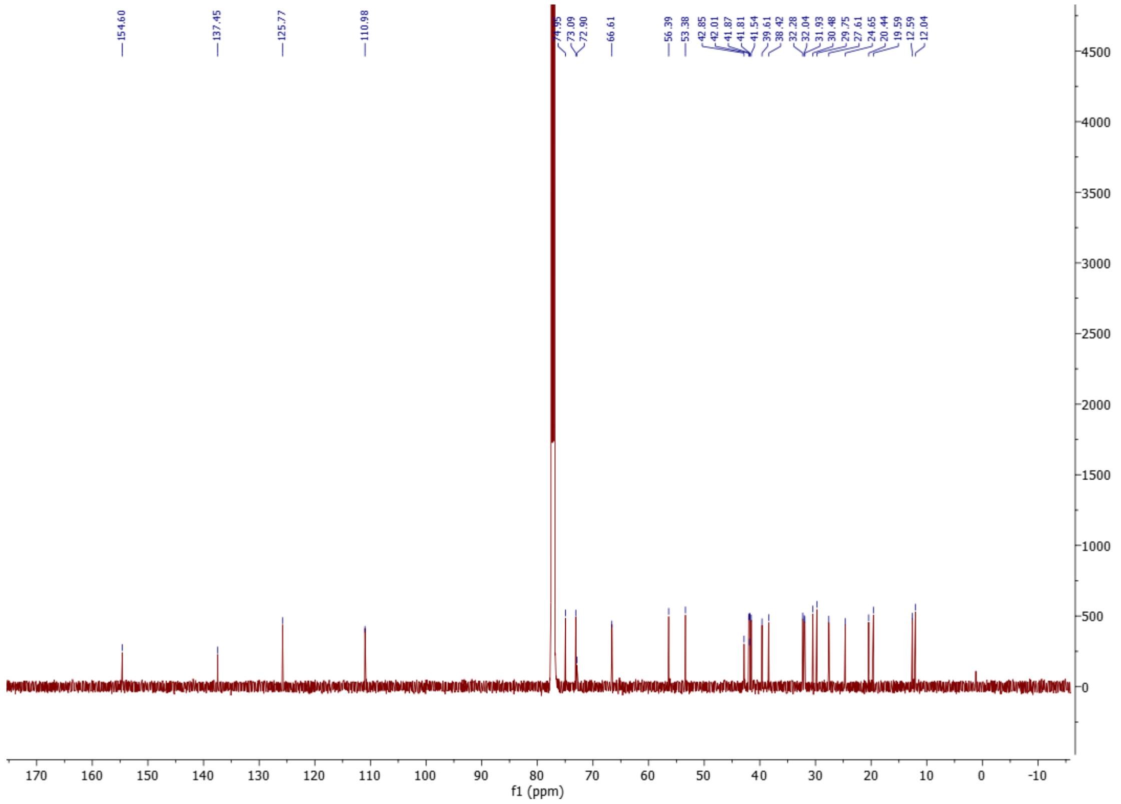


Supplementary Fig. 55. $^{13}C$ spectrum of (22*R*)-ergosta-5,24(28)-diene-1α,3β,22,25-tetraol (15) ($CDCl_3$, 298 K, 151 MHz).

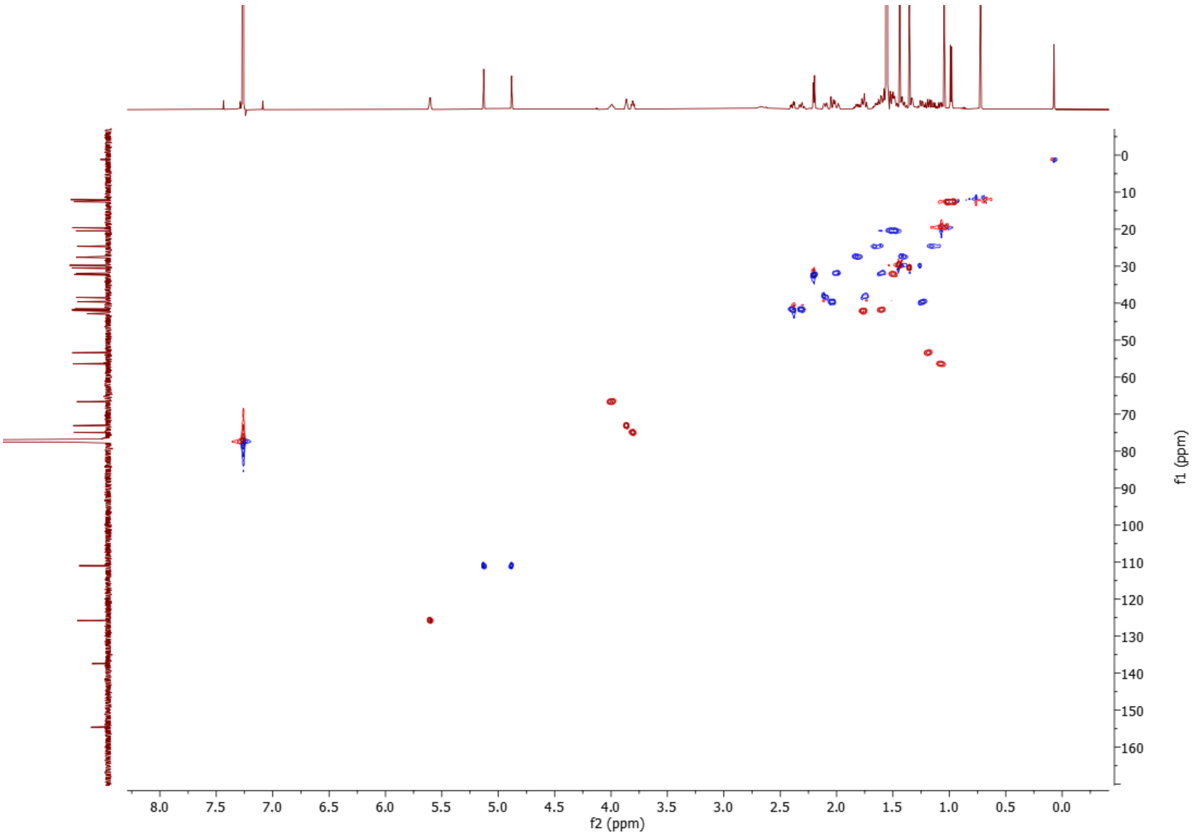


**Supplementary Fig. 56. HSQC spectrum of (22*R*)-ergosta-5,24(28)-diene-1α,3β,22,25-tetraol (15) ($CDCl_3$, 298 K, 600 MHz).**

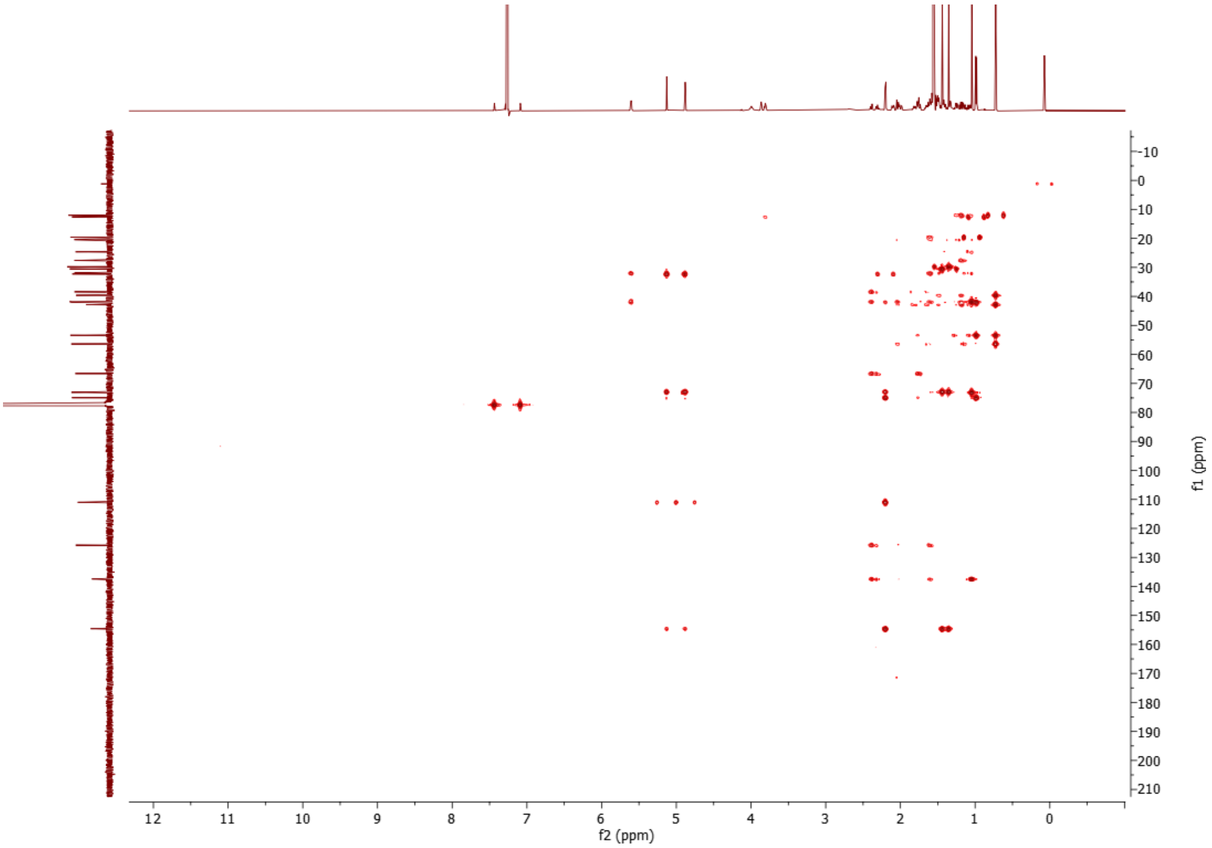


**Supplementary Fig. 57. HMBC spectrum of (22*R*)-ergosta-5,24(28)-diene-1α,3β,22,25-tetraol (15) ($CDCl_3$, 298 K, 600 MHz).**

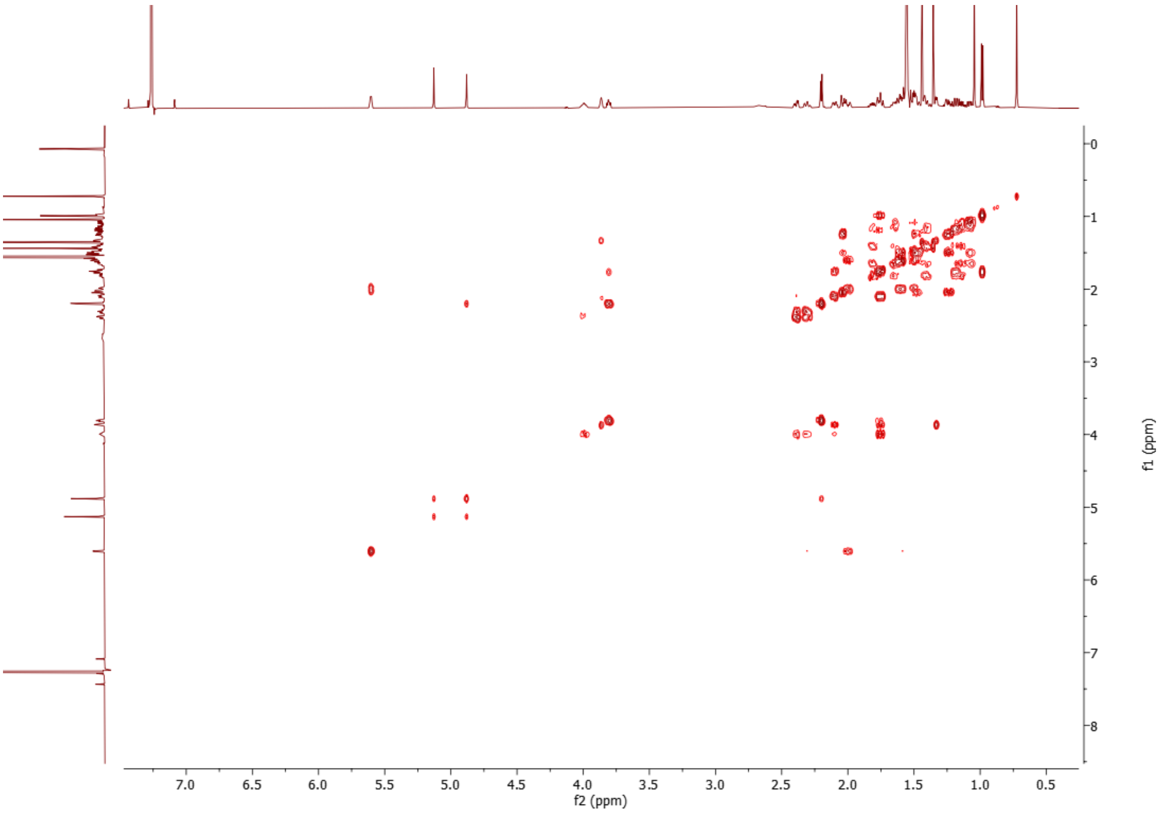


**Supplementary Fig. 58. COSY spectrum of (22*R*)-ergosta-5,24(28)-diene-1α,3β,22,25-tetraol (15) ($CDCl_3$, 298 K, 600 MHz).**

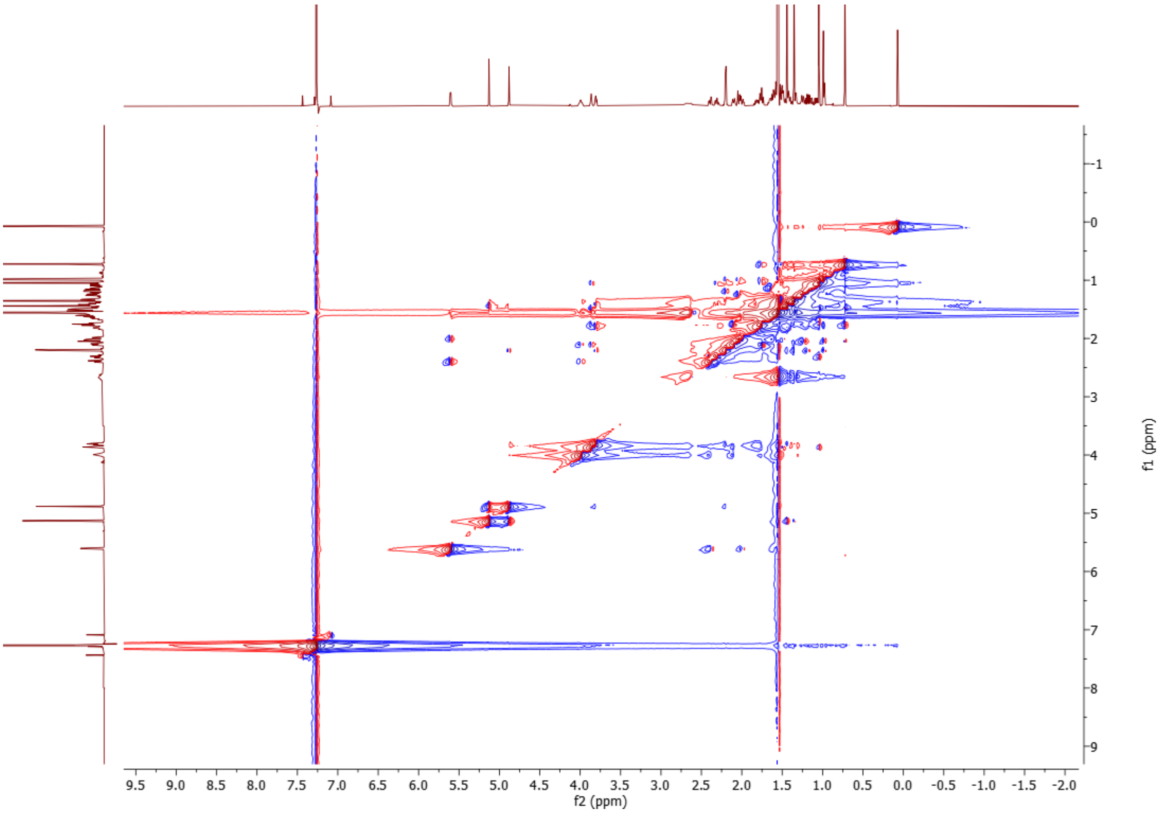


**Supplementary Fig. 59. NOESY spectrum of (22*R*)-ergosta-5,24(28)-diene-1α,3β,22,25-tetraol (15) ($CDCl_3$, 298 K, 600 MHz).**

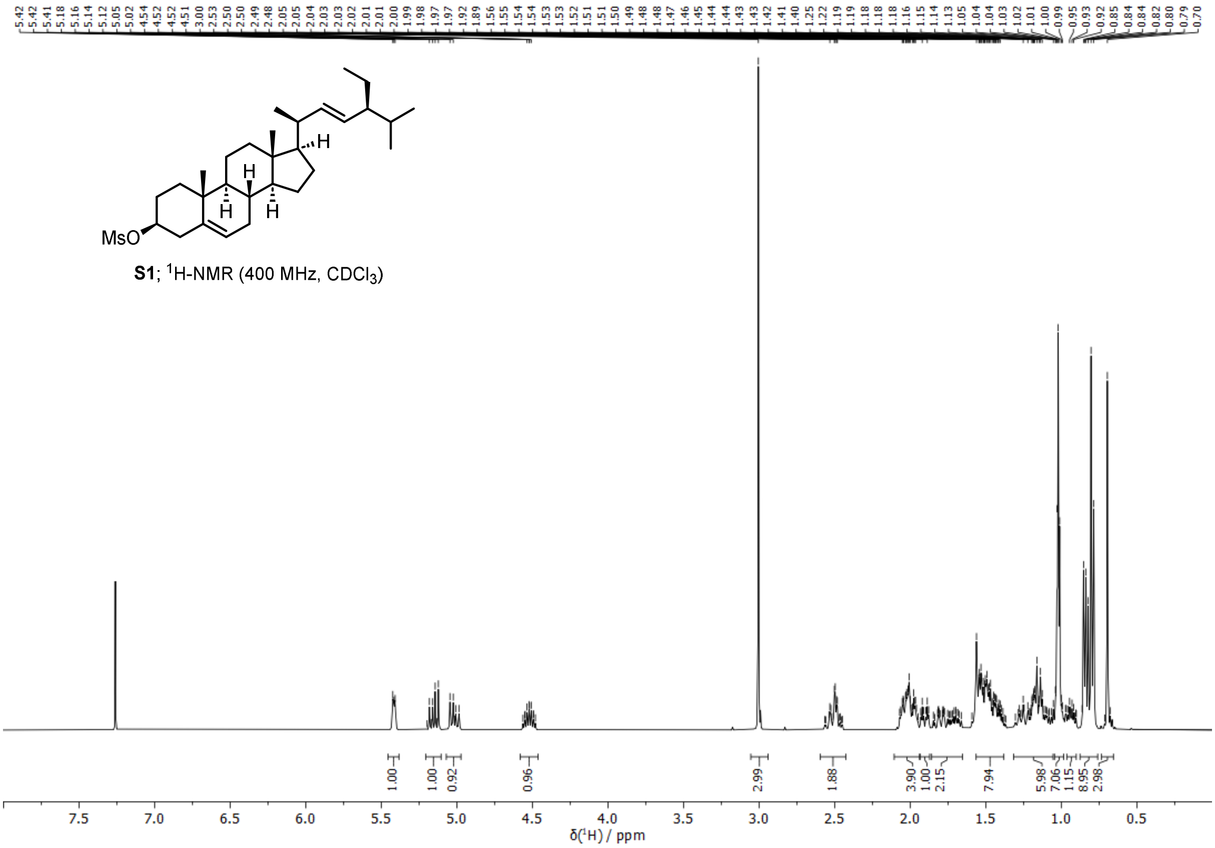


**Supplementary Fig. 60.** $^{1}$**H spectrum of stigmasteryl mesylate (S1) (400 MHz, CDCl$_3$, 298 K).**

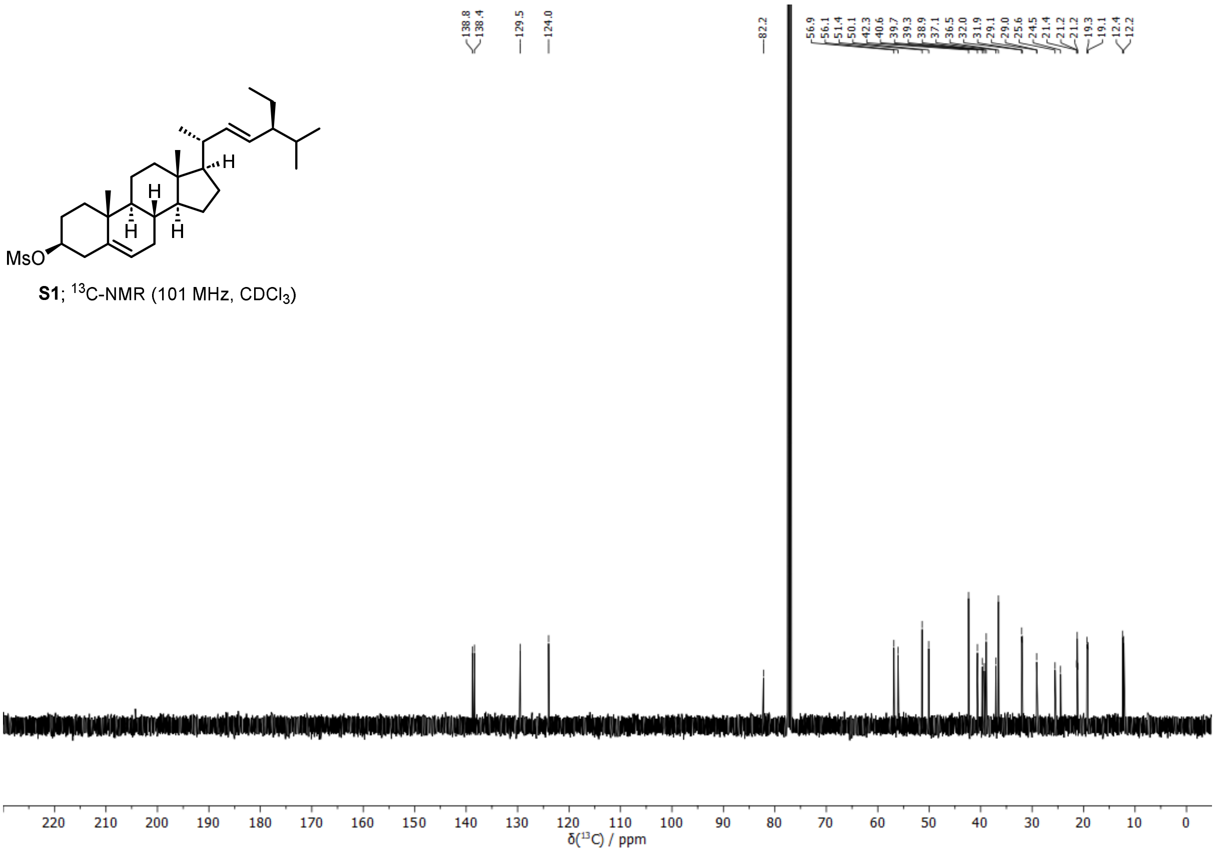


**Supplementary Fig. 61.** $^{13}$**C spectrum of stigmasteryl mesylate (S1) (101 MHz, CDCl$_3$, 298 K).**

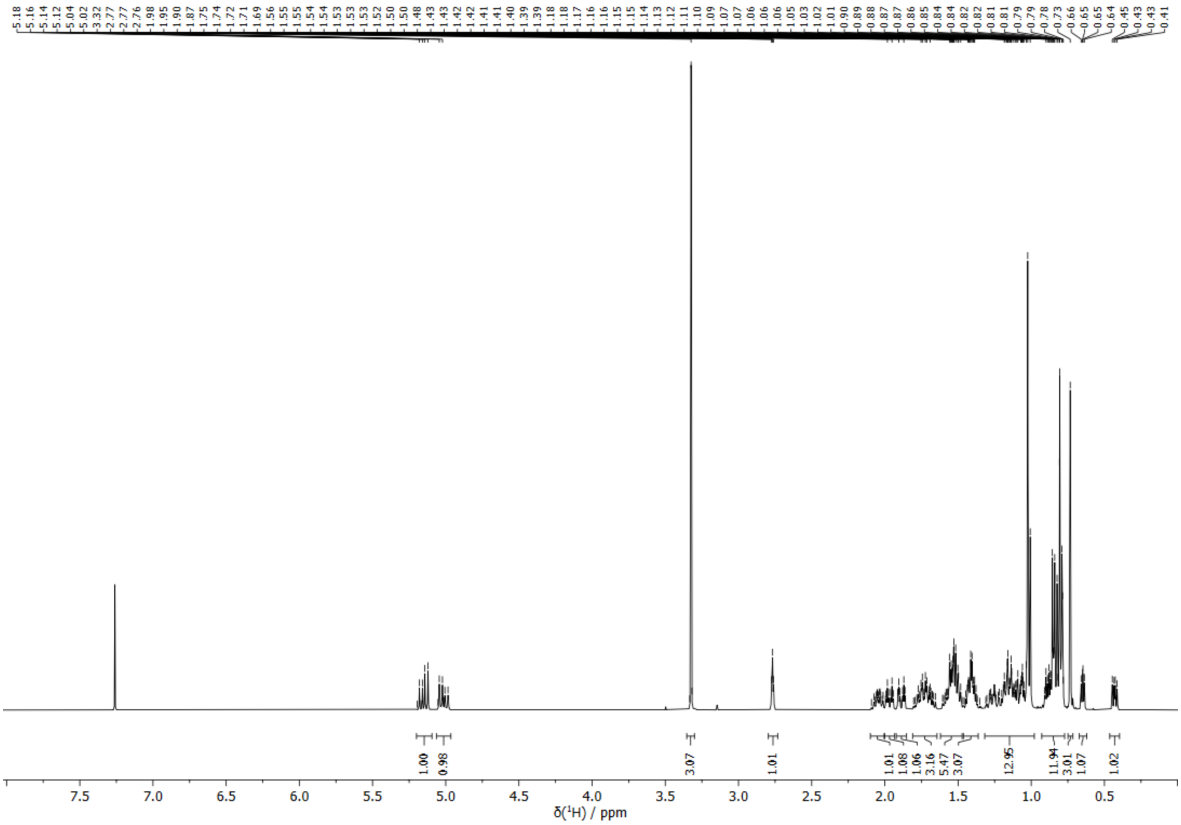


**Supplementary Fig. 62. $^{1}$H spectrum of *i*-stigmasteryl methyl ether (S2) (400 MHz, $CDCl_3$, 298 K).**

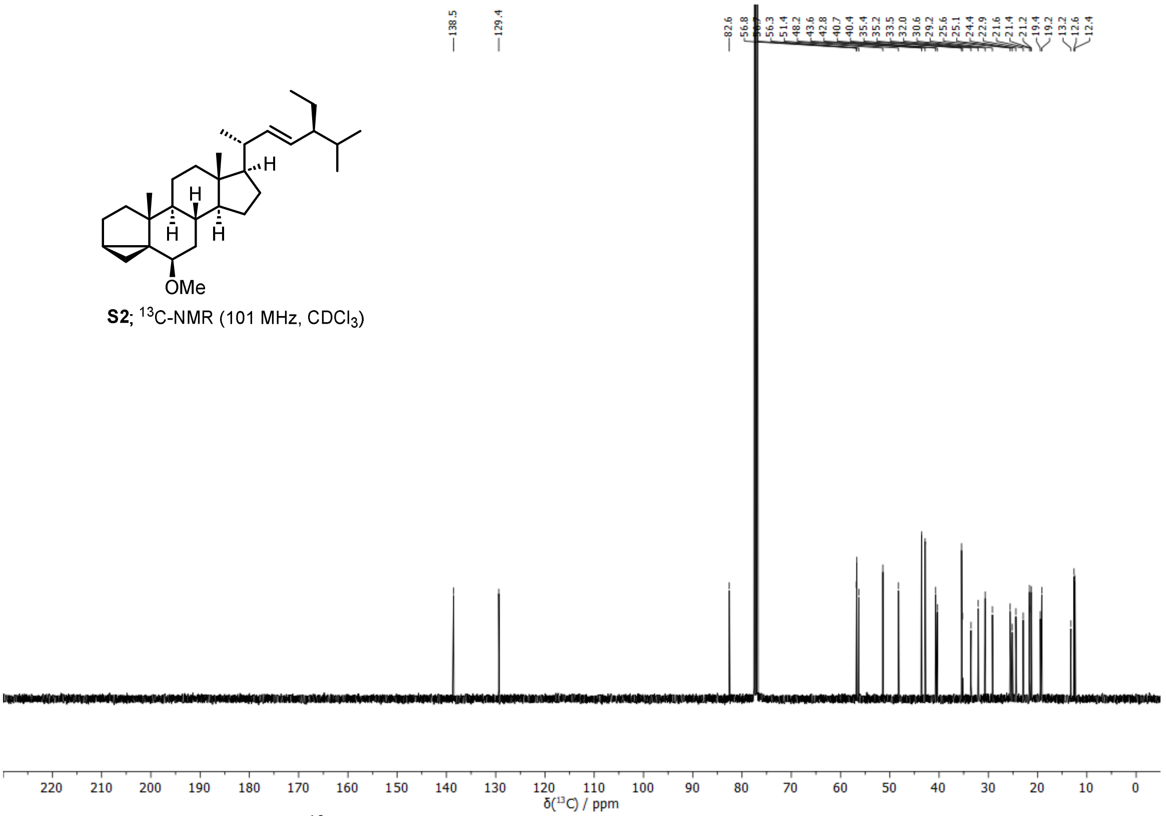


**Supplementary Fig. 63. $^{13}$C spectrum of *i*-stigmasteryl methyl ether (S2) (101 MHz, $CDCl_3$, 298 K).**

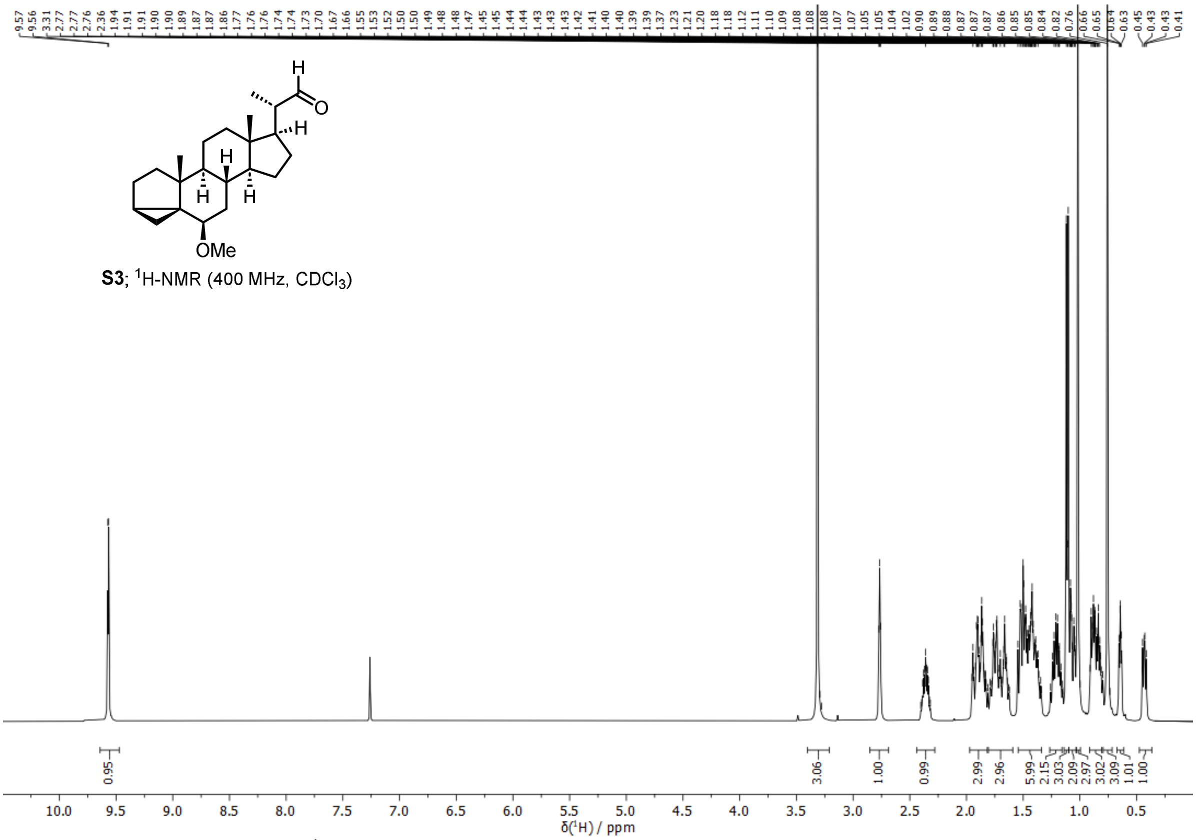


**Supplementary Fig. 64. ¹H spectrum of (20*S*)-6*β*-methoxy-3*α*,5-cyclopregnancarbaldehyde (S3) (400 MHz, $CDCl_3$, 298 K).**

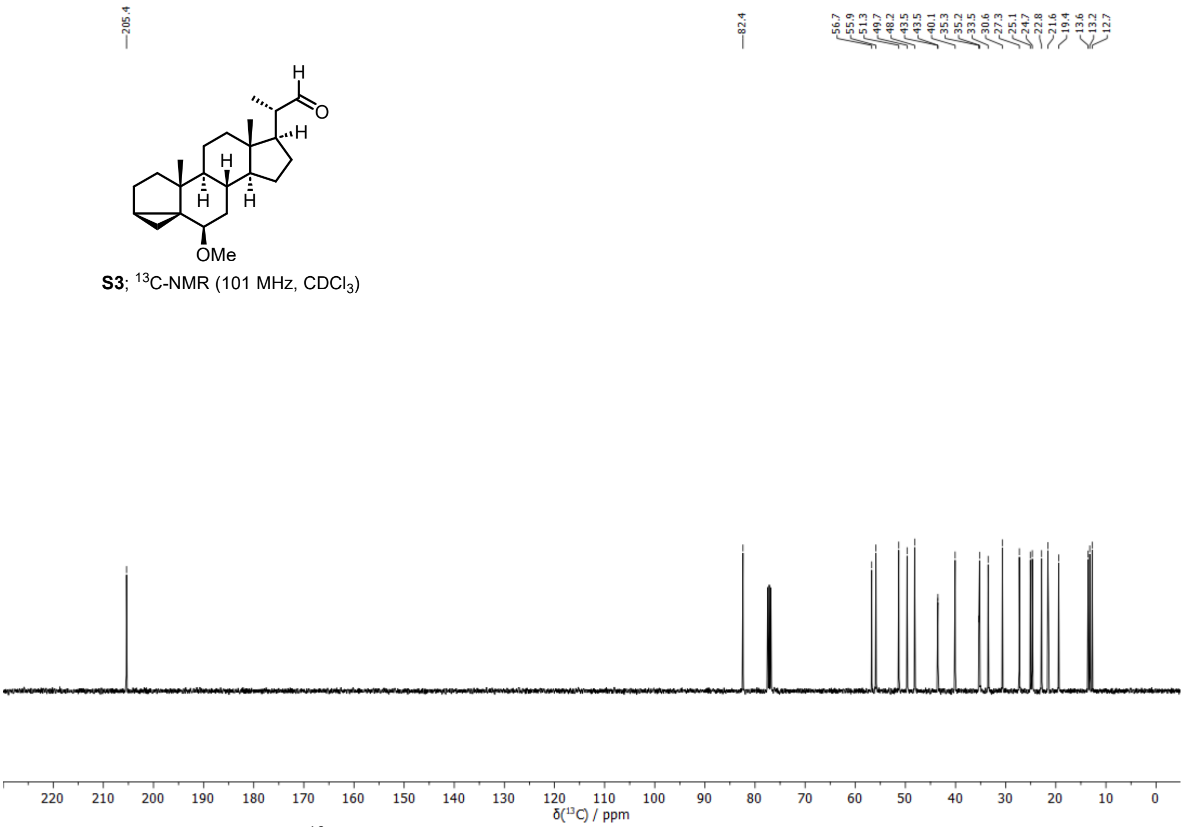


**Supplementary Fig. 65. ¹³C spectrum of (20*S*)-6*β*-methoxy-3*α*,5-cyclopregnancarbaldehyde (S3) (101 MHz, $CDCl_3$, 298 K).**

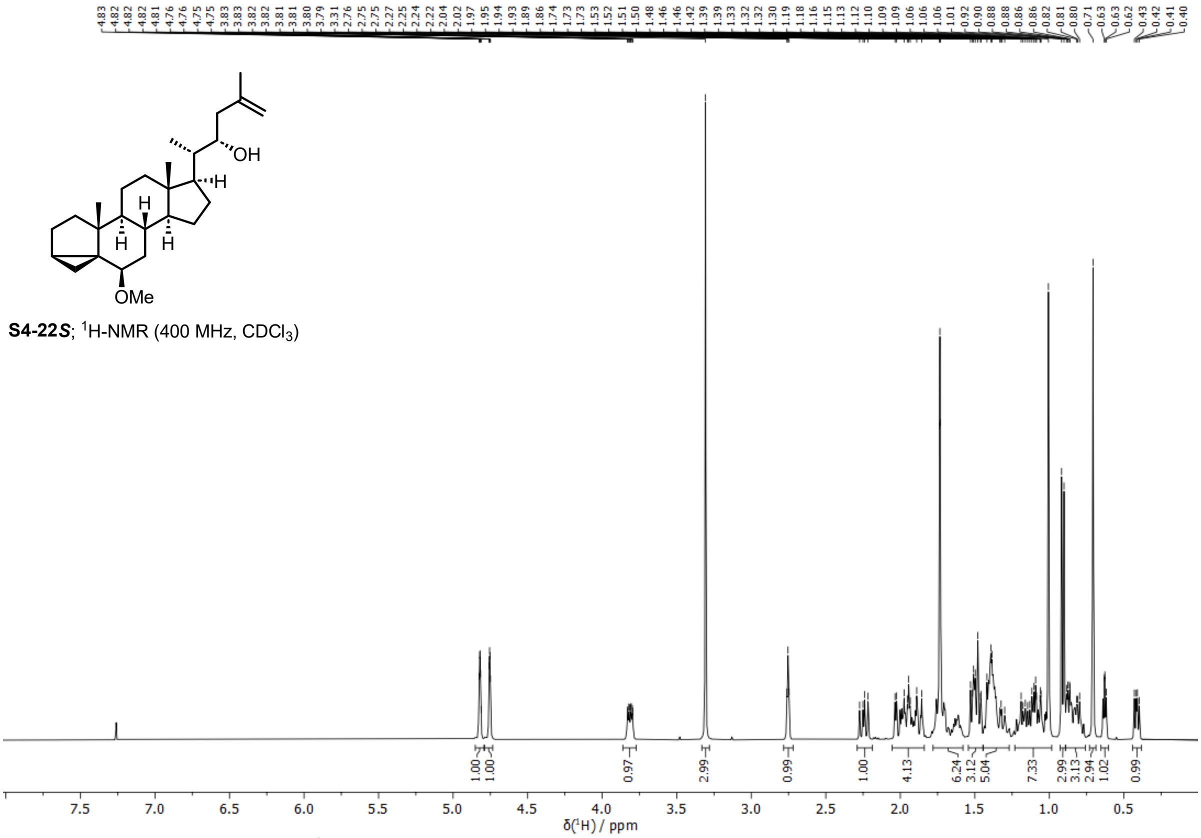


**Supplementary Fig. 66. $^{1}$H spectrum of (22*S*)-24-nor-6*β*-methoxy-3*α*,5-cyclocholest-25-en-22-ol (S4-(22*S*)) (400 MHz, CDCl$_3$, 298 K).**

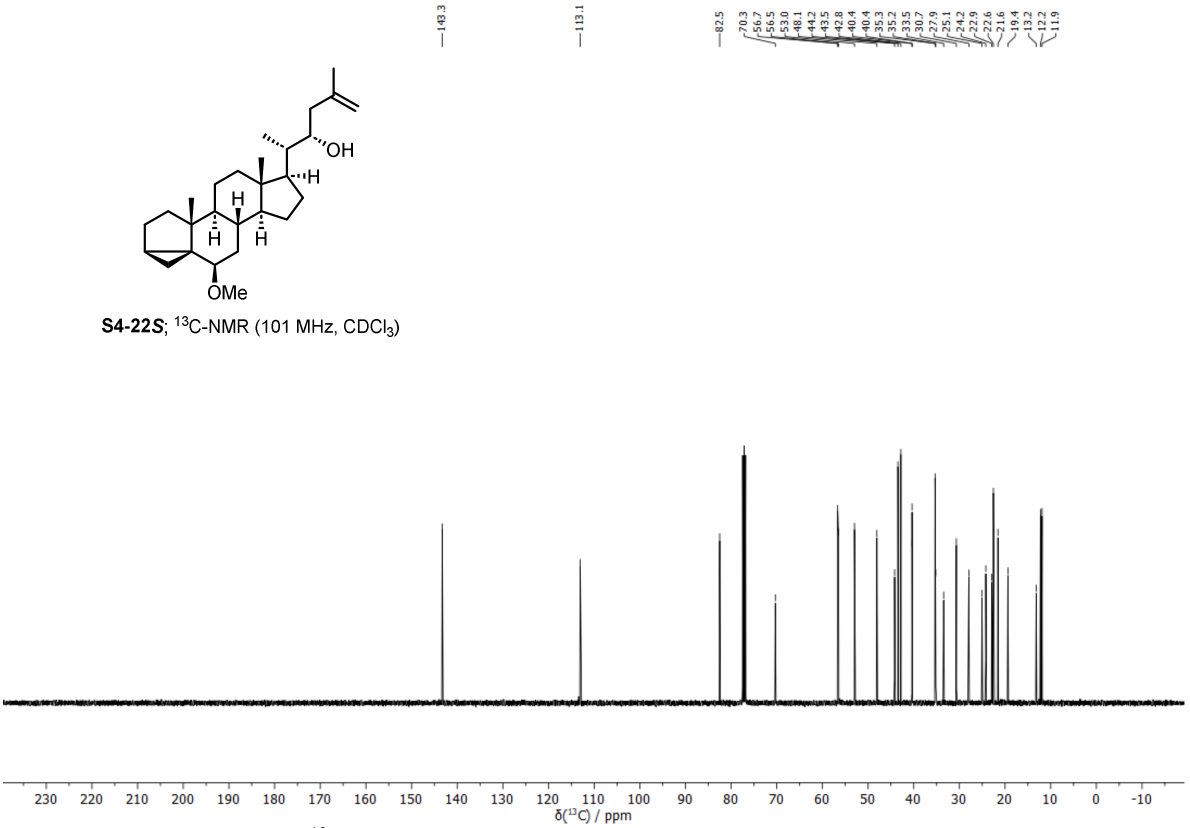


**Supplementary Fig. 67. $^{13}$C spectrum of (22*S*)-24-nor-6*β*-methoxy-3*α*,5-cyclocholest-25-en-22-ol (S4-(22*S*)) (101 MHz, CDCl$_3$, 298 K).**

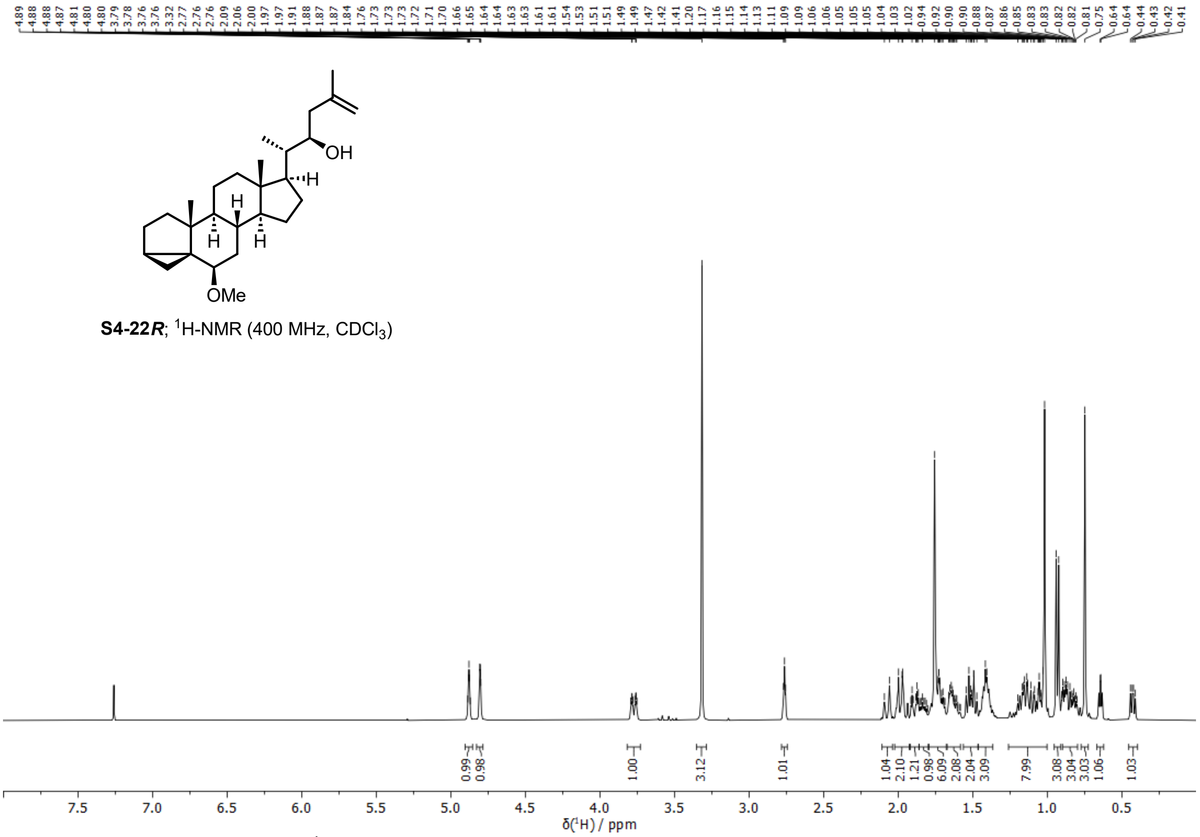


**Supplementary Fig. 68. $^{1}$H spectrum of (22*R*)-24-nor-6*β*-methoxy-3*α*,5-cyclocholest-25-en-22-ol (S4-(22*R*)) (400 MHz, $CDCl_3$, 298 K).**

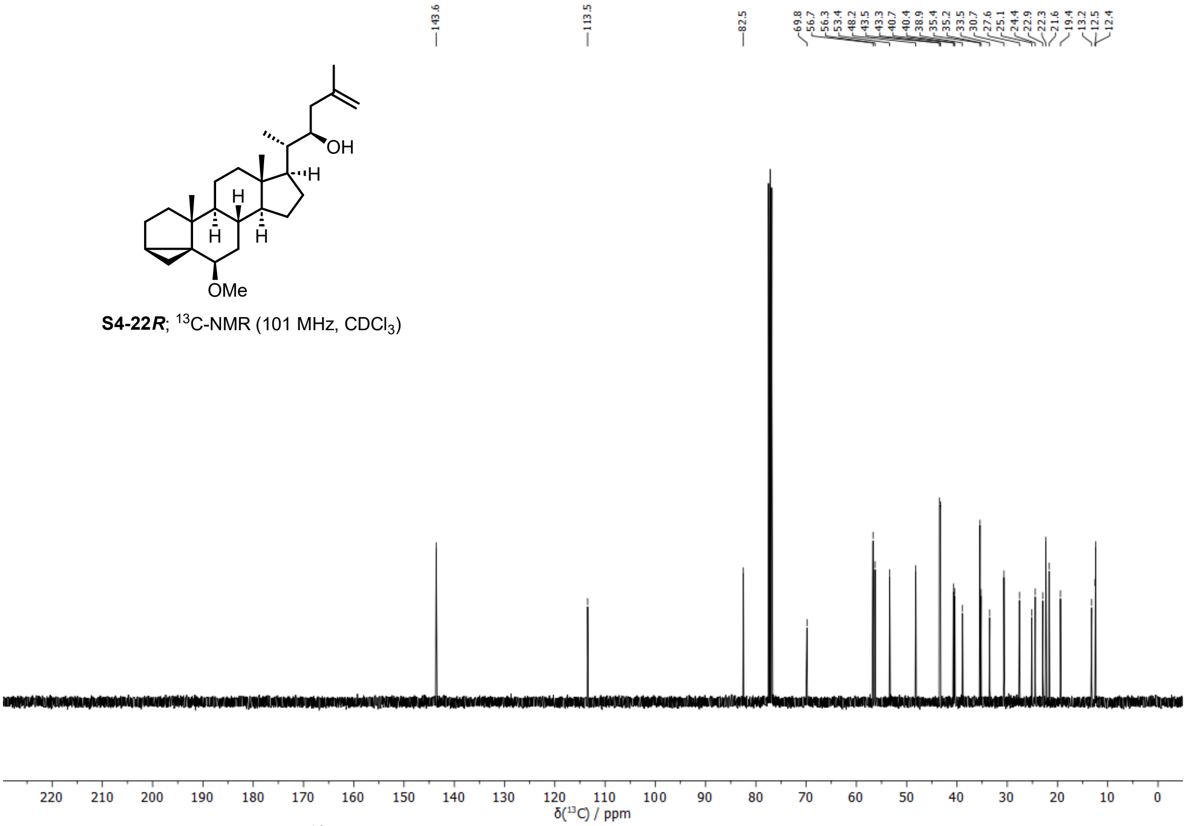


**Supplementary Fig. 69. $^{13}$C spectrum of (22*R*)-24-nor-6*β*-methoxy-3*α*,5-cyclocholest-25-en-22-ol (S4-(22*R*)) (101 MHz, $CDCl_3$, 298 K).**

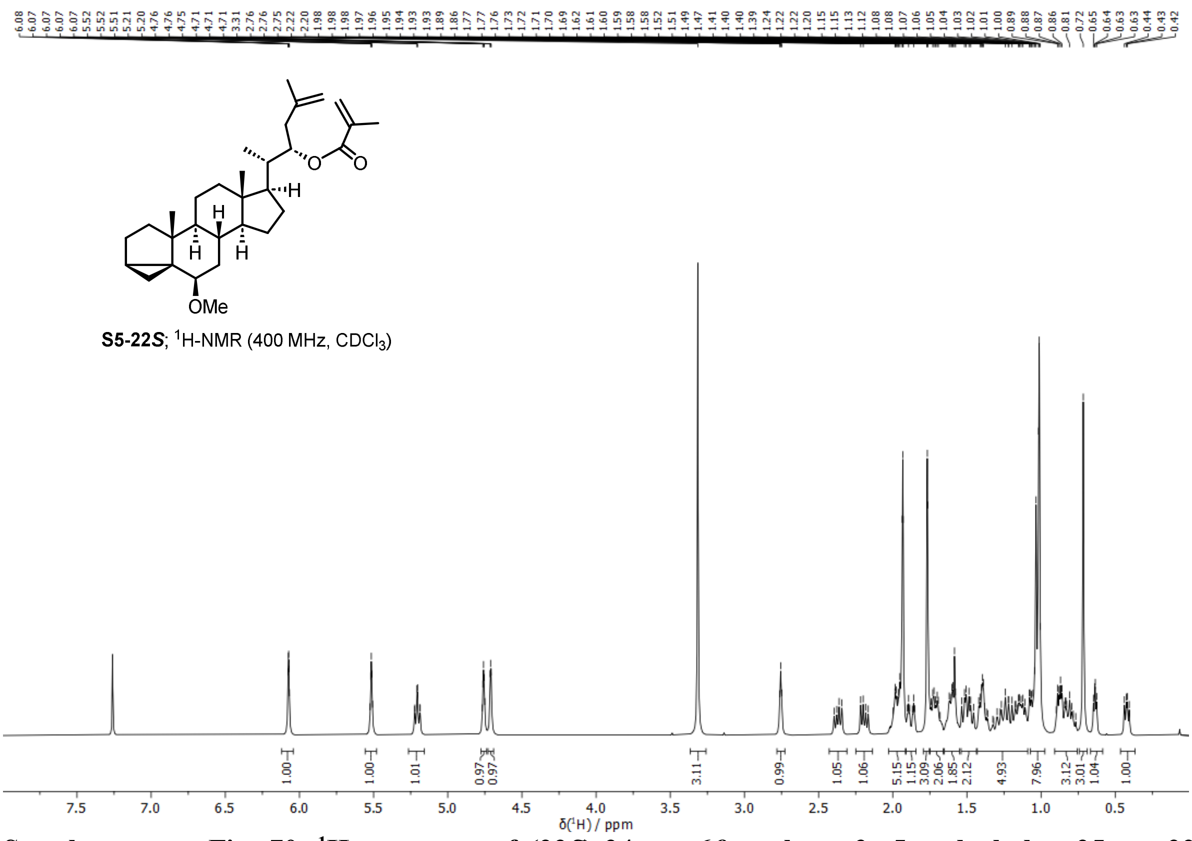


**Supplementary Fig. 70. [1]H spectrum of (22*S*)-24-nor-6*β*-methoxy-3*α*,5-cyclocholest-25-ene-22-methacrylate (S5-(22*S*)) (400 MHz, $CDCl_3$, 298 K).**

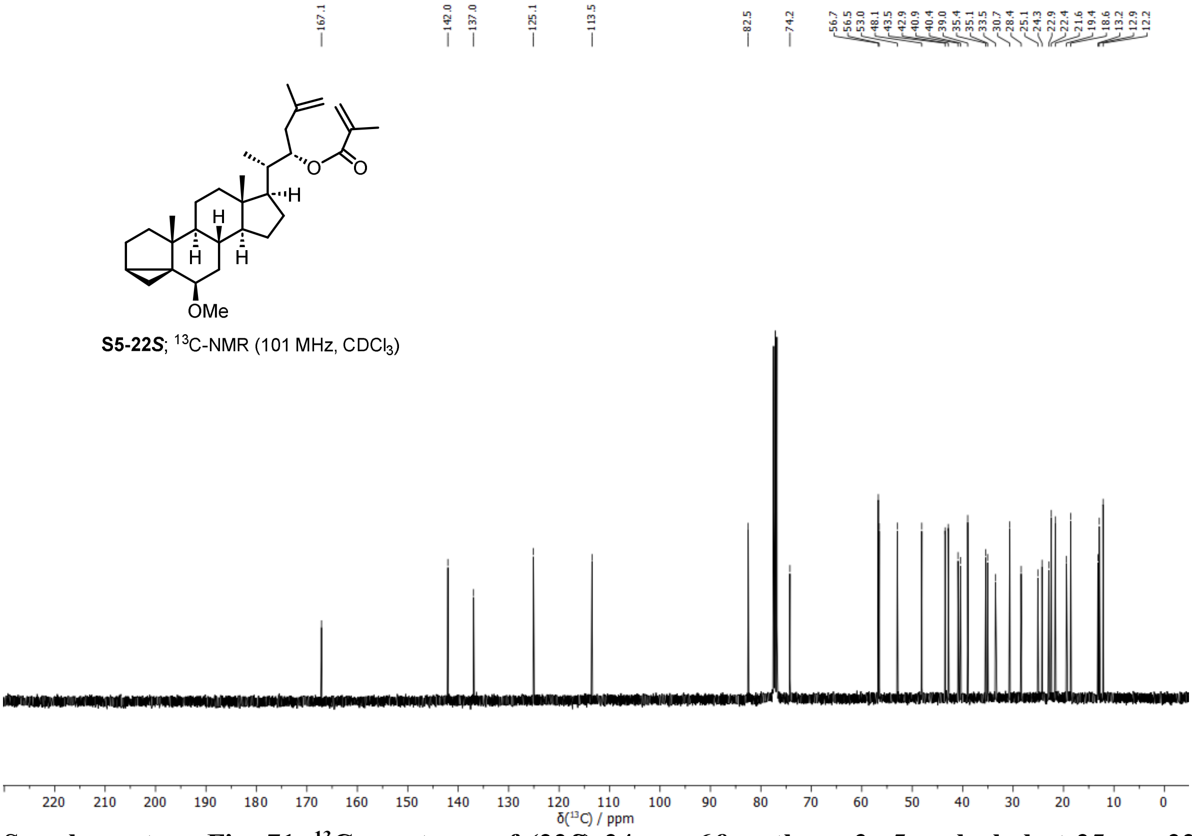


**Supplementary Fig. 71. [13]C spectrum of (22*S*)-24-nor-6*β*-methoxy-3*α*,5-cyclocholest-25-ene-22-methacrylate (S5-(22*S*)) (101 MHz, $CDCl_3$, 298 K).**

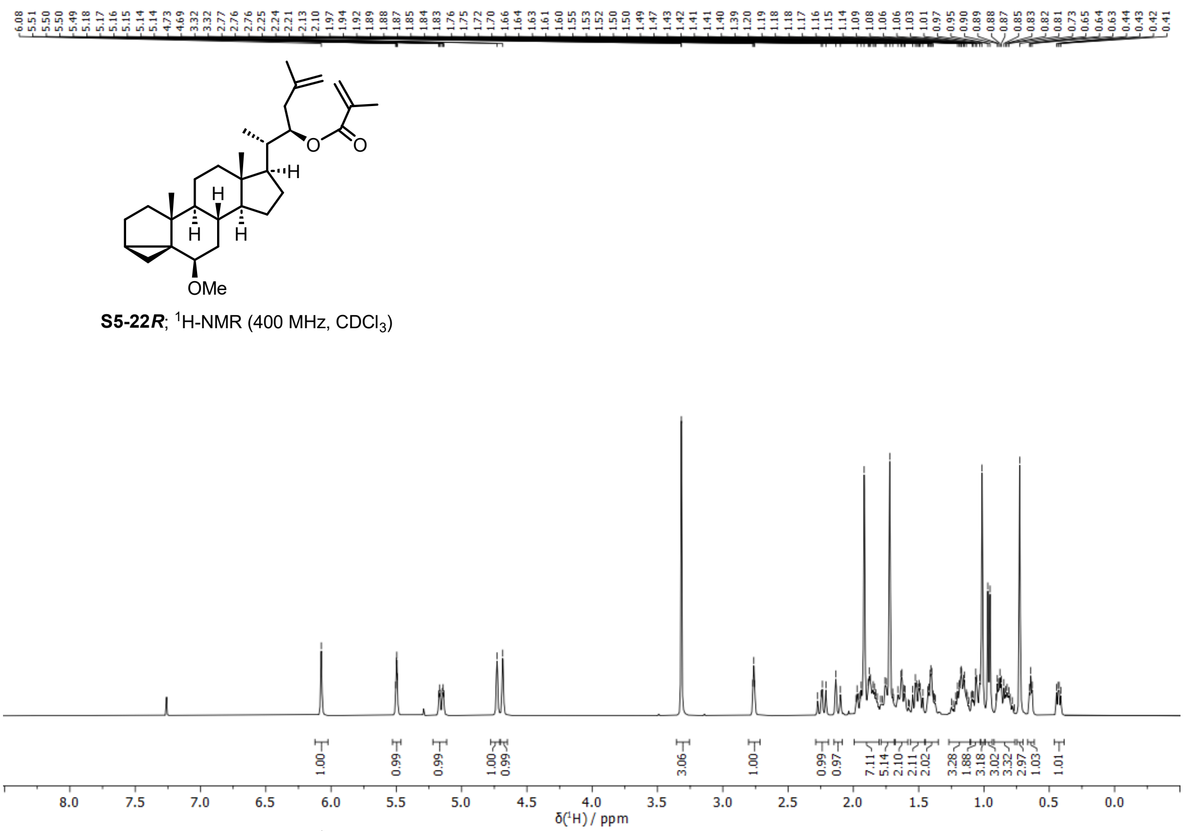


**Supplementary Fig. 72. $^1$H spectrum of (22*R*)-24-nor-6*β*-methoxy-3*α*,5-cyclocholest-25-ene-22-methacrylate (S5-(22*R*)) (400 MHz, $CDCl_3$, 298 K).**

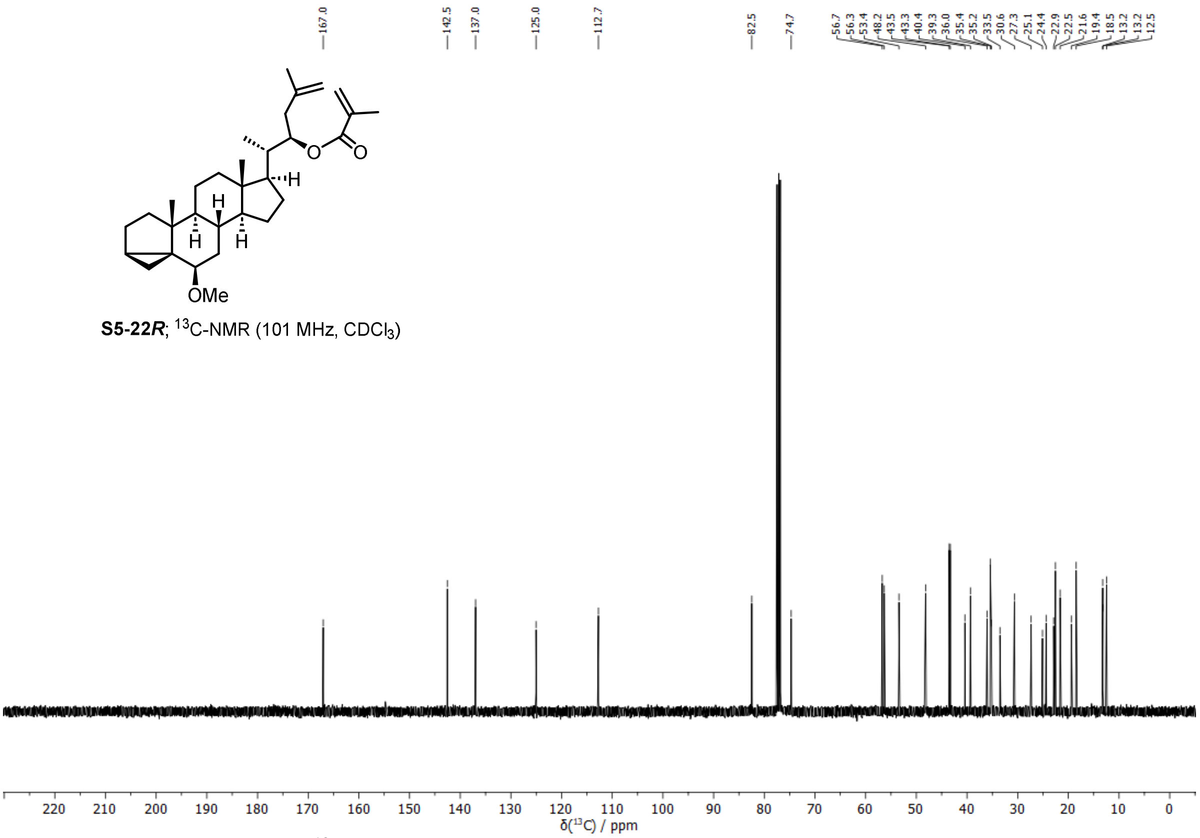


**Supplementary Fig. 73. $^{13}$C spectrum of (22*R*)-24-nor-6*β*-methoxy-3*α*,5-cyclocholest-25-ene-22-methacrylate (S5-(22*R*)) (101 MHz, $CDCl_3$, 298 K).**

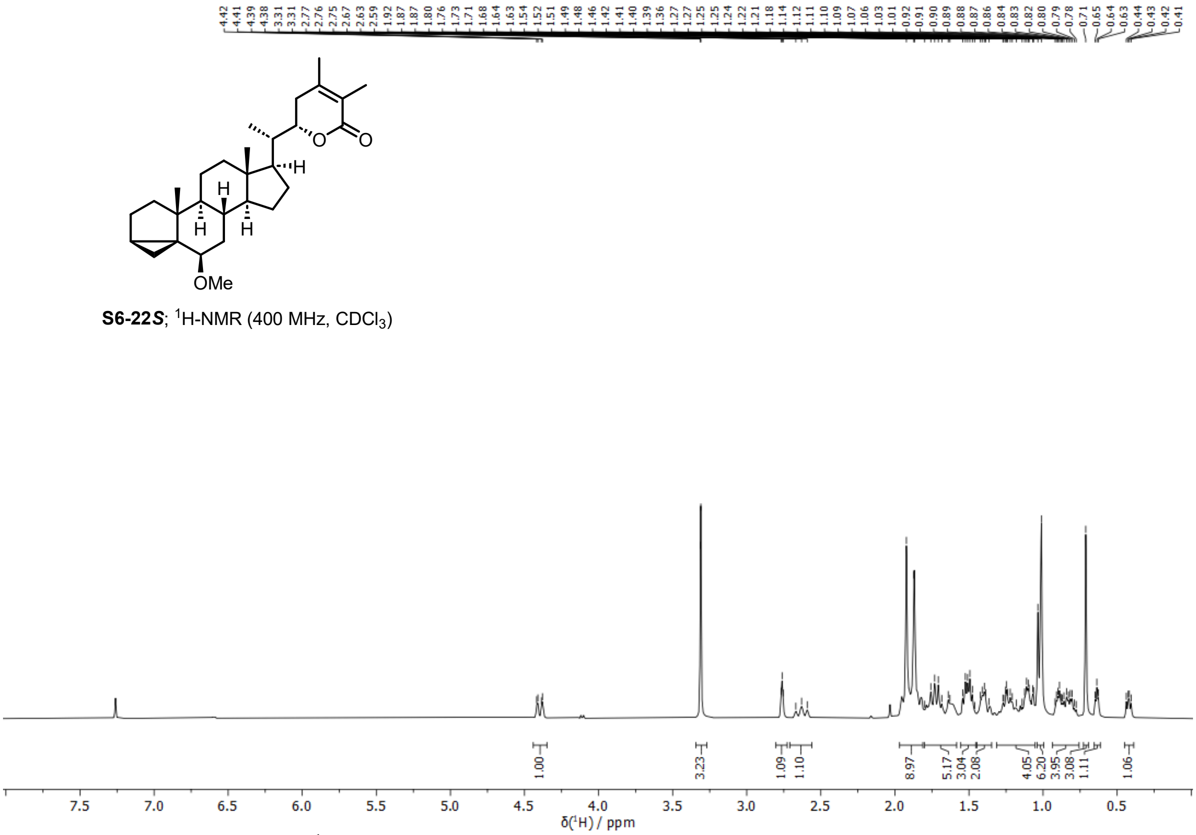


**Supplementary Fig. 74. $^{1}$H spectrum of (22*S*)-6*β*-methoxy-3*α*,5-cycloergosta-24,25-diene-26,22-lactone (S6-(22*S*)) (400 MHz, CDCl$_3$, 298 K).**

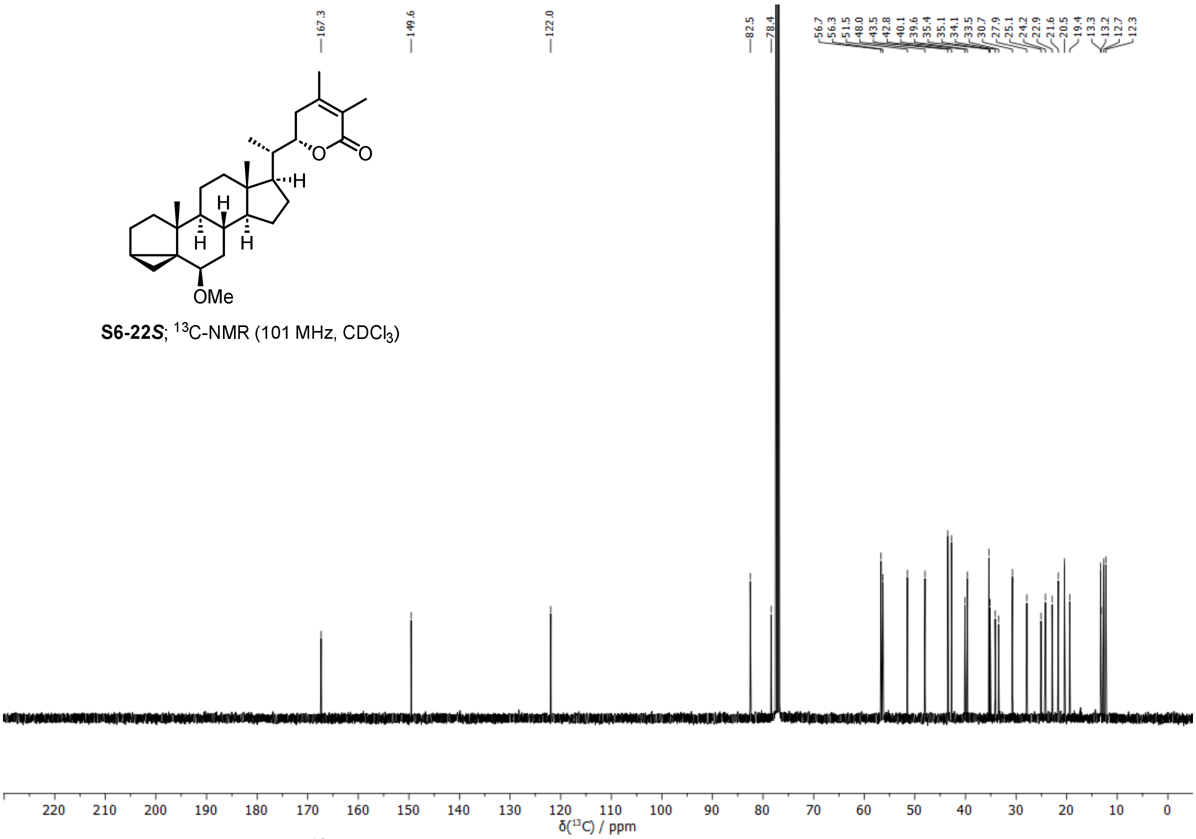


**Supplementary Fig. 75. $^{13}$C spectrum of (22*S*)-6*β*-methoxy-3*α*,5-cycloergosta-24,25-diene-26,22-lactone (S6-(22*S*)) (101 MHz, CDCl$_3$, 298 K).**

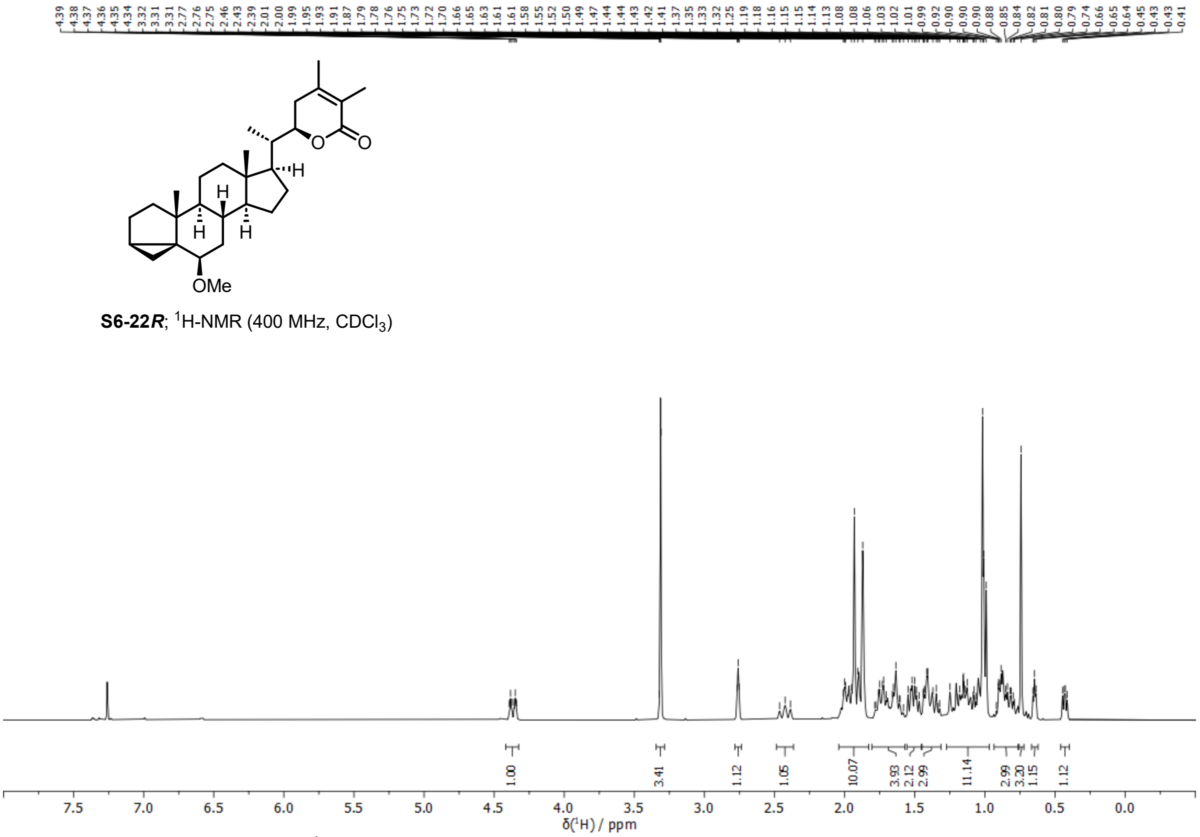


**Supplementary Fig. 76. $^{1}$H spectrum of (22*R*)-6*β*-methoxy-3*α*,5-cycloergosta-24,25-diene-26,22-lactone (S6-(22*R*)) (400 MHz, $CDCl_3$, 298 K).**

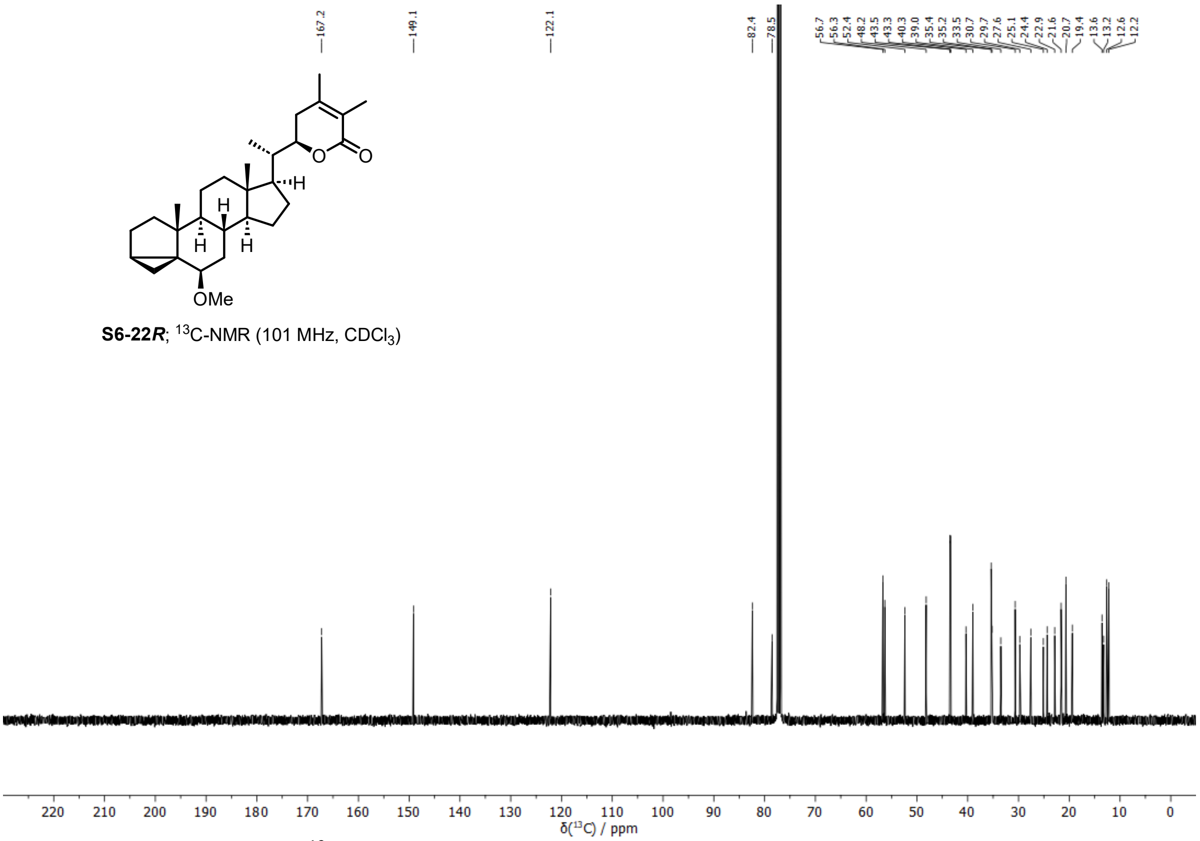


**Supplementary Fig. 77. $^{13}$C spectrum of (22*R*)-6*β*-methoxy-3*α*,5-cycloergosta-24,25-diene-26,22-lactone (S6-(22*R*)) (101 MHz, $CDCl_3$, 298 K).**

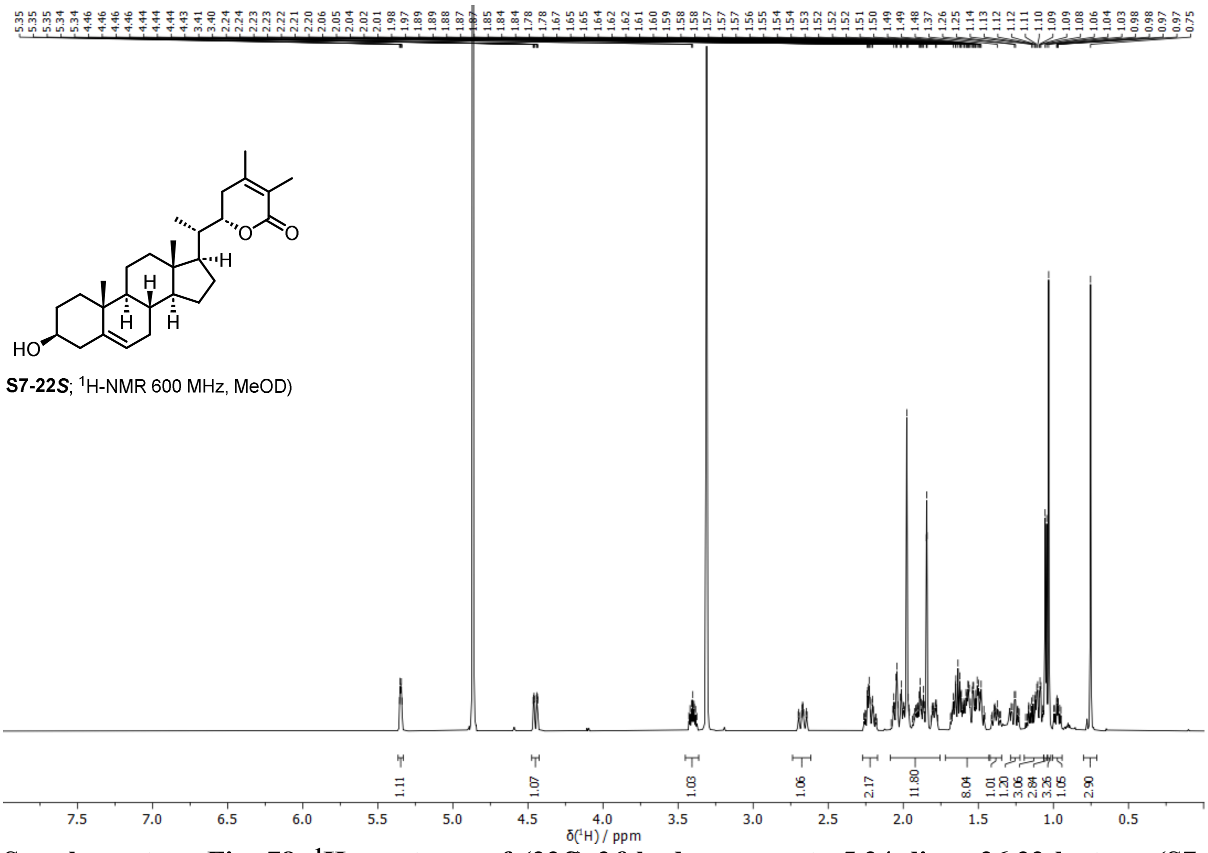


**Supplementary Fig. 78. $^{1}$H spectrum of (22*S*)-3*β*-hydroxyergosta-5,24-diene-26,22-lactone (S7-(22*S*)) (600 MHz, MeOD, 298 K).**

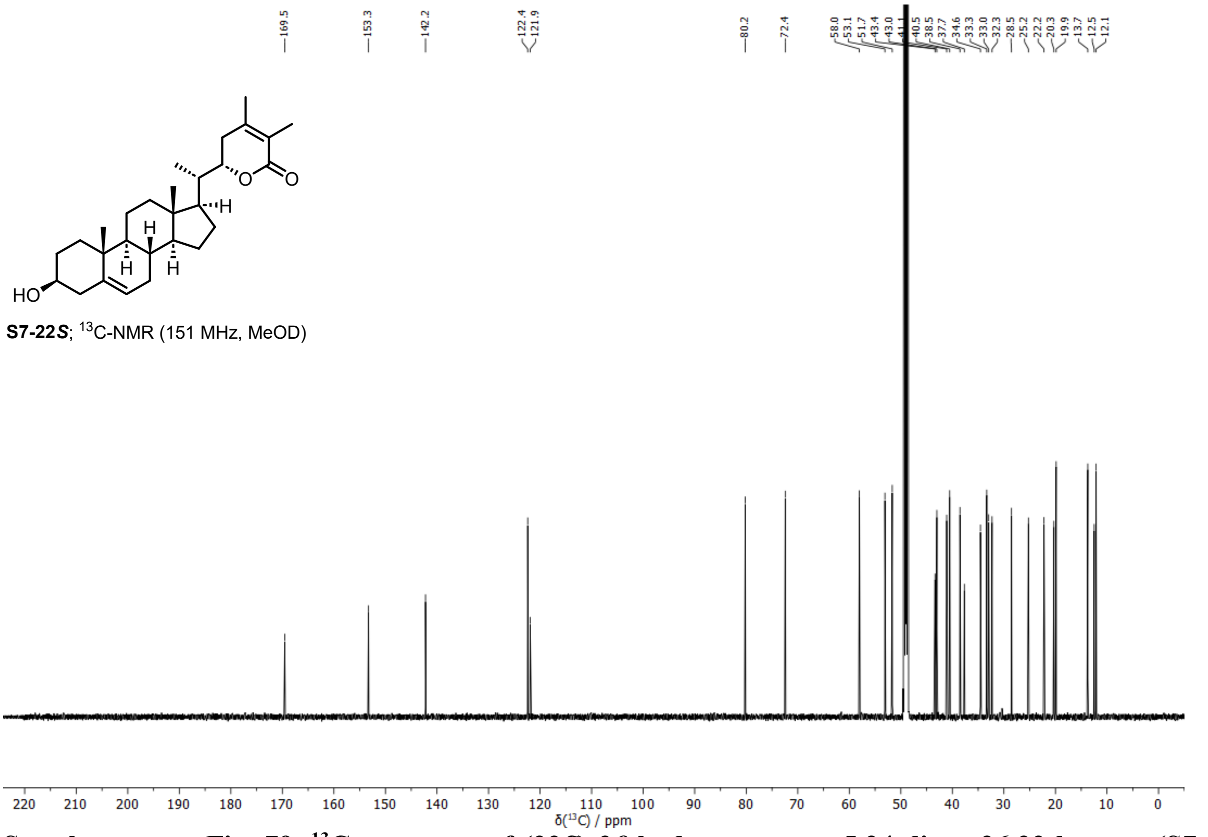


**Supplementary Fig. 79. $^{13}$C spectrum of (22*S*)-3*β*-hydroxyergosta-5,24-diene-26,22-lactone (S7-(22*S*)) (151 MHz, MeOD, 298 K).**

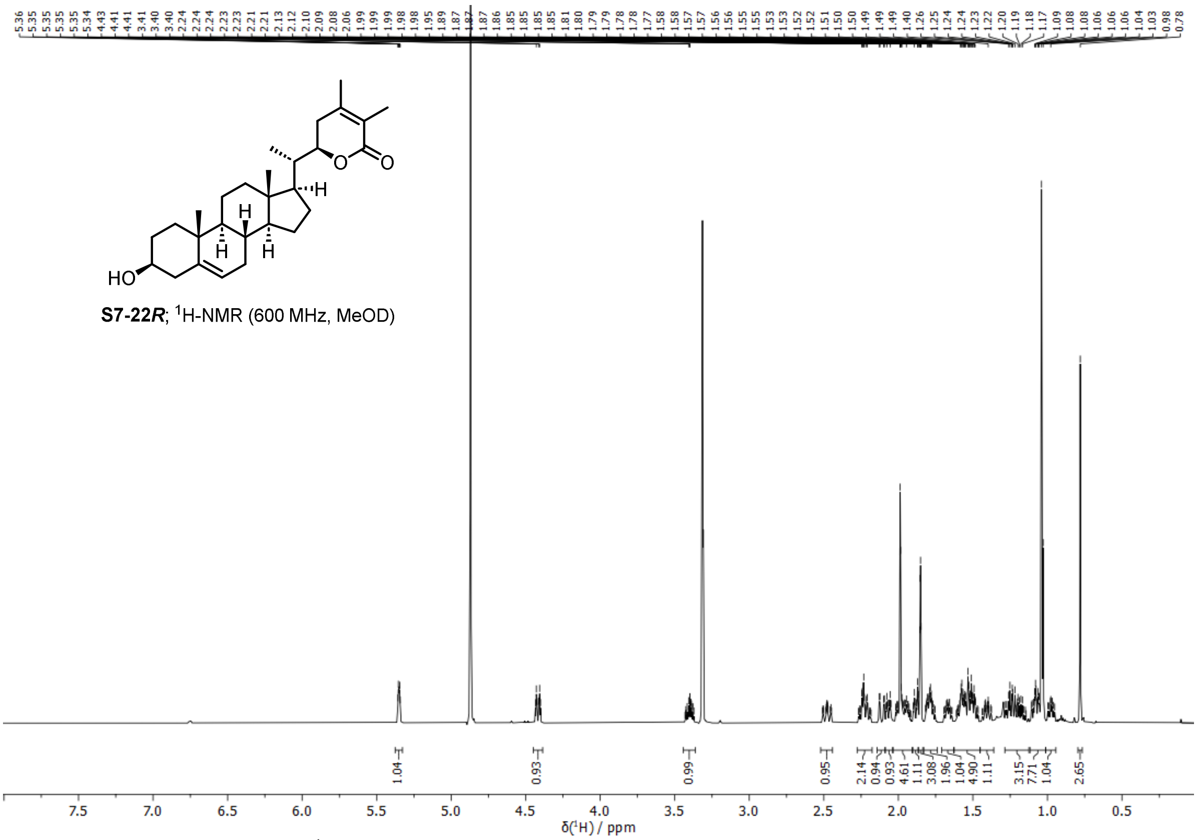


**Supplementary Fig. 80. $^{1}$H spectrum of (22*R*)-3*β*-hydroxyergosta-5,24-diene-26,22-lactone (S7-(22*R*)) (600 MHz, MeOD, 298 K).**

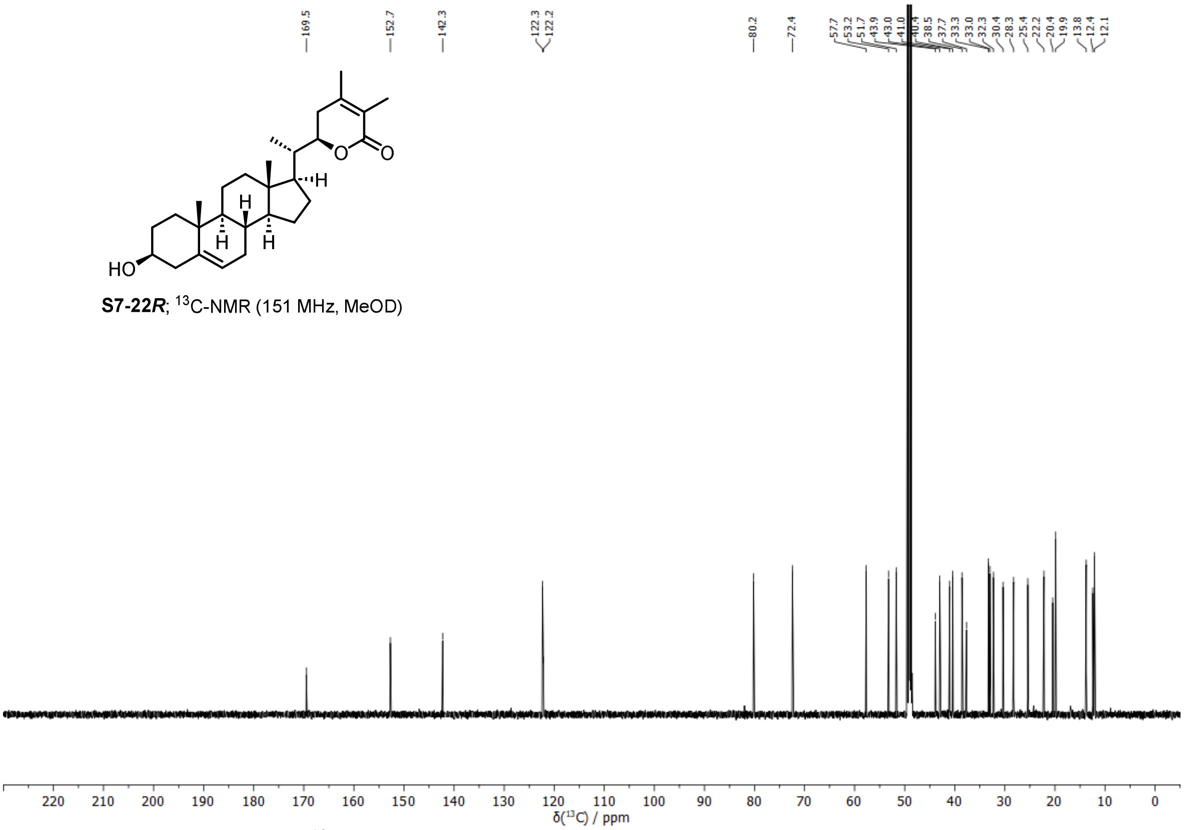


**Supplementary Fig. 81. $^{13}$C spectrum of (22*R*)-3*β*-hydroxyergosta-5,24-diene-26,22-lactone (S7-(22*R*)) (151 MHz, MeOD, 298 K).**

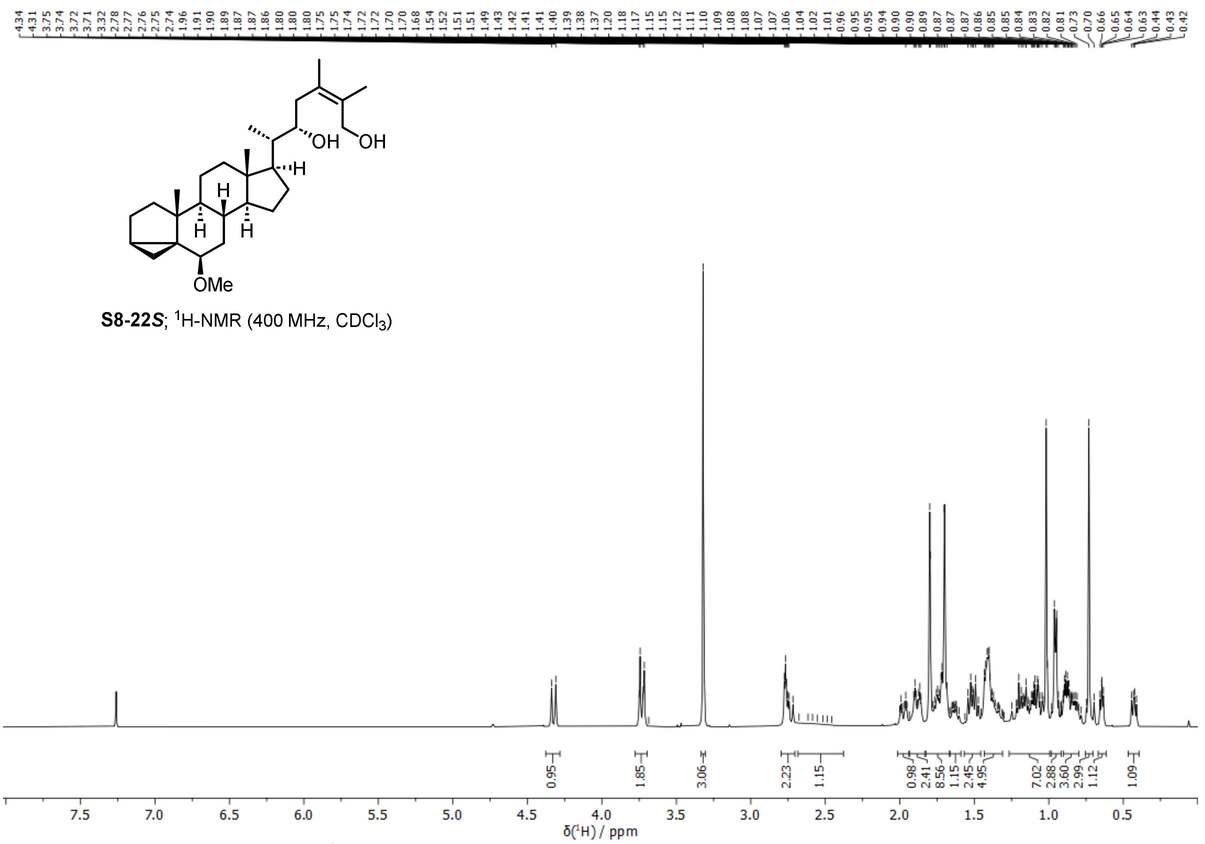

Supplementary Fig. 82. **$^{1}$H spectrum of (22*S*)-6*β*-methoxy-3*α*,5-cycloergost-24-ene-22,26-diol (S8-(22*S*)) (400 MHz, $CDCl_3$, 298 K).**

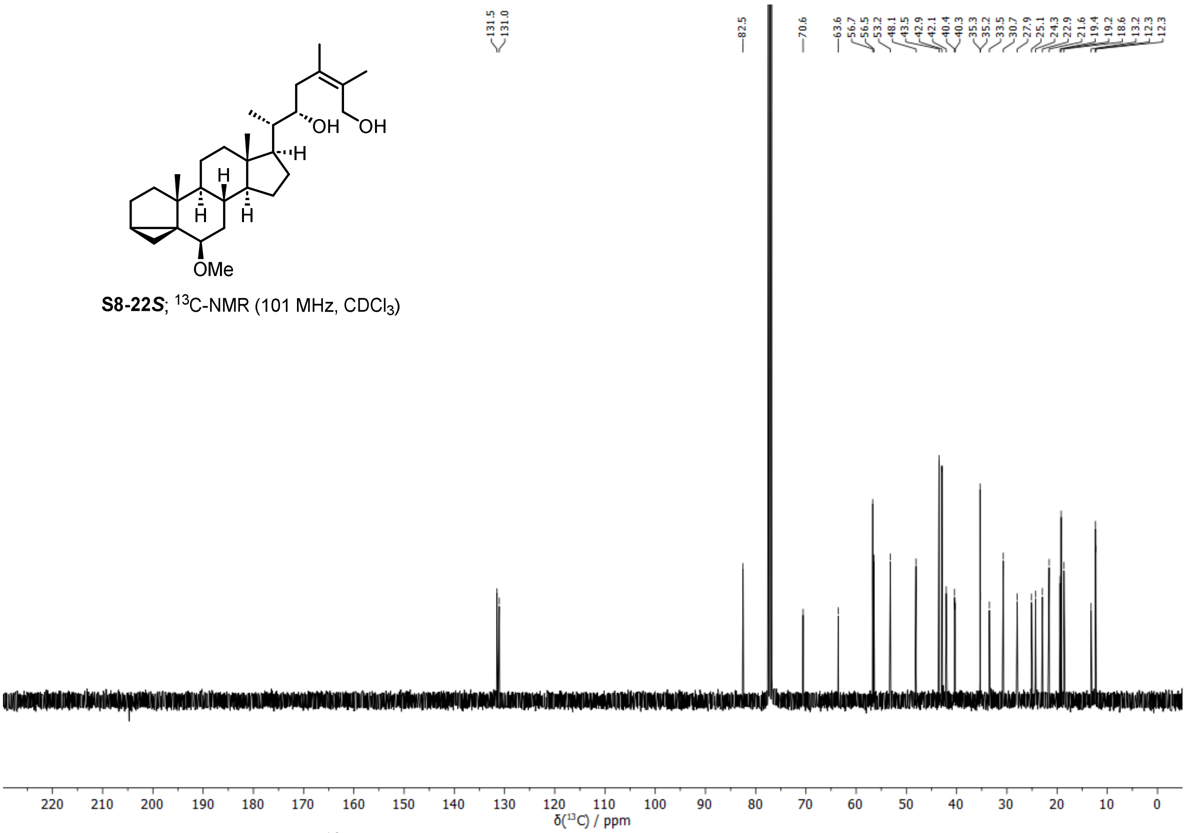

**Supplementary Fig. 83. $^{13}$C spectrum of (22*S*)-6*β*-methoxy-3*α*,5-cycloergost-24-ene-22,26-diol (S8-(22*S*)) (101 MHz, $CDCl_3$, 298 K).**

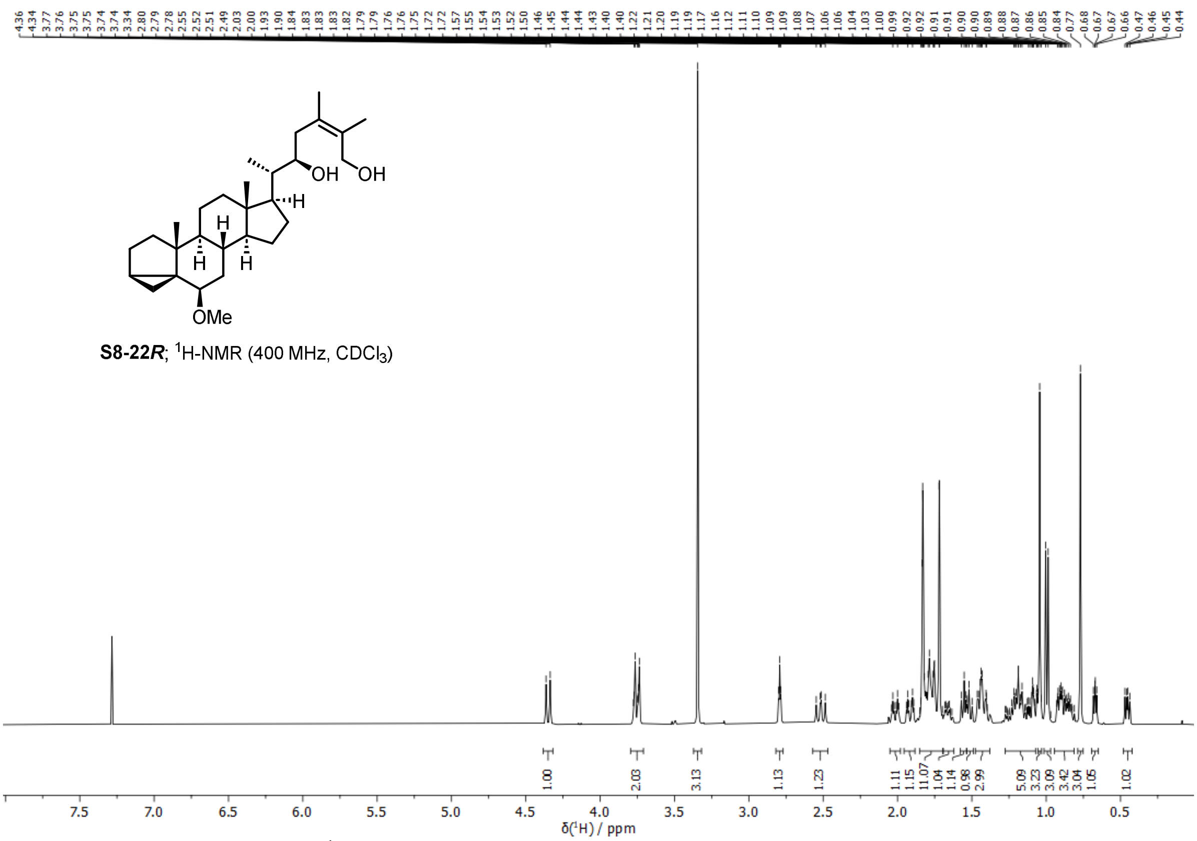


**Supplementary Fig. 84.** $^{1}$**H spectrum of (22*R*)-6*β*-methoxy-3*α*,5-cycloergost-24-ene-22,26-diol (S8-(22*R*)) (400 MHz, $CDCl_3$, 298 K).**

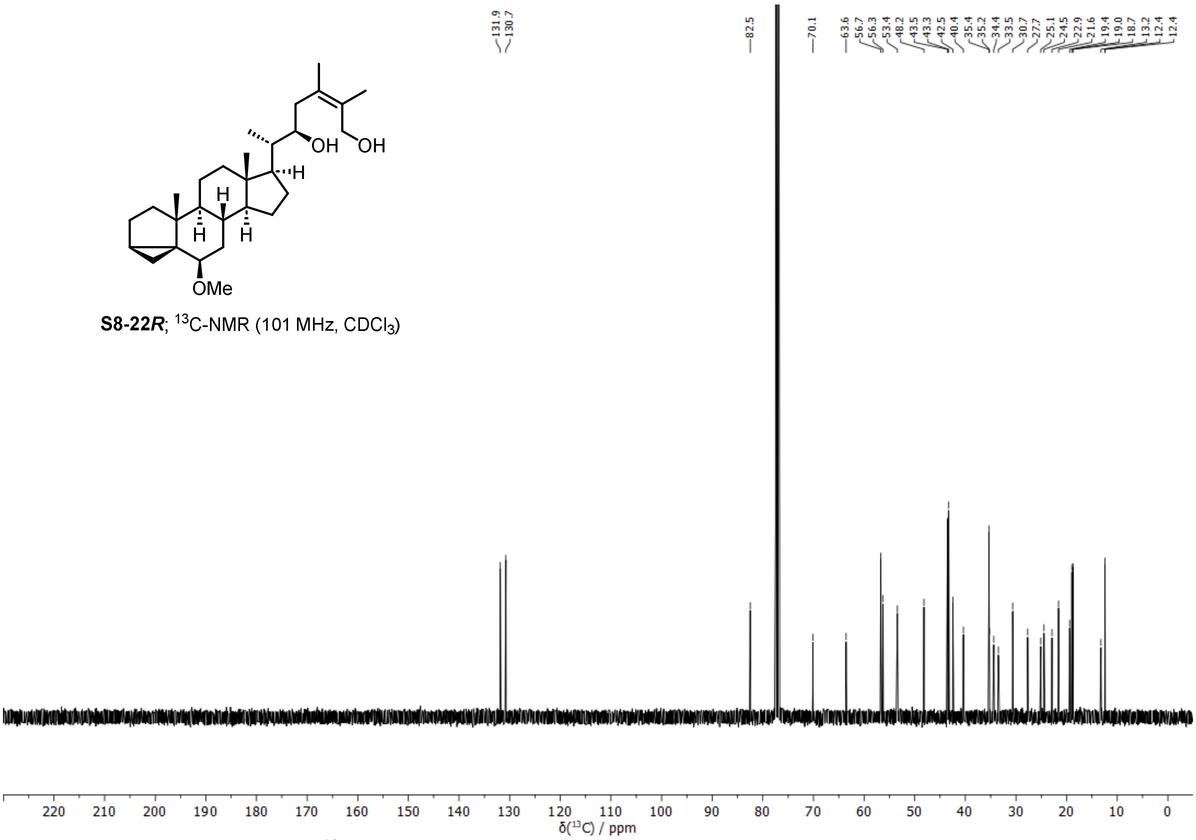


**Supplementary Fig. 85.** $^{13}$**C spectrum of (22*R*)-6*β*-methoxy-3*α*,5-cycloergost-24-ene-22,26-diol (S8-(22*R*)) (101 MHz, $CDCl_3$, 298 K).**

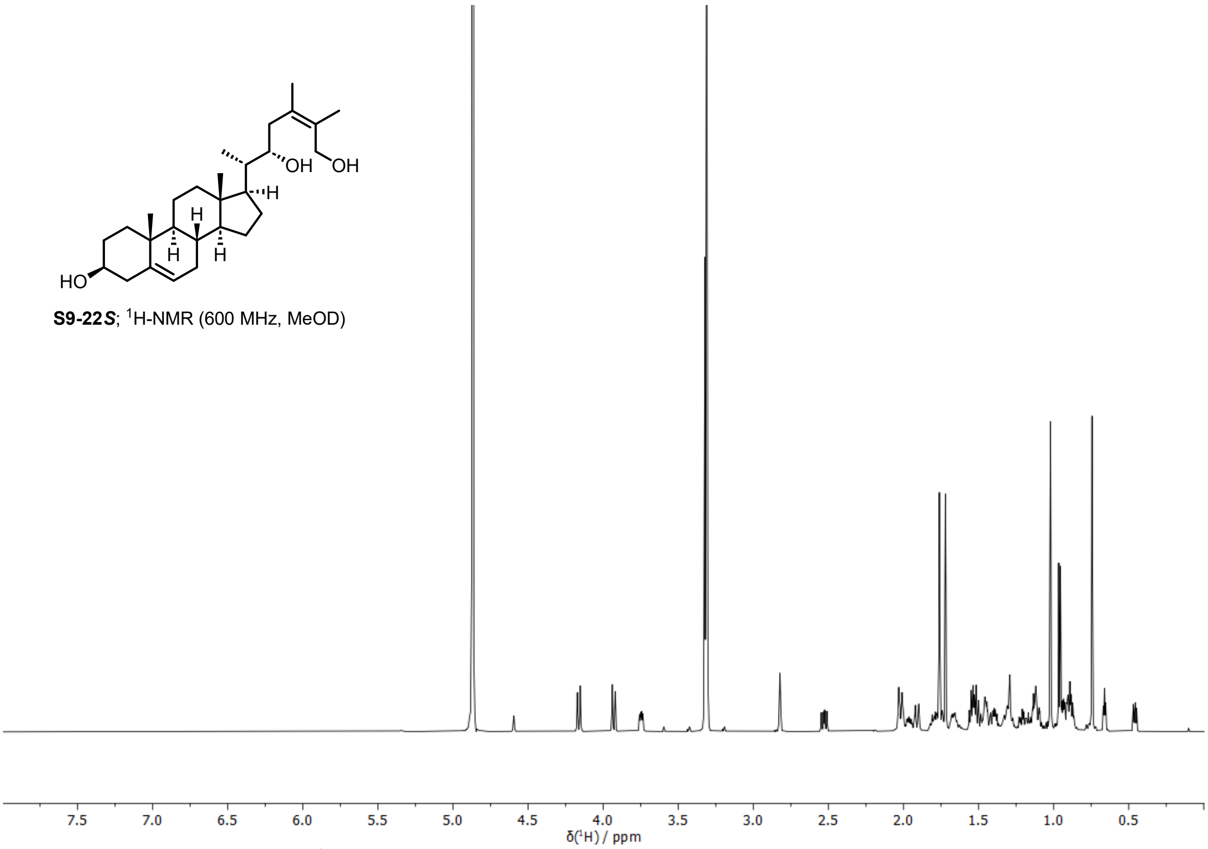


**Supplementary Fig. 86. $^{1}$H spectrum of (22*S*)-ergosta-5,24-diene-3*β*,22,26-triol (S9-(22*S*)) (600 MHz, MeOD, 298 K).**

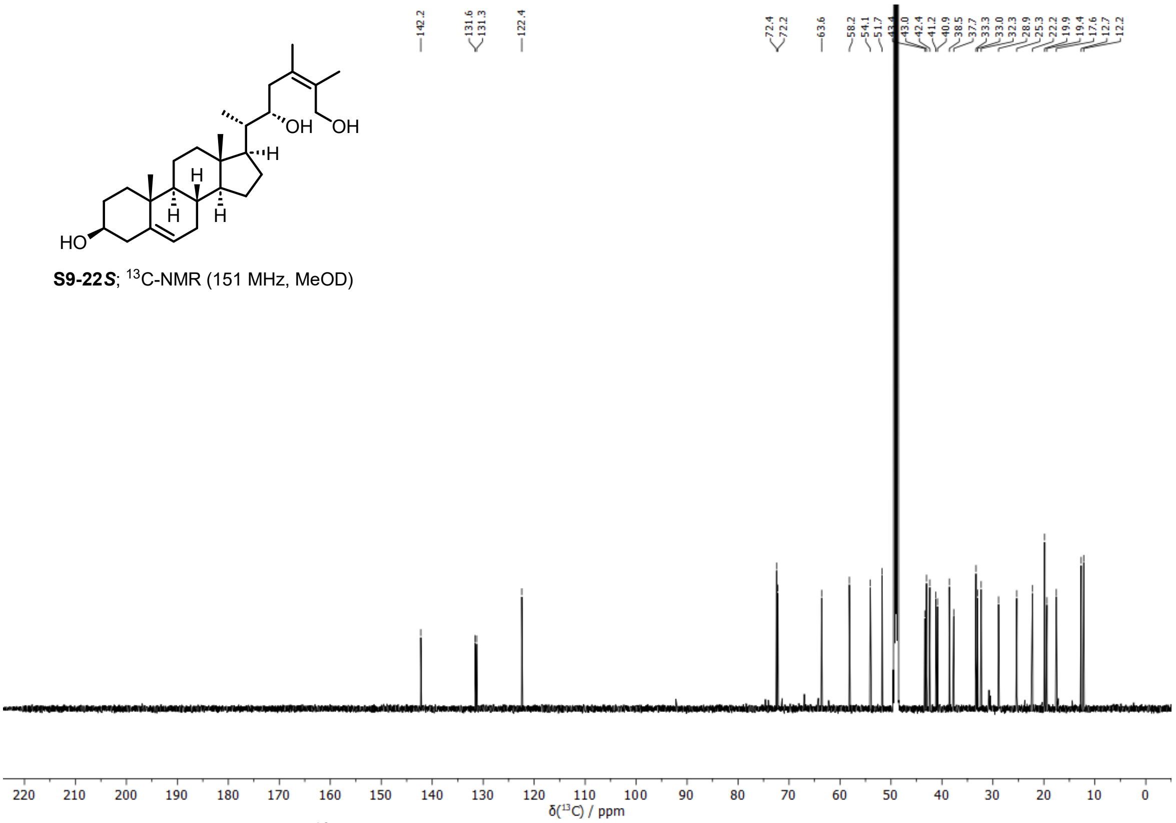


**Supplementary Fig. 87. $^{13}$C spectrum of (22*S*)-ergosta-5,24-diene-3*β*,22,26-triol (S9-(22*S*)) (151 MHz, MeOD, 298 K).**

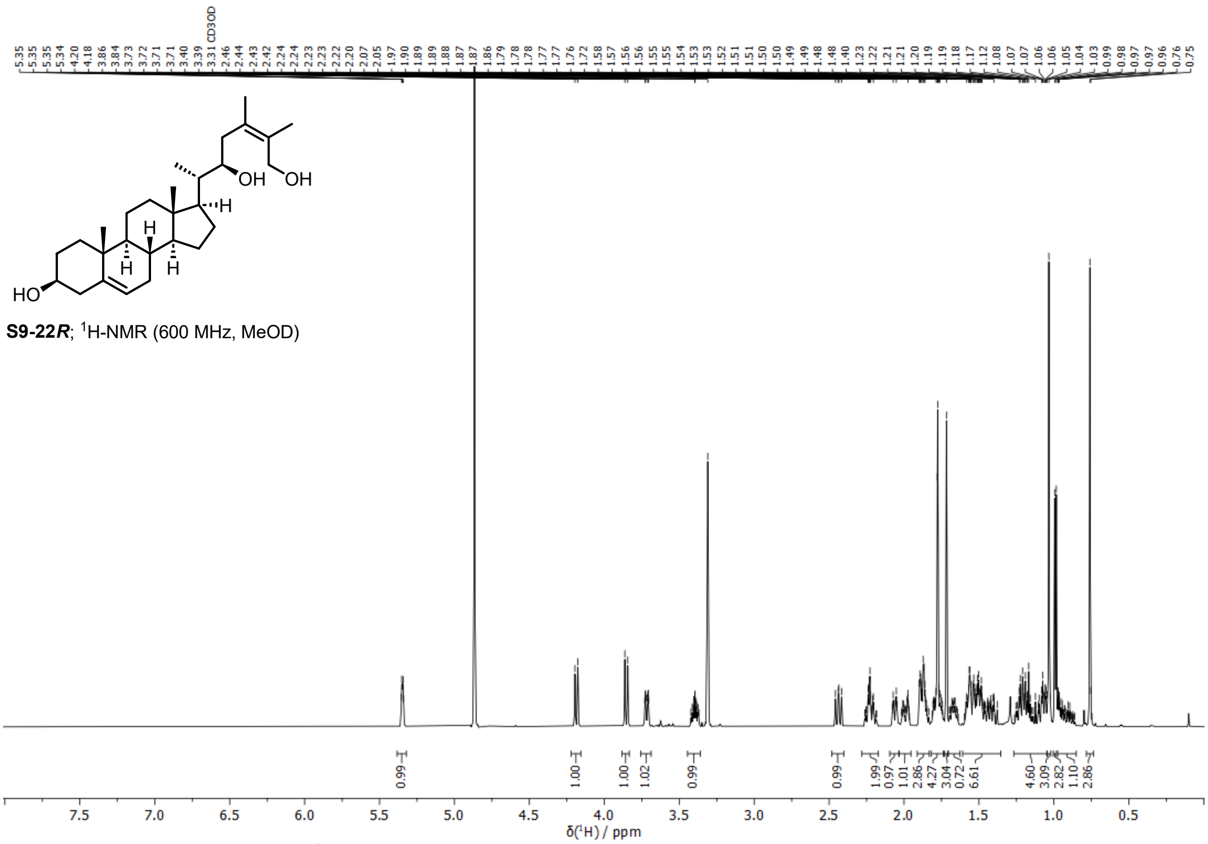


**Supplementary Fig. 88. ¹H spectrum of (22*R*)-ergosta-5,24-diene-3*β*,22,26-triol (S9-(22*R*)) (600 MHz, MeOD, 298 K).**

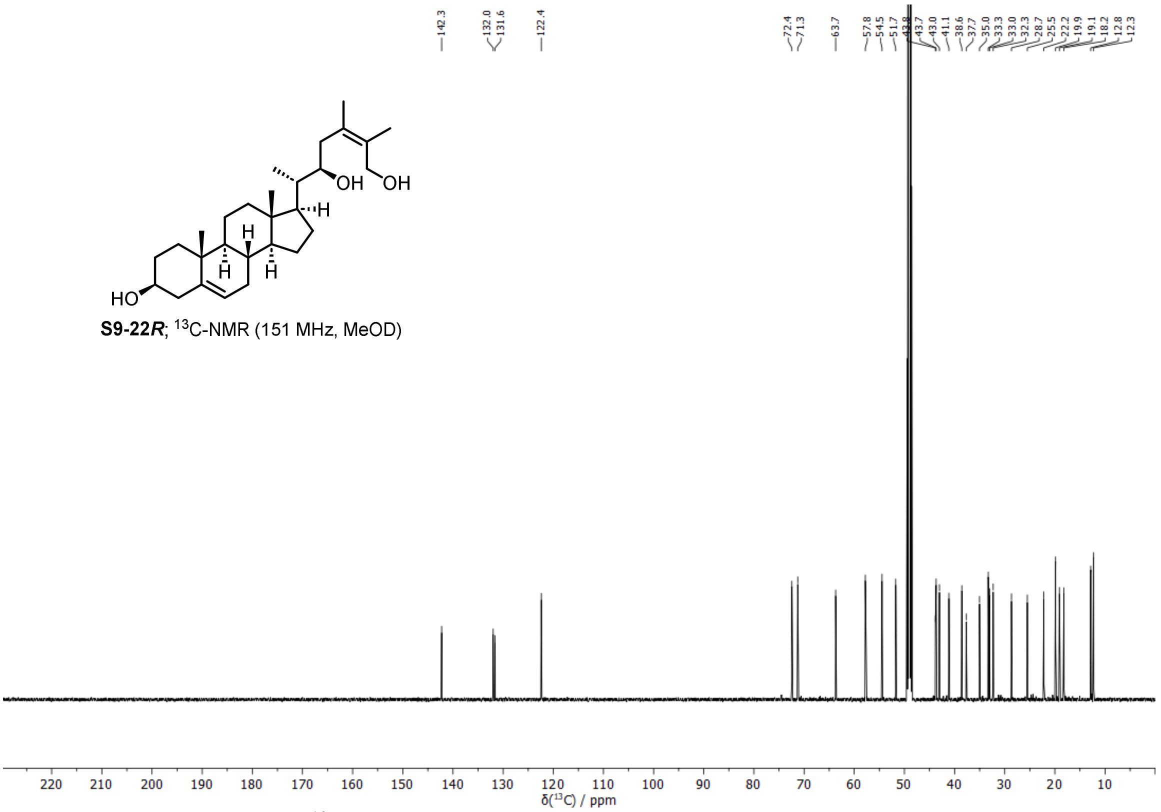


**Supplementary Fig. 89. ¹³C spectrum of (22*R*)-ergosta-5,24-diene-3*β*,22,26-triol (S9-(22*R*)) (151 MHz, MeOD, 298 K).**

**Supplementary Table 1. Assembly statistics of *W. somnifera* genome.**

| **Parameter** | **NextDenovo2 assembly** |
|---|---|
| Total number of contigs | 102 |
| Average contig length (bp) | 28,223,445 |
| Maximal contig length (bp) | 152,231,989 |
| N50 (bp) | 71,322,777 |
| N90 (bp) | 21,319,378 |
| GC Content (%) | 38.34 |
| Total length of contigs (bp) | 2,878,791,398 |
| BUSCO genes (Solanales_odb10) | 96.5% |

**Supplementary Table 2. The closest homologues of CYP genes putatively linked to withanolide biosynthesis based on gene silencing by others [22,23] are part of the withanolide gene clusters.**

| CYP gene (reference) | Match type | Bait sequence from literature | Best BLAST hit | | | | | |
|---|---|---|---|---|---|---|---|---|
| | | | Species | Position in genome | Gene ID | Gene name | Identity (%) | E-value |
| WsCYP749B1 (ref. [22]) | Protein sequence | MMIAVIAFSVFLIGVVVLGRYLYKSWWYPISLQHLMNSQGI<br>KGPRYEFGNGNSRATAEILMKFNNAPMDISHDIFPRLQPHF<br>RSWIKLYGSTFLYWMNSTKPQLVVSDVELIKEIFTNKQDSF<br>GKAKFDGILKRFVGDGSVFQKGHKWLKLRKVADNVFHAQSL<br>KDMLPAMVGRVESMLKTWKSYEGKEIEVFEEFRLLSLEMIS<br>NSVFGNDYSTGKHIFSMLDKIAYISAMSYGKSRNPIIDKLF<br>RSSEEIQADKILEELSDSFAGIIKKREDRVKAGEANNFGDD<br>FLGSLLEGRFNADENARISVDEIIEECKSFYFAGHKTVTSL<br>LSWSMLLLASNTDWQERAKNEVLEVLGQENPKAESISRLKT<br>VGMIINEALRLYPPFILLQRDVTKNTSLGKLKVPAGTEVII<br>AILAVHHNSEIWGEDAHLFKPERFAEGVSKATRDQVMAFLS<br>FGYGLRKCVGFNFATMEVKIALSMILQRYRLTVSPNYTHSP<br>IATFTLHPSNGIQIMLHPL | *Withania somnifera* | N/A | Wsom003G00477.1 | WsCYP749B-2b | 99.61 | 0.0 |
| | | | | N/A | Wsom003G00478.1 | WsCYP749B-2a | 98.83 | 0.0 |
| | | | | N/A | Wsom003G00475.1 | WsCYP749B-2c | 98.24 | 0.0 |
| | | | | N/A | Wsom090G00485.1 | WsCYP749B-a | 89.24 | 0.0 |
| | | | | N/A | Wsom090G00473.1 | WsCYP749B-b | 85.80 | 0.0 |
| PB.11591.2 (ref. [23]) | Protein sequence | MIPSDLVFDFTWKLVAVVLMVLLVRGFWRTYVSKFSFMYGN<br>REDVETDVEAGPVPRTPLLSLRYSLHHAPMLSANSDTKLAM<br>KGISHGACDYLVKLVRIEELRNIWQHIIRRKKVEPKMEYNL<br>VFLSTAFAVGILTLISVLKRANGWFYSMKFSSEKCRLPPGD<br>MGWPVVGNMLFFVKCLSAYDLKSFVSYFVTRFGQGGMYRTF<br>MFGKPSVIVTTPELCRKILMDDENFDLGFPSYILELLRKEP<br>IGGTSYQEDRLSRRLMTPIKSHALVSYFFDFLSETVQTTFE<br>KWATTGESLQLLFEMKKPTFKVLMQVLIGGDQVENKLLDTL<br>FKENNFRFAGLRSLPLDYPGSTYNRAMKGRGEIVKIYERII<br>NERKVMIAKTRGEPRTNLLDIMLDSQYDGEGKVLNDENIMK<br>VLLWYTFSGYESIAKVATQTIMLLEKHPECFKKAKEEQEEI<br>VKRRSSPDAGLTFGEIGQMKYVNNVINETLRLGSTETVLFR<br>DARTDVNINGYTIPKGWKVLALLGNLYMDPKTYVKPKEFNP<br>SRWDDFETKPNSFIPFGVGLRLCPGSNLVRLEVSVFLHYFL<br>LNYRLEQLNKDSKAEACIAKFKKISA | *Physalis grisea* | N/A | Pgri02031.1 | PgCYP88C-d | 99.17 | 0.0 |
| | | | *Physalis pruinosa* | N/A | Ppru02037.1 | PpCYP88C-d | 99.17 | 0.0 |
| PB.29095.11 (ref. [23]) | Primer (forward) | ATTGGAGCACTGGTTCTTGGTAGG | *Physalis grisea* | Chr1:114882829-114882806 | Pgri02022.1 | PgCYP749-a | 100.00 | 3.26e-05 |
| | | | | Chr1:115027450-115027473 | Pgri02030.1 | PgCYP749-b | 100.00 | 3.26e-05 |
| | | | *Physalis pruinosa* | Chr1:116023998-116023975 | Ppru02028.1 | PpCYP749-a | 100.00 | 3.28e-05 |
| | | | | Chr1:116168631-116168654 | Ppru02036.1 | PpCYP749-b | 100.00 | 3.28e-05 |

| | | | | | | | | |
|---|---|---|---|---|---|---|---|---|
| PB.29095.11 (ref. [23]) | Primer (reverse) | GTTCCCACTGCCAAATTCGTAAGG | *Physalis grisea* | Chr1:114882710-114882733 | Pgri02022.1 | PgCYP749-a | 100.00 | 3.26e-05 |
| | | | | Chr1:115027569-115027546 | Pgri02030.1 | PgCYP749-b | 100.00 | 3.26e-05 |
| | | | *Physalis pruinosa* | Chr1:116023879-11602390 | Ppru02028.1 | PpCYP749-a | 100.00 | 3.28e-05 |
| | | | | Chr1:116168750-116168727 | Ppru02036.1 | PpCYP749-b | 100.00 | 3.28e-05 |

**Supplementary Table 3. Overview of electron impact fragmentations and proposed fragments of compounds investigated in this work (TMS ethers).**

Relative intensities are provided in parentheses. Carbon atoms of proposed fragments and fragment names are highlighted in colour.

**M**=molecular ion; **F**=**M**−$CH_3^{\bullet}$; *= −TMSOH; **= −$(TMSOH)_2$; ***= −$(TMSOH)_3$; nd: not detected.

| Compound (TMS ether) | $[M]^{+\bullet}$ and $[M-CH_3]^+$ | $[M-(C4\text{-}C28)]^+$ | [M−side chain−$H_2]^+$ | [M−side chain$]^+$ | $[M-(C23\text{-}C28)]^{+\bullet}$ (McLafferty, [(a)]) or $[M-(C23\text{-}C28)]^+$ (α-cleavage, [(b)]) | $[M-(C1\text{-}C21)]^+$ | $[M-(C1\text{-}C23)]^+$ | $[M-(C1\text{-}C24)-C28]^+$ |
|---|---|---|---|---|---|---|---|---|
| | | 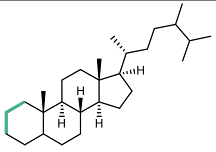 | 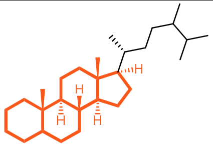 | | 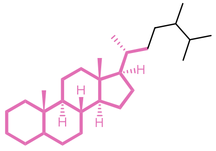 | 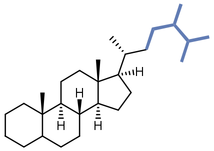 | 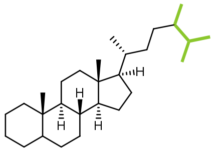 | 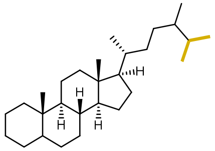 |
| **24-Methylene-cholesterol (1)** | 470 (13) **M**<br>455 (10) **F**<br>380 (24) **M***<br>365 (18) **F*** | 129 (100) **a**<br>341 (26) [M−**a**]$^+$<br>131 (24) **b**<br>217 (6) **c** | 343 (20) **d**<br>253 (18) **d*** | 345 (nd) **e**<br>255 (11) **e*** | 386 (35) **j**[(a)]<br>296 (25) **j***[(a)] | - | 69 (52) **r** | - |
| **24-Methyldesmosterol (2)** | 470 (17) **M**<br>455 (20) **F**<br>380 (11) **M***<br>365 (21) **F*** | 129 (91) **a**<br>341 (20) [M−**a**]$^+$<br>131 (26) **b**<br>217 (5) **c** | 343 (50) **d**<br>253 (20) **d*** | 345 (nd) **e**<br>255 (23) **e*** | 386 (60) **j**[(a)]<br>296 (34) **j***[(a)] | - | 69 (46) **s** | - |
| **Ergosta-5,7,24-trien-3β-ol (6)** | 468 (33) **M**<br>453 (3) **F**<br>378 (20) **M***<br>363 (100) **F*** | 129 (23) **a**<br>131 (23) **b**<br>337 (62) [M−**b**]$^+$<br>217 (1) **c** | 341 (1) **f**<br>251 (14) **f*** | 343 (1) **g**<br>253 (14) **g*** | 384 (<1) **k**[(a)]<br>294 (14) **k***[(a)] | - | 69 (16) **s** | - |
| **Ergosta-5,25-diene-3β,24ξ-diol (physalindicanol A) (7)** | 558 (1) **M**<br>543 (1) **F** | 129 (6) **a**<br>131 (2) **b**<br>217 (1) **c** | 343 (1) **d**<br>253 (<1) **d*** | 345 (nd) **e**<br>255 (<1) **e*** | 386 (<1) **j**[(a)]<br>296 (<1) **j***[(a)] | - | 157 (100) **t** | - |
| **Ergosta-5,24(28)-diene-3β,25-diol (physalindicanol B) (8)** | 558 (2) **M**<br>543 (6) **F**<br>468 (7) **M*** | 129 (42) **a** | 343 (7) **d**<br>253 (4) **d*** | 345 (nd) **e**<br>255 (5) **e*** | 386 (16) **j**[(a)]<br>296 (8) **j***[(a)] | - | 157 (100) **u** | 131 (74) **v** |
| **(22*R*)-Ergosta-5,24-diene-3β,22-diol (10)** | 558 (nd) **M**<br>543 (3) **F** | 129 (39) **a**<br>131 (18) **b**<br>217 (4) **c** | 343 (14) **d**<br>253 (29) **d*** | 345 (nd) **e**<br>255 (41) **e*** | 475 (25) **l**[(b)]<br>385 (21) **l***[(b]<br>295 (75) **l****[(b))] | 185 (40) **o**<br>95 (34) **o*** | - | - |
| **(22*R*)-Ergosta-5,7,24-triene-3β,22-diol (11)** | 556 (8) **M**<br>541 (3) **F**<br>451 (11) **F*** | 129 (22) **a**<br>131 (11) **b**<br>217 (2) **c** | 341 (22) **f**<br>251 (25) **f*** | 343 (nd) **g**<br>253 (25) **g*** | 473 (23) **m**[(b)]<br>383 (25) **m***[(b)]<br>293 (70) **m****[(b)] | 185 (33) **o**<br>95 (24) **o*** | - | - |

| | | | | | | | | |
|---|---|---|---|---|---|---|---|---|
| **(22*R*)-Ergosta-5,24(28)-diene-3β,22,25-triol (phyministerol A) (12)** | 646 (nd) **M**<br>631 (1) **F**<br>541 (<1) **F*** | 129 (10) **a**<br>217 (1) **c** | 343 (3) **d**<br>253 (5) **d*** | 345 (nd) **e**<br>255 (7) **e*** | 475 (6) **l**[(b)]<br>385 (3) **l***[(b)]<br>295 (9) **l****[(b)] | 273 (nd) **p**<br>183 (100) **p***<br>93 (15) **p**** | 157 (10) **u** | 131 (10) **v** |
| **(22*R*)-24,25-Epoxyergost-5-ene-3β,22-diol (13)** | 574 (nd) **M**<br>559 (1) **F** | 129 (17) **a**<br>217 (1) **c** | 343 (3) **d**<br>253 (6) **d*** | 345 (nd) **e**<br>255 (5) **e*** | 475 (3) **l**[(b)]<br>385 (2) **l***[(b)]<br>295 (9) **l****[(b)] | 201 (100) **q**<br>111 (25) **q*** | - | - |
| **(22*R*)-Ergosta-5,24-diene-1α,3β,22-triol (14)** | 646 (nd) **M**<br>631 (2) **F** | 129 (25) **a**<br>217 (17) **c** | 431 (nd) **h**<br>341 (7) **h***<br>251 (6) **h**** | 433 (nd) **i**<br>343 (7) **i***<br>253 (8) **i**** | 563 (14) **n**[(b)]<br>473 (8) **n***[(b)]<br>383 (33) **n****[(b)]<br>293 (32) **n*****[(b)] | 185 (25) **o**<br>95 (27) **o*** | - | - |
| **(22*R*)-Ergosta-5,24(28)-diene-1α,3β,22,25-tetraol (15)** | 734 (nd) **M**<br>644 (6) **M***<br>554 (6) **M**** | 129 (18) **a**<br>217 (14) **c** | 431 (nd) **h**<br>341 (4) **h***<br>251 (44) **h**** | 433 (nd) **i**<br>343 (4) **i***<br>253 (6) **i**** | 563 (7) **n**[(b)]<br>473 (3) **n***[(b)]<br>383 (6) **n****[(b)]<br>293 (8) **n*****[(b)] | 273 (nd) **p**<br>183 (67) **p***<br>93 (21) **p**** | 157 (22) **u** | 131 (16) **v** |

**Supplementary Table 4.** **$^{1}$H NMR data of isolated compounds compared with [26,27-$^{2}H_6$]24-methyldesmosterol[24] (2) (600 MHz, $CDCl_3$, 298 K).**

| | | | | | | | |
|---|---|---|---|---|---|---|---|
| | 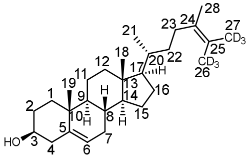 | 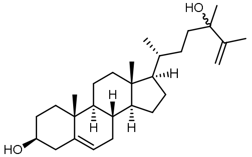 | 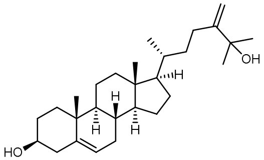 | 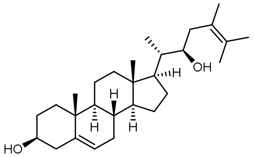 | 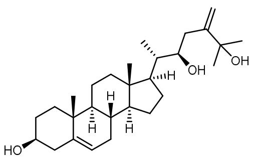 | 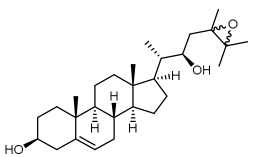 | 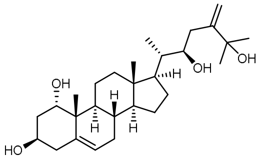 |
| **Pos.** | **[26,27-$^{2}H_6$]24-Methyldesmosterol reported data[24]** | **Ergosta-5,25-diene-3β,24ξ-diol (physalindicanol A) (7)** | **Ergosta-5,24(28)-diene-3β,25-diol (physalindicanol B) (8)** | **(22*R*)-Ergosta-5,24-diene-3β,22-diol (10)** | **(22*R*)-Ergosta-5,24(28)-diene-3β,22,25-triol (phyministerol A) (12)** | **(22*R*)-24,25-Epoxy-ergost-5-ene-3β,22-diol (13)** | **(22*R*)-Ergosta-5,24(28)-diene-1α,3β,22,25-tetraol (15)** |
| **1** | n.r. [a] | 1.08 m<br>1.84 m | 1.08 m<br>1.84 m | 1.08 m<br>1.85 m | 1.08 m<br>1.85 m | 1.08 m<br>1.85 m | 3.86 br s |
| **2** | n.r. [a] | 1.51 m<br>1.84 m | 1.50 m<br>1.83 m | 1.52 m<br>1.84 m | 1.52 m<br>1.84 m | 1.52 m<br>1.84 m | 1.75 m<br>2.10 m |
| **3** | 3.54 m | 3.52 m | 3.53 m | 3.53 m | 3.53 m | 3.52 m | 3.99 m |
| **4** | n.r. [a] | 2.24 m<br>2.30 ddd (13.1, 5.1, 2.1) | 2.25 m<br>2.30 ddd (13.1, 5.1, 2.2) | 2.25 m<br>2.30 ddd (12.9, 5.2, 2.1) | 2.25 m<br>2.30 ddd (13.0, 5.3, 2.1) | 2.25 m<br>2.30 ddd (13.1, 5.0, 2.0) | 2.32 m<br>2.39 ddd (13.5, 5.4, 2.3) |
| **5** | - | - | - | - | - | - | - |
| **6** | 5.35 br s | 5.35 d (5.2) | 5.35 d (5.2) | 5.35 d (5.1) | 5.35 d (4.9) | 5.35 d (5.4) | 5.60 d (5.1) |
| **7** | n.r. [a] | 1.53 m<br>1.97 m | 1.53 m<br>1.97 m | 1.54 m<br>1.99 m | 1.53 m<br>1.99 m | 1.53 m<br>1.99 m | 1.59 m<br>2.00 m |
| **8** | n.r. [a] | 1.45 m | 1.45 m | 1.47 m | 1.47 m | 1.47 m | 1.50 m |
| **9** | n.r. [a] | 0.93 m | 0.93 m | 0.94 m | 0.95 m | 0.94 m | 1.60 m |
| **10** | - | - | - | - | - | - | - |
| **11** | n.r. [a] | 1.46 m<br>1.50 m | 1.46 m<br>1.50 m | 1.47 m<br>1.52 m | 1.47 m<br>1.53 m | 1.46 m<br>1.52 m | 1.48 m<br>1.52 m |
| **12** | n.r. [a] | 1.14 m<br>2.00 m | 1.16 m<br>2.00 ddd (12.7, 3.4, 3.4) | 1.19 m<br>2.03 ddd (12.5, 3.3, 3.2) | 1.19 m<br>2.03 ddd (12.5, 3.5, 3.4) | 1.16 m<br>2.03 m | 1.25 m<br>2.03 ddd (13.2, 3.8, 3.7) |
| **13** | - | - | - | - | - | - | - |
| **14** | n.r. [a] | 0.98 m | 1.00 m | 1.00 m | 1.02 m | 0.99 m | 1.08 m |
| **15** | n.r. [a] | 1.08 m<br>1.59 m | 1.08 m<br>1.59 m | 1.14 m<br>1.63 m | 1.14 m<br>1.63 m | 1.14 m<br>1.63 m | 1.14 m<br>1.63 m |
| **16** | n.r. [a] | 1.25 m<br>1.83 m | 1.27 m<br>1.86 m | 1.42 m<br>1.78 m | 1.40 m<br>1.80 m | 1.47 m<br>1.75 m | 1.41 m<br>1.82 m |
| **17** | n.r. [a] | 1.11 m | 1.14 m | 1.15 m | 1.17 m | 1.06 m | 1.19 m |
| **18** | 0.68 s | 0.67 s | 0.69 s | 0.72 s | 0.72 s | 0.72 s | 0.72 s |
| **19** | 1.01 s | 1.00 s | 1.01 s | 1.01 s | 1.01 s | 1.01 s | 1.04 s |
| **20** | n.r. [a] | 1.38 m | 1.45 m | 1.86 m | 1.76 m | 1.81 m | 1.76 m |

| | | | | | | | |
|---|---|---|---|---|---|---|---|
| **21** | 0.96 d (6.6) | 0.92 d (6.6) | 0.97 d (6.5) | 0.99 d (6.7) | 0.98 d (6.7). | 0.92 d (6.8) | 0.98 d (6.8) |
| **22** | n.r. [a] | 1.25 m | 1.22 m<br>1.59 m | 3.76 ddd (11.2, 2.9, 2.9) | 3.80 td (7.0, 3.8) | 4.17 ddd (10.2, 2.6, 1.2) | 3.80 td (6.4, 3.2) |
| **23** | n.r. [a] | 1.43 m<br>1.64 m | 1.93 m<br>2.15 ddd (16.1, 11.7, 4.8) | 1.80 m<br>2.40 dd (13.5, 11.2) | 2.20 d (7.0) | 1.56 m | 2.20 d (6.4) |
| **24** | - | - | - | - | - | - | - |
| **25** | - | - | - | - | - | - | - |
| **26** | - | 4.82 br s<br>4.96 br s | 1.35 s | 1.71 s | 1.43 s | 1.35 s | 1.44 s |
| **27** | - | 1.74 s | 1.35 s | 1.72 s | 1.35 s | 1.37 s | 1.35 s |
| **28** | 1.61 s | 1.30 s | 4.76 br s<br>5.09 br s | 1.66 s | 4.88 s<br>5.12 s | 1.37 s | 4.88 s<br>5.12 s |

[a] n.r. = not reported

**Supplementary Table 5. $^{13}$C NMR data of isolated compounds compared with [26,27-$^{2}H_{6}$]24-methyldesmosterol[24] (151 MHz, $CDCl_3$, 298 K).**

| Pos. | [26,27-$^{2}H_{6}$]24-Methyldesmosterol reported data[24] | Ergosta-5,25-diene-3β,24ξ-diol (physalindicanol A) (7) | Ergosta-5,24(28)-diene-3β,25-diol (physalindicanol B) (8) | (22*R*)-Ergosta-5,24-diene-3β,22-diol (10) | (22*R*)-Ergosta-5,24(28)-diene-3β,22,25-triol (phyministerol A) (12) | (22*R*)-24,25-Epoxy-ergost-5-ene-3β,22-diol (13) | (22*R*)-Ergosta-5,24(28)-diene-1α,3β,22,25-tetraol (15) |
|---|---|---|---|---|---|---|---|
| | 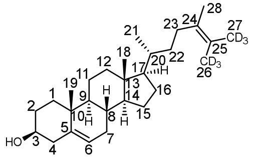 | 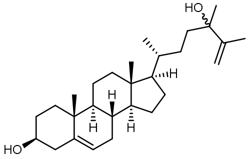 | 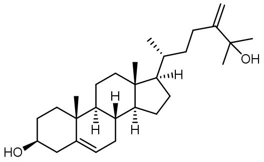 | 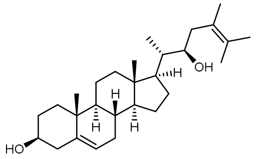 | 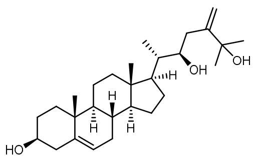 | 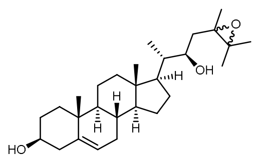 | 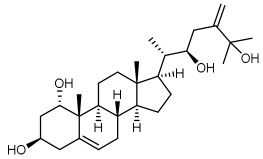 |
| **1** | 37.25 | 37.4 | 37.4 | 37.4 | 37.4 | 37.4 | 73.1 |
| **2** | 31.68 | 31.8 | 31.8 | 31.8 | 31.8 | 31.8 | 38.4 |
| **3** | 71.81 | 72.0 | 72.0 | 72.0 | 71.9 | 72.0 | 66.6 |
| **4** | 42.31 | 42.4 | 42.5 | 42.4 | 42.4 | 42.5 | 41.5 |
| **5** | 140.75 | 140.9 | 140.9 | 141.0 | 141.0 | 141.0 | 137.5 |
| **6** | 121.71 | 121.9 | 121.9 | 121.8 | 121.8 | 121.7 | 125.8 |
| **7** | 31.91 | 32.0 | 32.1 | 32.1 | 32.1 | 32.1 | 31.9 |
| **8** | 31.91 | 32.0 | 32.1 | 32.1 | 32.1 | 32.1 | 32.0 |
| **9** | 50.13 | 50.2 | 50.3 | 50.3 | 50.3 | 50.3 | 41.8 |
| **10** | 36.51 | 36.6 | 36.7 | 36.7 | 36.7 | 36.7 | 41.9 |
| **11** | 21.08 | 21.2 | 21.2 | 21.2 | 21.2 | 21.2 | 20.4 |
| **12** | 39.75 | 39.9 | 39.9 | 39.9 | 39.9 | 40.0 | 39.6 |
| **13** | 42.34 | 42.4 | 42.5 | 42.9 | 42.9 | 42.9 | 42.8 |
| **14** | 56.74 | 56.9 | 56.9 | 56.5 | 56.5 | 56.6 | 56.4 |
| **15** | 24.32 | 24.4 | 24.4 | 24.6 | 24.6 | 24.6 | 24.6 |
| **16** | 28.18 | 28.4 | 28.4 | 27.5 | 27.6 | 27.7 | 27.6 |
| **17** | 55.83 | 55.9 | 56.0 | 53.2 | 53.4 | 53.2 | 53.4 |
| **18** | 11.85 | 12.0 | 12.0 | 12.0 | 12.1 | 12.0 | 12.0 |
| **19** | 19.39 | 19.6 | 19.6 | 19.5 | 19.5 | 19.5 | 19.6 |
| **20** | 34.33 | 35.9 | 36.1 | 40.7 | 42.1 | 41.0 | 42.0 |
| **21** | 18.79 | 18.9 | 19.0 | 12.6 | 12.6 | 12.7 | 12.6 |
| **22** | 35.96 | 29.8 | 35.6 | 70.8 | 75.0 | 71.3 | 74.9 |
| **23** | 31.05 | 36.6 | 27.7 | 34.6 | 32.2 | 31.6 | 32.3 |
| **24** | 123.13 | 75.7 | 157.0 | 124.6 | 154.7 | 65.6 | 154.6 |
| **25** | 128.42 | 150.6 | 73.8 | 128.6 | 72.9 | 62.6 | 72.9 |
| **26** | 19.06 | 109.7 | 29.4/29.5 [a] | 20.8/21.1 [a] | 29.7/30.5 [a] | 21.0/21.6 [a] | 29.8/30.5 [a] |
| **27** | 19.62 | 19.6 | 29.4/29.5 [a] | 20.8/21.1 [a] | 29.7/30.5 [a] | 21.0/21.6 [a] | 29.8/30.5 [a] |
| **28** | 18.45 | 27.9 | 106.8 | 18.5 | 110.9 | 19.7 | 111.0 |

[a] Signals interchangeable.

**Supplementary Table 6. Comparison of NMR data of (22*R*)-ergosta-5,24(28)-diene-3β,22,25-triol (phyministerol A) (12) with literature data[25].**

Discrepancies besides systematic differences in chemical shifts are highlighted in red and described in the footnotes.

| Pos. | Our $^{13}C$ data $C_5D_5N$ 151 MHz, 298 K | Reference $^{13}C$ data[25] $C_5D_5N$ 125 MHz | Shift Δ (ppm) | Our $^{1}H$ data $C_5D_5N$ 600 MHz, 298 K | Reference $^{1}H$ data[25] $C_5D_5N$ 500 MHz | Shift Δ (ppm) |
|---|---|---|---|---|---|---|
| **1** | 38.3 | 37.6 | 0.7 | 1.17 m<br>1.87 m | 1.13 m<br>1.83 m | 0.04<br>0.04 |
| **2** | 33.1 | 32.4 | 0.7 | 1.85 m [b]<br>2.11 m | 1.11 m<br>2.07 m | 0.74<br>0.04 |
| **3** | 71.7 | 71.0 | 0.7 | 3.88 m | 3.83 m | 0.05 |
| **4** | 44.0 | 43.3 | 0.7 | <br>2.66 m | 2.59 m<br>2.61 m | <br>0.05 |
| **5** | 142.4 | 141.8 | 0.6 | | | |
| **6** | 121.6 | 121.0 | 0.6 | 5.45 d (5.0) | 5.41 d (5.0) | 0.04 |
| **7** | 32.6 | 32.0 | 0.6 | 1.62 m [b]<br>1.98 m | <br>1.96 m | <br>0.02 |
| **8** | 32.6 [a] | 32.7 [a] | 0.1 | 1.45 m [b] | 1.56 m | 0.11 |
| **9** | 51.0 | 50.3 | 0.7 | 1.01 m | 0.99 m | 0.02 |
| **10** | 37.3 | 36.9 | 0.4 | | | |
| **11** | 21.8 | 21.2 | 0.6 | 1.44 m<br>1.54 m | 1.39 m<br>1.49 m | 0.05<br>0.05 |
| **12** | 40.4 | 39.8 | 0.6 | 1.19 m<br>2.04 m | 1.17 m<br>2.01 m | 0.02<br>0.03 |
| **13** | 43.3 | 42.6 | 0.7 | | | |
| **14** | 56.9 | 56.3 | 0.6 | 0.97 m | 0.96 m | 0.02 |
| **15** | 25.1 | 24.5 | 0.6 | 1.07 m<br>1.60 m | 1.04 m<br>1.57 m | 0.03<br>0.03 |
| **16** | 28.3 | 27.6 | 0.7 | 1.42 m<br>1.94 m | 1.39 m<br>1.91 m | 0.03<br>0.03 |
| **17** | 54.1 | 53.5 | 0.6 | 1.30 m | 1.27 m | 0.03 |
| **18** | 12.4 | 11.8 | 0.6 | 0.67 s | 0.65 s | 0.02 |
| **19** | 20.0 | 19.4 | 0.6 | 1.07 s | 1.03 s | 0.04 |
| **20** | 43.4 | 42.7 | 0.7 | 2.04 m | 2.00 m | 0.04 |
| **21** | 13.4 | 12.8 | 0.6 | 1.24 d (6.7) | 1.21 d (7.0) | 0.03 |
| **22** | 75.1 | 74.4 | 0.7 | 4.19 m | 4.16 m | 0.03 |
| **23** | 33.3 [a] | 32.0 [a] | 1.3 | 2.49 d (14.2) [b]<br>2.64 m [b] | 1.86 m<br>1.96 m | 0.63<br>0.68 |
| **24** | 157.5 | 156.9 | 0.6 | | | |
| **25** | 72.5 | 71.9 | 0.6 | | | |
| **26** | 30.7 | 30.1 | 0.6 | 1.67 s | 1.64 s | 0.03 |
| **27** | 31.4 | 30.8 | 0.6 | 1.61 s | 1.58 s | 0.03 |
| **28** | 110.0 | 109.5 | 0.5 | 5.22 br s<br>5.36 br s | 5.19 br s<br>5.33 br s | 0.03<br>0.03 |

[a] Possibly swapped assignments of C8 and C23 in ref. [25]. Our assignments are supported by HMBC correlations.

[b] Our proton assignments are supported by HSQC and COSY correlations.

**Supplementary Table 7. RNA-seq SRA IDs used for structural annotation.**

| | |
|---|---|
| *Withania somnifera* | SRR8382569, SRR8382570, SRR8382571, SRR1012863, SRR1012864, , SRR1019197, SRR1019198, SRR1197573, SRR1197746, SRR8205409, SRR8205410, SRR14739595, SRR14739596, SRR14739597, SRR14739598, SRR520142, SRR520143, SRR10985100, SRR10985101, SRR10985102, SRR10985103, SRR10985104, SRR10985105, SRR12022208, SRR12022209, SRR12022210, SRR12022211, SRR12022212, SRR12022213, SRR12022214, SRR12022215, SRR12022216, SRR12022217, SRR12022218, SRR12022219, SRR12022220, SRR12022221, SRR12022222, SRR12022223, SRR12022224, SRR12022225, SRR12022226, SRR12022227, SRR12022228, SRR12022229, SRR12022230, SRR12022231, SRR12052462, SRR12052463, SRR12052464, SRR12052465, SRR12052466, SRR12052467 |
| *Physalis grisea* | SRR20680386, SRR20680387, SRR20680388, SRR20680389, SRR20680390, SRR20680391, SRR20680392, SRR20680393, SRR20680394, SRR20680395, SRR20680396, SRR20680397, SRR20680398, SRR20680399, SRR20680400, SRR20680401, SRR20680402, SRR20680403, SRR20680404, SRR20680405, SRR20680406, SRR20680407, SRR20680408, SRR20680409, SRR20680410, SRR20680411, SRR20680412, SRR20680413, SRR20680414, SRR20680415, SRR20680416, SRR20680417, SRR20680418, SRR20680419, SRR20680420, SRR20680421, SRR20680422, SRR20680423, SRR20680424, SRR2068042 |
| *Physalis pruinosa* | SRR7066585, SRR7066587, SRR7066588, SRR7066589, SRR7066590, SRR7066591, SRR7066592 |

**Supplementary Table 8. Source of genomic data sets used for gene prediction, synteny analysis and phylogenetic tree building.**

Note: The gene annotation sequences of *Withania somnifera*, *Physalis grisea* and *Physalis pruinosa* generated in this study are used in the synteny and phylogenetic analyses as well**.**

| Species | Used for | | | Reference |
|---|---|---|---|---|
| | Gene prediction | Synteny analysis | Phylogenetic trees | |
| *Atropa belladonna* | ✓ | | ✓ | [26] |
| *Capsicum annuum* | | | ✓ | [27] |
| *Capsicum baccatum* | | | ✓ | [28] |
| *Capsicum chinense* | | | ✓ | [28] |
| *Datura stramonium* | ✓ | ✓ | ✓ | [26] |
| *Datura wrightii* | | ✓ | ✓ | [29] |
| *Iochroma cyaneum* | ✓ | | ✓ | [30] |
| *Ipomoea nil* | | | ✓ | [31] |
| *Lycium barbarum* | ✓ | | ✓ | [32] |
| *Nicotiana attenuata* | | | ✓ | [33] |
| *Nicotiana otophora* | | | ✓ | [34] |
| *Nicotiana sylvestris* | | | ✓ | [35] |
| *Nicotiana tabacum* | | ✓ | ✓ | [34] |
| *Nicotiana tomentosiformis* | | | ✓ | [35] |
| *Petunia inflata* | | | ✓ | [36] |
| *Physalis floridana* | ✓ | ✓ | ✓ | [37] |
| *Solanum appendiculatum* | | | ✓ | [38] |
| *Solanum chilense* | | | ✓ | [39] |
| *Solanum clarkiae* | | | ✓ | GCA_011800125.2 |
| *Solanum dulcamara* | | | ✓ | [40] |
| *Solanum lycopersicum* | ✓ | ✓ | ✓ | [41] |
| *Solanum melongena* | | | ✓ | [42] |
| *Solanum okadae* | | | ✓ | [43] |
| *Solanum pennellii* | | | ✓ | [44] |
| *Solanum pimpinellifolium* | | | ✓ | [45] |
| *Solanum sitiens* | | | ✓ | [46] |
| *Solanum stenotomum* | | | ✓ | [47] |
| *Solanum tuberosum* | | | ✓ | [48] |
| *Solanum verrucosum* | | | ✓ | GCF_900185275.1 |

**Supplementary Table 9. List of yeast strains.**

*At*: *A. thaliana*; *Pp*: *P. pruinosa*; *Pper*: *P. peruviana*; $^{opt}$: codon-optimised for yeast.

| Strain | Genotype | Source |
|---|---|---|
| ST7574 | MATa; *HIS3; TRP1; LEU2; TRP1; URA3; MAL2-8C SUC2* + pCfB2312 (2µm Cas9 KanMX) | Euroscarf |
| KMY14 | ST7574 XI-1:: <-$T_{ADH1}$-*Pper7RED*-$P_{PGK1}$-$P_{TEF1}$-*Pper24ISO*-$T_{CYC1}$-> | This work |
| KMY23 | KMY14 + *erg5-Δ1 erg4-Δ1* | This work |
| KMY48 | ST7574 + *erg5-Δ1 erg4-Δ1* | This work |
| KMY50 | KMY23 + X-4:: <-$T_{ADH1}$-*Pper7RED*-$P_{PGK1}$-$P_{GPD1}$-*upc2-1*-$T_{CYC1}$-> | This work |
| KMY53 | KMY50 + XII-5:: <-$T_{ADH1}$-*YEH2*-$P_{FBA1}$-$P_{TDH3}$-*ARE2*-$T_{CYC1}$-> | This work |
| KMY54 | KMY50 + XII-5:: <-$T_{ADH1}$-*YEH1*-$P_{TEF1}$-$P_{TDH3}$-*ARE2*-$T_{CYC1}$-> | This work |
| KMY55 | KMY23 + XI-3:: <-$T_{ADH1}$-*PpCYP87G1*-$P_{ADH2}$-$P_{TEF1}$-*AtATR1*-$T_{CYC1}$-> | This work |
| KMY71 | KMY23 + X-4:: $P_{GAL1}$-*AtATR1*-$T_{CYC1}$-> | This work |
| KMY75 | KMY71 + XI-3:: <-$T_{ADH1}$-*PpCYP87G1*$^{opt}$-$P_{TEF1}$-$P_{TDH3}$-*PpCYP88C7*$^{opt}$-$T_{CYC1}$-> | This work |
| KMY77 | KMY23 + *Pper24iso-Δ1* | This work |
| KMY78 | KMY55 + *Pper24iso-Δ1* | This work |
| KMY83 | KMY71 + XII-2:: <-$T_{ADH1}$-*PpCYP87G1*$^{opt}$-$P_{TEF1}$ | This work |
| KMY84 | KMY71 + XII-2:: $P_{TDH3}$-*PpCYP88C7*$^{opt}$-$T_{CYC1}$-> | This work |

**Supplementary Table 10. List of primers for metabolic engineering in *S. cerevisiae*.**

Overhangs for USER-cloning assembly are bold and underlined.

| Name | Sequence | Purpose |
|---|---|---|
| PGK1-TEF1-1975-F | **ACCTGCACU**TTGTAATTAAAACTTAGATTAGATTG | Amplification of bidirectional promoter $P_{PGK1}$ and $P_{TEF1}$ from pCfB1975 [49] |
| PGK1-TEF1-1975-R | **ATGACAGATU**TGTTTTATATTTGTTGTAAAAAGTAG | |
| BB-PGK1-F | **ACCTGCACU**GAAGTACCTTCAAAGAATGG | Amplification of $P_{PGK1}$ promoter from plasmid pKME15 for USER assembly in pKME61 |
| BB-7RED-R | **CGTGCGAU**CTAGTAAATTCCGGGTACGAC | Amplification of *Pper7RED* from plasmid pKME15 for USER assembly in pKME61 |
| BB-GPD1-F | **AGTGCAGGU**CACTTCTACTTCTACATCG | Amplification of $P_{GPD1}$ promoter from *S. cerevisiae* WAT11 [50] genomic DNA for USER assembly in pKME61 |
| BB-GPD1-R | **ATCAGCAGU**CTTTATATTATCAATATTTGTGTTTGTGG | |
| BB-FBA1-F | **ACCTGCACU**TGGGTCATTACGTAAATAATGATAGG | Amplification of $P_{FBA1}$ promoter from *S. cerevisiae* WAT11 [50] genomic DNA for USER assembly in pKME62 |
| BB-FBA1-R | **ATGACAGATU**TTTGAATATGTATTACTTGGTTATGG | |
| BB-TDH3-TEF1-1977-F | **ACCTGCACU**TTTGTTTGTTTATGTGTGTTTATTC | Amplification of bidirectional promoter $P_{TDH3}$ and $P_{TEF1}$ from p1977[51] for USER assembly in pKME64 and pKME74 |
| BB-TDH3-TEF1-1977-F | **ATGACAGAU**TTGTAATTAAAACTTAGATTAGATTG | |
| BB-TDH3-F | **AGTGCAGGU**ATAAAAAACACGCTTTTTCAG | *E. coli*: Amplification of $P_{TDH3}$ promoter from p1977[51] for USER assembly in pKME62 |
| BB-TDH3-R | **ATCAGCAGU**TTTGTTTGTTTATGTGTGTTTATTCG | |
| BB-ADH2-F | **ACCTGCACU**TATCTAAAAATTGCCTTATGATCCGTC | Amplification of $P_{ADH2}$ promoter from *S. cerevisiae* WAT11 [50] genomic DNA for USER assembly in pKME55 |
| BB-ADH2-R | **ACTGCTGAU**TGTGTATTACGATATAGTTAATAGTTGATAG | |
| BB-Pper7RED-F | **AATCTGTCAU**ATGGGGGAGTCTCAGTTG | Amplification of *Pper7RED* from *P. peruviana* cDNA for USER assembly in KME15 |
| BB-Pper7RED-R | **CGTGCGAU**CTAGTAAATTCCGGGTACGAC | |
| BB-24ISO-F | **AGTGCAGGU**ATGTCAGGGGAGAAGGTGG | Amplification of *Pper24ISO* for USER assembly in KME15 |
| BB-24ISO-R | **CACGCGAU**TCAATCCGCGGGCTCATCG | |
| BB-ATR1-F | **ATCTGTCAU**ATGACTTCTGCTTTGTATGCTTCCGATTTG | Amplification of *ATR1* from *A. thaliana* cDNA for USER assembly in pKME73 |
| BB-ATR1-R | **CACGCGAU**TCACCAGACATCTCTGAGGTATCTTC | |
| BB-Gal1-F | **CGTGCGAU**ACGGATTAGAAGCCGCCGA | Amplification of $P_{GAL1}$ promoter from pYES2 vector for USER assembly in pKME73 |
| BB-Gal1-R | **ATGACAGAU**GGTTTTTTCTCCTTGACGTTAAAG | |
| BB-upc2-1-F | **ACTGCTGAU**ATGAGCGAAGTCGGTATACAGAATCACAAGAAAGCGGTG | Amplification of *upc2-1* from *S. cerevisiae* WAT11 [50] genomic DNA for USER assembly in pKME61 |
| BB-upc2-1-R | **CACGCGAU**TCATAACGAAAAATCAGAGAAATTTGTTGTTGTCATCGATGGTAATCCGCCACCGAGGAAATCTAGCATCATATGCATATCACCACCTCCACTGTATTCGTCAACATC | |
| BB-ARE2-F | **AGTGCAGGU**ATGGACAAGAAGAAGGATCTACTGG | *E. coli:* Amplification of *ARE2* from *S. cerevisiae* WAT11 [50] genomic DNA for USER assembly in pKME62 |
| BB-ARE2-R | **CACGCGAU**TTAGAATGTCAAGTACAACGTACACATG | |
| BB-YEH1-F | **ATCTGTCAU**ATGGGTGTTTCTGCGGTGTTGAAAAGAGC | Amplification of *YEH1* for USER assembly in pKME64 |
| BB-YEH1-R | **CGTGCGAU**TCAAGCCTTCTCAGCAACCATTTCC | |
| BB-YEH2-F | **AATCTGTCAU**ATGGTAAATAAGGTGGTTGATGAGG | Amplification of *YEH2* for USER assembly in pKME62 |
| BB-YEH2-R | **CGTGCGAU**TCACCTTGCATTAGGAAACCTCAAATTTTC | |
| BB- Pp-CYP87G1-F | **ATCTGTCAU**ATGTGGAATATTATTTTGTGTATTGTTGG | Amplification of *PpCYP87G1* for USER assembly in pKME74 |
| BB- Pp-CYP87G1-R | **CGTGCGAU**TTAAGCAACATTAATTTCTTTTTTATTATCTCTTTGG | |

| | | |
|---|---|---|
| BB- Pp-CYP88C7-F | **AGTGCAGGU**ATGGAATATAATTTGGTTTTTTTGTC TACC | Amplification of *PpCYP88C7* for USER assembly in pKME74 |
| BB- Pp-CYP88C7-R | **CACGCGAU**TTAAGCAGAAATTTTTTTAAATCTAGC AATAC | |
| ERG4-gRNA-F | ATTTACCCTGGTTTTAGAGCTAGAAATAGC | Primers for *ERG4* gRNA incorporation in pCfB8792 [51] |
| ERG4-gRNA-R | GGCTGAAGCCGATCATTTATCTTTCACTGCG | |
| ERG5-gRNA-F | GAATTATGAGGTTTTAGAGCTAGAAATAGC | Primers for *ERG5* gRNA incorporation in pCfB8792 [51] |
| ERG5-gRNA-R | TACGCTACACGATCATTTATCTTTCACTGCG | |
| 24ISO-gRNA-F | TAGATCTGCGGTTTTAGAGCTAGAAATAGC | Primers for *24ISO* gRNA incorporation in pCfB8792 [51] |
| 24ISO-gNRA-R | ACGTCCCCAACGATCATTTATCTTTCACTGCG | |
| ERG4-HDR-F | TAGATAGGCAGATACGGATATTTACGTAGTGTACA TAGATTAGCATCGCTTTATTTTC | Primers for donor-repair template generation for *ERG4* deletion |
| ERG4-HDR-R | ATATACAAACTGTAAAATAAGTTAATGAAGTGGA TAGAAAAAGAAAATAAAGCGATGCTAATC | |
| ERG5-HDR-F | AAAACATCACATTTTGCTATTCCAATAGACAATAA ATACCTTTTAACAAATG | Primers for donor-repair template generation for *ERG5* deletion |
| ERG5-HDR-R | GAAGTAAATATGATTTATTGTCTGGACAAAGTTCT GTTTTTCCCCATTTGTTAAAAGGTATTTATTG | |
| 24ISO-HDR-F | CATAGCAATCTAATCTAAGTTTTAATTACAAAGTG CAGGTATCGCGTGCATTCATCCGCT | Primers for donor-repair template generation for *Pper24ISO* deletion |
| 24ISO-HDR-R | AGGTTGTCTAACTCCTTCCTTTTCGGTTAGAGCGG ATGAATGCACGCGATACCTGCAC | |
| ERG4-del-F | TAGTGCTGAACGTGCACTCACATGAATATATC | Primers for confirming *ERG4* deletion |
| ERG4-del-R | ATGTATTGGCTAAGAGAATGGCTAACC | |
| ERG5-del-F | TACGAAGCAAGAGTAGCTAAGCGTGAAGC | Primers for confirming *ERG5* deletion |
| ERG5-del-R | ACGAACCTAGTGAAGCAGAACCTGTCAAG | |
| 24ISO-del-F | GCTCATTAGAAAGAAAGCATAGC | Primers for confirming *Pper24ISO* deletion |
| 24ISO-del-R | GAAGACCCATGGTTCCAAGGA | |

**Supplementary Table 11. List of plasmids for metabolic engineering in *S. cerevisiae*.**

| Strain | Description | Reference |
|---|---|---|
| pCfB8622 | Plasmid containing gRNA expression cassette targeting *ADE2* in *S. cerevisiae* (source plasmid for gRNA modifications) | [51] |
| pCfB3035 | Plasmid for integration of genes and promoters into site X-4 | [49] |
| pCfB3036 | Plasmid for integration of genes and promoters into site XI-1 | [49] |
| pCfB2904 | Plasmid for integration of genes and promoters into site XI-3 | [49] |
| pCfB2909 | Plasmid for integration of genes and promoters into site XII-5 | [49] |
| pCfB3039 | Plasmid for integration of genes and promoters into site XII-2 | [49] |
| pKME15 | pCfB3036- $T_{ADH1}$-*Pper7RED*-$P_{PGK1}$-$P_{TEF1}$-*Pper24ISO*-$T_{CYC1}$ | This work |
| pKME22 | Plasmid containing gRNA expression cassette targeting *ERG4* in ST7574 | This work |
| pKME24 | Plasmid containing gRNA expression cassette targeting *ERG5* in ST7574 | This work |
| pKME56 | Plasmid containing gRNA expression cassette targeting *Pper24ISO* in KMY23 | This work |
| pKME61 | pCfB3035-$T_{ADH1}$-*Pper7RED*-$P_{PGK1}$-$P_{GPD1}$-*upc2-1*-$T_{CYC1}$ | This work |
| pKME62 | pCfB2909-$T_{ADH1}$-*YEH2*-$P_{FBA1}$-$P_{TDH3}$-*ARE2*-$T_{CYC1}$ | This work |
| pKME63 | pCfB2904-$T_{ADH1}$-*PpCYP87G1*-$P_{ADH2}$-$P_{TEF1}$-*AtATR1*-$T_{CYC1}$ | This work |
| pKME64 | pCfB2909-$T_{ADH1}$-*YEH1*-$P_{TEF1}$-$P_{TDH3}$-*ARE2*-$T_{CYC1}$ | This work |
| pKME73 | pCfB3035-$P_{GAL1}$-*AtATR1*-$T_{CYC1}$ | This work |
| pKME74 | pCfB2904-$T_{ADH1}$-*PpCYP87G1*$^{opt}$-$P_{TEF1}$-$P_{TDH3}$-*PpCYP88C7*$^{opt}$ -$T_{CYC1}$ | This work |
| pKME81 | pCfB3039-$T_{ADH1}$-*PpCYP87G1*$^{opt}$ -$P_{TEF1}$ | This work |
| pKME82 | pCfB3039-$P_{TDH3}$-*PpCYP88C7*$^{opt}$-$T_{CYC1}$ | This work |

**Supplementary Table 12. Nucleotide sequences for metabolic engineering in *S. cerevisiae* used in this study.**

| Gene | Nucleotide sequence (5' - 3') |
|---|---|
| *P. peruviana 7RED* | ATGGGGGAGTCTCAGTTGGCACATCCTCCTTTATTCACTTATGTTTCAATGCTCACTCTTCTTACTATAGTACCCCCTTTTA<br>TTATTCTCATGTGGTATACAAATGTTCATGCTGATGGATCTATATTGAACACATTTAATTACCTAAAGGAGAATGGCCCGCA<br>AGGACTAATTGACATTTGGCCAAGACCCACGGCAGTTGCGGAAAAATAATAATTTGCTATGGTCTATTTGAGGCTGCACTT<br>CAGATTTTGTTGCCTGGTAAAAGGGTCGAAGGACCAATATCTCCAACTGGACACAGGCCTGTCTATAAGGCAAATGGAGTGG<br>CAGCATATACAGTGACGCTAATTACCTATCTCAGTCTTTGGTGGTTTGGAATATTCAATCCTACCATCGTGTATGATCATCT<br>GGGAGAGATTCTCTCTACACTAAATATTGGAAGCTTAATCTTCTGTCTCCTCTTATACATAAAAGGTCATGTTGCACCATCT<br>TCCACTGATCACGGTTCATCAGGAAACATAATAATCGACTTCTACTGGGGGATGGAGCTATATCCTCGCATTGGCAAACACT<br>TTGATATCAAGGTCTTCACAAACTGCAGATTTGGCTTGATTTCTTGGGCACTCCTCCCTATTACCTACTGTATAAAGCAGTA<br>TGAAGAATATGGAAGTCTCTCTGATTCCATGCTTGTACATACCATAATAACGTTGGTGTATGTCACGAAATTCTTTTGGTGG<br>GAAGCAGGTTACTGGAACACCATGGATATTGCACACGACAGAGCCGGTTTTTACATATGCTGGGGTTGCTTGGTATTTCTTC<br>CATGTATATATACTTCTCCTGGCATGTACCTTGTCAAACATCCTGTAAATCTTGGACCTCAGCTTGCACTTTCTATTCTAGC<br>AGCTGGAATTCTCTGTGTATACATAAACTATGATTGTGACAGACAGAGGCAAGAGTTTCGCAGAACAAATGGCAAAGCCCCT<br>GTCTGGGGAAAGGCTCCATCGAAGATTGTTGCCTCATACACTACCACTACTGGTGAGACCAAGACTAGCATTCTCCTGACAT<br>CTGGATGGTGGGGCTTAGCACGACACTTTCATTATGTTCCAGAAATACTAGCTTCATTTTTCTGGTGTGTGCCAGCTCTTTT<br>CAACCATTTCATTCCCTACTTCTACGTAGTCTATCTGACAGTCCTCCTTCTTGATCGAGCCAAAAGGGATGACGAGCGGTGC<br>AAGGCAAAGTACGGCAAATACTGGAAAAAGTATTGCGAAAAGGTGCCTTACCGGGTCGTACCCGGAATTTACTAG |
| *P. peruviana 24ISO* | ATGTCAGGGGAGAAGGTGGCGACTGTTCGTCCCAAGAGGGAGACGCGGTTGGTGGACTTTCTTGTCCAATTCAGATGGATCG<br>TTGTTATATTCTTCGTCCTTCCTTTCTCGTGCTTGTACTACTTCTCCATATACCTTGGGGACGTTAGATCTGCGCGGAAATC<br>TGAAAGCCAACGTAAGAAGGAACACGAAGAAAACGTTAAAGAGGTTGTGAAGCGCCTTACACAGAGAGATGCATCAAAGGAT<br>GGCCTTGTTTGCACAGCCAGGCCTCCCTGGACGGTTATTGGAATGAGAAATTGTGACTATAAGCGTGCTCGTAGTTTCAAAG<br>TTGATCTTTCCAAGTTTAGAAATATTCTTGAAATCAACAAGGAACGAATGGTTGCAAAAGTTGAACCTCTTGTCAATATGGC<br>CCAAATCTCTAGAGCTACTATACCCATGAATCTTTCGCTTGCAGTTATTGGCGAGCTTGATGATCTGACTGTTGGTGGTTTG<br>ATCAATGGATTCGGGATTGAAGGAAGCTCTCACGTCTATGGATTGTTCTCCGACACAATAGTGGCACTCGAGGTTGTTCTGG<br>CTGATGGACGGGTGGTTAGAGCTACAAAGGACAACGAGTACTCCGATCTTTTCTATGCTATCCCATGGTCACAGGGGACATT<br>GGGGCTTCTTGTTTCAGCCGAAATTAAGCTCATACCTGTGAAGGAATACATGAAGCTTACCTACAAACCTGTAACTGGTAAT<br>CTAAAAGAGCTAGCACAGGCCTTTGCAGATTCCTTTGCACCGAGAGACGGAGACCAGGATAATCCTAGTAAAGTTCCAGACT<br>TTGTTGAAGGCATGATTTACAGTCCCACGGAAGGTGTTATGATGACGGGTAGATACGCTTCAACAAAAGAGGCCAAGCAAAA<br>GGGCAACGTAATCAATAGTATCGGTTGGTGGTTCAAGCCATGGTTTTACCAGCATGCTCAAACTGCACTGAAAAGAGGGGAG<br>TTTGTAGAGTACATTCCAACTAGAGATTACTGTCACAGGCACACAAGATCCATCTACTGGGAAGGCAAATTAATTCTTCCAT<br>TTGGGGATCAGTTCTGGTTTAGGTATCTCTTTGGATGGATGATGCCTCCCAAGGTTGCTGTGCTTAAGGCCACTCAAAGTGA<br>GTATATCAGGAATTATTACCATGACAATCATGTCATTCAGGATTTGCTCGTTCCTCTTCACAAGGTCGCTGATACTCTCGAG<br>TGGGTCCACCGCGAGATGGAGGTATATCCTGTGTGGCTCTGCCCACACAGAATCTACAAGCTGCCTGTGAAACCTATGATCC<br>ATCCTGAACCAGGATTCGAGGCACACAAGAGACAGGGCGACACTAAATATGCCCAAATGTATACTGATGTTGGCGTCTACTA<br>TGCTCCTGGAGCAGTCTTAAGGGGCGAGCCCTATGATGGCGCAAAGAAATGCCATGACGTGGAGATTTTTCTAATCGAAAAC<br>CATGGATTCCAGCCTCAATACGCGGTGACTGAGCTATCAGAGAAGAACTTCTGGAGGATGTTTGATGCAACCTTATACCAGC<br>AATGTAGAAAAAAGTACAAAGCCATTGGAACATTCATGAACGTGTACTATAAATGCAAGAAAGGGAAGAAGACGGAGAAGGA<br>AGTGCAAGAAGCTGAGCAAGAGAAAGCTGAGCTTGACACTCCCGAGGTCGATGAGCCCGCGGATTGA |
| *A. thaliana ATR1* | ATGACTTCTGCTTTGTATGCTTCCGATTTGTTTAAGCAGCTCAAGTCAATTATGGGGACAGATTCGTTATCCGACGATGTTG<br>TACTTGTGATTGCAACGACGTCTTTGGCACTAGTAGCTGGATTTGTGGTGTTGTTATGGAAGAAAACGACGGCGGATCGGAG<br>CGGGGAGCTGAAGCCTTTGATGATCCCTAAGTCTCTTATGGCTAAGGACGAGGATGATGATTTGGATTTGGGATCCGGGAAG<br>ACTAGAGTCTCTATCTTCTTCGGTACGCAGACTGGAACAGCTGAGGGATTTGCTAAGGCATTATCCGAAGAAATCAAAGCGA<br>GATATGAAAAAGCAGCAGTCAAAGTCATTGACTTGGATGACTATGCTGCCGATGATGACCAGTATGAAGAGAAATTGAAGAA<br>GGAAACTTTGGCATTTTTCTGTGTTGCTACTTATGGAGATGGAGAGCCTACTGACAATGCTGCCAGATTTTACAAATGGTTT<br>ACGGAGGAAAATGAACGGGATATAAAGCTTCAACAACTAGCATATGGTGTGTTTGCTCTTGGTAATCGCCAATATGAACATT<br>TTAATAAGATCGGGATAGTTCTTGATGAAGAGTTATGTAAGAAAGGTGCAAAGCGTCTTATTGAAGTCGGTCTAGGAGATGA<br>TGATCAGAGCATTGAGGATGATTTTAATGCCTGGAAAGAATCACTATGGTCTGAGCTAGACAAGCTCCTCAAAGACGAGGAT<br>GATAAAAGTGTGGCAACTCCTTATACAGCTGTTATTCCTGAATACCGGGTGGTGACTCATGATCCTCGGTTTACAACTCAAA<br>AATCAATGGAATCAAATGTGGCCAATGGAAATACTACTATTGACATTCATCATCCCTGCAGAGTTGATGTTGCTGTGCAGAA<br>GGAGCTTCACACACATGAATCTGATCGGTCTTGCATTCATCTCGAGTTCGACATATCCAGGACGGGTATTACATATGAAACA<br>GGTGACCATGTAGGTGTATATGCTGAAAATCATGTTGAAATAGTTGAAGAAGCTGGAAAATTGCTTGGCCACTCTTTAGATT<br>TAGTATTTTCCATACATGCTGACAAGGAAGATGGCTCCCCATTGGAAAGCGCAGTGCCGCCTCCTTTCCCTGGTCCATGCAC<br>ACTTGGGACTGGTTTGGCAAGATACGCAGACCTTTTGAACCCTCCTCGAAAGTCTGCGTTAGTTGCCTTGGCGGCCTATGCC<br>ACTGAACCAAGTGAAGCCGAGAAACTTAAGCACCTGACATCACCTGATGGAAAGGATGAGTACTCACAATGGATTGTTGCAA<br>GTCAGAGAAGTCTTTTAGAGGTGATGGCTGCTTTTCCATCTGCAAAACCCCCACTAGGTGTATTTTTTGCTGCAATAGCTCC<br>TCGTCTACAACCTCGTTACTACTCCATCTCATCCTCGCCAAGATTGGCGCCAAGTAGAGTTCATGTTACATCCGCACTAGTA<br>TATGGTCCAACTCCTACTGGTAGAATCCACAAGGGTGTGTGTTCTACGTGGATGAAGAATGCAGTTCCTGCGGAGAAAAGTC<br>ATGAATGTAGTGGAGCCCCAATCTTTATTCGAGCATCTAATTTCAAGTTACCATCCAACCCTTCAACTCCAATCGTTATGGT<br>GGGACCTGGGACTGGGCTGGCACCTTTTAGAGGTTTTCTGCAGGAAAGGATGGCACTAAAAGAAGATGGAGAAGAACTAGGT<br>TCATCTTTGCTCTTCTTTGGGTGTAGAAATCGACAGATGGACTTTATATACGAGGATGAGCTCAATAATTTTGTTGATCAAG<br>GCGTAATATCTGAGCTCATCATGGCATTCTCCCGTGAAGGAGCTCAGAAGGAGTATGTTCAACATAAGATGATGGAGAAGGC<br>AGCACAAGTTTGGGATCTAATAAAGGAAGAAGGATATCTCTATGTATGCGGTGATGCTAAGGGCATGGCGAGGGACGTCCAC<br>CGAACTCTACACACCATTGTTCAGGAGCAGGAAGGTGTGAGTTCGTCAGAGGCAGAGGCTATAGTTAAGAAACTTCAAACCG<br>AAGGAAGATACCTCAGAGATGTCTGGTGA |
| *upc2-1* | ATGAGCGAAGTCGGTATACAGAATCACAAGAAAGCGGTGACAAAACCCAGAAGAAGAGAAAAAGTCATCGAGCTAATTGAAG<br>TGGACGGCAAAAAGGTGAGTACGACTTCAACCGGTAAACGTAAATTCCATAACAAATCAAAGAATGGGTGCGATAACTGTAA<br>AAGAAGAAGAGTTAAGTGTGATGAAGGGAAGCCAGCCTGTAGGAAGTGCACAAATATGAAGTTGGAATGTCAGTATACACCA<br>ATCCATTTAAGGAAAGGTAGAGGAGCAACAGTAGTGAAGTATGTCACGAGAAAGGCAGACGGTAGCGTGGAGTCTGATTCAT |

| | |
|---|---|
| | CGGTAGATTTACCTCCTACGATCAAGAAGGAGCAGACACCGTTCAATGATATCCAATCAGCGGTAAAAGCTTCAGGCTCATC<br>CAATGATTCCTTTCCATCAAGCGCCTCTACAACTAAGAGTGAGAGCGAGGAAAAGTCATCGGCCCCTATAGAGGACAAAAAC<br>AATATGACTCCTCTAAGTATGGGCCTCCAGGGTACCATCAATAAGAAAGATATGATGAATAACTTTTTCTCTCAAAATGGCA<br>CTATTGGTTTTGGTTCTCCTGAAAGATTGAATTCAGGTATCGATGGCTTACTATTACCGCCATTGCCTTCTGGAAATATGGG<br>TGCGTTCCAACTTCAGCAACAGCAGCAAGTGCAGCAGCAATCTCAACCACAGACCCAAGCGCAGCAAGCAAGTGGAACTCCA<br>AACGAGAGATATGGTTCATTCGATCTTGCGGGTAGTCCTGCATTGCAATCCACGGGAATGAGCTTATCAAATAGTCTAAGCG<br>GGATGTTACTATGTAACAGGATTCCTTCCGGCCAAAACTACACTCAACAACAATTACAATATCAATTACACCAGCAGCTGCA<br>ATTGCAACAGCATCAGCAAGTTCAGCTGCAGCAGTATCAACAATTACGTCAGGAACAACACCAACAAGTTCAGCAACAACAA<br>CAGGAACAACTCCAGCAATACCAACAACATTTTTTGCAACAGCAGCAACAAGTACTGCTTCAGCAAGAGCAACAACCTAACG<br>ATGAGGAAGGTGGCGTTCAGGAAGAAAACAGCAAAAAGGTAAAGGAAGGGCCTTTACAATCACAAACAAGCGAAACTACTTT<br>AAACAGCGATGCTGCTACATTACAAGCTGATGCATTATCTCAGTTAAGTAAGATGGGGCTAAGCCTAAAGTCGTTAAGTACC<br>TTTCCAACAGCTGGTATTGGTGGTGTTTCCTATGACTTTCAGGAACTGTTAGGTATTAAGTTTCCAATAAATAACGGCAATT<br>CAAGAGCTACTAAGGCCAGCAACGCAGAGGAAGCTTTGGCCAATATGCAAGAGCATCATGAACGTGCAGCTGCTTCTGTAAA<br>GGAGAATGATGGTCAGCTCTCTGATACGAAGAGTCCAGCGCCATCGAATAACGCCCAAGGGGGAAGTGCTAGTATTATGGAA<br>CCTCAGGCGGCTGATGCGGTTTCGACAATGGCGCCTATATCAATGATTGAAAGAAACATGAACAGAAACAGCAACATTTCTC<br>CATCAACGCCCTCTGCAGTGTTGAATGATAGGCAAGAGATGCAAGATTCTATAAGTTCTCTAGGAAATCTGACAAAAGCAGC<br>CTTGGAGAACAACGAACCAACGATAAGTTTACAAACATCACAGACAGAGAATGAAGACGATGCATCGCGGCAAGACATGACC<br>TCAAAAATTAATAACGAAGCTGACCGAAGTTCTGTTTCTGCTGGTACCAGTAACATCGCTAAGCTTTTAGATCTTTCTACCA<br>AAGGCAATCTGAACCTGATAGACATGAAACTGTTTCATCATTATTGCACAAAGGTCTGGCCTACGATTACAGCGGCCAAAGT<br>TTCTGGGCCTGAAATATGGAGGGACTACATACCGGAGTTAGCATTTGACTATCCATTTTTAATGCACGCTTTGTTGGCATTC<br>AGTGCCACCCATCTTTCGAGGACTGAAACTGGACTGGAGCAATACGTTTCATCTCACCGCCTAGACGCTCTGAGATTATTAA<br>GAGAAGCTGTTTTAGAAATATCTGAGAATAACACCGATGCGCTAGTTGCCAGCGCCCTGATACTAATCATGGACTCGTTAGC<br>AAATGCTAGTGGTAACGGCACTGTAGGAAACCAAAGTTTGAATAGCATGTCACCAAGCGCTTGGATCTTTCATGTCAAAGGT<br>GCTGCAACAATTTTAACCGCTGTGTGGCCTTTGAGTGAAAGATCTAAATTTCATAACATTATATCTGTTGATCTTAGCGATT<br>TAGGCGATGTCATTAACCCTGATGTTGGAACAATTACTGAATTGGTATGTTTTGATGAAAGTATTGCCGATTTGTATCCTGT<br>CGGCTTAGATTCGCCATATTTGATAACACTAGCTTATTTAGATAAATTGCACCGTGAAAAAAACCAGGGTGATTTTATTCTG<br>CGGGTATTTACATTTCCAGCATTGCTAGACAAGACATTCCTGGCATTACTGATGACAGGTGATTTAGGTGCAATGAGAATTA<br>TGAGATCATATTATAAACTACTTCGAGGATTTGCCACAGAGGTCAAGGATAAAGTCTGGTTTCTCGAAGGAGTCACGCAGGT<br>GCTGCCTCAAGATGTTGACGAATACAGTGGAGGTGGTGGTATGCATATGATGCTAGATTTCCTCGGTGGCGGATTACCATCG<br>ATGACAACAACAAATTTCTCTGATTTTTCGTTATGA |
| *ARE2* | ATGGACAAGAAGAAGGATCTACTGGAGAACGAACAATTTCTCCGCATCCAAAAGCTCAACGCTGCCGATGCGGGCAAAAGAC<br>AATCTATAACAGTGGACGACGAGGGCGAACTATATGGGTTAGACACCTCCGGCAACTCACCAGCCAATGAACACACAGCTAC<br>CACAATTACACAGAATCACAGCGTGGTGGCCTCAAACGGAGACGTCGCATTCATCCCAGGAACTGCTACCGAAGGCAATACA<br>GAGATTGTAACTGAAGAAGTGATTGAGACCGATGATAACATGTTCAAGACCCATGTGAAGACTTTAAGCTCCAAAGAGAAGG<br>CACGGTATAGGCAAGGGTCCTCTAACTTTATATCGTATTTCGATGATATGTCATTTGAACACAGGCCCAGTATATTAGATGG<br>GTCAGTTAACGAGCCCTTCAAGACCAAATTCGTGGGACCTACTTTAGAAAAGGAGATCAGAAGAAGGGAGAAAGAGCTAATG<br>GCCATGCGCAAAAATTTACACCACCGCAAGTCCTCCCCAGATGCTGTCGACTCAGTAGGGAAAAATGATGGCGCCGCCCCAA<br>CTACTGTTCCAACTGCCGCCACCTCAGAAACGGTGGTCACCGTTGAAACCACCATAATTTCATCCAATTTCTCCGGGTTGTA<br>CGTGGCGTTTTGGATGGCTATTGCATTTGGTGCTGTCAAGGCTTTAATAGACTATTATTACCAGCATAATGGTAGCTTCAAG<br>GATTCGGAGATCTTGAAATTTATGACTACGAATTTGTTCACTGTGGCATCCGTAGATCTTTTGATGTATTTGAGCACTTATT<br>TTGTCGTTGGAATACAATACTTATGCAAGTGGGGGGTCTTGAAATGGGGCACTACCGGCTGGATCTTCACCTCAATTTACGA<br>GTTTTTGTTTGTTATCTTCTACATGTATTTAACAGAAAACATCCTAAAACTACACTGGCTGTCCAAGATCTTCCTTTTTTTG<br>CATTCTTTAGTTTTATTGATGAAAATGCATTCTTTCGCCTTCTACAATGGCTATCTATGGGGTATAAAGGAAGAACTACAAT<br>TTTCCAAAAGCGCTCTTGCCAAATACAAGGATTCTATAAATGATCCAAAAGTTATTGGTGCTCTTGAGAAAAGCTGTGAGTT<br>TTGTAGTTTTGAATTGAGCTCTCAGTCTTTAAGCGACCAAACTCAAAAATTCCCCAACAATATCAGTGCAAAAAGCTTTTTT<br>TGGTTCACCATGTTTCCAACCCTAATTTACCAAATTGAATATCCAAGAACTAAGGAAATCAGATGGAGCTACGTATTAGAAA<br>AGATCTGCGCCATCTTCGGTACCATTTTCTTAATGATGATAGATGCTCAAATCTTGATGTATCCTGTAGCAATGAGAGCATT<br>GGCTGTGCGCAATTCTGAATGGACTGGTATATTGGATAGATTATTGAAATGGGTTGGATTGCTCGTTGATATCGTCCCAGGG<br>TTTATCGTGATGTACATCTTGGACTTCTATTTGATTTGGGATGCCATTTTGAACTGTGTGGCTGAATTGACAAGATTTGGCG<br>ACAGATATTTCTACGGTGACTGGTGGAATTGTGTTAGTTGGGCAGACTTCAGTAGAATTTGGAACATCCCAGTGCATAAGTT<br>TTTGTTAAGACATGTTTACCATAGTTCAATGAGTTCATTCAAATTGAACAAGAGTCAAGCAACTTTGATGACCTTTTTCTTA<br>AGTTCCGTCGTTCATGAATTAGCAATGTACGTTATCTTCAAGAAATTGAGGTTTTACTTGTTCTTCTTCCAAATGCTGCAAA<br>TGCCATTAGTAGCTTTAACAAATACTAAATTCATGAGGAACAGAACCATAATCGGAAATGTTATTTTCTGGCTCGGTATCTG<br>CATGGGACCAAGTGTCATGTGTACGTTGTACTTGACATTCTAA |
| *YEH1* | ATGGGTGTTTCTGCGGTGTTGAAAAGAGCTAGGAATTTACTAGCAACGTTCATAGTCTGCTGTTTTATGGCAGTAGTGCTTG<br>TTCTGGCGCTGGCACACCATTTTATAAATGAGCACAGAGACACTAGGAGTTCATCGACCCAAATCGAAGTTGATGACGAAAG<br>TAAAAGAAACGTACACCATGACCATGTTCTCACTAGAACCAATGCATACGCGACGCCATACCTCGATCTGGAACATGACAAG<br>AAAAACGGTATCGTCTACGATCATACAAGAACGGTTGTCCGTAAAAAGAACCACGAAGTGGGGTCCTCGTCACTGCATAAAA<br>ACCTTTTTCACAAATTTTTGACAAAGCTTATTTTTAGGTTTATCGAGAAGGAAAAAGTTACCGAAGGCGTAACGCAAGGAAA<br>GTTCAATAATAGTAGCAATGAAATTGCCAATCATGAACCGGTTTTTGAAAAAATTCCTGTACAGTGCGACAATCCATTACAG<br>AATCTAATCTTATCGGAAGACTTGACATTAGTTGCGGATCTTAATTATTATTTTAACCAGTACAATATTCAAATAGAGGAAT<br>TTAGATTGGAGACCGAAGATGGGTTTGTTATAGATTTGTGGCACTTGATACCAAAATATAGAACGACAGATTCTGACAAGAA<br>GAAGAGGCCACCCATTTTGATGCTACATGGCCTTTTGCAAAGCAGTGGTTCGTTCGCATCCAATGGTAGAAAATCTCTGGCA<br>TATTTTCTGTATCAATCCGGTTACGACATATGGTTAGGGAATAACAGATGCGGGTTTAGGCCGGAATGGAACGAAGCGAAAG<br>TACCGACACTAGCTTCCAGGTGGGACTGGGACCTTCGCGAGATGGTTAAGTACGATCTGACCCTTTTGATTGATACCGTGTT<br>AGCTAAGACGCAGTTTGAAAAGCTTACTTTGATCTCGCATTCTCAGGGCACTACACAGGGGTTTATGGGCTTGGTCAACGAA<br>GATAAGTTTTTCCCTCCCGGTTCGGGATCTAAAGAATCTTTTTTCACTTCTAAGATCGCAAACTATATTGCCTTGGCCCCCG<br>CAGTGTATCCTGGTCCCTTACTTAACGAGAAATTGTTTGTTAAGCTTATGACAAAGGAAATCGAAAATCCCTGGTTCTTTGG<br>TGAAACGAGCTTTTTTGAGATAATGATGATTGTAAGAAACTTGTGCGTTGGTGAGAGCTTGTTCTCCTTTGTTTGTTACACC<br>ATCTTCAATTACCTGTTTGATTGGAACGATACCCTTTGGGATACCGCATTAAGAGATCGCCATTTCCTGTTTTCGCCAGTCC<br>ATGTTTCAGTGAAGTTGATGCAATGGTGGCTGTCACCCGACCCCAACAAGGTAAGTTTAAATTTGGTTCCCATAAGATGTT<br>CCCCGACAATGTTAAGTGGTTTTCAGACGCATCAAAGGCCCCAAATATCTACTTGTTTGTTCCAAAGCAAGATAGATTGGTG<br>GACGGAGAAAGACTAATCAATCATTTCGTCAATGTGGAGTCGAATGTCAACTACAAGATCTGGTACATTGATGAGTATGCCC<br>ATATTGATGTCCTATGGGCACATGATGTCATAGAGAGAATTGGTAAACCAATTTTACAGAATTTGAACAACTATTACTCCAA<br>GAAGCCATCCAGCGCCTTTGAAAGTGATTGTTCGGACACAGAGGTGGAAACGGAGCTGGAAATGGTTGCTGAGAAGGCTTGA |

| | |
|---|---|
| *YEH2* | ATGGTAAATAAGGTGGTTGATGAGGTTCAGCGGTTGGTGAGCGCAATAATTCTCACTTCGTTTATGACGGGGTTATTCATTT<br>TGTCACTATGGAAGAATTACGTGACAGTTCATTTCCAACATAAAAATGATCCAAGAGATACAAGAAGTTCGAGAACCAAGAT<br>CCAGCCAAATGACAAGAAGAAGAAACGACCAGCACGTCACTCTAGACCTTTGTCAATATCTTCCACAACGCCCTTAGATTTA<br>CAGCGGGATCAGGAAAATAATATTGAATATGACCGTACTGTGACTAGCAAGCTGTCCATGACTAGTAATGCCTCTTTATCAG<br>AGAACGGTGATGGAAATGCTAATATTAAAATGGAAACGAATGTGAACCAAGCCCCATATGCTGCAGAGAATCCCTTCCAAAA<br>CATTGCGCTTGCGGAGGATACCAAGCTGGTACCTGACTTAAAGTACTATTATAAAGAGTATGGTATTGATATTGAAGAATTT<br>GAGGTGGAGACAGACGACGGTTTTATCATTGATTTGTGGCACTTCAAGTCCAGGTTAAACGATGGCGTAGAGGAAGTGAAAC<br>GTGAACCCATTTTGCTATTACACGGTCTTTTGCAAAGTTGCGGGGCCTTCGCATCTTCAGGTAGAAAATCTCTGGCATACTT<br>CCTTTATGAATCTGGATTTGATGTTTGGTTGGGGAATAACAGATGTGGCTTAAATGCCAAGTGGAACATGAAAAAATTAGGT<br>AATGATCATTCCAAAAAATGGGATTGGGATATGCACCAGATGGTTCAATACGATTTAAAAGCACTTATAAATTATGTACTTG<br>ATAGTACCGGATATGCTAAACTATCGCTCGTGGCGCATTCACAAGGTACAACACAAGGGTTTATGGGGCTTGTCAACGGTGA<br>GAAACTATACGCAAGTGATTTTAAATTAGTCGATAAGCTGGAAAACTTTGTTGCTTTGGCACCTGCAGTGTACCCTGGACCT<br>TTATTGGATGAAAAAGCATTTGTGAGGCTAATGGCAAAAGGTATTGATTCTCCGTGGTACTTTGGTAGAAGATCGTTTATAC<br>CTCTTATGATGACAATGAGAAAACTAATGGTTGGGACCAAGATTTTTTCGTTCCTCTCATACATCATGTTTAACTACCTATT<br>TGATTGGAACGATGTCTTATGGGATAGAGTTTTAAGGGACCGAAATTTCTTATTTTCACCAGTCCATATTTCCGTCAAACTA<br>ATGCAATGGTGGTTGTCACCTCTTCCTAATAAGCTAAGTTTTAAGAAGGGTGCAGAGAAAATTTTTCCAGATAAAAAGACAT<br>GGTTTCCTATTGCCAAGAATGATGATGACAGTGGCAATAATTTGGATAACAACAAACTACACTTAAATCCGAAAAGACAAAA<br>TAGTGAAGAATTTCCACACATTATAATGTTTATTCCAAAGCAAGATAGATTAGTAGATGGTGAACGATTAATTAATCACTTC<br>ATTAATCATGAAGCAAATGCAGTGTACAAAATCTGGTATATTGATGAGTATTCCCATCTGGACGTTTTATGGGCGCATGATG<br>TTATAGATAGAATTGGTAAGCCAATGATAGAAAATTTGAGGTTTCCTAATGCAAGGTGA |
| *PpCYP87G1* | ATGTGGAACATAATCTTATGCATAGTAGGGCTAGTAGTAGTAGGAATCACCCATTGGGTGTACAGATGGAGGAATCCAAAAT<br>GCAAAGGAGTATTGCCTCCTGGCTCTATGGGTCTTCCCCTTATTGGAGAAACCCTTCAATACTTCTCCAAGAGTCCCTATGA<br>AGGAATTCCCCCTTTCATTGCTGAAAGAAAAGCAAAGTATGGGACGCTGTTTAAAACGAGCTTAGTTGGACAGCCTATTATA<br>ATATCGACAGATCCAGATGTGAATTACTATGTGTTCCAACAAGAAGATAAGTTATTTCAATGTTCTTACACAAAGAGTGCTG<br>TTGAACTGTCTGGGAAAAAAGGTTTGATGGGTAATGGTGGATCTGCCCATAAATACCTTAGGAACTTAGTGCTATCTCTTAC<br>TGGACCAGATAAGCTAAGGACTAAGTTAGTCTCTGATGTTGATATCATAACTCGTGAACACTTGCATAGGTGGACTACTCAA<br>GGTGATGTTGAAGTCAAAGATGCTTCTGAAATTATGTTGTTCATATTTATTGCGGCAAAGATTCTGGGTATGAATGAAAAAG<br>AGGCATTGACTCTAAGAGGACACTACAAAGCCTTTGTCAAAGGCTTTCTCTCATTTCCTATCAATTTGCCTGGAACAGCTTT<br>TCATTCTGGTTTACAGGGGCGTAAAAGCGCTATAAAGATGATCAAAGACATTTTCGAGAAAAGGAGGTCATCAAAGGATAAA<br>GCAAATGAGCAAGACTTCATAGATCACCTACTTCAAGAAATAGACAAGGAGGACACATTTATAACAGAAGACACAGCAGTGG<br>ATCTAATCTTCTTTGTTATATTTGCTGCCCATGAAACCACTTCTTCAACCATGACACTACTCTTTAAATACTTCACTGAGCA<br>TCCTGATGTTGTTAAAAAACTAAAGGAAGAACATGAAAACATCATTAGAAATCGTGAAGATAAGAACGCTCCAATTTCATGG<br>TCTGAATACAAGTCCATGAACTTCACACATAAGGTTACAAATGAAACAGTAAGACTTGCCAATATCGCCCCAGGAATTTTCA<br>GAAAAGTATTAAAGGATGTGGAAATCAAAGGATATACGATTCCAGAAGGATGGACAATGGTTGTATGTTCCCCATCAGTTCA<br>TCTAGATGAAAACATGTATGACAATCCACTTGCGTTTAACCCATCCCGATGGAAGGATGATGAGCTTGGTGCATCCAAAAAG<br>TTCATGGCGTTCGGAGGAGGAAATAGAATGTGTGTTGGAGCTGATCTCTCAAAGATGCAAACATCCATTTTTATTCACTACT<br>TGATGACAAGTTTTAAGTGGAAGGTTGCAAATAAAGGAAACATTATACGACAACCCTATTTGAACTTCAACGATGGGATATG<br>CATTCGAGTCGATGAGATTCAAAGAGATAACAAAAAGGAAATTAATGTTGCTTGA |
| *PpCYP87G1*$^{opt}$<br>codon-optimised | ATGTGGAATATTATTTTGTGTATTGTTGGTTTGGTTGTTGTTGGTATTACTCATTGGGTTTATAGATGGAGAAATCCAAAAT<br>GTAAAGGTGTTTTGCCACCTGGTTCTATGGGTTTGCCATTGATTGGTGAAACTTTGCAATATTTTTCTAAATCTCCATATGA<br>AGGTATTCCACCATTTATTGCTGAAAGAAAAGCTAAATATGGTACTTTGTTTAAAACTTCTTTGGTTGGGCAACCAATTATT<br>ATTTCTACTGATCCTGATGTAAATTACTATGTGTTCCAACAAGAAGATAAGCTTTTTCAATGTTCTTATACTAAATCTGCTG<br>TTGAATTGTCTGGTAAAAAAGGTTTGATGGGTAATGGTGGTTCTGCTCATAAATATTTGAGAAATTTGGTTTTGTCTTTGAC<br>TGGTCCTGATAAATTGAGAACTAAATTGGTTTCTGATGTTGATATTATTACTAGAGAACATTTGCATAGATGGACTACTCAA<br>GGTGACGTTGAGGTAAAAGATGCTTCTGAAATTATGCTGTTTATTTTCATAGCTGCTAAAATTTTGGGTATGAATGAAAAAG<br>AAGCTTTGACTTTGAGAGGTCATTATAAAGCTTTTGTTAAAGGTTTTTTGTCTTTTCCAATTAATTTGCCTGGTACTGCTTT<br>TCACTCTGGACTTCAAGGGAGGAAATCTGCTATAAAGATGATTAAAGATATTTTCGAAAAAAGACGTTCTTCTAAAGATAAA<br>GCTAACGAGCAAGATTTTATAGACCACTTATTGCAAGAAATTGATAAGGAAGACACTTTCATTACTGAAGATACCGCTGTGG<br>ATTTGATCTTTTTTGTTATTTTTGCTGCTCATGAAACTACTTCTTCTACTATGACATTGTTGTTTAAGTATTTTACCGAACA<br>CCCGGATGTGGTTAAGAAGTTGAAGGAGGAACATGAAAACATTATTAGGAATCGAGAAGATAAAAATGCTCCAATCTCATGG<br>TCTGAATACAAATCTATGAACTTTACGCACAAAGTTACTAATGAAACTGTTAGATTGGCTAATATTGCTCCTGGTATTTTCA<br>GAAAAGTTTTGAAAGACGTTGAAATTAAAGGTTATACTATTCCTGAAGGTTGGACTATGGTTGTTTGTTCTCCATCTGTTCA<br>TTTGGATGAAAATATGTATGATAATCCATTGGCTTTTAATCCATCTAGATGGAAAGATGATGAATTGGGTGCTTCTAAAAAA<br>TTTATGGCTTTTGGTGGTGGTAATAGAATGTGTGTTGGTGCTGATTTGTCTAAAATGCAAACTTCTATTTTCATTCATTACT<br>TGATGACTTCTTTCAAATGGAAGGTTGCTAATAAAGGTAACATAATTAGACAACCATACTTGAATTTCAATGACGGTATTTG<br>CATTAGAGTTGACGAGATCCAAAGAGATAATAAAAAAGAAATTAATGTTGCTTAA |
| *PpCYP88C7*$^{opt}$<br>codon-optimised | ATGGAATATAATTTGGTTTTTTTGTCTACCGCTTTTGCTGTTGGTATTTTGACACTTATTTCAGTATTGAAAAGAGCAAACG<br>GTTGGTTTTATTCTATGAAATTTTCTTCTGAAAAATGTAGATTGCCACCTGGTGATATGGGTTGGCCTGTTGTTGGCAATAT<br>GTTGTTTTTTGTTAAGTGCTTATCTGCTTACGACCTGAAGTCTTTTGTTTCATATTTTGTTACTAGATTTGGTCAAGGTGGT<br>ATGTATAGAACTTTTATGTTTGGTAAACCATCTGTTATTGTTACTACTCCTGAACTGTGTAGGAAAATATTGATGGATGATG<br>AGAATTTCGATTTGGGGTTTCCATCTTATATTTTGGAATTGTTGAGAAAAGAACCAATTGGTGGTACTTCTTATCAAGAAGA<br>TAGATTGTCTAGAAGATTGATGACTCCAATTAAATCTCATGCATTGGTTTCTTACTTTTTTGACTTTTTGTCTGAAACTGTT<br>CAAACTACTTTTGAAAAATGGGCTACTACTGGTGAATCTTTGCAATTGTTGTTTGAAATGAAAAAACCAACTTTTAAAGTTT<br>TGATGAAAGTTTTAATTGGCGGTGATCAAGTTGAAAACAAGCTGTTAGATTCTCTATTTAAAGAGAATAATTTTAGATTTGC<br>TGGTTTGAGATCTTTGCCATTGGATTATCCTGGTTCTACGTATAATAGAGCTATGAAAGGGAGGGGTGAGATTGTCAAAATT<br>TATGAGAGAATCATTAATGAAAGAAAGGTTATGATTGCTAAAACTAGAGGTGAACCAAGAACTAATTTGTTGGATATTATGT<br>TGGATTCCCAATACGACGGTGAAGGTAAGGTTCTGAATGATGAAAATATTATGAAAGTTTTGTTGTGGTACACTTTTTCTGG<br>TTATGAATCTATTGCTAAAGTTGCTACTCAAACTATTATGTTGTTGGAAAAACATCCTGAATGTTTCAAAAAGGCTAAAGAG<br>GAACAAGAAGAGATTGTTAAAAGAAGATCTTCTCCTGATGCTGGTTTGACTTTTTCAGAAATTGGTCAAATGAAATATGTTA<br>ACAATGTCATAAACGAAACTTTGAGATTGGGTTCTACTGAAACAGTTTTGTTTCGAGACGCCCGTACTGATGTGAATATCAA<br>CGGTTATACTATTCCAAAAGGTTGGAAAGTTTTGGCATTGCTAGGTAATTTGTACATGGATCCAAAGACTTATGTTAAACCA<br>AAAGAGTTTAACCCATCTAGATGGGATGATTTTGAAACTAAACCAAATTCTTTTATTCCATTTGGTGTTGGTTTGAGATTGT<br>GTCCTGGTAGCAATCTAGTGAGATTGGAAGTCTCTGTGTTTCTCCACTATTTCTTGCTTAATTATCGACTCGAACAATTGAA<br>TAAAGATTCTAAAGCTGAAGCTTGTATTGCTAGATTTAAAAAAATTTCTGCTTAA |

| | |
|---|---|
| $P_{TEF1}$ promoter | CACACACCATAGCTTCAAAATGTTTCTACTCCTTTTTTACTCTTCCAGATTTTCTCGGACTCCGCGCATCGCCGTACCACTT<br>CAAAACACCCAAGCACAGCATACTAAATTTCCCCTCTTTCTTCCTCTAGGGTGTCGTTAATTACCCGTACTAAAGGTTTGGA<br>AAAGAAAAAAGAGACCGCCTCGTTTCTTTTTCTTCGTCGAAAAAGGCAATAAAAATTTTTATCACGTTTCTTTTTCTTGAAA<br>ATTTTTTTTTTTGATTTTTTTCTCTTTCGATGACCTCCCATTGATATTTAAGTTAATAAACGGTCTTCAATTTCTCAAGTTT<br>CAGTTTCATTTTTCTTGTTCTATTACAACTTTTTTTACTTCTTGCTCATTAGAAAGAAAGCATAGCAATCTAATCTAAG |
| $P_{PGK1}$ promoter | GAAGTACCTTCAAAGAATGGGGTCTTATCTTGTTTTGCAAGTACCACTGAGCAGGATAATAATAGAAATGATAATATACTAT<br>AGTAGAGATAACGTCGATGACTTCCCATACTGTAATTGCTTTTAGTTGTGTATTTTTAGTGTGCAAGTTTCTGTAAATCGAT<br>TAATTTTTTTTTCTTTCCTCTTTTTATTAACCTTAATTTTTATTTTAGATTCCTGACTTCAACTCAAGACGCACAGATATTA<br>TAACATCTGCATAATAGGCATTTGCAAGAATTACTCGTGAGTAAGGAAAGAGTGAGGAACTATCGCATACCTGCATTTAAAG<br>ATGCCGATTTGGGCGCGAATCCTTTATTTTGGCTTCACCCTCATACTATTATCAGGGCCAGAAAAAGGAAGTGTTTCCCTCC<br>TTCTTGAATTGATGTTACCCTCATAAAGCACGTGGCCTCTTATCGAGAAAGAAATTACCGTCGCTCGTGATTTGTTTGCAAA<br>AAGAACAAAACTGAAAAAACCCAGACACGCTCGACTTCCTGTCTTCCTATTGATTGCAGCTTCCAATTTCGTCACACAACAA<br>GGTCCTAGCGACGGCTCACAGGTTTTGTAACAAGCAATCGAAGGTTCTGGAATGGCGGGAAAGGGTTTAGTACCACATGCTA<br>TGATGCCCACTGTGATCTCCAGAGCAAAGTTCGTTCGATCGTACTGTTACTCTCTCTCTTTCAAACAGAATTGTCCGAATCG<br>TGTGACAACAACAGCCTGTTCTCACACACTCTTTTCTTCTAACCAAGGGGGTGGTTTAGTTTAGTAGAACCTCGTGAAACTT<br>ACATTTACATATATATAAACTTGCATAAATTGGTCAATGCAAGAAATACATATTTGGTCTTTTCTAATTCGTAGTTTTTCAA<br>GTTCTTAGATGCTTTCTTTTTCTCTTTTTTACAGATCATCAAGGAAGTAATTATCTACTTTTTACAACAAATATAAAACA |
| $P_{TDH3}$ promoter | ATAAAAAACACGCTTTTTCAGTTCGAGTTTATCATTATCAATACTGCCATTTCAAAGAATACGTAAATAATTAATAGTAGTG<br>ATTTTCCTAACTTTATTTAGTCAAAAAATTAGCCTTTTAATTCTGCTGTAACCCGTACATGCCCAAAATAGGGGGCGGGTTA<br>CACAGAATATATAACATCGTAGGTGTCTGGGTGAACAGTTTATTCCTGGCATCCACTAAATATAATGGAGCCCGCTTTTTAA<br>GCTGGCATCCAGAAAAAAAAAGAATCCCAGCACCAAAATATTGTTTTCTTCACCAACCATCAGTTCATAGGTCCATTCTCTT<br>AGCGCAACTACAGAGAACAGGGGCACAAACAGGCAAAAAACGGGCACAACCTCAATGGAGTGATGCAACCTGCCTGGAGTAA<br>ATGATGACACAAGGCAATTGACCCACGCATGTATCTATCTCATTTTCTTACACCTTCTATTACCTTCTGCTCTCTCTGATTT<br>GGAAAAAGCTGAAAAAAAAGGTTGAAACCAGTTCCCTGAAATTATTCCCCTACTTGACTAATAAGTATATAAAGACGGTAGG<br>TATTGATTGTAATTCTGTAAATCTATTTCTTAAACTTCTTAAATTCTACTTTTATAGTTAGTCTTTTTTTTAGTTTTAAAAC<br>ACCAAGAACTTAGTTTCGAATAAACACACATAAACAAACAAA |
| $P_{FBA1}$ promoter | TGGGTCATTACGTAAATAATGATAGGAATGGGATTCTTCTATTTTTCCTTTTTCCATTCTAGCAGCCGTCGGGAAAACGTGG<br>CATCCTCTCTTTCGGGCTCAATTGGAGTCACGCTGCCGTGAGCATCCTCTCTTTCCATATCTAACAACTGAGCACGTAACCA<br>ATGGAAAAGCATGAGCTTAGCGTTGCTCCAAAAAAGTATTGGATGGTTAATACCATTTGTCTGTTCTCTTCTGACTTTGACT<br>CCTCAAAAAAAAAAAATCTACAATCAACAGATCGCTTCAATTACGCCCTCACAAAAACTTTTTTCCTTCTTCTTCGCCCACG<br>TTAAATTTTATCCCTCATGTTGTCTAACGGATTTCTGCACTTGATTTATTATAAAAAGACAAAGACATAATACTTCTCTATC<br>AATTTCAGTTATTGTTCTTCCTTGCGTTATTCTTCTGTTCTTCTTTTTCTTTTGTCATATATAACCATAACCAAGTAATACA<br>TATTCAAA |
| $P_{GPD1}$ promoter | CACTTCTACTTCTACATCGGAAAAACATTCCATTCACATATCGTCTTTGGCCTATCTTGTTTTGTCCTCGGTAGATCAGGTC<br>AGTACAAACGCAACACGAAAGAACAAAAAAAGAAGAAAACAGAAGGCCAAGACAGGGTCAATGAGACTGTTGTCCTCCTACT<br>GTCCCTATGTCTCTGGCCGATCACGCGCCATTGTCCCTCAGAAACAAATCAAACACCCACACCCCGGGCACCCAAAGTCCCC<br>ACCCACACCACCAATACGTAAACGGGGCGCCCCCTGCAGGCCCTCCTGCGCGCGGCCTCCCGCCTTGCTTCTCTCCCCTTCC<br>TTTTCTTTTTCCAGTTTTCCCTATTTTGTCCCTTTTTCCGCACAACAAGTATCAGAATGGGTTCATCAAATCTATCCAACCT<br>AATTCGCACGTAGACTGGCTTGGTATTGGCAGTTTCGTAGTTATATATATACTACCATGAGTGAAACTGTTACGTTACCTTA<br>AATTCTTTCTCCCTTTAATTTTCTTTTATCTTACTCTCCTACATAAGACATCAAGAAACAATTGTATATTGTACACCCCCCC<br>CCTCCACAAACACAAATATTGATAATATAAAG |
| $P_{GAL1}$ promoter | ACGGATTAGAAGCCGCCGAGCGGGTGACAGCCCTCCGAAGGAAGACTCTCCTCCGTGCGTCCTCGTCTTCACCGGTCGCGTT<br>CCTGAAACGCAGATGTGCCTCGCGCCGCACTGCTCCGAACAATAAAGATTCTACAATACTAGCTTTTATGGTTATGAAGAGG<br>AAAAATTGGCAGTAACCTGGCCCCACAAACCTTCAAATGAACGAATCAAATTAACAACCATAGGATGATAATGCGATTAGTT<br>TTTTAGCCTTATTTCTGGGGTAATTAATCAGCGAAGCGATGATTTTTGATCTATTAACAGATATATAAATGCAAAAACTGCA<br>TAACCACTTTAACTAATACTTTCAACATTTTCGGTTTGTATTACTTCTTATTCAAATGTAATAAAAGTATCAACAAAAAATT<br>GTTAATATACCTCTATACTTTAACGTCAAGGAGAAAAAAC |
| $P_{ADH2}$ promoter | TATCTAAAAATTGCCTTATGATCCGTCTCTCCGGTTACAGCCTGTGTAACTGATTAATCCTGCCTTTCTAATCACCATTCTA<br>ATGTTTTAATTAAGGGATTTTGTCTTCATTAACGGCTTTCGCTCATAAAAATGTTATGACGTTTTGCCCGCAGGCGGGAAAC<br>CATCCACTTCACGAGACTGATCTCCTCTGCCGGAACACCGGGCATCTCCAACTTATAAGTTGGAGAAATAAGAGAATTTCAG<br>ATTGAGAGAATGAAAAAAAAAAAAAAAAAAAAAGGCAGAGGAGAGCATAGAAATGGGGTTCACTTTTTGGTAAAGCTATAGCA<br>TGCCTATCACATATAAATAGAGTGCCAGTAGCGACTTTTTTCACACTCGAAATACTCTTACTACTGCTCTCTTGTTGTTTTT<br>ATCACTTCTTGTTTCTTCTTGGTAAATAGAATATCAAGCTACAAAAAGCATACAATCAACTATCAACTATTAACTATATCGT<br>AATACACA |

**Supplementary Table 13. List of guide RNA sequences for gene deletions in yeast.**

| **Target gene name** | **gRNA sequence 5' – 3'** |
|---|---|
| *ERG4* (*YGL012W*) | GGCTTCAGCCATTTACCCTG |
| *ERG5* (*YMR015C*) | GTGTAGCGTAGAATTATGAG |
| *Pper24ISO* | TTCGTCCCAAGAGGGAGACG |

**Supplementary Table 14. List of primers for metabolic engineering and transient expression in *N. benthamiana*.**

Overhangs for cloning are bold and underlined.

| Name | Sequence | Purpose |
|---|---|---|
| AtCAS-F | **CACCACAGGTCTCGAAAA**ATGTGGAAACTGAAGATCGCGG | Cloning *AtCAS* into pHREAC |
| AtCAS-R | **CACCACAGGTCTCGAGCGT**CATTCTCCTTGTTGCAATAATACCT | Cloning *AtCAS* into pHREAC |
| AtSMT1-Fr1-F | **TTGGTCTCAAAAA**ATGGATCTCGCGTCGAATCTTG | Cloning *AtSMT1* into pHREAC |
| AtSMT1-Fr1-R | **TTGGTCTCTGCGT**TGGCTTCCTTGAGGTGCAAG | Cloning *AtSMT1* into pHREAC |
| AtSMT1-Fr2-F | **TTGGTCTCAACGC**GTCTCAAATTTCCTGGAGCAGGC | Cloning *AtSMT1* into pHREAC |
| AtSMT1-Fr2-R | **TTGGTCTCTAGCGT**TACTCTGGCTTCCGGGCC | Cloning *AtSMT1* into pHREAC |
| AtSMO1-1-F | **CACCACAGGTCTCGAAAA**ATGATTCCTTACGCTACAGTCGAAG | Cloning *AtSMO1-1* into pHREAC |
| AtSMO1-1-R | **CACCACAGGTCTCGAGCG**CTAATCGGATTTTATTCCTCCGTTATGC | Cloning *AtSMO1-1* into pHREAC |
| At3βHSD1-F | **CACCACAGGTCTCGAAAA**ATGGTGATGGAAGTTACAGAGACTG | Cloning *At3βHSD1* into pHREAC |
| At3βHSD1-R | **CACCACAGGTCTCGAGCG**TTAGTCGATCTTCTTGCTCCCG | Cloning *At3βHSD1* into pHREAC |
| AtERG28-F | **CACCACAGGTCTCGAAAA**ATGAAGGCGTTGGGGTATTGGT | Cloning *AtERG28* into pHREAC |
| AtERG28-R | **CACCACAGGTCTCGAGCG**TCAAGAAAGTTTGGAGTGTGGTTGT | Cloning *AtERG28* into pHREAC |
| AtCPI-F | **CACCACAGGTCTCGAAAA**ATGTCAGGATCTTCTTCACCGAG | Cloning *AtCPI* into pHREAC |
| AtCPI-R | **CACCACAGGTCTCGAGCG**TCAGCAGTTGGAGAAGAAAAGC | Cloning *AtCPI* into pHREAC |
| AtCYP51-F | **CACCACAGGTCTCGAAAA**ATGGAATTGGATTCGGAGAACAAAT | Cloning *AtCYP51* into pHREAC |
| AtCYP51-R | **CACCACAGGTCTCGAGCG**TTAAGAAAGCTGGCGCCTCTT | Cloning *AtCYP51* into pHREAC |
| AtC14R-F | **CACCACAGGTCTCGAAAA**ATGCTGCTAGATATGGATCTCGG | Cloning *AtC14R* into pHREAC |
| AtC14R-R | **CACCACAGGTCTCGAGCG**CTAATAAACATAAGGAAGTATTCTCCAGG | Cloning *AtC14R* into pHREAC |
| At8,7SI-Fr1-F | **TTGGTCTCAAAAA**ATGGAGGAGTTGGCGCATCC | Cloning *At8,7SI* into pHREAC |
| At8,7SI-Fr1-R | **TTGGTCTCTCAGA**CCAGTGAATGTCCACCAACACATAAG | Cloning *At8,7SI* into pHREAC |
| At8,7SI-Fr2-F | **TTGGTCTCATCTG**ACTCATGTTATTCTCGAGGGCTATTTC | Cloning *At8,7SI* into pHREAC |
| At8,7SI-Fr2-R | **TTGGTCTCTAGCG**TCAACGGGTTTTCTTCTTCGTCTTTG | Cloning *At8,7SI* into pHREAC |
| AtSMO2-1-F | **CACCACAGGTCTCGAAAA**ATGGCTTCCTTCGTGGAATCTG | Cloning *AtSMO2-1* into pHREAC |
| AtSMO2-1-R | **CACCACAGGTCTCGAGCG**TCACGTTTGTTTCATGTCACCG | Cloning *AtSMO2-1* into pHREAC |
| AtSMO2-2-F | **CACCACAGGTCTCGAAAA**ATGGATTCTCTCGTTGAATCCGG | Cloning *AtSMO2-2* into pHREAC |
| AtSMO2-2-R | **CACCACAGGTCTCGAGCG**TCAGGTTTCTTTTAGGGCCTTAAGT | Cloning *AtSMO2-2* into pHREAC |
| AtDWARF7-F | **CACCACAGGTCTCGAAAA**ATGGCGGCGGATAATGCTTAT | Cloning *AtDWARF7* into pHREAC |
| AtDWARF7-R | **CACCACAGGTCTCGAGCG**TCACTCTGCTTTCTTGAAGCTGT | Cloning *AtDWARF7* into pHREAC |
| AtDWARF5-F | **CACCACAGGTCTCGAAAA**ATGGCGGAGACTGTACATTCTC | Cloning *AtDWARF5* into pHREAC |
| AtDWARF5-R | **CACCACAGGTCTCGAGCG**TCAATAAATTCCCGGAATGATCCTG | Cloning *AtDWARF5* into pHREAC |
| pHREAC_F | CTGTCACTTTATTGAGAAGATAGTGG | Colony PCR and sequencing |
| pHREAC_R | CCTTGCTGAAGGGACGACCTG | Colony PCR and sequencing |
| Pp24ISO-F | **CAATCTACATTATATTAAACGTCTCTAAAA**ATGTCAGGGGAGAAGGTGGC | Amplification of *Physalis peruviana 24ISO* for In-Fusion cloning into pHREAC |
| Pp24ISO-R | **AACTAAAGAAAATTTAATGAAACCAGAGCG**TCAATCCGCGGGCTCATC | Amplification of *Physalis peruviana 24ISO* for In-Fusion cloning into |

| | | |
|---|---|---|
| | | pHREAC |
| CYP87G1-F | **TTGGTCTCAAAA**AATGTGGAACATAATCTTATGCATAGTA | Cloning *PpCYP87G1* into pHREAC |
| CYP87G1-R | **TTGGTCTCTAGCG**TCAAGCAACATTAATTTCCTTTTTGTT | Cloning *PpCYP87G1* into pHREAC |
| CYP87G1-SeqR | CCAGATGCATAAGTATCGTAAC | Sequencing |
| CYP88C7-F | **TTGGTCTCAAAA**AATGGAGTACAACTTAGTGTTCTTGTCC | Cloning *PpCYP88C7* into pHREAC |
| CYP88C7-R | **TTGGTCTCTAGCG**TTATGCCGAGATCTTCTTGAACCTCGC | Cloning *PpCYP88C7* into pHREAC |
| CYP88C7-SeqR | GTCTGTGTGGCAACCTTAGC | Sequencing |
| CYP749B2-F | **TTGGTCTCAAAA**AATGATGATAGCACTAATAGCTCTCTCT | Cloning *CYP749B2* into pHREAC |
| CYP749B2-R | **TTGGTCTCTAGCG**TTACAGTGGATTAAGAATGATTTGAAT | Cloning *CYP749B2* into pHREAC |
| CYP749B2-SeqR | GCAGGCTCCAGCTTAGCAAAC | Sequencing |
| DWF1silfwd_040918 0.5 kb | **TTTTGAATTC**AATGGTTGCAAGAGTTGA | Cloning a fragment of *NbDWF1* into pTRV2 |
| DWF1silrev_040918 0.5 kb | **TTTTCTCGAGC**GAAGCATATCTACCGG | Cloning a fragment of *NbDWF1* into pTRV2 |
| PDSsilfwd_040918 0.6 kb | **TTTTGAATTC**TCAAATTTGCTATTGGACTC | Cloning a fragment of *NbPDS* into pTRV2 |
| PDSsilrev_040918 0.6 kb | **TTTTCTCGAGC**CCACTAGCTTCTCCA | Cloning a fragment of *NbPDS* into pTRV2 |

**Supplementary Table 15. Purification of ergosta-5,25-diene-3β,24ξ-diol (7) heterologously produced in *N. benthamiana*.**

| | | | | | |
|---|---|---|---|---|---|
| Ergosta-5,25-diene-3β,24ξ-diol (**7**) | **Extraction Solvent:** | | Hexane | **Plant Dry Weight** | 78 g |
| | **Volume:** | | 1.8 L | **Crude extract:** | 1,080 mg |
| | **Instrument** | **Column** | **Solvents** | **Gradient** | **Yield** |
| | Biotage | SNAP KP-Sil 25g | A: Petroleum ether<br>B: Ethyl acetate | 0-25% B (10 CV)<br>25-30% B (3 CV)<br>0% B (8 CV)<br>30-100% B (3 CV)<br>100% B (5 CV) | 9 mg |
| | LC-MS | Kinetex 5 µm C18 100 Å 250 x 10 mm | A: $H_2O/NH_4OAc$<br>B: Methanol/$NH_4OAc$<br>Flowrate: 5 mL/min | 85% B (4 min)<br>85-90% B (10 min)<br>100% B (2 min)<br>85% B (2 min) | 2 mg |

**Supplementary Table 16. Purification of ergosta-5,24(28)-diene-3β,25-diol (8) heterologously produced in *N. benthamiana*.**

| | | | | | |
|---|---|---|---|---|---|
| Ergosta-5,24(28)-diene-3β,25-diol **(8)** | **Extraction Solvent:** | | Hexane | **Plant Dry Weight** | 78 g |
| | **Volume:** | | 1.8 L | **Crude extract:** | 1,080 mg |
| | **Instrument** | **Column** | **Solvents** | **Gradient** | **Yield** |
| | Biotage | SNAP KP-Sil 25g | A: Petroleum ether<br>B: Ethyl acetate | 0-25% B (10 CV)<br>25-30% B (3 CV)<br>30% B (8 CV)<br>30-100% B (3 CV)<br>100% B (5 CV) | 34 mg |
| | LC-MS | Kinetex 5 µm C18 100 Å 250 x 10 mm | A: $H_2O/NH_4OAc$<br>B: Methanol/$NH_4OAc$<br>Flowrate: 4.5 mL/min | 85% B (3 min)<br>85-90% B (10 min)<br>100% B (3 min)<br>85% B (3 min) | 2 mg |

**Supplementary Table 17. Purification of (22*R*)-ergosta-5,24-diene-3β,22-diol (10) heterologously produced in *N. benthamiana*.**

| | | | | | |
|---|---|---|---|---|---|
| (22*R*)-Ergosta-5,24-diene-3β,22-diol **(10)** | **Extraction Solvent:** | | Hexane | **Plant Dry Weight** | 78 g |
| | **Volume:** | | 1.8 L | **Crude extract:** | 1,080 mg |
| | **Instrument** | **Column** | **Solvents** | **Gradient** | **Yield** |
| | Biotage | SNAP KP-Sil 25g | A: Petroleum ether<br>B: Ethyl acetate | 0-25% B (10 CV)<br>25-30% B (3 CV)<br>30% B (8 CV)<br>30-100% B (3 CV)<br>100% B (5 CV) | 16 mg |
| | LC-MS | Luna 5 µm C8 100 Å 250 x 10 mm | A: $H_2O/NH_4OAc$<br>B: Methanol/$NH_4OAc$<br>Flowrate: 5 mL/min | 85-90% B (8 min)<br>90%-100% B (3 min)<br>100% B (2 min)<br>85% B (2 min) | 3 mg |

**Supplementary Table 18. Purification of (22*R*)-ergosta-5,24(28)-diene-3β,22,25-triol (phyministerol A) (12) heterologously produced in *N. benthamiana*.**

| | | | | | |
|---|---|---|---|---|---|
| (22*R*)-Ergosta-5,24(28)-diene-3β,22,25-triol (phyministerol A) (**12**) | **Extraction Solvent:** | | Ethyl acetate | **Plant dry weight:** | 45 g |
| | **Volume:** | | 1.2 L | **Crude extract:** | 1,847 mg |
| | **Instrument** | **Column** | **Solvents** | **Gradient** | **Yield** |
| | Biotage | Sfaer Silica HC D 10g<br>Sfaer Silica HC D 5g (2x) | A: Petroleum ether<br>B: Ethyl acetate | 0-30% B (10 CV)<br>30% B (10 CV)<br>30-100% B (3 CV)<br>100% B (5 CV) | 33 mg |
| | Biotage | Sfaer Silica HC D 5g | A: Petroleum ether<br>B: Ethyl acetate | 70% B (12 CV)<br>70-100% B (2 CV)<br>100% B (3 CV) | 14 mg |
| | LC-MS | Luna 5 µm C8 100 Å 250 x 10 mm | A: $H_2O/NH_4OAc$<br>B: Methanol/$NH_4OAc$<br>Flowrate: 5 mL/min | 85-100% B (10 min)<br>100% B (3 min)<br>100-85% B (0.1 min)<br>85% B (2 min) | 2 mg |

**Supplementary Table 19. Purification of (22*R*)-24,25-epoxy-ergost-5-ene-3β,22-diol (13) heterologously produced in *N. benthamiana*.**

| | | | | | |
|---|---|---|---|---|---|
| | **Extraction Solvent:** | | Hexane | **Plant Dry Weight** | 78 g |
| | | | 1.8 L | **Crude extract:** | 1,080 mg |
| | **Instrument** | **Column** | **Solvents** | **Gradient** | **Yield** |
| (22*R*)-24,25-Epoxy-ergost-5-ene-3β,22-diol (**13**) | Biotage | SNAP KP-Sil 25g | A: Petroleum ether<br>B: Ethyl acetate | 0-25% B (10 CV)<br>25-30% B (3 CV)<br>30% B (8 CV)<br>30-100% B (3 CV)<br>100% B (5 CV) | 17 mg |
| | LC-MS | Kinetex 5 µm C18 100 Å<br>250 x 10 mm | A: $H_2O/NH_4OAc$<br>B: Methanol/$NH_4OAc$<br>Flowrate: 5 mL/min | 85% B (4 min)<br>85-90% B (10 min)<br>100% B (2 min)<br>85% B (2 min) | 1 mg |

**Supplementary Table 20. Purification of (22*R*)-ergosta-5,24(28)-diene-1α,3β,22,25-tetraol (15) heterologously produced in *N. benthamiana*.**

FA: Formic acid

| | | | | | |
|---|---|---|---|---|---|
| (22*R*)-Ergosta-5,24(28)-diene-1α,3β,22,25-tetraol (**15**) | **Extraction Solvent:** | | Ethyl acetate | **Plant dry weight:** | 78 g |
| | **Volume:** | | 1 L | **Crude extract:** | 4,453 mg |
| | **Instrument** | **Column** | **Solvents** | **Gradient** | **Yield** |
| | Biotage | SNAP KP Sil 100g | A: Petroleum ether<br>B: Ethyl acetate | 0-100% B (11 CV)<br>100% B (3 CV) | 206 mg |
| | Biotage | Sfaer C18 D 12g | A: Water<br>B: Acetonitrile | 10-90% B (15 CV)<br>90-100% B (1 CV)<br>100% B (5 CV) | 10 mg |
| | LC-MS | Kinetex 5 µm C18 100 Å 250 x 10 mm | A: $H_2O$/FA<br>B: Acetonitrile/FA<br>Flowrate: 5 mL/min | 40-78.6% B (10 min)<br>78.6-90% B (1 min)<br>90% B (2 min)<br>90-40% B (0.1 min)<br>40% B (2 min) | 1 mg |

**Supplementary Table 21. X-ray crystallographic data of (22*S*)-6*β*-methoxy-3*α*,5-cycloergosta-24,25-diene-26,22-lactone (S6-(22*S*)).**

| | |
|---|---|
| Identification code | MB293twin_twin1_hklf4 |
| ccdc deposition number | 2369097 |
| Empirical formula | $C_{29}H_{44}O_3$ |
| Formula weight | 440.64 |
| Temperature/K | 100.00(10) |
| Crystal system | monoclinic |
| Space group | $P2_1$ |
| a/Å | 7.57280(10) |
| b/Å | 20.7677(5) |
| c/Å | 15.8495(4) |
| α/° | 90 |
| β/° | 93.261(2) |
| γ/° | 90 |
| Volume/Å$^3$ | 2488.61(9) |
| Z | 4 |
| $\rho_{calc}$g/cm$^3$ | 1.176 |
| μ/mm$^{-1}$ | 0.570 |
| F(000) | 968.0 |
| Crystal size/mm$^3$ | 0.466 × 0.362 × 0.08 |
| Radiation | Cu Kα (λ = 1.54184) |
| 2Θ range for data collection/° | 5.584 to 159.238 |
| Index ranges | $-9 \le h \le 9$, $-25 \le k \le 25$, $-20 \le l \le 20$ |
| Reflections collected | 14886 |
| Independent reflections | 14886 [$R_{int}$ = 0.048, $R_{sigma}$ = 0.0209] |
| Data/restraints/parameters | 14886/1/590 |
| Goodness-of-fit on $F^2$ | 1.109 |
| Final R indexes [I>=2σ (I)] | $R_1$ = 0.0452, w$R_2$ = 0.1238 |
| Final R indexes [all data] | $R_1$ = 0.0483, w$R_2$ = 0.1262 |
| Largest diff. peak/hole / e Å$^{-3}$ | 0.20/-0.26 |
| Flack parameter | -0.11(14) |
